# Non-Carbohydrate Inhibitors of Sialic Acid-binding Immunomodulatory-type Lectin-7 (Siglec-7) Discovered from Genetically Encoded Bicyclic Peptide Libraries

**DOI:** 10.1101/2025.10.10.681534

**Authors:** Danial Yazdan, A. Michael Downey, Ana Gimeno, Caishun Li, Edward N. Schmidt, Jeffrey Y. K. Wong, Caleb Loo, Jaesoo Jung, Ryan Qiu, Ewa Lis, Lily Lindmeier, June Ereño-Orbea, Jesús Ji-ménez-Barbero, Matthew S. Macauley, Ratmir Derda

## Abstract

Glycan-binding proteins (GBP) are among the most difficult to drug targets. This deficiency delays clinical progress for therapeutically important GBPs. We employed bicyclic genetically encoded libraries (BiGELs), produced by chemical modification of phage-displayed libraries of peptides with two-fold symmetric linchpins, to discover inhibitors of therapeutically relevant Siglec-7:GD3 interactions. Next-generation sequencing (NGS) analysis of panning of BiGEL against Siglec-7 yielded 815 candidates from which 23 hits yielded *K*_D_ = 1−100 µM as determined by surface plasmon resonance (SPR). Competitive enzyme-linked immunosorbent assays (ELISA) identified a subset of leads that disrupted the Siglec-7:GD3 interaction with *IC*_50_= 3−300 µM. Machine learning models trained on NGS datasets identified additional inhibitors with equivalent potency. Alanine scans of **8c** (SWCRPATVNC, *IC*_50_ = 3.8 µM) and **12c** (SFCHYPTHVC, *IC*_50_= 11 µM), identified key residues as crucial for activity. Ring reshaping studies of compound **8c** highlighted the critical role of bicyclic topology produced by analogue **46e** (SAAAAAWCRPATVNC, *IC*_50_= 9.5 µM). Multivalent display of the lead bicycles alongside ∼100 glycans in Liquid glycan Array (LiGA), made it possible to compare the binding of bicycles and glycans to Siglec-7 expressed on CHO, Jurkat, and Raji cells. LiGA assays confirmed binding of the bicycles to Siglec-7 but revealed considerable non-specific interactions with receptor-negative cells. Saturation transfer difference nuclear magnetic resonance (STD-NMR) revealed **46e** binds to Siglec-7 at a site distinct from the V-Ig domain, suggesting it might inhibit binding of glycans to the glycan-binding site of Siglec-7 via an allosteric site. Together these results demonstrate that BiGEL enables the discovery of bicyclic peptides for undruggable Siglec targets but highlights future challenges in molecular discoveries that aim to identify small, non-carbohydrate inhibitors of GBPs.

## Introduction

Altered cell-surface glycosylation allows cancer cells to escape immune responce.^1, 2^ Sialic acid-binding immunodulatory-type lectins (Siglecs) are sialic acid-binding receptors expressed on the surface of immune cells and interactions. Sialic acid-containing ligands can dampen the immune response of tumour-infiltrating immune cells.^3^ Hypersialylation, a hallmark of many cancers, results in interaction with myeloid cell surface Siglecs that facilitates evasion of the immune response.^4-7^ Given the prominence of hypersialylation in cancer and reported roles of Siglecs in CD8^+^ T cells,^8^ NK cells,^9, 10^ and macrophages,^11^ elimination of the dampening effects emanating from Siglec-sialic acid interaction is a viable therapeutic avenue for repriming the immune cells in the tumour microenvironments. Bertozzi and coworkers^12^ proposed targeted removal of sialic acids by targeted sialidases as one of the strategies for ablation of undesired Siglec-sialic acid interactions. Classical antagonism of specific Siglecs by anti-Siglec antibodies is another strategy. Indeed anti-Siglec-7 or anti-Siglec-9 antibodies have been shown to augment immunity toward carcinomas through suppression of Siglec-7/9 activity.^13-15^ Carbohydrate-binding pockets of GBPs often exhibit only a modest affinity for glycan substrates,^16^ but overall avidity of glycan:GBP is augmented by multivalent interactions.^17^ Multivalent interactions between GBP and native ligands exacerbate the complexity of targeting GBPs with non-antibody modalities (e.g., small molecules, peptides, aptamers).^18^ To explore therapeutic and targeting opportunities complimentary to biologics, we performed a comprehensive discovery campaign against Siglec-7 using small-footprint peptide macrocycles.

Glycan-derived FDA-approved drugs and FDA-approved targeting modalities are known^19^. Due to the susceptibility of glycosidic linkages to degradation and the weak affinity of glycans, a rebuilding of the glycan core is often necessary^19^. Multi-step rebuilding of the oligosaccharide cores to yield clinical candidates for Galectins and Selectins by GalectoBiotech^20^ and GlycoMimetics^21^ provide excellent examples of medicinal chemistry efforts needed for such drug design. When compared to small-molecules targeting non-GBP targets, relatively few GBP-targeting glycomimetic hits progressed though clinical trials; some very promising candidates were withdrawn. Siglec-binding molecules combining a glycan core with small molecule fragments have been pioneered by Paulson and coworkers (Figure 1).^3, 13, 22-26^ Synthetic inhibitors of Siglec-7 based on the sialyllactose core show millimolar to low-micromolar affinity,^22^ and for preclinical applications, the activity of these molecules is often boosted by multivalent display on liposomes, nanoparticles, or proteins. To date, these glycan-derived inhibitors represent the most robust modality for selective targeting of Siglec proteins. As in the case of other glycomimetic modalities, optimization of glycan-derived Siglec inhibitors requires significant medicinal chemistry efforts. We recognized that the availability of a peptide-based scaffold of even comparable affinity may offer a significantly simplified path to optimization. Libraries of peptides and peptide macrocycles previously served as source of leads for GBPs.^27^ There are more than 60 FDA-approved peptide-based therapeutics and >140 more in clinical trials.^28^ However, as therapeutics, peptides also come with their shortcomings, including poor bioavailability and susceptibility to protease degradation.^29^ Strategies to combat these deficiencies include utilization of macrocyclic peptides with condensed secondary structure to reduce surface area and exposure to augment permeability and hinder degradation.^30, 31^

**Figure 1.**
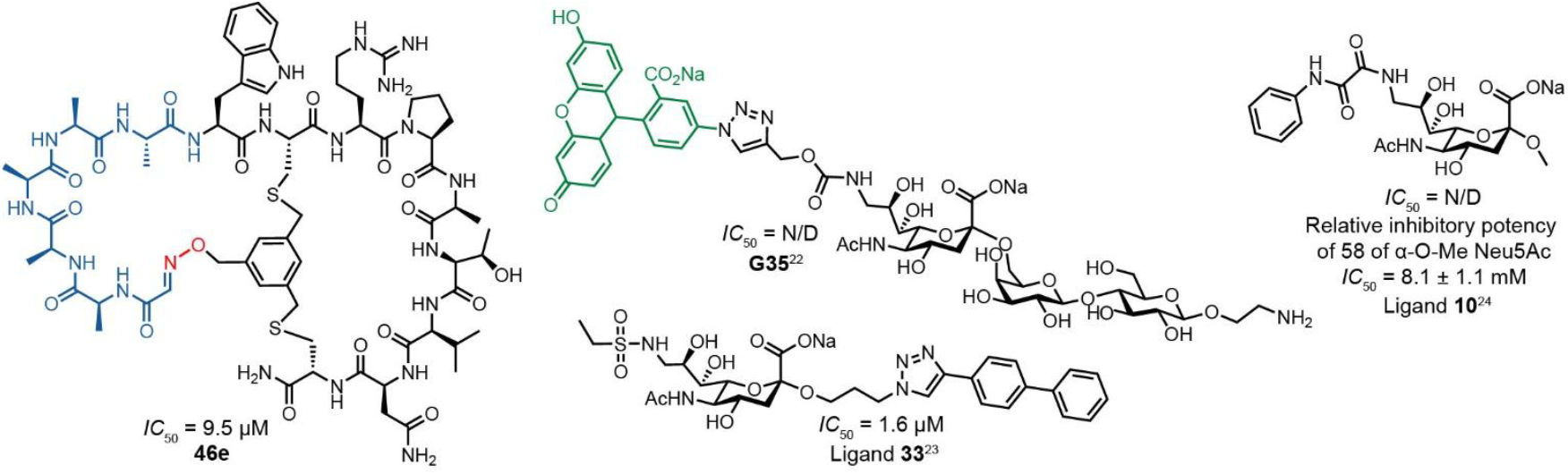
Synthetic inhibitors of Siglec-7. Our peptide-based inhibitor (**46e**) against glycan-derived inhibitors from the Paulson (**G35**)^22^ and Brossmer (Ligand **10** and Ligand **33**) groups.^23, 24^

Phage display is a powerful tool in molecular discovery,^32^ and late-stage functionalization of phage-displayed peptides in aqueous media converts readily available million-to-billion scale genetically encoded chemical space to incorporate unnatural chemotypes and pharmacophores not present in the original peptide libraries.^33-46^ Such an expanded chemical space can drive the discovery of macrocyclic peptide-based inhibitors and may offer leads in the development of Siglec inhibitors. Towards this goal, we elected to use bicyclic scaffolds with masked N-termini, due to their inherent stability over conventional macrocycles^44^ to identify bicyclic peptides that bind to Siglec-7.

## Results and Discussion

### Siglec-7 panning campaign

We employed previously reported bicyclic genetically encoded libraries based on twofold symmetric linkers (BiGEL2)^44^, and purified Fc^47^ fusions of ectodomains of Siglec-7 and Siglec-9 as baits for panning. BiGEL2s were used in four rounds of panning to Siglec-7, with next-generation sequencing (NGS) of each round. Alternated panning against Siglec-7-Fc^47^ and Siglec-7-biotin immobilized on protein G and streptavidin beads steered selection away from peptides that bind to Fc, protein G or streptavidin (Figure 2a). Panning depleted modified phage library against blank support and then immobilized Siglec-7, followed by washing, elution, and amplification to subsequent rounds (Figure 2a). A differential enrichment (DE) analysis of NGS data, conducted as described previously, identified 535 macrocyclic sequences for downstream evaluation. These peptides, plus their alanine-scanned variants (∼8000 total) were re-synthesized using semiconductor-based electrochemical oligo synthesis (Genescript) to yield a synthetic oligo pool, which was used to clone a focused phage-displayed library (Figure 2b). Round 3 enrichment (Figure 2c) and panning the focused library against Siglec-7-Fc or Siglec-7-biotin resulted in nomination of ∼30 candidates for synthesis based on their combined enrichment in the focused library against Siglec-7-Fc and Siglec-7-biotin (Figure S9). A similar multi-round panning campaign on Siglec-9-Fc/Siglec-9-biotin targets nominated 100 sequences for the focused library construction and panning which were narrowed down to 8 + 12 = 20 sequences for synthesis and validation (Figure S8). From the pooled nominees, 25 were synthesized as TSL6-bicycles and the outstanding 12 were deprioritized due to low yields in synthesis or poor solubility.

**Figure 2.**
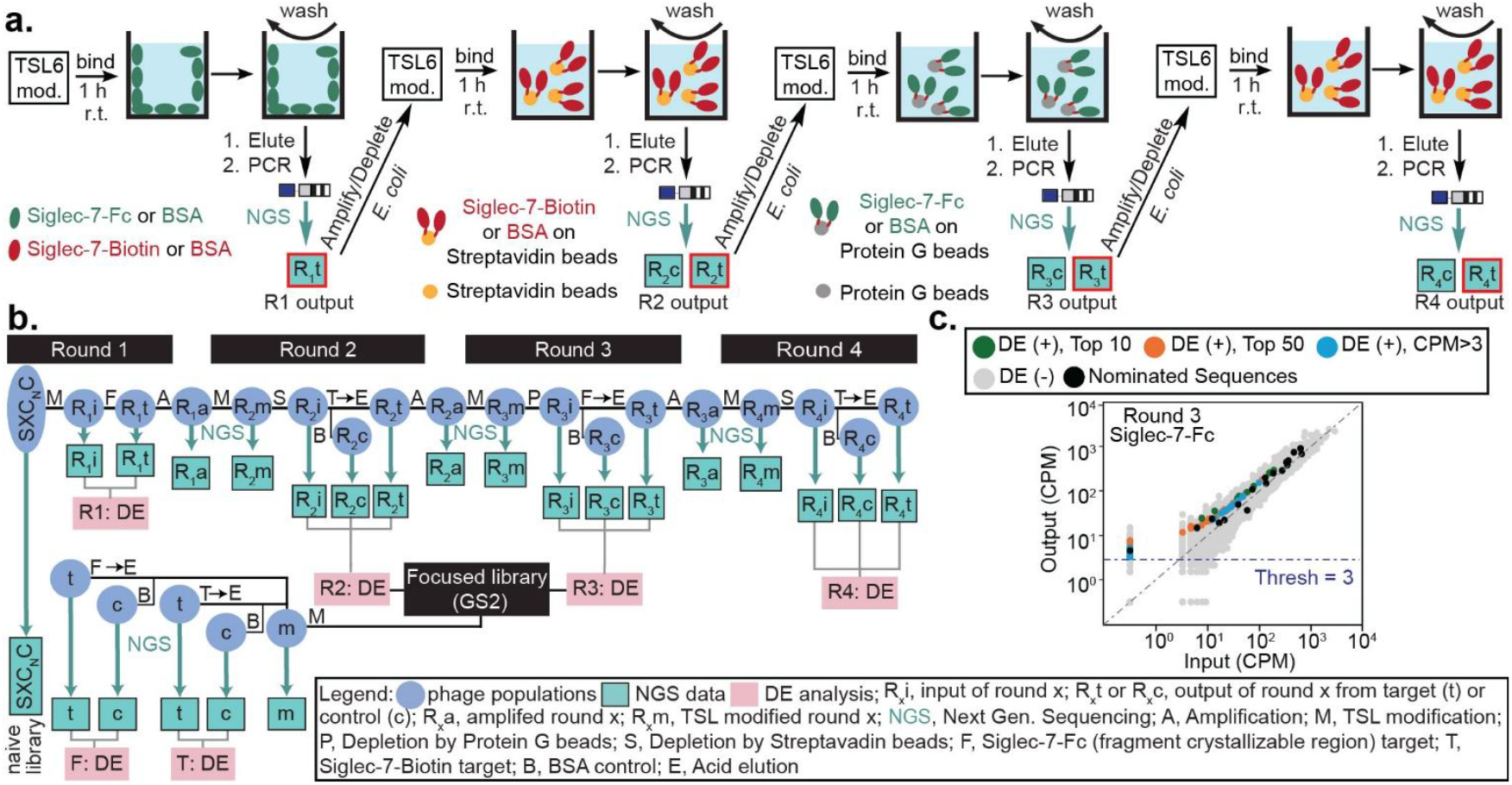
Multi-round panning campaign against Siglec-7. a) Panning procedure for each round alternating between Siglec-7-Fc and Siglec-7-biotin. b) A flow chart of the panning campaign against Siglec-7. c) DE analysis of Round 3 panning against Siglec-7-Fc.

### Analysis of interaction with Siglec-7

Synthesis of the 25 TSL6-bicycles and analysis of their interaction with Siglec-7-Fc by SPR identified **3c, 8c**, and **12c** as promising leads with *K*_D_ = 1.5−3.0 µM (Figure 3a). A subset of 14 TSL6-bicycles were re-tested using a competitive ELISA to determine the ability of bicycles to compete with binding of Siglec-7 to sialoside GD3. From 14 bicycles, 11 were deprioritized due to poor/inconclusive SPR data, poor solubility, or lack of detectable inhibition. **3c** showed affinity *K*_D_ = 2−3 µM as determined by SPR but no detectable inhibition of GD3:Siglec-7 binding at any concentration was observed (Figure 3b). A trend between SPR affinity and FC for the target, and lack of such trend for FC on BSA coated beads was encouraging. The three sequences that broke the trend and exhibited a higher than population average enrichment on BSA coincidentally were the same sequences that exhibited strong SPR affinity (**3c, 22c**, and **18c**) but weak inhibition (*IC*_50_ >100 µM) in the ELISA (Figure 3b). However, a relatively low number of the compounds in this category (*n* = 3) precluded evaluation of the significance of this correlation. Based on the SPR and ELISA data, we focused on the downstream evaluation of the two lead sequences **8c** and **12c** which exhibited SPR and ELISA responses in the low micromolar range (Figure 3a).

**Figure 3.**
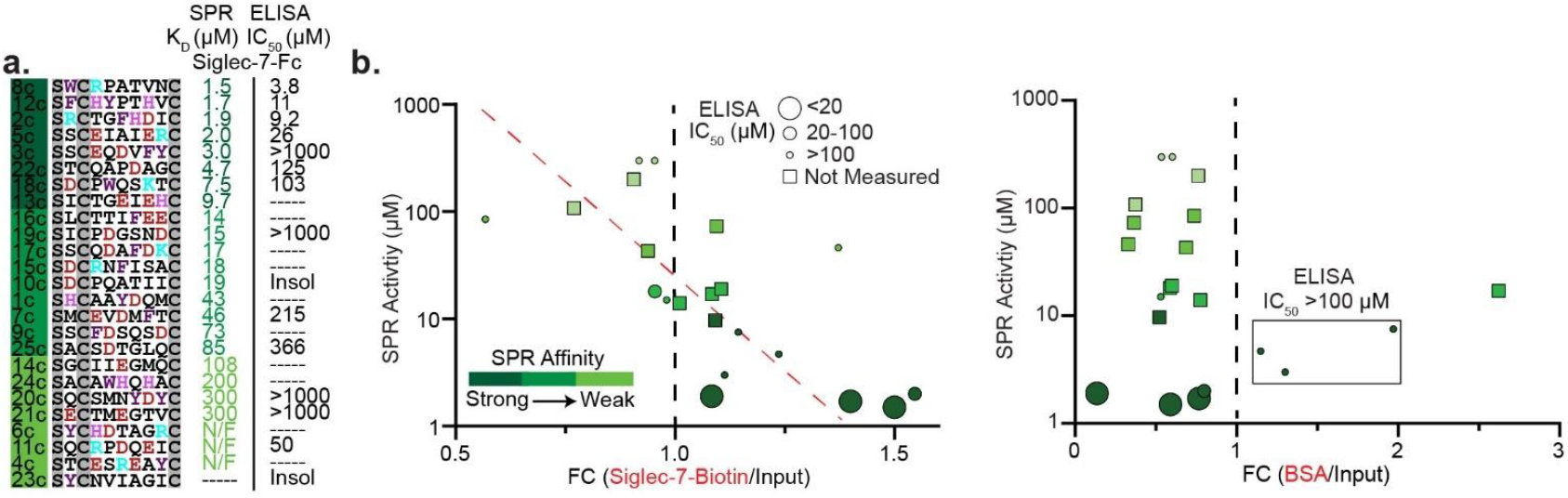
The 25 nominated sequences and their biochemical interactions with Siglec-7. a) SPR and ELISA evaluation of binding to Siglec-7-Fc and inhibition of the GD3:Siglec-7 interaction measured by ELISA, ordered in terms of SPR affinity and colour coded to highlight charged or aromatic residues. b) Trends in SPR data vs. the enrichment of the sequence in a focused library panned against Siglec-7-Biotin or BSA coated streptavidin beads. A downward slope highlights that SPR affinity improves with higher FC (fold change) for Siglec-7 but the three sequences with high FC towards BSA, corresponding to the three sequences that show >100 µM affinity in ELISA, despite their strong SPR affinity.

We synthesized the alanine-scan variants of **8c** and **12c** to build a preliminary SAR profile as well as a linker scan to explore effect on geometry (Figure 4a). The Ala-scan identified the five underlined amino acids in **8c** (S<u>W</u>C<u>R</u>PA<u>T</u>V<u>N</u>C) as important for binding, and the R4A mutant was completely inactive. The Alascan of **12c** identified His 4 (underlined SFC<u>H</u>YPTHVC) as the sole critical residue; interestingly, all other Ala mutants had either no loss of activity or a modest gain of activity, and a triple mutant SFCH<u>YPT</u>HVC → SFCH<u>AAA</u>HVC exhibited only a two-fold loss in activity (**40c**, Figure 4a). As in our previous reports,^46^ we appended the C-terminal triglycine linker (GGG) with propargyl-glycine (Z) to introduce a chemically orthogonal handle for conjugation in downstream testing. We observed that **12c** tolerated C-terminal GGGZ linkers analogous to the Gly-rich linkers in the phage-displayed library. Similar tolerance to C-terminal modification was observed for compound **5c** (Figure S3). Unexpectedly, the same C-terminal GGGZ linker led to a tenfold decrease in potency of **8c** (*IC*_50_ 3.8 µM → 31 µM for **8c**→ **41c**, Figure 4a).

**Figure 4.**
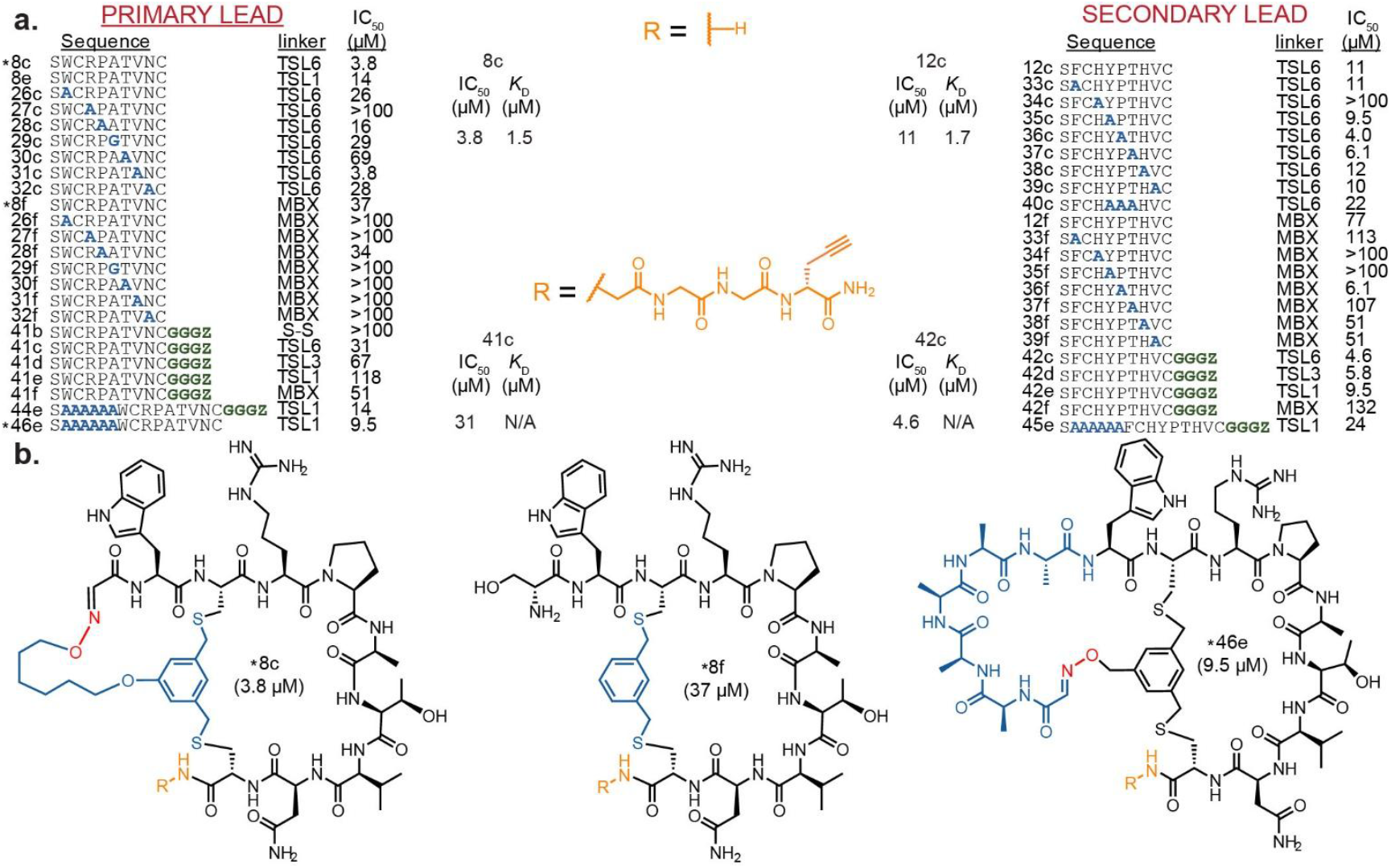
Linker and positional scan of lead sequences. a) IC_50_ values of the alanine-scanned and linker-modified variants of our lead binders, **8c** and **12c**, as determined by our ELSIAs to build an SAR profile, highlighting the effect that C-terminal elongation has on our lead sequences (Z = L-propargylglycine). b) Structures of reshaped variants of peptide **8** and their activities in ELISA.

The combined SAR data and synthetic availability positioned **8c** as the primary lead and **12c** as the secondary lead due to greater sensitivity of **8c** to structural modifications. We then systematically tested the role of the N-terminal ring and the resulting bicyclic architecture by replacing the TSL-6 linker with the dibromo-*m*-xylene (MBX) linker. The resulting monocyclic analogs of **8c** and **12c** exhibited a 7 to 10-fold decrease in activity (Figure 4a). The analogous TSL-6 to MBX replacement in the Ala-variants further highlighted the interplay of amino acid residues and bicyclic topology in binding both in **8c** and **12c** (Figure 4a). Systematic changes in ring geometry of the variants with the appended GGGZ terminal handle demonstrated a mild deterioration of binding upon contraction of the first ring from six to three to one carbon atoms using the TSL-6, TSL-3, and TSL-1 linchpins (**41c, 41d**, and **41e**) (**42c, 42d**, and **42e**), respectively; opening of the first ring while maintaining the geometry of the second ring significantly decreased the binding (MBX analogs: **41f** and **42f**). The disulfide analog of **8c** exhibited no measurable inhibition (Figure 4a, **41b**). Reshaping the N-terminal rings from the TSL-6 geometry to the Ala5-TSL1 geometry replaces the alkyl chain with the peptide chain, and we observed that both **8c** and **12c** tolerated such reshaping. Although not explored further, bicyclic geometries with eight amino acid positions in the N-terminal ring open the possibility for permutation of the amino acids in the N-terminal ring, similar to systematic dialing of the potency in the independent loops of disulfide-rich peptides^48, 49^. Due to the increased solubility of the Ala5-TSL1 analog of **8c** and the affinity being similar on ELISA, we decided to focus our investigation on this lead analog (*IC*_50_ 3.8 µM → 9.5 µM for **8c** → **46e**, Figure 4b).

### Embedding the NGS data and predicting new classes of binders

While classical analysis of leads in multi-round screening yielded productive leads, it restricted discovery to the vicinity of chemical space found within peptide **8c**. To explore broader chemical space, we re-examined the NGS data using machine learning (ML), training classifiers on peptide sequences enriched in Siglec-7 selections versus controls. Models trained on pooled rounds 2–4 achieved average AUC values of ∼0.64 and were used to rescore and rank enriched sequences. Of the top 20 predictions, 10 were synthesized and tested, yielding inhibitors with *IC*_50_ values ranging from 14 to 347 µM (**54c** and **49c** respectively), including two hits with potency comparable to **8c** (**51c** and **54c**, Figure 5). These results demonstrate that ML-guided prioritization can capture SAR trends in NGS data and nominate binders outside the space accessible to classical funneling.

**Figure 5.**
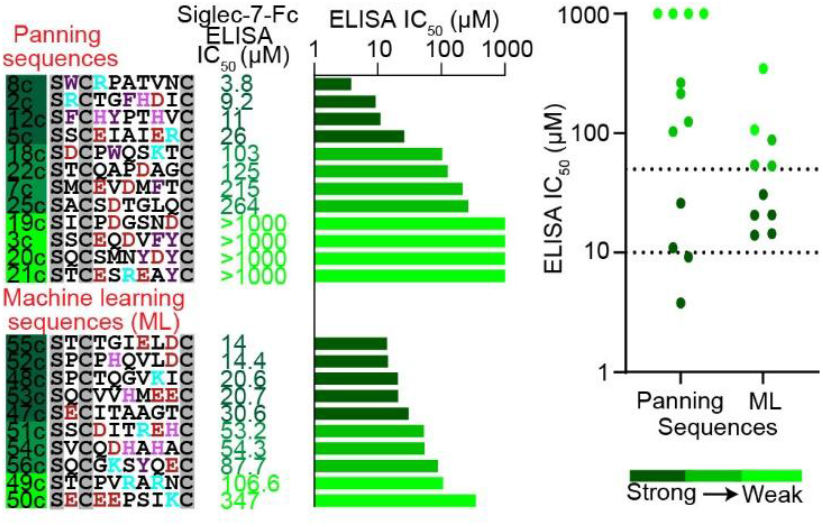
Prediction of new binders through ML of the 10 sequences synthesized with the TSL-6 linker and validated with competitive ELISA organized by affinity. Comparing the ELISA *IC*_50_s of the sequences we obtained through classical panning funneling against those predicted by ML.

### Structural biology investigation

To gain insight into the interaction of **46e** with Siglec-7, we employed high-resolution NMR spectroscopy and computational modeling. The analysis of the homonuclear (TOCSY, NOESY) and heteronuclear (HSQC) NMR spectra (Figure 6a) for the peptide enabled identification of the spin systems corresponding to the different amino acid types (Trp, Arg, Ala, Pro, Thr, Val, Asn, Cys) as well as those bearing an aromatic moiety. Interestingly, many NMR signals were duplicated in the NMR spectra, although with different intensities, especially those located on the “east side” of the molecule (marked in red in Figure 6d). This observation suggests the presence of a major and a minor species in solution. The possibility of the presence of two different molecules was assessed by using diffusion ordered (DOSY NMR) experiments (Figure S13). However, the observed diffusion coefficient was identical for both sets of NMR signals, strongly suggesting that they all belong to molecular entities with similar size. The analysis of variable temperature NMR experiments allowed the observation of significant changes in the shape of the NMR signals (Figure S14). The coalescence temperature was observed in ex-cess of 80 ^°^C, affirming the presence of rotamers (Figure S14). Moreover, NOESY (EXSY) experiments performed at a variety of mixing times (up to 2 s) did not show any exchange crosspeaks between the duplicated signals. Provided that conformers coexist in solution, this evidence suggests that the energy barrier for interconversion is rather large but attainable at high temperatures. Inter-residue Pro(d)/Arg(a) NOE cross peaks observed in the major species (Figure 6b) indicate a *trans*-Pro residue, as these protons are only spatially close in the *trans* geometry. In contrast, the minor species shows no such NOE, and its Pro signals appear at higher field, a hallmark of *cis*-Pro rotamers in small peptides (Figure 6b). This experimental evidence strongly suggests that the two species are due to *cis*/*trans* Pro isomerization.

**Figure 6.**
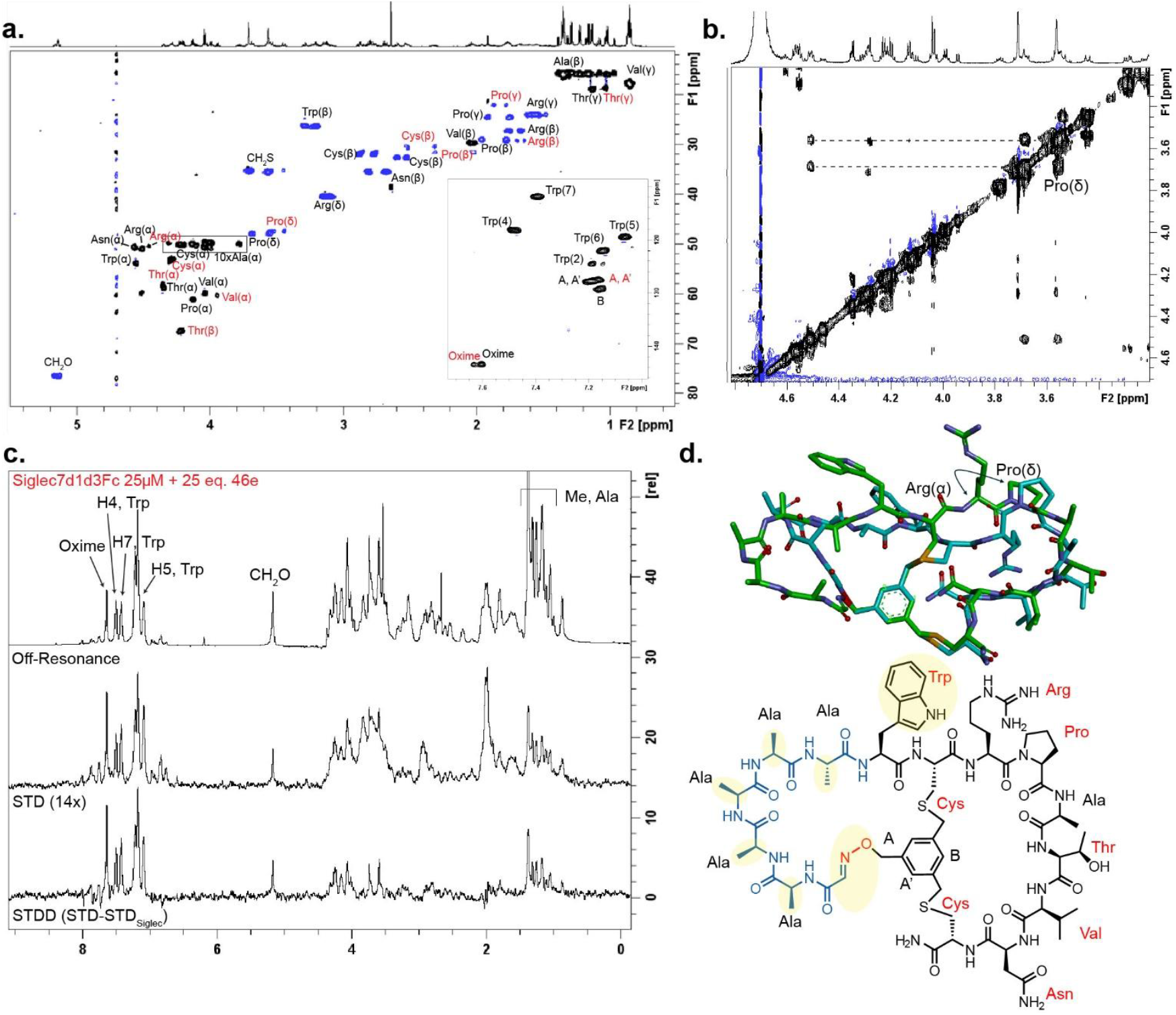
Structural NMR analysis of **46e** in D_2_O. a) The identification of the cross-peaks belonging to the different amino acid residues are indicated. Duplication of some signals suggest the presence of two distinct entities. The cross-peaks corresponding to the less populated *cis*-Pro rotamer are indicated in red. The aromatic region is highlighted in the bottom-right corner. b) Section of the 2D NOESY spectrum (200 ms mixing time) recorded for **46e** in D_2_O. NOE contacts between Pro(d) and Arg(a), characteristic of the *trans*-isomer, were found for the major species, but not for the minor one. c) ^1^H STD-NMR spectra recorded for the mixture of Siglec-7_d1d3_ and the bicyclic peptide in a 1:25 molar ratio. STD signals are clearly identified for the Trp side chain and olefinic protons, along with many methyl groups. The blank STD experiment recorded for the peptide ligand alone did not show an STD response. The *off-resonance* spectrum for the mixture is shown in the top trace. The STD experiment obtained for Siglec-7_d1d3_ alone was subtracted from the STD spectrum (middle trace) to provide the final STDD spectrum, free of interferences, which shows the peptide NMR signals corresponding to the amino acid residues highlighted in the molecular scheme on the right-hand side. Identical results were obtained when the R124A mutant of Siglec-7_d1d3_ was employed as a receptor (Figure S16). d) The superimposed global minima identified for the *cis*- and *trans*-Pro rotamers and chemical structure of **46e**.

Saturation transfer difference nuclear magnetic resonance (STD-NMR) experiments were then used to probe peptide–receptor recognition. The interaction of the peptide with distinct Siglec-7 variants was then monitored at the molecular level. Specifically, the full-length extracellular domain (ECD) of Siglec-7d1d3 (residues 19-357) and its mutant R124A, fused to the human IgG1 Fc region, as well as the single V-Ig set domain (Siglec-7d1) were employed for the molecular recognition studies using STD-NMR experiments. R124 has been demonstrated to be located at the canonical binding site of the V-Ig set domain and is essential for sialic acid binding. Interestingly, STD-NMR responses were only observed for different peptide NMR signals in the presence of either the full-length extracellular domain (ECD) of Siglec-7d1d3 or of its R124A mutant (Figure 6c). No STD response was observed when the isolated V-Ig set domain (Siglec-7d1) was used as a putative receptor (Figure S16). In contrast, identical STD-NMR responses of the peptide (Figure 6c) were obtained in the presence of either Siglec-7d1d3 variants. Clear STD intensities were observed for different methyl groups, as well as for the proton signals of the Trp moiety and for the olefinic and *O*-methylene protons, among others (Figure 6d). STD responses for the major and minor species were identified for the olefinic proton closest to the oxime moiety. These results reveal that the two rotamers of the peptide indeed interact with Siglec-7. However, according to the data obtained for the experiments carried out with the three Siglec-7 variants, the interaction does not take place at the canonical sialic acid binding site of Siglec-7 at the V-Ig set domain (Siglec-7d1), but at another domain. Together, these studies establish that bicyclic peptide **46e** adopts distinct rotameric conformations in solution and recognizes Siglec-7 through a non-canonical binding interface, providing structural rationale for the inhibitory activity observed by our biochemical assays.

### Validation of lead peptide using multivalent display in liquid glycan arrays

We sought to evaluate the performance of the lead bicycle in cell-based assays alongside previously reported glycan binders to Siglec-7 on the surface of cells measured by Liquid Glycan Array (LiGA) technology^50^, we re-synthesized **46e** with a C-terminal PEG-azide handle and conjugated it to dibenzocyclooctyne installed on major coat proteins (pVIII) on M13 phage analogous to the synthesis of the LiGA components.^43^ This liquid arrays allowed the display of glycans alongside bicyclic peptides, and systematic control of the valency of the display of both structures was encoded by DNA. In preliminary tests, we observed unexpected binding of wild-type M13 virion to purified Siglec-7 protein (Figure S19). To prevent this undesired binding, we conjugated equal concentrations of **57e** and PEG8-azide to M13 virion. Matrix-assisted laser desorption-ionization (MALDI) by time-of-flight (TOF) mass spectrometry confirmed this conjugation (Figure 7a). The disappearance of the DBCO peak in MALDI experiments allowed us to approximate conjugation, as previously reported,^43^ we approximated the density of 320 copies of **57e** and PEG8-azide. The resulting phage construct exhibited significantly higher binding to purified Siglec-7 protein when compared to all controls (BSA-coated well or phage conjugated to PEG alone, Figure 7b).

**Figure 7.**
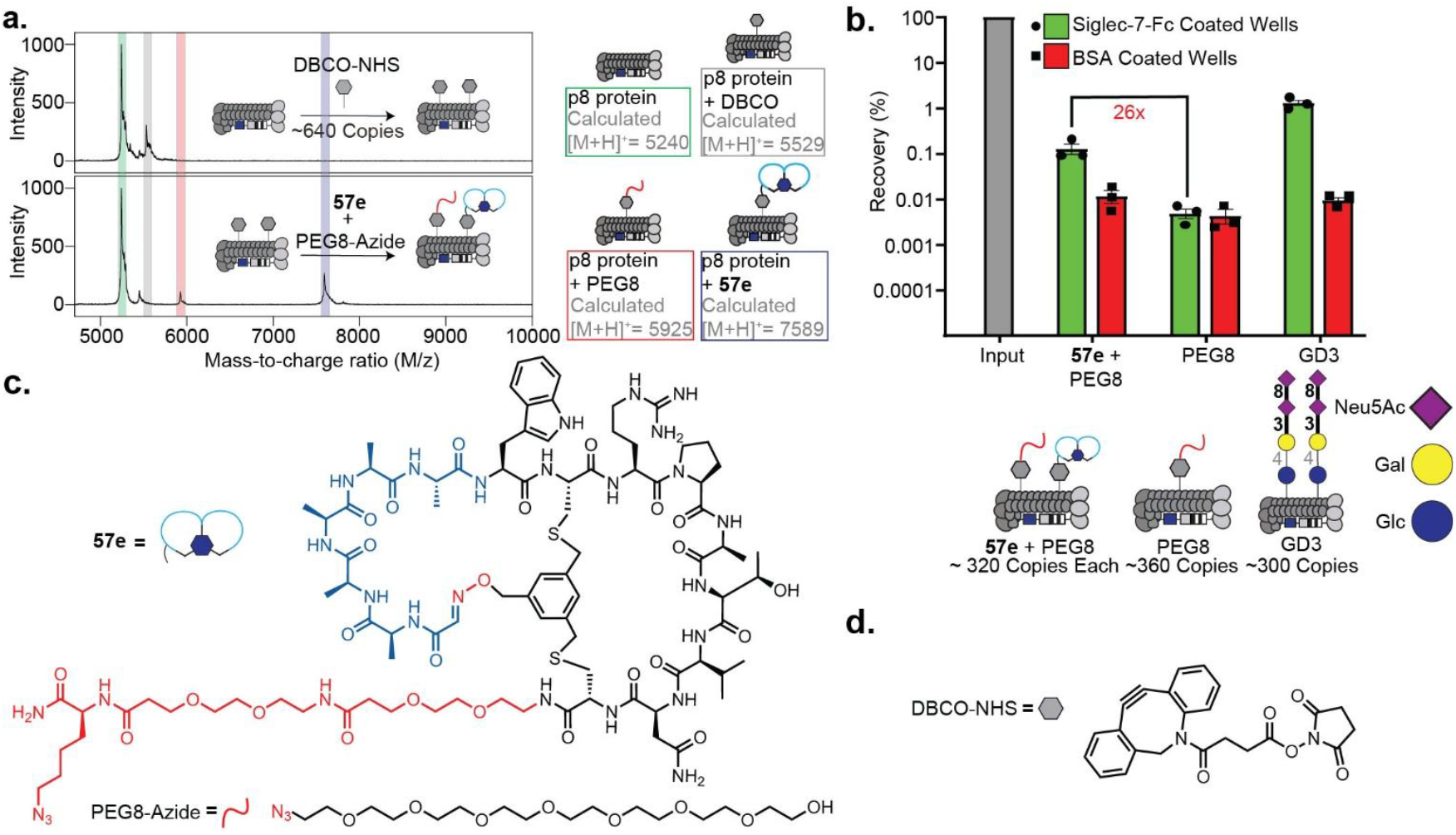
Conjugation of **57e** with PEG8-azide and validation. a) To pVIII proteins on M13 phage, DBCO was conjugated at a density of roughly 640 copies verified and monitored via MALDI-TOF. Equal concentrations of both **57e** and PEG8-azide were added resulting in the peptide-conjugated phage construct. b) Phage recovery assay (Figure S18) of the phage constructs shown below the graph against Siglec. The recovery assay highlights the affinity of the peptide construct to the positive wells (wells coated by Siglec-7-Fc) shown in green against the affinity of the construct to negative wells (wells only coated by BSA) shown in red. The peptide-conjugated phage shows a measurable (26-fold) enrichment over the negative control. c) The chemical structures of both **57e** and PEG8-Azide used in the conjugation. d) The chemical structure of commercially available DBCO-NHS used for conjugation.

Following the confirmation of binding of the multivalent display of bicyclic peptides to purified Siglec-7, we tested binding of the same construct to several cell types that express Siglec-7. Binding of the bicycle-displaying construct to Siglec-7^+^ CHO cells and Jurkat cells engineered to overexpress Siglec-7 was stronger than binding of PEG-blocked phage. However, the same bicycle construct also bound to CHO cells and Jurkat cells that did not express Siglec-7 (Figure S22). In contrast, phage constructs displaying glycan GD3 exhibited selective binding only to Siglec-7^+^ but not Siglec-7^-^ cells (Figure S22). To test whether *cis*-inhibition by sialic acid residues in the glycocalyx play any role in these observations, we tested binding to U937 CMAS^KO^ cells^51^ as well as to their wild-type counterparts. Again, GD3 exhibited significantly enhanced binding to “unmasked” Siglec-7 on U937 CMAS^KO^ cells, whereas bicycle peptides exhibited comparable binding to both cell types (Figure S22).

We next employed NGS to perform a side-by-side comparison of the bicycle and 100 glycans, components of LiGA,^43^ in the same cell-based assay. Multivalent display of bicyclic peptide and PEGylated phage as a negative control was mixed with LiGA and screened against hSiglec-7^+^ Jurkat and CHO cells, and their siglec-7^-^ counterparts as well as U937 CMAS^KO^ (positive or wild-type) our peptide-conjugated phage construct showed measurable binding, while sialylated glycans only showed measurable binding to the positive lines (Figure 8).

**Figure 8.**
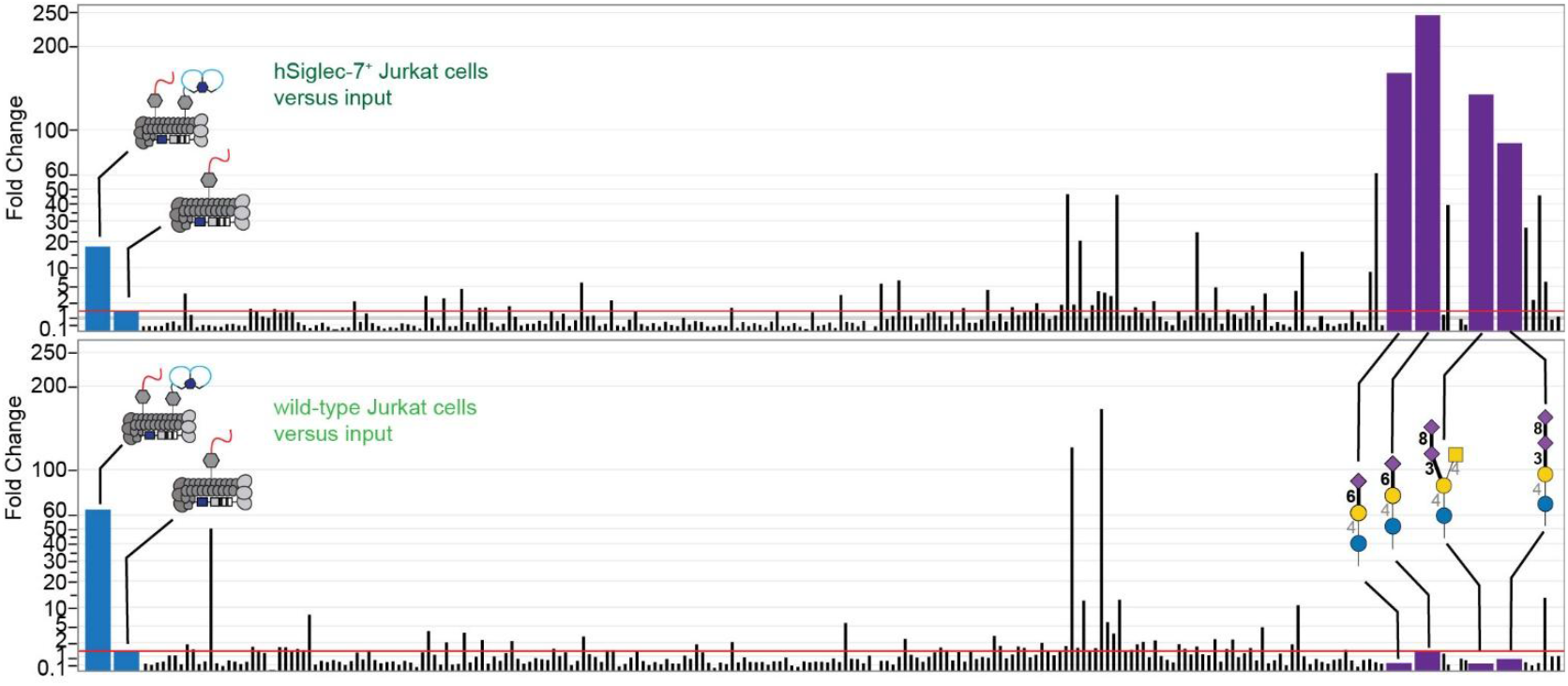
Bar plot of the LiGA mixture against Jurkat cells. The bar plots highlight the fold change of a phage construct against hSiglec-7^+^ or wild-type cells as a control. Relevant phage constructs (sialylated glycans, peptide-conjugated phage, and PEGylated phage) have been expanded and colored purple and blue, respectively. Fully labeled plots for Jurkat, CHO, and U937 cells are shown in Figure S23, S24, and S25, respectively.

## Conclusion

Previously, our group^33, 36, 37, 40^ and other groups^27, 52, 53^ directed the selection of GBP inhibitors toward the glycan-binding site using DNA-encoded glycopeptide libraries. In this study, we deliberately omitted the complex sialo-glycan warhead to simplify downstream synthesis and optimization. This design is not new *per se* and identification of peptide-based binders for GBP using glycan-free (aglycon) libraries has been attempted multiple times^52, 54, 55^. To the best of our knowledge, the feasibility of identifying aglycone inhibitors of Siglec has not been reported, although unpublished attempts to discover Siglec-10 antagonists exist in mRNA display fields^56^. Our screening efforts yielded bicyclic peptide binders that we classified into inhibitory and non-inhibitory groups. In parallel, application of ML to NGS datasets enabled us to prioritize additional sequences beyond the chemical space of the **8c** family. Synthesis and testing of these ML-predicted peptides yielded new binders with low-micromolar potencies, including two with activities comparable to our best experimental leads. Although not stronger than the primary hits, these results validate ML-guided approaches as a complementary tool to expand the pool of validated Siglec-7 inhibitors. Structural investigations by NMR revealed that our most potent inhibitor **46e** exists as a *cis*/*trans* rotameric mixture; it interacts with Siglec-7 at a noncanonical binding interface rather than at the sialic acid–binding pocket via an allosteric mechanism of inhibition. Multivalent presentation by conjugation of our lead peptide **57e** to M13 phage using LiGA methodology^43^ results in binding to pure Siglec-7 but not cells. This demonstrates that peptide affinity in their monovalent form, leveraging their geometry and presentation can be used to achieve robust recognition in a multivalent context. In conclusion, this work provides the first demonstration of bicyclic peptide macrocycles as inhibitors of Siglec-7 and highlights both the opportunities and challenges in designing peptide-based inhibitors of GBPs. Our findings underscore the importance of non-canonical binding sites, the potential of multivalent display to boost functional recognition, and the value of ML to expand chemical diversity. While these current leads are allosteric inhibitors and face solubility-related challenges, strategies such as ring expansion, physicochemical optimization, and guided multivalent design offer promising routes toward higher-affinity and more developable inhibitors.

## ASSOCIATED CONTENT

**Supporting Information 1**. Workflow, abbreviations, and chemical protocols (Section S1-S3); Siglec-7 expression protocol, panning and deep sequencing protocol, NMR analysis, and machine learning information (Section S4); SPR, ELISA, and phage recovery measurements protocols (Section S5); summary of sequences synthesized (Table S1); Data summary pages (Section S6).

**Supporting information 2**. Synthesis summaries of peptides listed in Table S1.

## AUTHOR INFORMATION

## Notes

The authors declare the following competing financial interest(s): R.D. is the C.S.O. and a shareholder of 48Hour Discovery Inc., the company that licensed US9958437B2 and WO2020107118AI patents that protect modification and analysis strategies used in this report. 48HD maintains https://48hd.cloud/ used for deep sequencing analysis. Data Availability: All raw deep-sequencing data is publicly available on https://48hd.cloud/ with dataspecific URL listed in Supplementary Figure S10, S11 and Table S2, S3.

## ACKNOWLEDGMENT

We thank Mark Miskolzie and Nupur Dabral at the University of Alberta NMR facility.

CIC bioGUNE members thank Agencia Estatal de Investigacio n of Spain for grants PID2024-157610OB-100 (J. J.-B.), PID2023-150777O B-I00 (J. E.-O.), PID2023-150779OA-I00 (A. G.), and the Severo Ochoa Center of Excellence Accreditation CEX2021-001136-S, funded through MCIN/AEI/10.13039/501100011033. They also thankfully acknowledge the NMR resources and the technical support provided by the LRE of the Spanish ICTS Red de Laboratorios de RMN de Biomole culas (R-LRB) and CIBERES, an initiative of Instituto de Salud Carlos III (ISCIII, Madrid, Spain).

Matthew S. Macauley thanks NSERC and GlycoNet for funding.

