## Supporting Information 1 for "Non-Carbohydrate Inhibitors of Sialic Acid-binding Immunomodulatory-type Lectin-7 (Siglec-7) Discovered from Genetically Encoded Bicyclic Peptide Libraries"

4.7.3.1 Panning on a Kingfisher instrument ... 21
4.7.3.2 Panning of purified recombinant Siglec-7 protein ... 22
**Figure S7.** Visual flowchart of panning done on Siglec-7 ... 22
4.7.3.3 Panning of purified recombinant Siglec-9 protein ... 25
**Figure S8.** Visual flowchart of panning done on Siglec-9 ... 25
4.7.3.4 Focused library panning ... 27
**Figure S9**. FC scatter plot ... 29
4.7.3.5 PCR amplification protocol for deep sequencing on Illumina ... 29
4.7.4 Illumina deep sequencing ... 30
4.7.4.1 Processing of the Illumina data ... 30
**Table S2**. Illumina data for rounds 1–4 against Siglec-7 ... 31
**Figure S10**. Screenshot of NGS flowchart for Siglec-7 ... 31
**Table S3**. Illumina data for the TSL6-modified focused library against Siglec-7 ... 32
**Figure S11.** Screenshot of NGS flowchart for Siglec-7 GS2 focus library ... 32
**Table S4**. Illumina data for rounds 1–4 against Siglec-9 ... 33
**Figure S12**. Screenshot of the representative data entry web address. ... 33
4.8 NMR analysis of **46e** ... 33
4.8.1 General workflow of **46e** ... 33
1.1.1 2D DOSY spectrum ... 35
**Figure S13.** 2D DOSY spectrum recorded for the bicyclic peptide in $D_2O$. ... 35
4.8.2 Variable temperature NMR ... 35
**Figure S14.** Variable temperature NMR of **46e** at various temperature points. ... 36
**Figure S15.** Variable temperature NMR of **58c** at various temperature points. ... 37
4.8.3 NOESY analysis ... 37
4.8.4 Expression of Siglec-7 for STD-NMR ... 38
**Figure S16.** $^1$H STD NMR of isolated V-Ig set domain (Siglec-7$d_1$) ... 39
4.8.5 Computational modelling ... 39
4.9 Machine learning predications ... 40
4.9.1 Prediction of new classes of binders ... 40
**5 Measurements ... 41**
5.1 General procedure for SPR ... 41

#### 1 Project flowchart

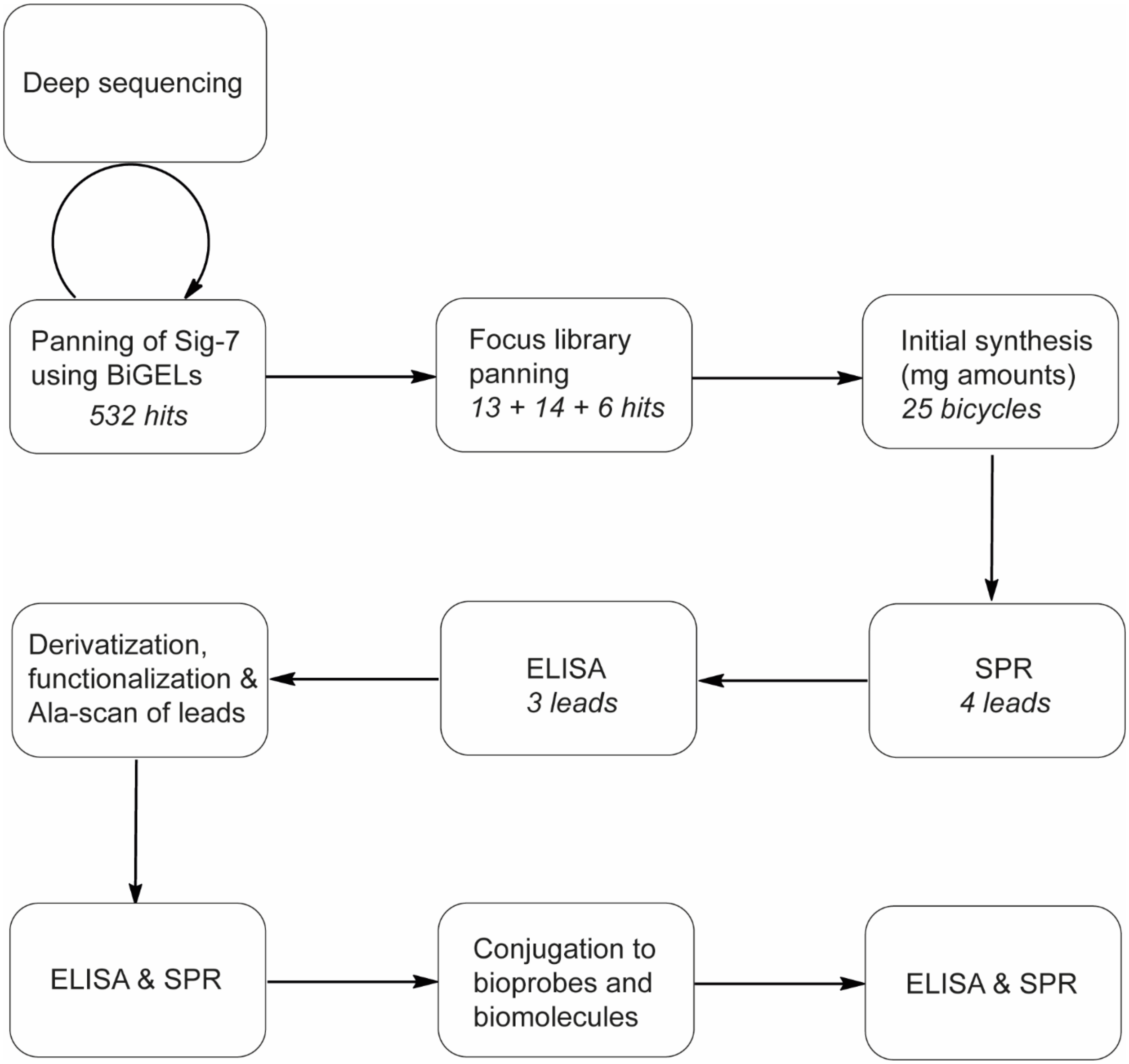

**Figure S1**. Overall project workflow

#### 2 List of abbreviations

| | |
|---|---|
| AOB | aminooxybiotin |
| B | fmoc-AEEP; (Fmoc-9-amino-4,7-dioxanonanoic acid) |
| BCA | bicinchoninic acid |
| BiGELs | bicyclic genetically encoded libraries |
| Boc | *tert*-butyloxycarbonyl |
| DBCO | dibenzocyclooctyne |
| DE | differential enrichment |
| DIPEA | *N*,*N*-diisopropylethyl amine |
| DMF | *N*,*N*-dimethylformamide |
| DMSO | dimethyl sulfoxide |
| ECD | extracellular domain |
| ELISA | enzyme-linked immunosorbent assay |
| EDT | ethanedithiol |
| EDC | 1-ethyl-3-(3-dimethylaminopropyl)carbodiimide hydrochloride |
| ESI | electrospray ionization |
| Fc | fragment crystallizable region |
| FC | fold change |
| Fmoc | fluorenylmethoxycarbonyl |
| GBP | glycan-binding proteins |
| HR | high resolution |
| HBTU | hexafluorophosphate benzotriazole tetramethyl uronium |
| HPLC | high-performance liquid chromatography |
| HRP | horse radish peroxidase |
| LC–MS | liquid chromatography–mass spectrometry |
| LiGA | liquid glycan array |
| MALDI-TOF | matrix-assisted laser desorption-ionization by time-of-flight |
| MBX | $\alpha,\alpha$-dibromo-*m*-xylene |
| ML | machine learning |
| MWCO | molecular weight cut-off |
| NGS | next-generation sequencing |
| NHS | *N*-hydroxysuccinimide |

| | |
|---|---|
| PBS | phosphate-buffered saline |
| PCR | polymerase chain reaction |
| PEG | polyethylene glycol |
| pfu | plaque-forming unit |
| propG | propargyl glycine |
| RP | reverse phase |
| rt | room temperature |
| SAR | structure-activity relationship |
| SDS-PAGE | sodium dodecyl sulfate–polyacrylamide gel electrophoresis |
| Siglecs | sialic acid-binding immunodulatory-type lectins |
| SPR | surface plasmon resonance |
| STD-NMR | saturation transfer difference nuclear magnetic resonance |
| TCEP | tris(2-carboxyethyl)phosphine |
| TEV | tobacco etch virus |
| TFA | trifluoroacetic acid |
| TSL | twofold-symmetric linker |
| TIPS | triisopropylsilane |
| THPTA | tris(3-hydroxypropyltriazolylmethyl)amine |
| TMB | 3,3′,5,5′-Tetramethylbenzidine |
| TRIS | tris(hydroxymethyl)aminomethane |
| X | L-azidolysine |
| Z | L-propargylglycine |

#### 3 Chemistry

##### 3.1 General chemistry information

Chemical reagents and solvents were purchased from Sigma-Aldrich or Fisher Scientific unless noted otherwise. TSL-1, TSL-3 and TSL-6 were prepared as described previously.[1] TCEP was purchased from Soltech Ventures. The appropriately Fmoc-protected amino acid reagents for automated solid-phase peptide synthesis were purchased from ChemPep.

LC-MS analysis of the reported peptides was obtained on Agilent Technologies 6130 LC-MS. A gradient of solvent A (dd$H_2O$) and solvent B (MeCN/dd$H_2O$ 95/5) was run at a flow rate of 0.5 mL/min (0–4.0 min 5% B; 4.0–5.0 min 5%–60% B; 5.0–5.5 min 60–100% B; 5.5–7.5 100% B, 7.5-11 min 100–5% B) Purity was estimated using the resulting spectra.

Semipreparative RP-HPLC was conducted using a Waters Symmetry prep 19×50 mm C18 column and a Waters 2489 UV detector. Two gradients, or very slight variations thereof as delineated on their respective data summary pages, at a flow rate of 10 mL/min, were used to purify the peptides on flash RP chromatography over 30 min. Gradient A was 0–2 min, 2%–5% MeCN/TFA (1000/1) in dd$H_2O$/TFA (1000:1); 2–21 min 2–50% MeCN/TFA (1000:1) in dd$H_2O$/TFA (1000:1); 21–23 min, 50–100% MeCN/TFA (1000:1) in dd$H_2O$/TFA (1000:1); 23–26 min 100% MeCN/TFA (1000:1); 26–30 min 100%–2% MeCN/TFA (1000:1) in dd$H_2O$/TFA (1000:1). Gradient B was 0–2 min, 2%–5% MeCN/TFA (1000/1) in dd$H_2O$/TFA (1000:1); 2–21 min 2–70% MeCN/TFA (1000:1) in dd$H_2O$/TFA (1000:1); 21–23 min, 70–100% MeCN/TFA (1000:1) in dd$H_2O$/TFA (1000:1); 23–27 min 100% MeCN/TFA (1000:1); 27–30 min 100%–2% MeCN/TFA (1000:1) in dd$H_2O$/TFA (1000:1).

Flash RP chromatography was carried out using C-18 as the solid phase performed on a Teledyne Isco Combiflash system. Two gradients, or very slight variations thereof as delineated on their respective data summary pages, were used to purify the peptides on flash RP chromatography over 12 min: gradient A or gradient B. Gradient A was 0–1 min 100% dd$H_2O$/TFA (1000:1), 1–10 min 0–50% MeCN/TFA (1000:1) in dd$H_2O$/TFA (1000:1), 10–12 min 100% MeCN/TFA (1000:1); gradient B was 0–1 min 100% dd$H_2O$/TFA (1000:1), 1–10 min 0–70% MeCN/TFA (1000:1) in dd$H_2O$/TFA (1000:1), 10–12 min 100% MeCN/TFA (1000:1). Further, three column sizes containing C-18 were used: column A (50 g), column B (15.5 g) or column C (5.5 g) with flow rates of 40 mL/min, 30 mL/min and 18 mL/min, respectively. Removal of aqueous solvents was performed using a Labconco Freezone 2.5w lyophilizer. Throughout this supplementary information, **Z** denotes L-propargylglycine.

Exact masses of the precursors used, and their overall yields are listed for each peptide on their respective data summary pages (Supplementary Information 2); these data summary pages appear in the order the sequence is described in this section. All the sequencing results are available on the

48Hour Discovery cloud: https://48hd.cloud/. All the 20 × 20 plots were generated on the 48Hour Discovery cloud: https://48hd.cloud/.

**Table S1.** A summary of every peptide synthesized and/or screened for binding.

| # | Sequence | **a** | **b** | **c** | **d** | **e** | **f** | **g** | **h** | **i** | **j** | **k** | **l** |
|---|---|---|---|---|---|---|---|---|---|---|---|---|---|
| | | Linear | S-S | TSL-6 | TSL-3 | TSL-1 | MBX | TSL6-PEG$_n$-Biotin | TSL3-PEG$_n$-Biotin | TSL1-PEG$_n$-Biotin | MBX-PEG$_n$-Biotin | TSL6-PEG$_n$-N$_3$ | TSL6-PEG3-N$_3$-PEG3-Biotin |
| **1** | SHCAAYDQMC | | | **1c** | | | | | | | | | |
| **2** | SRCTGFHDIC | | | **2c** | | | | | | | | | |
| **3** | SSCEQDVFYC | | | **3c** | | | | | | | | | |
| **4** | SQCRPDQEIC | | | **4c** | | | | | | | | | |
| **5** | SSCEIAIERC | | | **5c** | | | | | | | | | |
| **6** | SECTMEGTVC | | | **6c** | | | | | | | | | |
| **7** | SMCEVDMFTC | | | **7c** | | | | | | | | | |
| **8** | SWCRPATVNC | | | **8c** | | **8e** | **8f** | | | | | | |
| **9** | SSCFDSQSDC | | | **9c** | | | | | | | | | |
| **10** | SDCPQATIIC | | | **10c** | | | | | | | | | |
| **11** | SYCHDTAGRC | | | **11c** | | | | | | | | | |
| **12** | SFCHYPTHVC | | | **12c** | | | **12f** | | | | | | |
| **13** | SICTGEIEHC | | | **13c** | | | | | | | | | |
| **14** | SGCIIEGMQC | | | **14c** | | | | | | | | | |
| **15** | SDCRNFISAC | | | **15c** | | | | | | | | | |
| **16** | SLCTTIFEEC | | | **16c** | | | | | | | | | |
| **17** | SSCQDAFDKC | | | **17c** | | | | | | | | | |
| **18** | SDCPWQSKTC | | | **18c** | | | | | | | | | |
| **19** | SICPDGSNDC | | | **19c** | | | | | | | | | |
| **20** | SQCSMNYDYC | | | **20c** | | | | | | | | | |
| **21** | STCESREAYC | | | **21c** | | | | | | | | | |
| **22** | STCQAPDAGC | | | **22c** | | | | | | | | | |
| **23** | SYCNVIAGIC | | | **23c** | | | | | | | | | |
| **24** | SACAWHQHAC | | | **24c** | | | | | | | | | |
| **25** | SACSDTGLQC | | | **25c** | | | | | | | | | |
| **26** | S**A**CRPATVNC | | | **26c** | | | **26f** | | | | | | |
| **27** | SWC**A**PATVNC | | | **27c** | | | **27f** | | | | | | |
| **28** | SWCR**A**ATVNC | | | **28c** | | | **28f** | | | | | | |
| **29** | SWCRP**G**TVNC | | | **29c** | | | **29f** | | | | | | |

| | | | | | | | | | | | | | |
|---|---|---|---|---|---|---|---|---|---|---|---|---|---|
| **30** | SWCRPA**A**VNC | | | **30c** | | | **30f** | | | | | | |
| **31** | SWCRPAT**A**NC | | | **31c** | | | **31f** | | | | | | |
| **32** | SWCRPATV**A**C | | | **32c** | | | **32f** | | | | | | |
| **33** | S**A**CHYPTHVC | | | **33c** | | | **33f** | | | | | | |
| **34** | SFC**A**YPTHVC | | | **34c** | | | **34f** | | | | | | |
| **35** | SFCH**A**PTHVC | | | **35c** | | | **35f** | | | | | | |
| **36** | SFCHY**A**THVC | | | **36c** | | | **36f** | | | | | | |
| **37** | SFCHYP**A**HVC | | | **37c** | | | **37f** | | | | | | |
| **38** | SFCHYPT**A**VC | | | **38c** | | | **38f** | | | | | | |
| **39** | SFCHYPTH**A**C | | | **39c** | | | **39f** | | | | | | |
| **40** | SFCH**AAA**HVC | | | **40c** | | | | | | | | | |
| **41** | SWCRPATVNCGGGZ | **41a** | **41b** | **41c** | **41d** | **41e** | **41f** | **41g** | **41h** | **41i** | **41j** | **41k** | **41l** |
| **42** | SFCHYPTHVCGGGZ | **42a** | **42b** | **42c** | **42d** | **42e** | **42f** | **42g** | | | | **42k** | **42l** |
| **43** | SSCEIAIERCGZ | **43a** | | **43c** | | | | **43g** | | | | | |
| **44** | S**AAAAA**WCRPATVNCGGGZ | | | | | **44e** | | | | | | | |
| **45** | S**AAAAA**FCHYPTHVCGGGZ | | | | | **45e** | | | | | | | |
| **46** | S**AAAAA**WCRPATVNC | **46a** | | | | **46e** | | | | | | | |
| **47** | SECITAAGTC | | | **47c** | | | | | | | | | |
| **48** | SPCTQGVKIC | | | **48c** | | | | | | | | | |
| **49** | STCPVRARNC | | | **49c** | | | | | | | | | |
| **50** | SECEEPSIKC | | | **50c** | | | | | | | | | |
| **51** | SSCDITREHC | | | **51c** | | | | | | | | | |
| **52** | SPCPHQVLDC | | | **52c** | | | | | | | | | |
| **53** | SQCVVHMEEC | | | **53c** | | | | | | | | | |
| **54** | SVCQDHAHAC | | | **54c** | | | | | | | | | |
| **55** | STCTGIELDC | | | **55c** | | | | | | | | | |
| **56** | SQCGKSYQEC | | | **56c** | | | | | | | | | |
| **57** | S**AAAAA**WCRPATVNCBBX | | | | | **57e** | | | | | | | |
| **58** | SFCHYPTHVCGGGX | | | **58c** | | | | | | | | | |

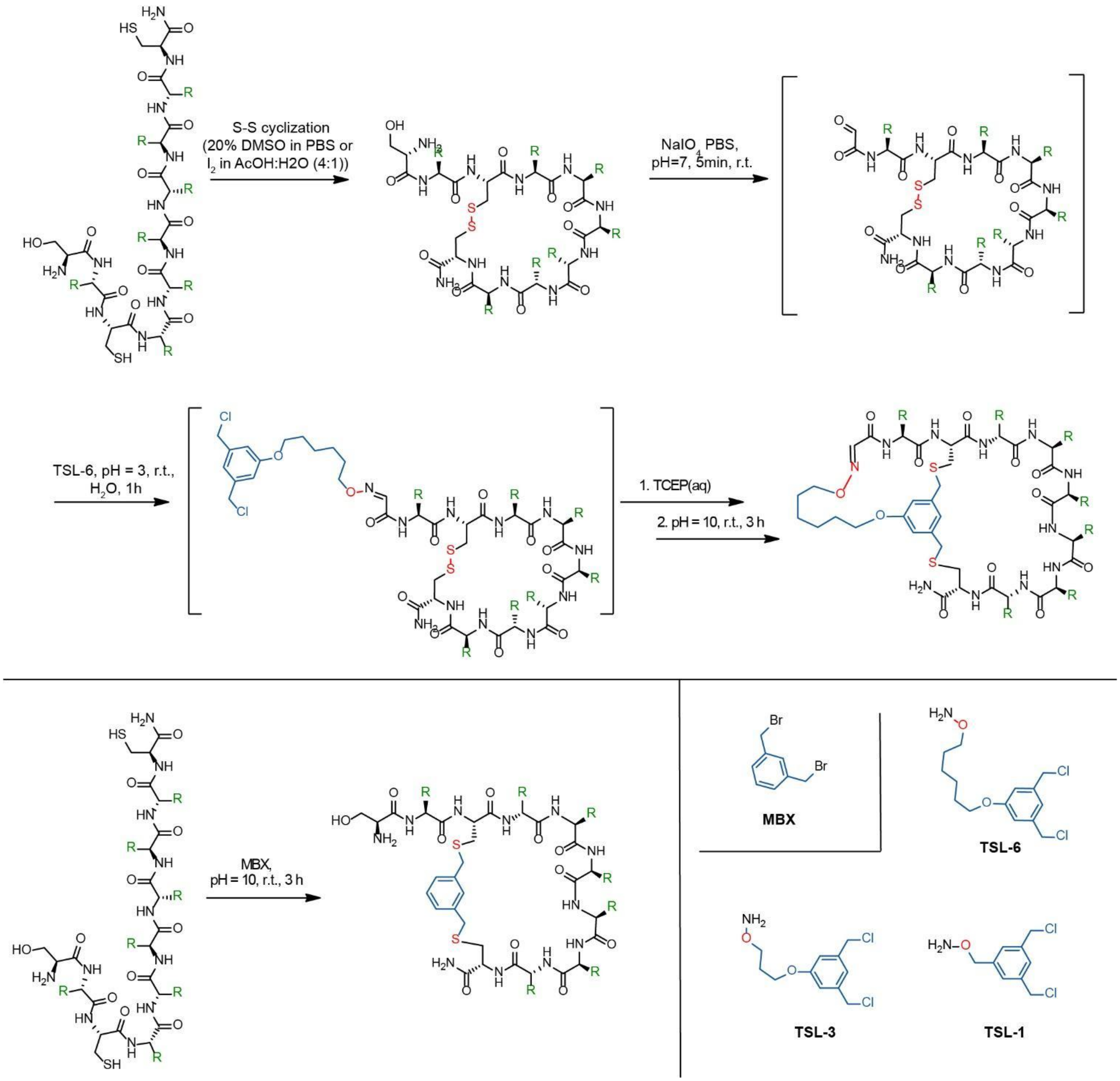

**Figure S2.** General synthetic access to mono- and bicyclic peptides.

3.2 General procedure for automated solid-phase peptide synthesis

In a 50-mL polypropylene centrifuge tube, linear peptide (5–20 mg, ~0.003–0.01 mmol) was dissolved in 8 mL DMSO and to this, PBS (32 mL) was added. **Note:** The maximum concentration ever reached was 0.5 mg peptide/mL oxidizing solution. The reaction mixture was subsequently agitated for 48 h on a shaker at rt. After 48 h, the crude disulfide bridge-bearing peptide was directly purified by flash RP chromatography (column A, B or C; gradient A). The fractions containing the product were pooled, flash frozen and lyophilized to provide the product as a fluffy white powder.

3.3 General procedure for the five-step synthesis of the macrobicycles (Procedure A)

In a 50-mL polypropylene centrifuge tube, linear peptide (5–20 mg, ~0.003–0.01 mmol) was dissolved in 8 mL DMSO and to this, PBS (32 mL) was added. **Note:** The maximum concentration ever reached was 0.5 mg peptide/mL oxidizing solution. The reaction mixture was subsequently agitated for 48 h on a shaker at rt. After 48 h, the crude disulfide bridge-bearing peptide was directly purified by flash RP chromatography (column A, B or C; gradient A). The fractions containing the product, as confirmed by LC-MS analysis, were pooled in a 50-mL polypropylene centrifuge tube. The pH of the reaction mixture was adjusted to ~7 using $NaHCO_3/Na_2CO_3$(aq) buffer (pH = 10). To this reaction mixture, freshly prepared* $NaIO_4$(aq) (1.2 equiv, 100 mM stock) was added, and the mixture was agitated for 5 min in the dark on a shaker. After 5 min, the reaction mixture was directly purified using flash RP chromatography (column A, B or C; gradient A). As determined by LC-MS, the fractions containing the purified aldehyde were pooled in a 50-mL polypropylene centrifuge tube. The pH was checked to ensure it was ~4; if it was not, the pH was decreased to 3–4 using dilute TFA(aq). To the reaction mixture, **TSLx** (2 equiv, 30 mM stock in MeCN) was added, and the mixture was shaken for 1 h. After 1 h, TCEP (4 equiv, 100 mM stock) was added, and the reaction mixture was further shaken for 30 min. After 15 min, the pH was increased to 9–10 using $NaHCO_3/Na_2CO_3$(aq) buffer. Shaking was continued for another 3 h. Complete bicyclization can be confirmed LC-MS analysis. The reaction mixture was subsequently purified by flash RP chromatography (column A, B or C; gradient B) or HPLC. The fractions containing the product were pooled, flash frozen and lyophilized to provide the product as a fluffy white powder.

*The $NaIO_4$(aq) stock solution must be prepared freshly and used within 4–6 h; all other reagents are stable in solution for prolonged periods (at least one month) at 4 °C.

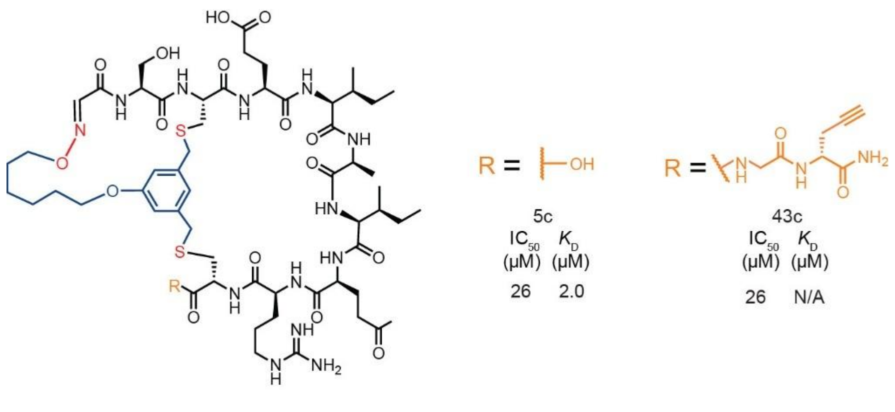

**Figure S3**. Effect of C-terminal elongation on **5c**

3.4 General procedure for the five-step synthesis of the Met-containing macrobicycles (Procedure B)

In a 50-mL polypropylene centrifuge tube, linear peptide (5–20 mg, ~0.003–0.01 mmol) was dissolved in 8 mL DMSO and to this, PBS (32 mL) was added. **Note:** The maximum concentration ever reached was 0.5 mg peptide/mL oxidizing solution. The reaction mixture was subsequently agitated for 48 h on a shaker at rt. After 48 h, the crude disulfide bridge-bearing peptide was directly purified by flash RP chromatography (column A, B or C; gradient A). The fractions containing the product, as confirmed by LC-MS analysis, were pooled in a 50-mL polypropylene centrifuge tube. The pH of the reaction mixture was adjusted to ~7 using $NaHCO_3$/$Na_2CO_3$(aq) buffer (pH = 10). To this reaction mixture, freshly prepared* $NaIO_4$(aq) (1.2 equiv, 100 mM stock) was added, and the mixture was agitated for 5 min in the dark on a shaker. After 5 min, the reaction mixture was quenched using methionine (5.0 equiv, 100 mM). The mixture was then purified using flash RP chromatography (column A, B or C; gradient A). As determined by LC-MS, the fractions containing the purified aldehyde were pooled in a 50-mL polypropylene centrifuge tube. The pH was checked to ensure it was ~4; if it was not, the pH was decreased to 3–4 using dilute TFA(aq). To the reaction mixture, **TSLx** (2 equiv, 30 mM stock in MeCN) was added, and the mixture was shaken for 1 h. After 1 h, TCEP (4 equiv, 100 mM stock) was added, and the reaction mixture was further shaken for 30 min. After 15 min, the pH was increased to 9–10 using $NaHCO_3$/$Na_2CO_3$(aq) buffer. Shaking was continued for another 3 h. Complete bicyclization can be confirmed by LC-MS analysis. The reaction mixture was subsequently purified by flash RP chromatography (column A, B or C; gradient B) or HPLC. The fractions containing the product were pooled, flash frozen and lyophilized to provide the product as a fluffy white powder.

*The $NaIO_4$(aq) stock solution must be prepared freshly and used within 4–6 h; all other reagents are stable in solution for prolonged periods (at least one month) if stored at 4 °C.

3.5 General procedure for cyclization using MBX

Pure linear peptide (5–10 mg, 0.03–0.07 mmol) was dissolved in $H_2O$/MeCN (1:1, maximum 1 mg peptide/mL solute) in a 15-mL centrifuge tube and to it a solution of MBX (1.2 equiv, 100 mM stock solution in MeCN) was added. Tris-HCl buffer (500 $\mu$L, 500 mM, pH 8.5, final concentration of Tris-HCl buffer was 50 mM) was added into the tube. The mixture was vortexed for 30 s and then shaken at rt for 1 h. After 1 h, the reaction mixture was purified directly using flash RP chromatography (column B or C, gradient A); the fractions containing the product were pooled, flash frozen and lyophilized to obtain the unicyclic peptide as a fluffy white powder.

3.6 General procedure for modification of peptides not containing Arg via CuAAC

In a 1.5-mL centrifuge tube, the propargylated peptide (1–2 mg, ~0.6–1.2 $\mu$mol) was dissolved in DMF (0.5 mL) to achieve a final concentration of 1–2.5 mM. To this tube, azide-PEG3-azide (10 equiv, 200 mM stock solution in DMF), biotin-PEG4-alkyne (3.0 equiv, 100 mM stock solution in DMF) or biotin azide (3.0 equiv, 100 mM stock solution in DMF); THPTA (1.67 equiv, 100 mM stock solution in ddH$_2$O); sodium ascorbate (8.3 equiv, 100 mM freshly prepared* stock solution in ddH$_2$O); and copper(II) sulfate (1.5 equiv, 200 mM stock solution in ddH$_2$O) were added in that order. The reaction mixture was shaken for 12 h. After 12 h, the reaction mixture was added to 4 mL of ddH$_2$O and purified by RP-HPLC. The fractions containing the triazole product were pooled, flash frozen and lyophilized to provide the product as a fluffy, white powder.

*The sodium ascorbate stock solution must be prepared freshly just prior to use; all other reagents are stable for prolonged periods (at least one month) at 4 °C in solution.

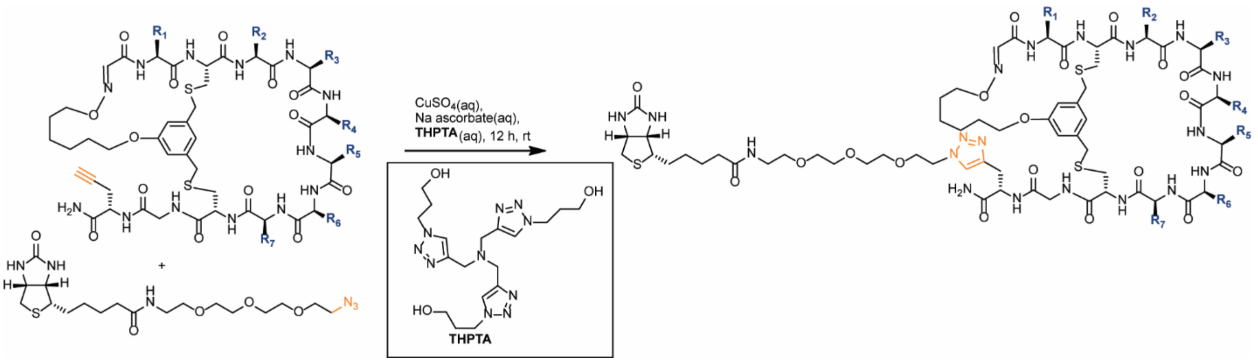

**Figure S4**. Modification of peptides not containing arginine via CuAAC

3.7 General procedure for the modification of peptides containing Arg via CuAAC

In a 1.5-mL centrifuge tube, the propargylated peptide (1–2 mg, ~0.6–1.2 $\mu$mol) was dissolved in DMF (0.5 mL) to achieve a final concentration of 1–2.5 mM. To this tube, biotin-PEG3-azide (3 equiv, 100 mM stock solution in DMF), THPTA (1.67 equiv, 100 mM stock solution in ddH$_2$O), aminoguanidine (10 equiv, 100 mM freshly prepared* stock solution in ddH$_2$O), sodium ascorbate (8.3 equiv, 100 mM freshly prepared* stock solution in ddH$_2$O) and copper(II) sulfate (1.5 equiv, 200 mM stock solution in ddH$_2$O) were added in that order. The reaction mixture was agitated for 12 h. After 12 h, the reaction mixture was added to 4 mL of ddH$_2$O and purified by RP-HPLC. The fractions containing the triazole product were pooled, flash frozen and lyophilized to provide the product as a fluffy, white powder.

*The sodium ascorbate and amino guanidine stock solutions must be prepared freshly just prior to use; all other reagents are stable for prolonged periods (at least one month) at 4 °C in solution.

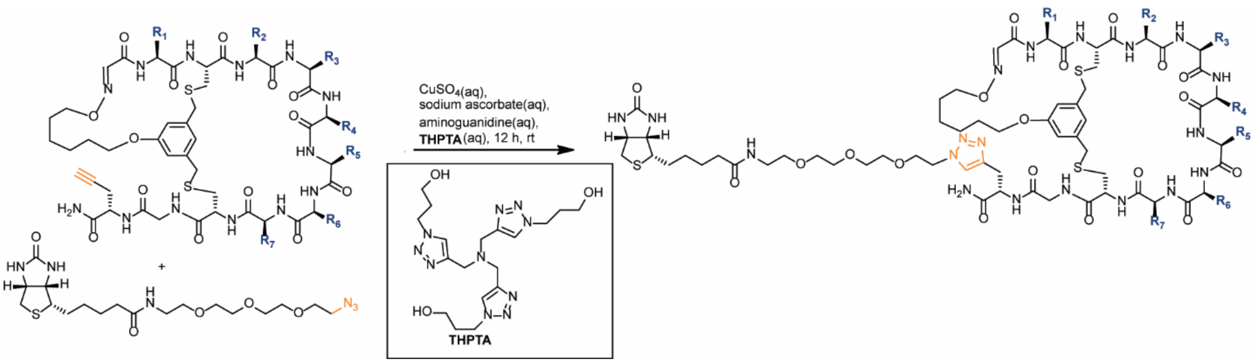

**Figure S5**. Modification of peptides that contain arginine via CuAAC

### 4 Biochemistry

#### 4.1 Protein synthesis

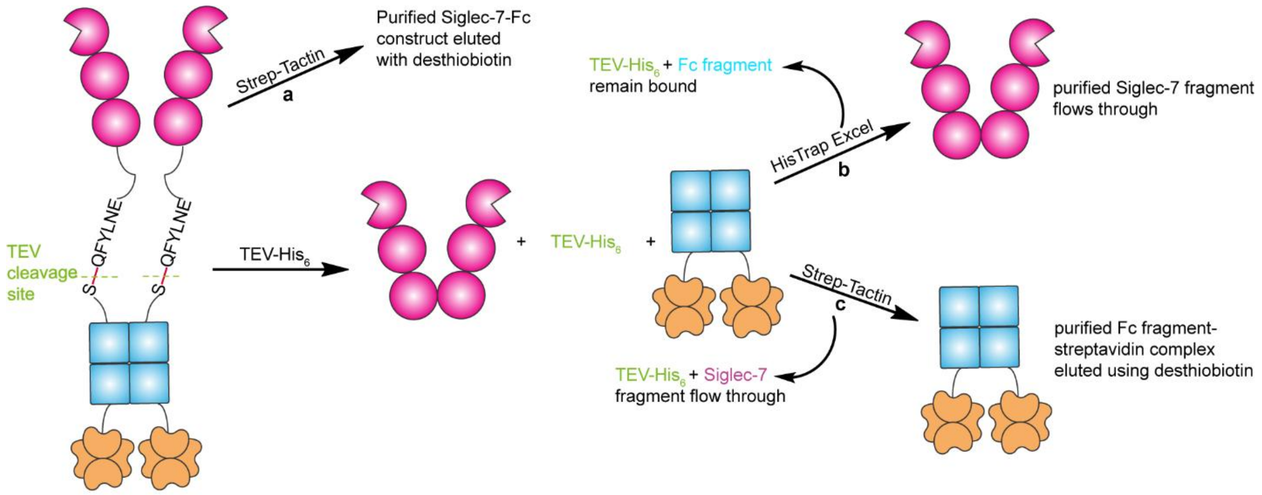

**Figure S6.** Workflow of protein synthesis.

The various chromatographic and biochemical sequences used to isolate a) the purified Siglec-7–Fc fragment construct, b) the purified Siglec-7 fragment and c) the purified Fc fragment–streptavidin construct.

#### 4.2 Expression and purification of Siglec-7–Fc

Siglec-7-Fc was expressed identically to as described previously.[2, 3] The Siglec-7 supernatant was first purified by $Ni^{2+}$ affinity chromatography using a HisTrap Excel (GE LifeSciences) column. After equilibration of the column with 20 column volumes of equilibrium buffer (500 mM NaCl(aq), 20 mM $NaPO_4H_2$(aq), pH 7.4), the supernatant (~250 mL) was loaded. The column was washed with 20 column volumes of wash buffer (500 mM NaCl(aq), 30 mM imidazole(aq), 20 mM $NaPO_4H_2$(aq), pH 7.4). The purified supernatant was subsequently eluted with 20 column volumes of elution buffer (500 mM NaCl(aq), 500 mM imidazole(aq), 20 mM $NaPO_4H_2$(aq), pH 7.4).

The fractions containing Siglec-7-Fc were consolidated and diluted 1:4 with wash buffer; crude Siglec-7-Fc was purified using a Strep-Tactin column thusly. After equilibration with 15 column volumes of wash buffer (100 mM Tris-HCl(aq), 150 mM NaCl(aq), 1 mM EDTA(aq), pH 8.0), the sample (~100 mL) was loaded. The column was then washed with 15 column volumes of wash buffer, and the purified protein sample eluted with 15 column volumes of elution buffer (100 mM Tris-HCl(aq), 150 mM NaCl(aq), 1 mM EDTA(aq), 10 mM desthiobiotin, pH = 8.0). The elution fractions containing purified Siglec-7–Fc were then dialysed against PBS at 4 °C for approximately 48 hours. The dialysed protein was then concentrated using Amicon ultracentrifugal filters (MWCO 30 kDa) to approximately 0.15

mg/mL via nanodrop and then aliquoted in 5 μg increments, frozen and lyophilized. The freeze-dried samples were subsequently assessed for purity via SDS-PAGE; the final concentration of Siglec-7–Fc was determined by the BCA protein or Bradford assay.

##### 4.3 Synthesis and purification of Siglec-7–biotin

The Fc-fragment cleavage portion of this procedure has been described previously,[2] but it is reiterated here with slight modifications as pertinent to this study for clarity.

Purified Siglec-7–Fc was incubated with a tenfold molar excess of TEV protease at 30 °C for 8 h; SDS-PAGE was used to verify that no intact Siglec-7–Fc remained. The reaction mixture was purified by passing it over a HisTrap Excel (GE LifeSciences) column preequilibrated with 15 column volumes of equilibrium buffer (20 mM $Na_2PO_4$(aq), 0.5 M NaCl(aq), pH 7.4); the column was subsequently washed with 5 column volumes of washing buffer (20 mM $Na_2PO_4$(aq), 0.5 M NaCl(aq), 30 mM imidazole(aq), pH 7.4). The flow-through containing Siglec-7–Fc was combined, and to it biotin NHS ester (10 mM in DMSO, 20 equiv) was added; the reaction mixture was then incubated on ice for 2 h, which was sufficiently long enough to ensure complete biotinylation.

The biotinylated protein was then concentrated using an ultracentrifugal device (MWCO 30 kDa) with iterative washes of PBS. The partially concentrated protein was aliquoted, frozen and lyophilized. The freeze-dried samples were subsequently assessed for purity via SDS-PAGE; the final concentration of Siglec-7–biotin was determined by the BCA protein or Bradford assay.

##### 4.4 Isolation of the Fc fragment

Purified Siglec-7–Fc was incubated with a tenfold molar excess of TEV protease at 30 °C for 8 h; SDS-PAGE was used to verify that no intact Siglec-7–Fc remained. The reaction mixture was purified by passing it over a Strep-Tactin column (IBA LifeSciences) pre-equilibrated with 5 column volumes of wash buffer (100 mM Tris-HCl(aq), 150 mM NaCl(aq), 1 mM EDTA(aq), pH 8). After loading, the column was washed with 5 column volumes of the same wash buffer (NOTE: the Siglec-7 fragment and TEV protease flow through); the purified Fc fragment was then eluted with 10 column volumes of elution buffer (100 mM Tris-HCl(aq), 150 mM NaCl(aq), 1 mM EDTA(aq), pH 8, 5 mM desthiobiotin(aq)). The fractions that contained the protein were combined and concentrated down to ~1 mL by Amicon ultracentrifugal filters (Thermo Fisher).

The partially concentrated protein was aliquoted, frozen and lyophilized. The freeze-dried samples were subsequently assessed for purity via SDS-PAGE; the final concentration of Fc–biotin was determined by the BCA protein or Bradford assay.

##### 4.5 Expression and purification of Siglec-9–Fc

Siglec-9–Fc was expressed and purified exactly as described in Sections 4.2 for Siglec-7–Fc.

##### 4.6 Synthesis and purification of Siglec-9–biotin

Siglec-9–biotin was synthesized exactly as described in Section 4.3 for Siglec-7-biotin.

##### 4.7 Panning and deep sequencing

###### 4.7.1 General protocol for modification of the $SXCX_6C$ phage library

The $SXCX_6C$ phage-displayed peptide library was cloned using trinucleotide codon libraries and purified by PEG precipitation; this phage library $SXCX_6C$ was further cleaned up with Triton X-100 and PEG-NaCl precipitation as described previously.[1] The resuspended phage were then dialyzed at 4 °C against 4 L of PBS (50 mM, pH 7.4) for 16 h with two buffer changes at 4 h and 12 h using a 10K MWCO membrane. All incubations steps that included chemical modification were performed by gentle agitation on a slow rotator, as prolonged agitation of phage is detrimental to the infectivity of phage.[4]

###### 4.7.2 Chemical modifications

**Oxidative cleavage:** To a cleaned phage library (300 $\mu$L, >5 × $10^{12}$ pfu/mL), aqueous sodium periodate (3 $\mu$L of a 6 mM solution; final concentration was 60 $\mu$M) was added and incubated on ice in the dark for 8 min. The oxidation was quenched with aqueous methionine (3 $\mu$L of a 50 mM aq solution; final concentration of 0.5 mM) and incubated for 20 min at rt. The phage were purified using a Zeba desalting column (7K MWCO, 0.5 mL, cat# 89882) equilibrated with 1 × PBS.

**Ligation:** To 300 $\mu$L of the oxidized library, TSL-6 (15 $\mu$L of 20 mM TSL-6(aq) and 1% HCl(aq) (15 $\mu$L) were added, and the phage were incubated for 40 min at rt. The reaction mixture was diluted twofold with sodium acetate buffer (10 mM, pH = 5) to increase the percentage of phage passing through a Zeba desalting column equilibrated with sodium acetate buffer (350 $\mu$L, 10 mM, pH = 5). To monitor the progress of the ligation, an AOB chase was used as described previously.[1] Briefly, 5 $\mu$L of the oxidized or 5 $\mu$L of the oxime-ligated phage solutions were combined with 1 mM (5 $\mu$L of 2 mM AOB in 200 mM aniline acetate buffer, pH 4.6) for 1 h. the AOB-modified phage was subsequently diluted 106-fold and captured with streptavidin magnetic beads; the supernatant was titered before and after capture.

**Reduction and bicyclization:** To 100 µL of the purified library, TCEP (1 $\mu$L of 100 mM TCEP in water, final concentration 0.5 mM) was added, and the reaction mixture was incubated for 30 min. The pH was increased to 10 by addition of bicarbonate buffer (25 $\mu$L of 1 M bicarbonate buffer, pH 10) and incubation for 3 h led to cyclization. The modified library was supplemented with PBS (20 $\mu$L of 500 mM PBS, pH 7.4) and purified using a Zeba column prior to storage or panning. To monitor the cyclization reaction, 5 µL of the reaction mixture was sampled at various stages (before and after cyclization) and combined thiol-biotin (BSH) at pH 8.5 (2 $\mu$L of 4 mM BSH in $ddH_2O$), supplemented with Tris-HCl(aq) (5 $\mu$L, 500 mM, pH 8.5) and $ddH_2O$ (38 $\mu$L) and incubated for 3 h. The phage treated with BSH were captured using the biotin-capture assay as described above. Typically, at least 40% of the phage library was successfully bicyclized.

4.7.3 General selection and validation method

4.7.3.1 Panning on a Kingfisher instrument

The protein-immobilized beads from round 2, round 3 and round 4 panning suspensions, and the other reagents were added to a 96-deepwell plate (Thermo Fisher, catalogue # 95040450) as follows:

Row A: protein-coated magnetic beads (transfer-coated beads with buffer into one well, PBS was added up to 1 mL)

Row B: 12-tip deep well magnetic comb (Thermo Fisher, catalogue # 97003500)

Row C: wash buffer (1 mL in each well of 1 x PBS)

Row D: blocking buffer (1 mL in each well of 2% BSA (w/v) in 1 x PBS)

Row E: a solution of TSL-6-modified $SXCX_6C$ libraries after depletion for each round (1 mL in PBS buffer, $10^{10}$ pfu for round 2 and $10^9$ pfu for round 3 and round 4)

Row F: wash buffer (1 mL, 0.1 % Tween-20 (v/v) in PBS for round 2, round 3 and round 4)

Row G: wash buffer (1 mL, 0.1 % Tween-20 (v/v) in PBS for round 3 and round 4)

Row H: wash buffer (1 mL, 0.1 % Tween-20 (v/v) in PBS for round 3 and round 4)

The following steps were performed using a KingFisherTM Duo Prime purification system with a 12-tip magnetic comb to transfer the beads.

The program was as follows:

1) Collect comb from row B.

2) Collect beads from row A on comb.

3) Wash beads in row C for 30 s.

d) Block in row D for 1 h.

e) Bind phage in row E for 1.5 h.

f) Wash beads in row F for 1 min (for round 2, round 3 and round 4).

g) Wash beads in row G for 1 min (for round 3 and round 4)

h) Wash beads in row H for 1 min (for round 3 and round 4)

At the end of the program, the protein-coated beads with phage bound were in the wells in row F or row H. The contents of each well from their respective row was transferred to an Eppendorf tube and processed for library amplification in the next round of panning or Illumina PCR.

##### 4.7.3.2 Panning of purified recombinant Siglec-7 protein

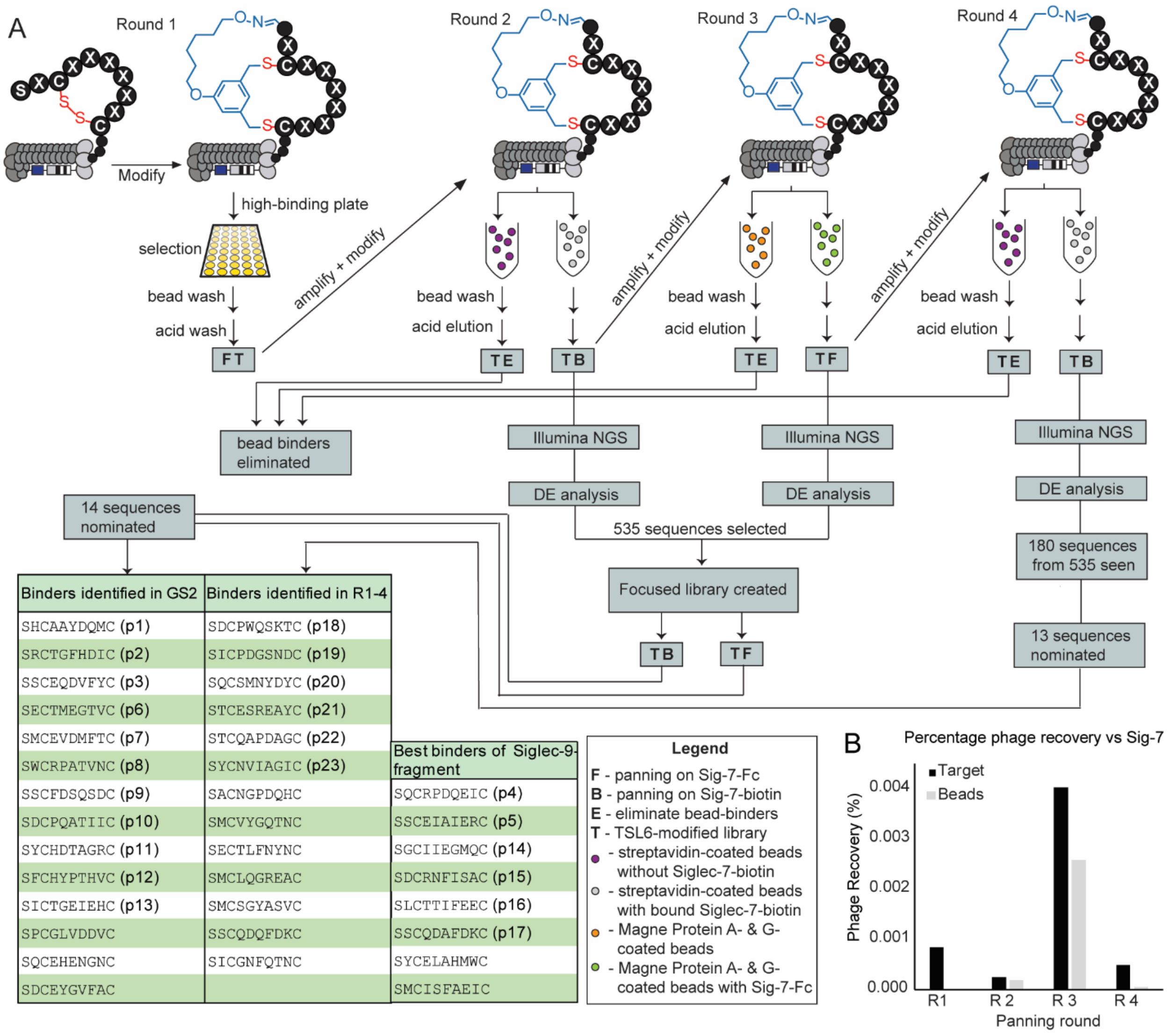

**Figure S7.** Visual flowchart of panning done on Siglec-7.

A) Workflow of the panning experiments completed against Siglec-7-Fc. B) Percentage of the phage recovered through rounds 1–3.

*Round 1 of selection*: 10 $\mu$g of purified Siglec-7–FC in 100 $\mu$L 1 · PBS was added to one well of a Corning high-binding 96-well plate; a total of four wells were used for panning in round 1. The plate was incubated for 12 h at 4 °C to allow the protein to link to the surface of the plate. The coated wells were blocked with 2% BSA in 1 · PBS (200 $\mu$L/well) for 1 h at rt. After removing the blocking buffer, each well received $10^{11}$ pfu of TSL6-modified phage library in 100 $\mu$L of blocking buffer (2% BSA in 1 · PBS); the wells were then incubated at rt for 1 h. After the 1-h incubation, the phage solution is discarded. The non-specifically bound phage particles were removed by washing three times with 1 · PBS buffer containing 0.1% BSA and 0.1% Tween 20. For each of the four wells, the phage particles were eluted by adding 200 $\mu$L of elution buffer (0.2 M glycine-HCl + 1 mg/mL BSA, pH = 2.2) and incubated at rt for 9 min. These elutions were then transferred into a 1.7-mL centrifuge tube containing 30 $\mu$L neutralization buffer (1 M Tris HCl, pH = 9.1). The recovered output phage solution (230 $\mu$L) was amplified for the next round of biopanning and PCR for deep sequencing.

*Round 2 of selection*: Amplified phage recovered from round 1 were modified with TSL6. In a 1.7-mL centrifuge tube, 30 $\mu$L of streptavidin-coated magnetic beads (Streptavidin MagneSphere® Paramagnetic Particles, Promega, catalogue #: Z5481) were incubated with 10 $\mu$g of the purified Siglec-7–biotin protein overnight in 100 $\mu$L of 1× PBS at 4 °C. Eight incubation tubes with protein (target) were prepared for round 2, and another four tubes without protein (control) were used as a negative control. In parallel, to the 1.7-mL centrifuge tube used for depletion, the TSL-6-modified round 2 library was incubated with 100 $\mu$L of the protein-free streptavidin-coated magnetic beads at 4 °C to remove the non-specific bead binding sequences. To the protein-immobilized bead suspension, the depleted phage library ($10^{10}$ pfu for each replicate; there were eight replicates for the target and four for the negative control) and other reagents (see Section 4.3.1) were added to a 96-deepwell plate (Thermo Fisher Scientific, catalogue #: 95040450) for panning in round 2. There was only one wash after incubation with phage in round 2. The washed beads were used for DNA extraction for Illumina PCR or phage elution for phage amplification. For 4 out of the 8 target replicates and 4 control replicates, the washed beads were transferred into individual 1.7-mL centrifuge tubes, with the supernatant removed. Each of the tubes received 30 $\mu$L dd$H_2O$ and were boiled at 95 °C for 10 min, followed by centrifugation at 20 000*g* for 2 min. The supernatant was transferred into a new tube and stored at –20 °C. Two Illumina PCRs were completed for each of these replicates (target and control), and 13.5 $\mu$L of the extractions was used as template for each PCR tube. For the remaining four target replicates, binding phage were eluted from the beads for phage amplification. The beads were incubated with 200 $\mu$L of elution buffer (0.2 M glycine-HCl + 1 mg/mL BSA, pH = 2.2) and incubated at rt for 9 min. These elutions were then transferred into a new 1.7-mL centrifuge tube containing 30 $\mu$L neutralization buffer (1 M Tris HCl, pH = 9.1). The recovered output phage solution (230 $\mu$L) was amplified (200 $\mu$L) with ER2738 bacteria for the next round of biopanning and PCR (two PCR tubes each replicate, 10 $\mu$L elutions for each PCR tube) for deep sequencing.

*Round 3 of selection*: Amplified phage recovered from round 2 were modified with TSL6. Round 3 panning was done with Siglec-7–Fc protein-coated Magne® Protein G beads (Promega, Catalogue #: G7472) following the panning steps for round 2 with the following modifications: the number of input phage particles was reduced to $10^9$ pfu for each replicate, and washing was carried out thrice. The washed beads were treated the same way as round 2.

*Round 4 of selection*: Round 4 panning was done with Siglec-7–biotin coated with Streptavidin MagneSphere® Paramagnetic Particles following the same steps as in round 3. There were 4 replicates for target (Siglec-7–biotin-coated beads) and 4 replicates for control (empty beads). The washed beads were transferred into a 1.7 mL centrifuge tube and phage particles were eluted out by adding 200 $\mu$L of elution buffer (0.2 M glycine-HCl + 1 mg/mL BSA, pH = 2.2) and incubated at rt for 9 min. These tubes were placed on a magnetic rack and the supernatant were transferred into a new 1.7-mL centrifuge tube containing 30 $\mu$L neutralization buffer (1 M Tris HCl, pH = 9.1). All of the recovered output phage solution (230 $\mu$L) was used for phage amplification with ER2738 bacteria. Illumina PCR for deep sequencing was done with the amplified phage library.

4.7.3.3 Panning of purified recombinant Siglec-9 protein

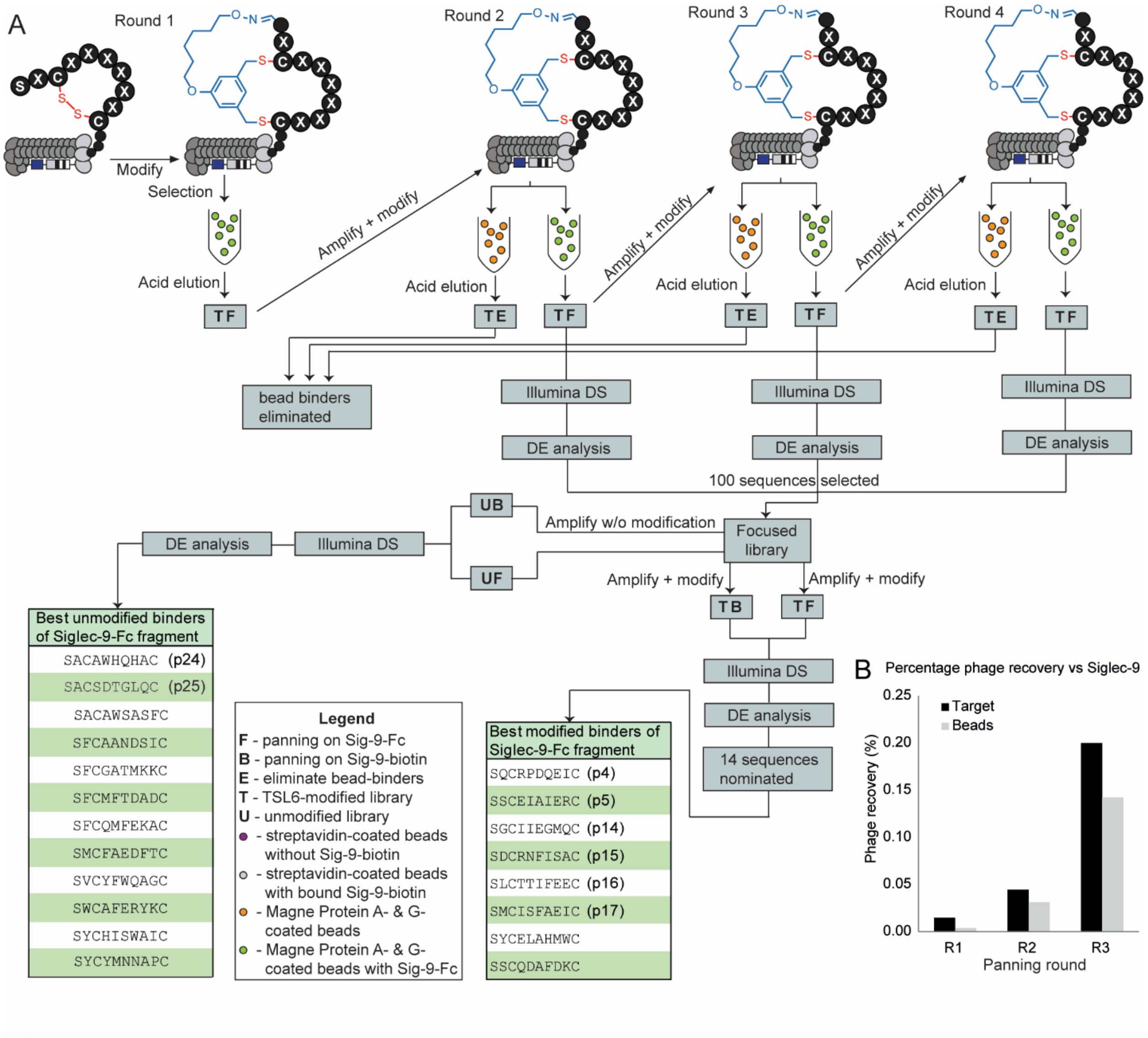

**Figure S8.** Visual flowchart of panning done on Siglec-9

A) Workflow of the panning experiments completed against Siglec-9-Fc. B) Percentage of the phage recovered through rounds 1–3.

*Round 1 of selection*: 10 $\mu$g of purified Siglec-7–Fc in 100 $\mu$L 1 · PBS was added to one well of a Corning high-binding 96-well plate; a total of four wells were used for panning in round 1. The plate was incubated for 12 h at 4 °C to allow the protein to link to the surface of the plate. The coated wells were blocked with 2% BSA in 1 · PBS (200 $\mu$L/well) for 1 h at rt. After removing the blocking buffer, each well received $10^{11}$ pfu of TSL6-modified phage library in 100 $\mu$L of blocking buffer (2% BSA in 1 · PBS); the wells were then incubated at rt for 1 h. After the 1-h incubation, the phage solution is discarded.

The non-specifically bound phage particles were removed by washing three times with 1 · PBS buffer containing 0.1% BSA and 0.1% Tween 20. For each of the four wells, the phage particles were eluted by adding 200 $\mu$L of elution buffer (0.2 M glycine-HCl + 1 mg/mL BSA, pH = 2.2) and incubated at rt for 9 min. These elutions were then transferred into a 1.7-mL centrifuge tube containing 30 $\mu$L neutralization buffer (1 M Tris HCl, pH = 9.1). The recovered output phage solution (230 $\mu$L) was amplified for the next round of biopanning and PCR for deep sequencing.

*Round 2 of selection*: Amplified phage recovered from round 1 were modified with TSL6. In a 1.7-mL centrifuge tube, 30 $\mu$L of streptavidin-coated magnetic beads (Streptavidin MagneSphere® Paramagnetic Particles, Promega, catalogue #: Z5481) were incubated with 10 $\mu$g of the purified Siglec-9–biotin protein overnight in 100 $\mu$L of 1× PBS at 4 °C. Eight incubation tubes with protein (target) were prepared for round 2, and another four tubes without protein (control) were used as a negative control. In parallel, to the 1.7-mL centrifuge tube used for depletion, the TSL-6-modified round 2 library was incubated with 100 $\mu$L of the protein-free streptavidin-coated magnetic beads at 4 °C to remove the non-specific bead binding sequences. To the protein-immobilized bead suspension, the depleted phage library ($10^{10}$ pfu for each replicate; there were eight replicates for the target and four for the negative control) and other reagents (see Section 4.3.1) were added to a 96-deepwell plate (Thermo Fisher Scientific, catalogue #: 95040450) for panning in round 2. There was only one wash after incubation with phage in round 2. The washed beads were used for DNA extraction for Illumina PCR or phage elution for phage amplification. For 4 out of the 8 target replicates and 4 control replicates, the washed beads were transferred into individual 1.7-mL centrifuge tubes, with the supernatant removed. Each of the tubes received 30 $\mu$L dd$H_2O$ and were boiled at 95 °C for 10 min, followed by centrifugation at 20 000*g* for 2 min. The supernatant was transferred into a new tube and stored at –20 °C. Two Illumina PCRs were completed for each of these replicates (target and control), and 13.5 $\mu$L of the extractions was used as template for each PCR tube. For the remaining four target replicates, binding phage were eluted from the beads for phage amplification. The beads were incubated with 200 $\mu$L of elution buffer (0.2 M glycine-HCl + 1 mg/mL BSA, pH = 2.2) and incubated at rt for 9 min. These elutions were then transferred into a new 1.7-mL centrifuge tube containing 30 $\mu$L neutralization buffer (1 M Tris HCl, pH = 9.1). The recovered output phage solution (230 $\mu$L) was amplified (200 $\mu$L) with ER2738 bacteria for the next round of biopanning and PCR (two PCR tubes each replicate, 10 $\mu$L elutions for each PCR tube) for deep sequencing.

*Round 3 of selection*: Amplified phage recovered from round 2 were modified with TSL6. Round 3 panning was done with Siglec-9–Fc protein-coated Magne® Protein G beads (Promega, Catalogue #:

G7472) following the panning steps for round 2 with the following modifications: the number of input phage particles was reduced to $10^9$ pfu for each replicate, and washing was carried out thrice. The washed beads were treated the same way as round 2. A DE analysis plot of round 3 selection is shown against Siglec-7-Fc (Figure 2C) and BSA coated beads (Figure S8a).

*Round 4 of selection*: Round 4 panning was done with Siglec-9–biotin coated with Streptavidin MagneSphere® Paramagnetic Particles following the same steps as in round 3. There were 4 replicates for target (Siglec-7–biotin-coated beads) and 4 replicates for control (empty beads). The washed beads were transferred into a 1.7 mL centrifuge tube and phage particles were eluted out by adding 200 $\mu$L of elution buffer (0.2 M glycine-HCl + 1 mg/mL BSA, pH = 2.2) and incubated at rt for 9 min. These tubes were placed on a magnetic rack and the supernatants were transferred into a new 1.7-mL centrifuge tube containing 30 $\mu$L neutralization buffer (1 M Tris HCl, pH = 9.1). All of the recovered output phage solution (230 $\mu$L) was used for phage amplification with ER2738 bacteria. Illumina PCR for deep sequencing was done with the amplified phage library.

4.7.3.4 Focused library panning

A total of 535 peptide sequences enriched from Siglec 7 protein-panning round 2 and round 3 and 100 peptide sequences enriched from Siglec 9 protein-panning round 1, round 2 and round 3 were selected for the focused library (GS2) panning with Ala scan. (Note: Ala is used to replace positions x SxCxxxxxxC sequentially. For example, SWCRHTNIVC has 8 sequences to be sequentially replaced with alanine.) The DNA oligonucleotides of the peptides with an enzyme adaptor were ordered from Genescript. The oligonucleotides were amplified by PCR with Klenow (cat# M0210S, New England Biolabs) in the presence of maturation primer, dNTPs, and buffer. The amplified PCR products (dsDNA, inserts) were digested with BsaI (cat# R3733S , New England Biolabs) for 5 h. The SDB phage M13KE-BsaI vector was digested with BsaI overnight and phenol extracted and purified on Sephacryl S300 syringe column. (Note: The BsaI cutting sites were carefully designed to let the library insert into the vector. You can see there are two recognition sites of the enzyme and they make different cuts and provide different types of sticky ends. The recognition site on the vector used NGAGACC and the recognition site on oligonucleotides used GGTCTCN; these two will be able to ligate after cut. Otherwise, they will not ligate. The digested inserts were then ligated to the digested vector with T4 DNA ligase (cat# M0202S New England Biolabs) at 3:1 molar excess (inserts:vector). Electroporation of the ligates was done with ecG10prlA4 fresh electrocompetent cells. Conditions were: 2.1 kV, 25 uF, 200 Ω and $\tau$ = 4.5 ms with 0.1 cm cuvettes (cat# 1652083, Bio-Rad). Focused library was Illumina sequenced to confirm that the planned sequences/peptides were present in the cloned library (https://48hd.cloud/file/7462). The cloned GS2 focused library was modified with TSL6.

The modified GS2 library was used for protein panning for Siglec 7-Fc and Siglec 7-biotin. In a 1.7-mL centrifuge tube, 30 $\mu$L of streptavidin-coated magnetic beads (Streptavidin MagneSphere® Paramagnetic Particles, Promega, catalogue #: Z5481) were incubated with 10 $\mu$g of the purified Siglec-7–biotin protein overnight in 100 $\mu$L of 1× PBS at 4 °C. Four incubation tubes with protein (target) were prepared for the focused library panning, and another four tubes without protein (control) were used as negative controls. In parallel, in a 1.7-mL centrifuge tube, 30 $\mu$L of Magne® Protein G magnetic beads (Promega, Catalogue #: G7472) were incubated with 10 $\mu$g of the purified Siglec-7–Fc protein overnight in 100 $\mu$L of 1× PBS at 4 °C. Four incubation tubes with protein (target) were prepared for focused library panning, and another four tubes without protein (control) were used as negative controls.

To the protein-immobilized bead suspension, the focused library ($10^9$ pfu for each replicate; there were four replicates for the target and four for the negative control for each protein type) and other reagents (see Section 4.3.1) were added to a 96-deepwell plate (Thermo Fisher Scientific, catalogue #: 95040450) for panning. There were three washes after incubation with phage in the focused library panning. The washed beads were used for DNA extraction for Illumina PCR. The washed beads were transferred into individual 1.7-mL centrifuge tubes, with the supernatant removed. Each of the tubes received 30 $\mu$L dd$H_2O$ and were boiled at 95 °C for 10 min, followed by centrifugation at 20 000*g* for 2 min. The supernatant was transferred into a new tube and stored at –20 °C before Illumina PCR. Two Illumina PCRs were completed for each of these replicates (target and control), and 13.5 $\mu$L of the extractions was used as template for each Illumina PCR tube for deep sequencing. The streptavidin-coated MagneSphere® Paramagnetic Particles without protein were used as control for the panning; both the experimental and control were replicated in quadruplicate. The panning protocol was the same as in round 3 without the 4 target replicates for phage amplification. The FC scatter plot of the focused library showed the nominated sequences showing higher enrichment for Siglec-7-biotin (Figure S8b).

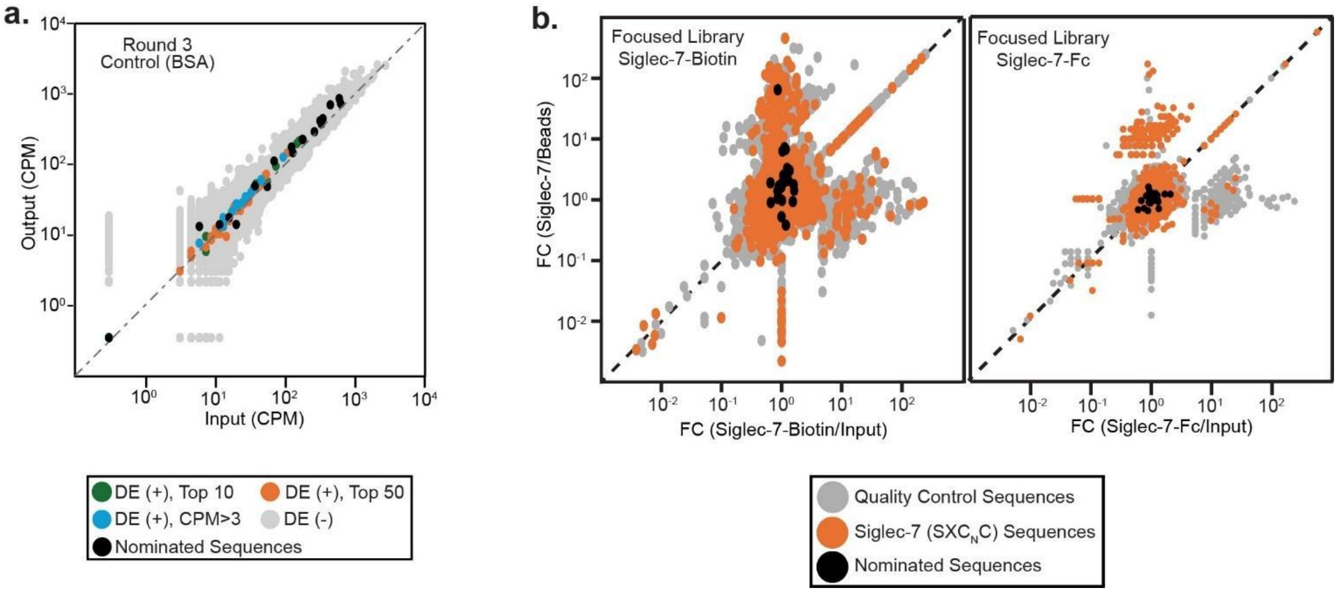

**Figure S9**. FC scatter plot

a) DE analysis plot of round 3 panning against BSA coated beads. b) Scatter plot of FC of Siglec-7 sequences in both Siglec-7-biotin and Siglec-7-Fc against input, with nominated sequences in black.

4.7.3.5 PCR amplification protocol for deep sequencing on Illumina

Either an eluted phage or diluted phage solution (10 $\mu$L) was used as the template for PCR with the total volume being 50 µL; there were two PCR tubes used in each replicate. This 50 µL Illumina PCR system consisted of 5 x Phusion buffer (10 µL), 10 mM dNTPs(aq) (1 µL), Phusion high-fidelity DNA polymerase (0.5 µL, 2U/µL NEB, catalogue # M0530S), forward primer (3'-CAAGCAGAAGACGGCATACGAGATCGGTCTCGGCATTCCTGCTGAACCGCTCTTCCGATCTXXXXTTGGAGATTTTCAACGTG-5' (10 µM, 2.5 µL), reverse primer (3'-AATGATACGGCGACCACCGAGATCTACACTCTTTCCCTACACGACGCTCTTCCGATCTXXXXACAGTTTCGGCCGA-5'), template solution (10 µM, 2.5 µL) and nuclease-free water (23.5 µL). Thermocycling was preformed using the following settings:

1) 98 °C for 3 min
2) 98 °C for 10 s
3) 50 °C for 20 s
4) 72 °C for 30 s
5) repeat steps 2–4 10 times
6) 98 °C for 10 s
7) 65 °C for 20 s

8) 72 °C for 30 s

9) repeat steps 6–8 20 times

10) 72 °C for 5 min

11) 4 °C indefinitely

##### 4.7.4 Illumina deep sequencing

Illumina deep sequencing was completed with the support of MBSU at the University of Alberta. The products were provided by PCR as described in Section 4.7.3.5 with one exception: in the amplification of the libraries before panning (input), the volume of the template (phage solution) was 1 μL ($10^9$ pfu). All Illumina PCR products were quantified by 2% (w/v) agarose gel in tris-borate-EDTA buffer at 100 V for ~35 min using a low molecular weight DNA ladder as standard (NEB, catalogue # N3233S). The PCR products that contained different indexing barcodes (the four XXXX DNA barcodes used for separation of different PCR tubes) were pooled, yielding 20 ng of each product in the mixture. The mixture was purified with 2% (w/v) agarose gel, the cut-out bands were purified with a Monarch® DNA gel extraction kit (New England Biolabs, Catalogue # T1020S), quantified by QuBit and sequenced using the Illumina NextSeq paired-end 500/550 high-output kit v2.5 (2x75 Cycles). The data were automatically uploaded to BaseSpace™ Sequence Hub. Processing of the data was carried out as described .

###### 4.7.4.1 Processing of the Illumina data

The compressed FASTQ files from Illumina were downloaded from BaseSpace™ Sequence Hub. The files were converted into tables of DNA sequences and their counts per experiment. Briefly, FASTQ files were parsed based on unique multiplexing barcodes (XXXX in the primer sequences) within the reads; any reads that contained a low-quality score were discarded. Mapping the forward (F) and reverse (R) barcoding regions, mapping of the F and R priming regions allowing no more than one base substitution each and F-R read alignment allowing no mismatches between F and R reads yielded DNA sequences located between the priming regions.

The files with DNA reads, peptide sequences, raw counts, and mapped peptide modifications were uploaded to http://48hd.cloud/ server. Each experiment has a unique alphanumeric name (e.g., 20191002-297TSpxBI-ZO) and a unique static URL.

**Table S2**. Illumina data for rounds 1–4 against Siglec-7

| | Unmodified | TSL-6-modified | Depleted | Target output | Control output |
|---|---|---|---|---|---|
| | 20191002-296OOooOO-ZO | 20191002-297TSooBI-ZO | 20191002-297TSooOO-ZO | 20191002-297TSpxBI-ZO | N/A |
| | 20191002-298OOooOO-ZO | 20191002-298TSooOO-ZO | 20191002-299TSooOO-ZO | 20191002-299TSpxBD-ZO | 20191002-299TSpxMK-ZO |
| | 20191002-300OOooOO-ZO | 20191002-300TSooOO-ZO | 20191002-301TSooOO-ZO | 20191002-301TSpxAL-ZO | 20191002-301TSpxBS-ZO |
| | 20191107-302OOooOO-ZO | 20191107-302TSooOO-ZO | 20191107-303TSooOO-ZO | 20191107-303TSpxOO-ZO | 20191107-303TSsbOO-ZO |

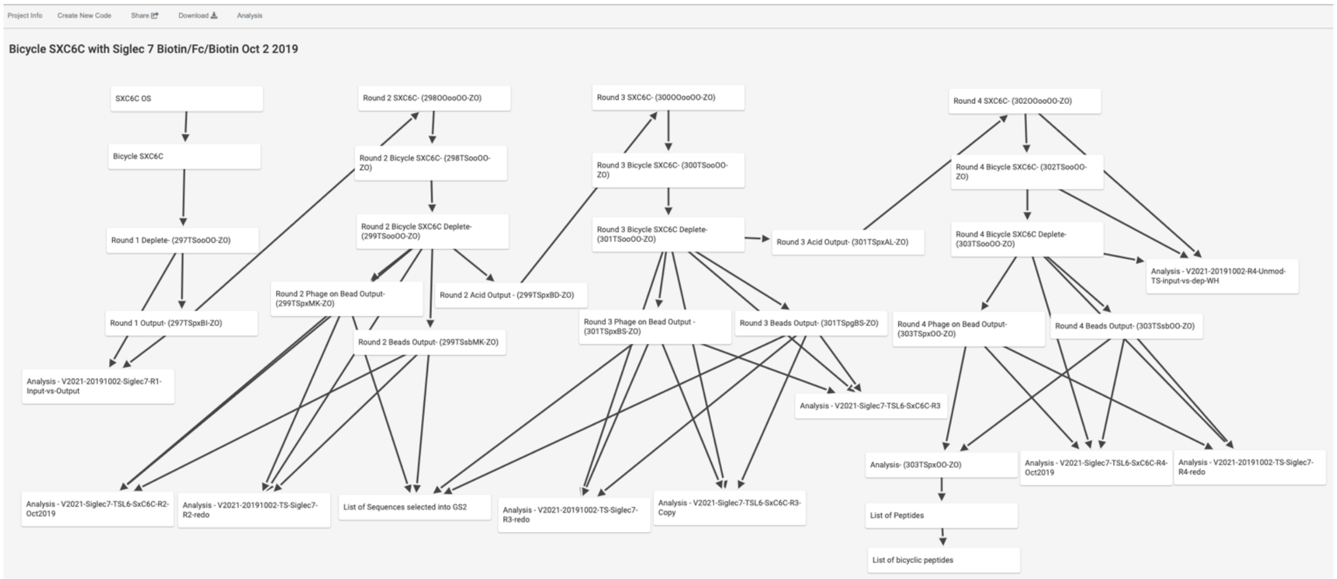

**Figure S10**. Screenshot of NGS flowchart for Siglec-7

Describing various experiments and analyses performed using multiple rounds of selection starting from a Naive phage displayed library. Boxes describe either a naïve or selected library produced as a result of specific experiment and an NGS sequencing dataset associated with each library. Boxes marked as "analysis" contain the results of the differential enrichment (DE) analyses performed by comparing several NSG sequencing samples. Boxes marked as "list of [..]" contain lists of molecules/sequences nominated for further evaluation. The flow chart is an interactive web object hosted on the secure server at https://48hd.cloud/flowchart/edit/81# (to be made public at the time of publication)

**Table S3**. Illumina data for the TSL6-modified focused library against Siglec-7

| | Unmodified | TSL-6-modified | Depleted | Target output | Control output |
|---|---|---|---|---|---|
| | 20200129-366TSnaFO-ZO | 20200129-366TSooOO-ZO | 20200129-366TSpgFO-ZO | 20200129-366TSsbFO-ZO | 20200129-366TSxdFO-ZO |

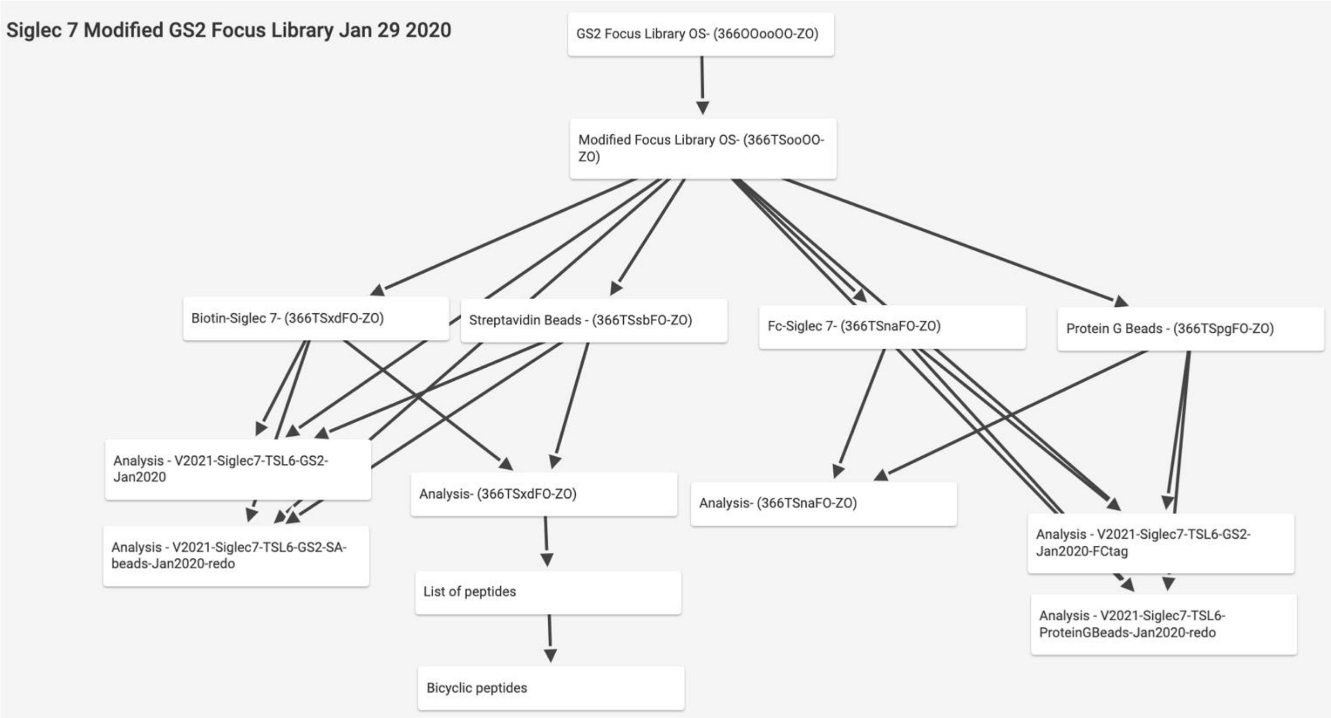

**Figure S11.** Screenshot of NGS flowchart for Siglec-7 GS2 focus library

Describing various experiments and analyses performed using focused phage displayed library. Boxes in the first three layers describe either a naïve or selected library produced as a result of specific experiment and an NGS sequencing dataset associated with each library. Boxes marked as "analysis" contain the results of the differential enrichment (DE) analyses performed by comparing several NSG sequencing samples: The flow chart is hosted on the secure server at https://48hd.cloud/flowchart/edit/140 (to be made public at the time of publication)

**Table S4**. Illumina data for rounds 1–4 against Siglec-9

| | Unmodified | TSL-6-modified | Depleted | Target output | Control output |
|---|---|---|---|---|---|
| | 20190430-262OOlqJK-ZO | | | | |
| | 20190430-268OOooOO-ZO | 20190815-283TSooOO-ZO | | | |
| | 20190703-270OOooOO-ZO | | | | |
| | 20190703-272OOooOO-ZO | | | | |

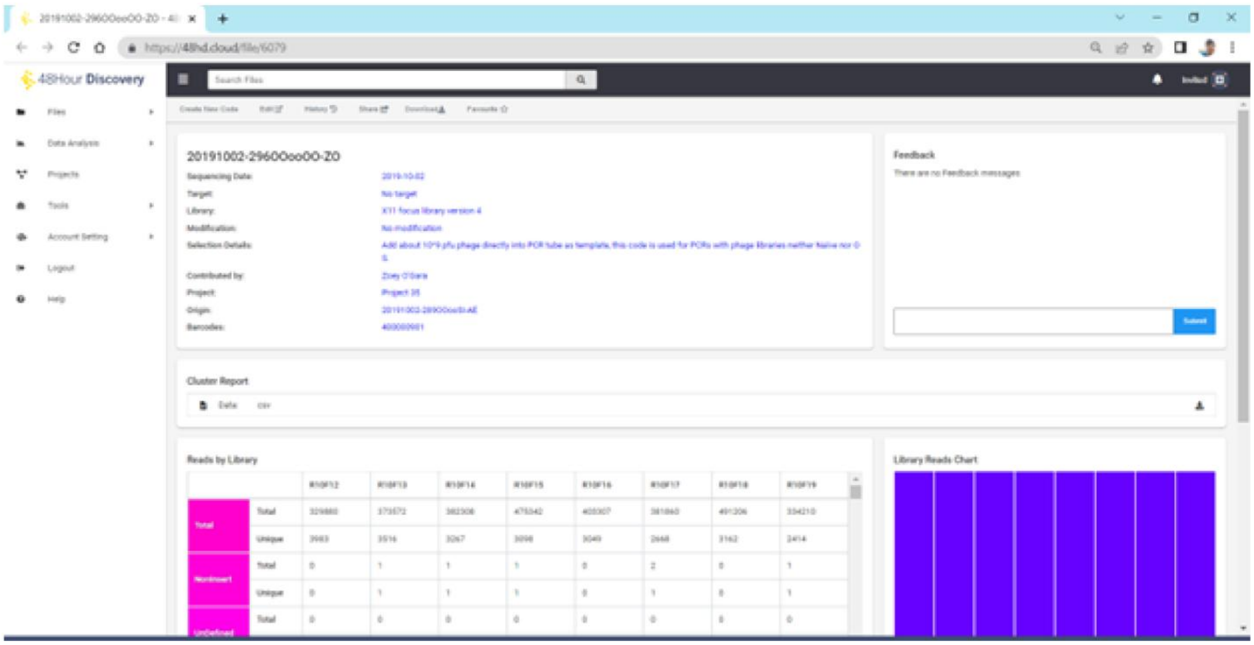

**Figure S12**. Screenshot of the representative data entry web address.

Links to differential analysis report (https://48hd.cloud/file/6079). Both the next-generation sequencing data and analysis reports have been included in supplementary data

#### 4.8 NMR analysis of **46e**

##### 4.8.1 General workflow of **46e**

NMR spectra of the cyclic peptide were recorded on Bruker 800 MHz Avance II spectrometer equipped with a cryo-probes in 90:10 $H_2O$:$D_2O$ at 300 K. Unfortunately, the amide NH protons displayed fast exchange and were not visible in the NMR spectrum. Thus, the spectra were recorded in $D_2O$ using a concentration of 0.5 mM, where only the non-exchangeable protons were observed. Standard homonuclear 2D-NMR experiments were then carried out to assign the amino acid types depending on the spin systems. Thus, TOCSY and NOESY experiments were performed in the phase-sensitive mode using the TPPI method for quadrature detection in F1. Typically, a data matrix of 512 × 2K points was used to digitize a spectral width of 8000 Hz. Before Fourier transformation, zero filling was used in F1 to expand the data to 2K × 2K followed by automated baseline- and phase correction. 2D-TOCSY experiments were acquired with 16 scans per increment, 30 and 80 ms mixing time and a relaxation delay of 1.0 s. NOESY experiments were acquired employing 64 scans per increment, a relaxation delay of 1 s and mixing times of 200 and 300 ms.

2D-HSQC spectra in $D_2O$ were also acquired. A data matrix of 2K*1K was used to digitize a spectral width of 6000 Hz in F2 and 15000 Hz in F1. 32 scans per increment were employed with a relaxation delay of 1 s and a delay corresponding to a J value of 145 Hz.

STD-NMR (Bruker pulse sequence stddiffesgp) experiments[5] were acquired (300K with 1728 scans and 64 K data points) at 800 MHz and 300 K for the peptide in the presence of Siglec7d1d3Fc, its R124A mutant, or Siglec7d1, always at 25uM concentration. The molar ratio was 25:1 peptide:Siglec7 in all cases. An excitation sculpting module with gradients was used to minimize the water signal. Selective saturation of the protein resonances (on-resonance spectrum) was performed by irradiating at d 0.2 ppm using a series of 40 Eburp2.1000-shaped 90° pulses (50 ms) for a total saturation time of 2 s, using a relaxation delay of 4 s. For the reference spectrum (*off- resonance*), the samples were irradiated at d 100 ppm. Control experiments for the isolated Siglec-7 domains and for the free peptide were acquired using the same experimental conditions and using the same experimental setup. The blank STD experiment recorded for the peptide alone did not show any STD signal, while the minor STD intensities observed for the Siglec7 variants were subtracted from those obtained for their corresponding complexes with the peptide. No STD responses were observed for the peptide:Siglec7d1 mixture, strongly suggesting that the peptide does not interact with the isolated Siglec7d1 domain, at least under these experimental conditions.

1.1.1 2D DOSY spectrum

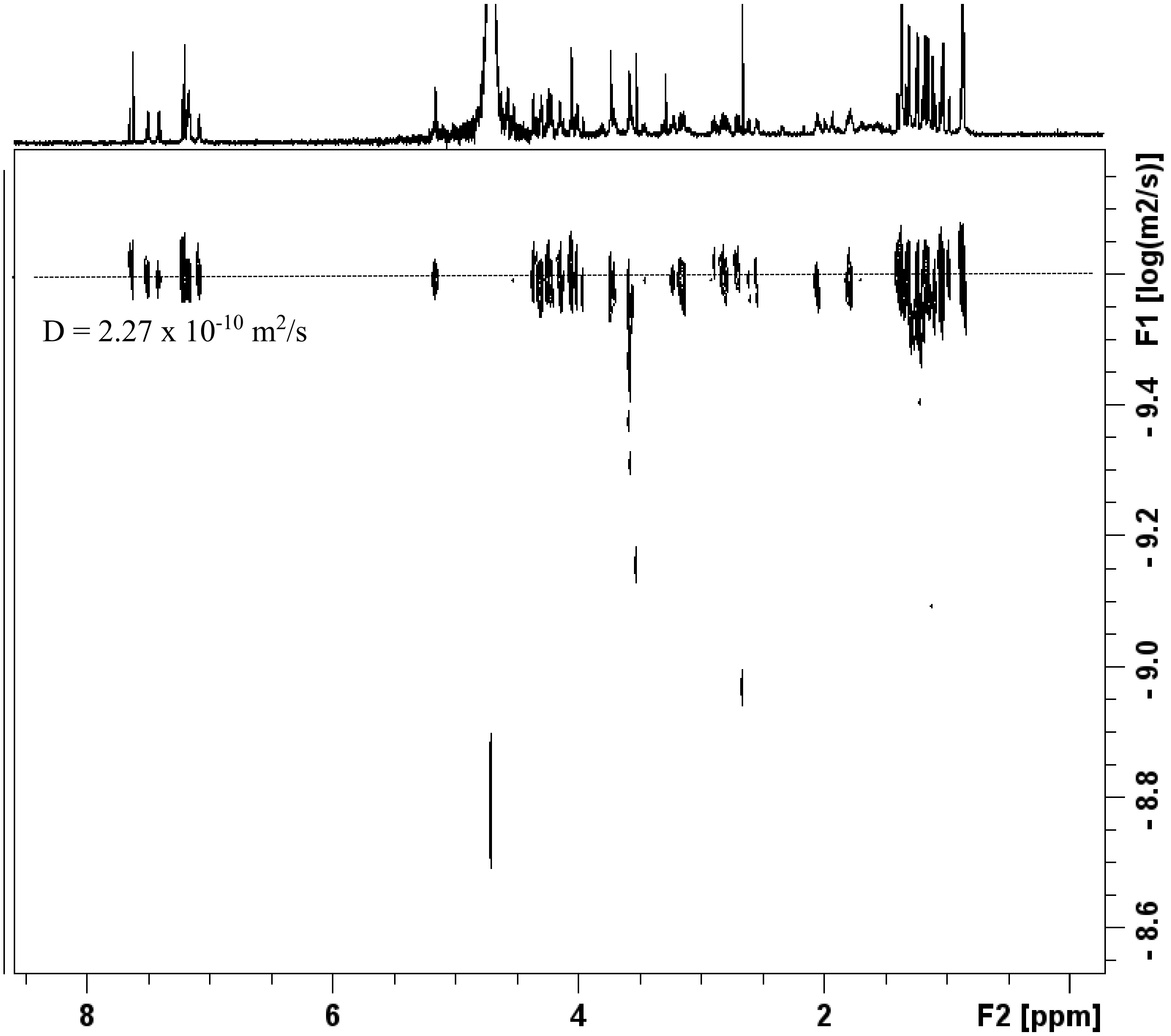

**Figure S13.** 2D DOSY spectrum recorded for the bicyclic peptide in $D_2O$.

The estimated diffusion coefficients for different distinguishable protons of the two species are basically identical (ca. 2.27 x 10 $^{-10}$ $m^2/s$). This result suggests that both species display very similar molecular weight sand shapes.

The possibility of the presence of two different molecules was assessed by using diffusion ordered (DOSY[6] NMR) experiments (Figure S13). However, the observed diffusion coefficient was identical for all the NMR signals, strongly suggesting that they all belong to the same molecular entity.

4.8.2 Variable temperature NMR

The analysis of variable temperature NMR experiments showed significant changes in the shape of the NMR signals. Coalescence of peaks could be observed for **46e** at temperatures exceeding 80 ºC, highlighting the presence of rotamers (Figure S14). A similar trend was also observed with a different parental sequence 57c, showing coalescence at temperatures exceeding 80 ºC (Figure S15).

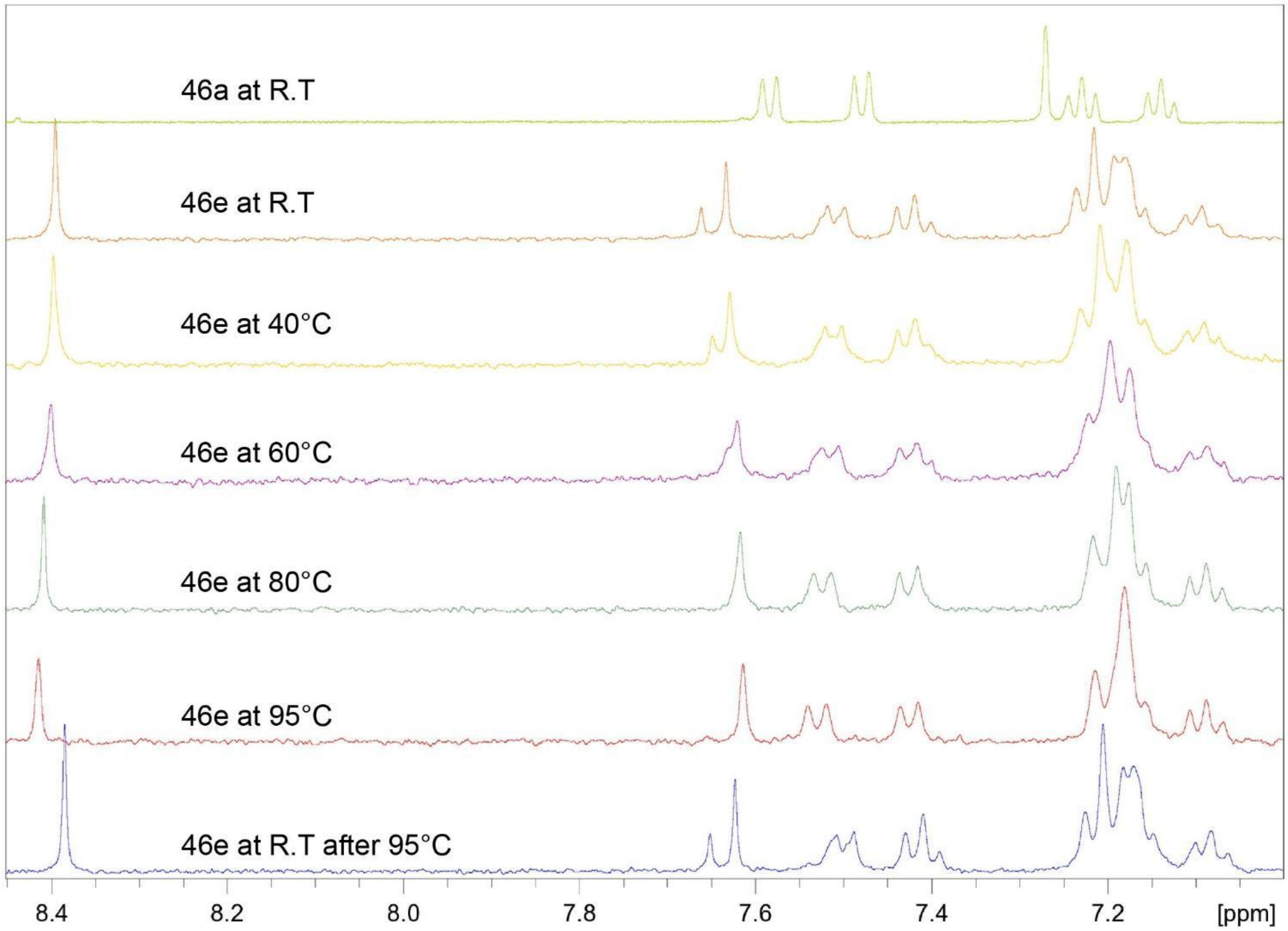

**Figure S14.** Variable temperature NMR of **46e** at various temperature points.

Coalescence of peaks at 7.7-7.6, 7.4, and 7.2-7.1 ppm could be observed at 95 ºC. After heating to such temperatures and cooling back to R.T we can observe an almost identical spectra to R.T highlighting the stability of **46e**.

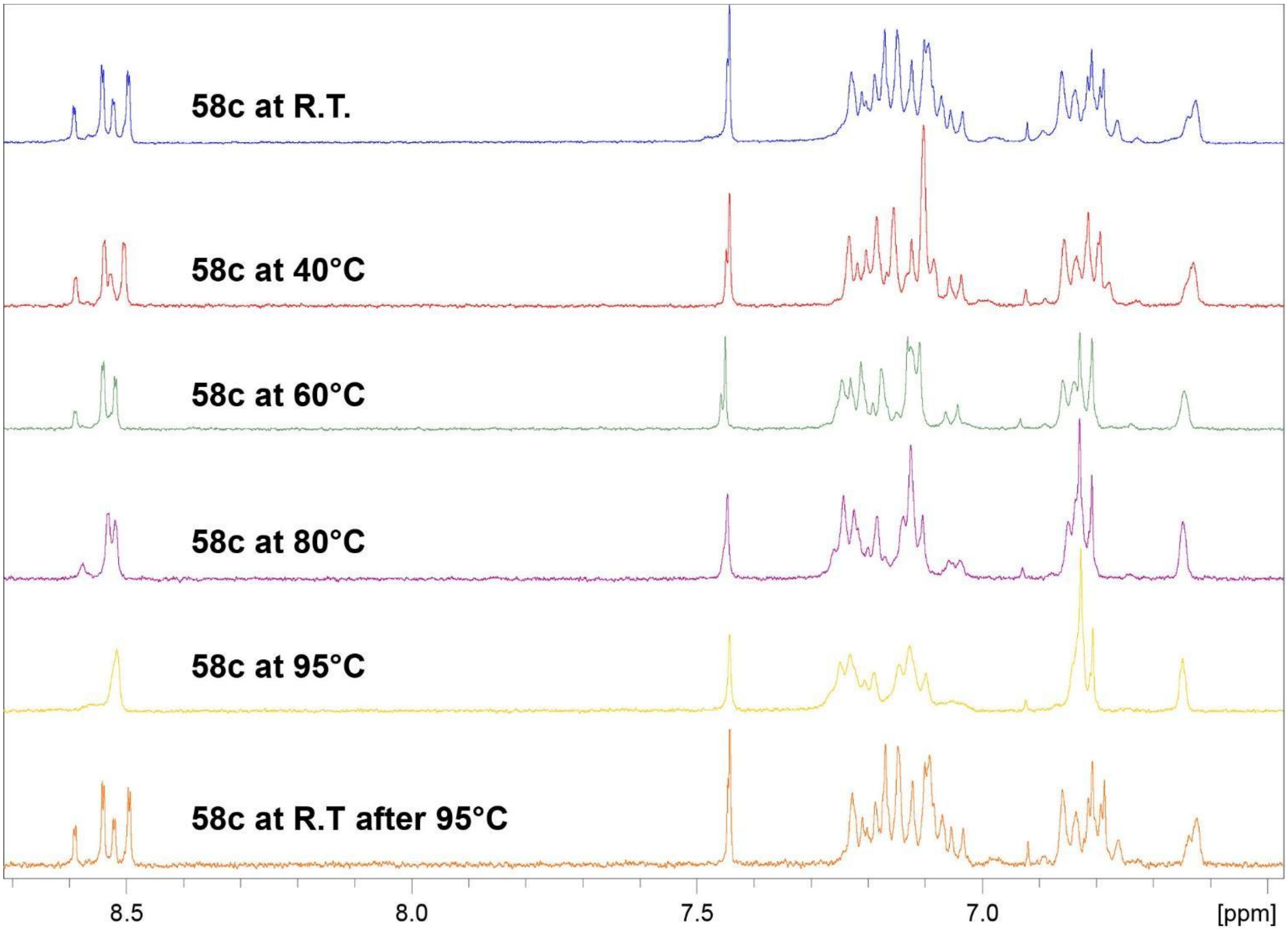

**Figure S15.** Variable temperature NMR of **58c** at various temperature points.

Coalescence of peaks at 8.6-8.5, 7.44, 7.23-7.0, 6.87-6.77, and 6.62 ppm could be observed at 95 ºC. After heating to such temperatures and cooling back to R.T we can observe an almost identical spectra to R.T highlighting the stability of **58c**.

##### 4.8.3 NOESY analysis

Closer inspection of the NOE spectrum revealed inter-residue Pro(d)/Arg(a) cross peaks (Figure 5b) for the major species, consistent with the presence of a trans-Pro residue, since only in this geometry are the two protons in close spatial proximity. The alternative Pro(d) protons of the minor species did not show this NOE and were shifted to higher field relative to those of the major species, as typically reported for cis-Pro rotamers in small peptides. Given the aromatic moiety bridging the two Cys residues, the energy barrier for cis–trans isomerization is predicted to be very high and cannot be overcome under the NMR experimental conditions. To further investigate this, a molecular modeling protocol was implemented. Using the Macromodel suite, two initial structures of the cyclic peptide (cis-Pro and trans-Pro conformers) were generated and subjected to a Monte Carlo search. In both cases, no interconversion between cis and trans states was observed; instead, the simulations produced families of structures reflecting variations in torsional angles at flexible linkages. The global minima identified for the cis- and trans-Pro rotamers were superimposed and are displayed in Figure 5d.

##### 4.8.4 Expression of Siglec-7 for STD-NMR

The pET-43.1(a) plasmid coding Siglec-7 (UniprotKB Q9Y286) V-Ig set domain (amino acid residues 19−148) (Siglec-$7_{d1}$) containing C41S mutation and 6×His tag, was synthesized and purchased from GenScript. The expression and purification in Rosetta-gami B (DE3) *E.coli* competent cells (Novagen) was conducted as described elsewhere[7].

The full-length extracellular domain (ECD) of Siglec-$7_{d1d3}$ (residues 19-357) and the mutant R124A, were fused to human IgG1 Fc region (UniprotKB P01857, residues 99–330) and with a C terminal 6x His tag, were subcloned between XbaI and AfeI restriction sites into pcDNA 3.4 (Invitrogen) and codon optimized for expression in human cells. Plasmids were transiently transfected into HEK293F (Thermo Fisher Scientific) suspension cells with the transfection reagent FectoPRO (Polyplus Transfections). Cells were incubated at 37 °C, 130 rpm, 8% CO2 and 70 % humidity for 6–7 days. After this period, cells were harvested by centrifugation at 5000× g for 20 min, and supernatants were retained and filtered using a 0.45 µm Steritop filter (EMD Millipore). Supernatants were passed through a HisTrap Ni-NTA column (GE Healthcare) and eluted in 20 mM Tris pH 8.0, 300 mM NaCl buffer with an increasing gradient of imidazole (up to 500 mM). The fractions containing Siglec-7-Fc were pooled and separated on a Superdex 200 Increase size exclusion column (GE Healthcare).

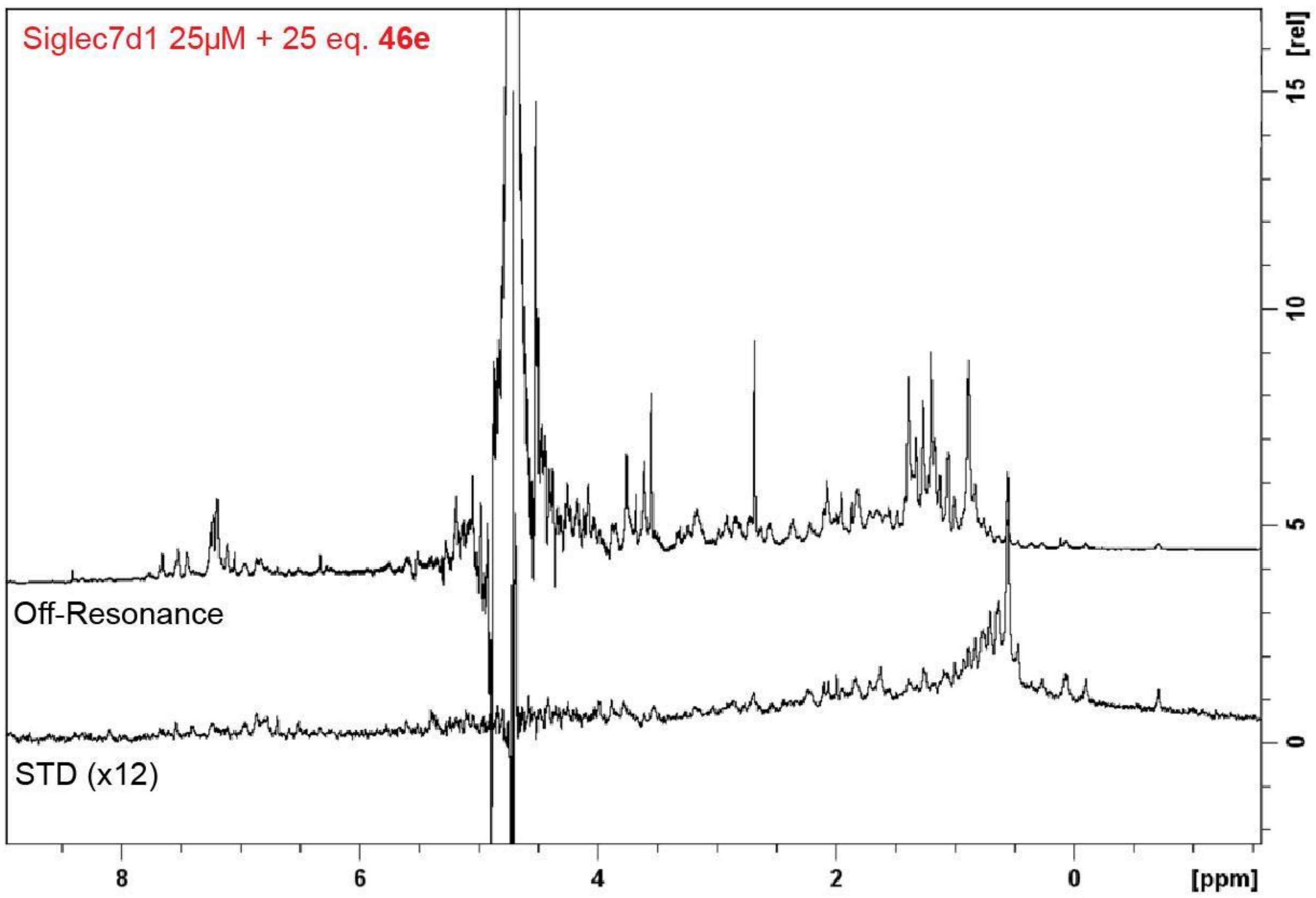

**Figure S16.** $^{1}$H STD NMR of isolated V-Ig set domain (Siglec-7d$_1$)

Mixture of the single isolated V-Ig set domain (Siglec-7d$_1$) and the bicyclic peptide in a 1:25 molar ratio. No STD responses of the peptide were observed. Only STD signals of the protein are present. This result indicates that the peptide does not interact with this domain, but with another domain at the Siglec-7$_{d1d3}$ construct.

4.8.5 Computational modelling

A Low-Mode Conformational Search (LMOD[8]) consisting of 100,000 Monte Carlo steps was conducted using Schrödinger Macromodel software[9] to explore all the possible conformers of the peptide, starting from either the cis- or trans-rotamers of the Pro residue. This method has proved to be efficient at locating low-energy minima on the conformational energy surface of both cyclic and acyclic molecules[8]. The lowest energy conformer in each case was analyzed. Each Monte Carlo cycle begins from the preceding structure only if its energy is in the window of 10 kcal mol$^{-1}$. The force field (FF) employed for the conformational search was OPLS3e. Water was used as solvent with the GB/SA (Generalized Born Surface Area) solvation model, whereas the truncated Newton (TNCG) method was used for minimization.

#### 4.9 Machine learning predications

##### 4.9.1 Prediction of new classes of binders

Classical funneling from multi-round screening to focused libraries and SAR analysis of ~80 synthetic compounds yielded productive leads but confined exploration to chemical space near **8c** peptides. To test whether new Siglec-7 inhibitors could be identified, we applied machine learning (ML) to our NGS datasets. Prior studies from Pentelute[10], Keating[11], He[12], Suga[13], Bowers[14], and even our own lab[15] have shown that ML models can be trained directly on amino acid (AA) strings from NGS data without embedding unnatural chemistries, as long as such entities are consistently present. Following these precedents, we embedded AA strings from rounds 2–4 of selection and labeled positives (class = 1) using logFC thresholds (>1 or >2), with negatives (class = 0) sampled from the pre-selection library. Equal numbers of positives and negatives were used to train classifier models (Koliber AI software) across a range of embeddings, evaluated via cross-validation.

Models trained on pooled rounds 2–4 performed best (AUC ~0.64), while round 4 alone was least effective. Enriched peptides were rescored via leave-one-peptide-out validation, and ranked by differential Siglec-7 vs control scores. From the top 20 predictions, 10 macrocyclic bicycles were synthesized. Eight showed measurable activity ($IC_{50}$ = 14–1000 μM), including two (**51c**, **54c**) with potency (~14 μM) comparable to the best **8c** derivatives, while only two sequences were inactive. Compared to panning-derived sequences, the ML-predicted set displayed a narrower $IC_{50}$ distribution centered around the median potency of experimentally enriched binders. Thus, while higher-affinity inhibitors were not identified, ML successfully extended the landscape of validated Siglec-7 binders beyond conventional hits, highlighting the value of ML-guided prioritization as a complement to experimental screening.

#### 5 Measurements

##### 5.1 General procedure for SPR

All experiments were done on a commercial SPR instrument (Biacore 3000, Biacore International). The amine coupling kits were purchased from Biacore International. The peptide solutions were preprepared in HBS-EP buffer. The highest concentration prepared was 50 $\mu$M; twofold dilutions were carried out until a final concentration of 3.1 $\mu$M was reached (five dilutions: 50 $\mu$M, 25 $\mu$M, 12.5 $\mu$M, 6.2 $\mu$M and 3.1 $\mu$M). These solutions were transferred into SPR tubes. **Note:** For initial quick screening, only three concentrations (50 $\mu$M, 25 $\mu$M and 12.5 $\mu$M were used). Simultaneously, 200 $\mu$L solutions of Siglec-7–Fc–biotin for the SA chip solution at 10 $\mu$g/mL in HBS-EP buffer were prepared.

New chips were washed with HBS-EP buffer before any immobilization was done. The Siglec-Fc-biotin protein was then immobilized onto the chip. There were four FCs (FC1–4) on one chip; usually, FC1 was kept empty as control. Thus, immobilization of Siglec 7-Fc-biotin occurred on FC2, enabling another protein of interest to be immobilized on FC3 if required. All immobilized FCs, including FC1, were blocked with 10 $\mu$M biotin at a flowrate of 10 $\mu$L/min over 30 s. The system was washed again with HBS-EP buffer for 20 min at 10 $\mu$L/min. One NaOH (10 mM, 10 $\mu$L/min, 30 s) injection was done before the real sample injections, followed by a 30 min HBS-EP buffer wash. With the protein immobilized onto the chip, the Customized Application Wizard was employed to setup the peptide injections. Glycine-HCl buffer (10 mM, pH 3.0) was used for regeneration. Briefly, our settings were flow speed were 10 $\mu$L/min; time before injection, 3 min; injection volume, 20 $\mu$L; time after injection, 5 min; regeneration once for 30 s; wash after regeneration for 5 min. All peptides were injected from low concentration to high concentration. After the run, a global fitting for each peptide concentration was obtained separately with the BIAevaluation Software. After a $K_D$ value for each concentration could be determined, the average of all $K_D$ values at all of the concentrations was reported as the final $K_D$. The fitted curves were exported separately as .txt files. The SPR curves and the fitting curves were combined using GraphPad Prism 8.

##### 5.2 General procedure for the ELISAs

Lyophilized GD3 (20 ng, 5 µL from 4 $\mu$g/µL stock) was dissolved in 100% ethanol (2.8 mL), and 50 $\mu$L of this solution were added to a 96-well ELISA plate to achieve a final concentration of 250 pmol/well. As a negative control, 50 µL of 100% ethanol were added instead. The ethanol was subsequently evaporated at room temperature over 12 h or at 37 °C for 2 h. The plate was washed five times with 1x PBS and tamped dry.

BSA solution (5% in PBS, 200 µL) was added to each well; the plate was left to stand for 1 h. During this time, a 3 $\mu$g/mL Siglec Fc solution in PBS was prepared; then, a dilute stock of HRP (Strep-Tactin® conjugate, IBA life science, Cat#: 2-1502-001) was prepared and added into the Siglec Fc solution to achieve a 1:5000 dilution. This mixture was incubated at r.t. for 30 min in the absence of light. The contents of the 96-well ELISA plate were discarded, and the wells washed with 1· PBS five times and tamped dry. Our macrocyclic stock solutions were prepared in 1· PBS at a starting concentration of 600 µM, diluting threefold to a final concentration of 0.09 $\mu$M (nine dilution sequences). Each peptide concentration was added to the wells in triplicate. To do this, (3 · 50 + 10) $\mu$L of the (Siglec-Fc + HRP) and (3 · 10 + 5) µL of each concentration of stock peptide separately were mixed. To achieve a starting final macrocyclic concentration, we commenced at 100 $\mu$M and ended at of 0.03 $\mu$M (nine dilutions). For the control wells (positive and negative), (3 · 10 +5) $\mu$L 1x PBS instead of peptide solution were used. The mixture of Siglec-Fc protein, HRP and macrocycle was incubated at rt for 15 min. To each analyzed well, 60 µL of the mixture were transferred, and the plate was left to stand further at rt for 2 h. 1-Step™ Ultra TMB-ELISA Substrate Solution (Thermo Scientific, Cat#: PI34028) were left to warm at r.t. during this final incubation. After 2 h, the contents of the plate were discarded, and the plate was washed five times with 1 x PBS and tamped dry.

To the wells, 75 $\mu$L of the prewarmed TMB-ELISA substrate solution were added. The reaction progress was gauged by the eye, and typically took 5 to 15 min to complete; after the blue colour no longer seemed to increase in intensity, the reaction in each well was quenched via the addition of 75 $\mu$L of 1 M $H_3PO_4$(aq). The absorbance in each well was measured using an Agilent Cytation 5 plate reader (Agilent Technologies Inc., Mississauga, ON, Canada) at 450 nm. The data were analyzed using GraphPad Prism 8.

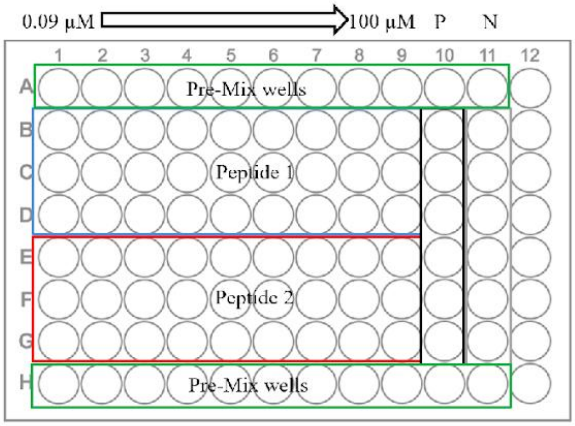

**Figure S17.** Visual representation of 96-well plate analysis for ELISA.

#### 5.3 General procedure for phage recovery assay

##### 5.3.1 Phage recovery on plate

Individual phage constructs and their concentrations were first verified through quantitative polymerase chain reaction (qPCR) using a calibration curve made of phages at known concentrations. Then the individual phage constructs are diluted as needed to reach ~$10^8$ Plaque-Forming Units (PFU) as a working input. To plates previously coated with target protein, and blocked with a blocking agent such as BSA, was added individual phage constructs (test, positive control, or negative control) and left to incubate for an hour at room temperature. The wells were then washed at minimum three times using 1x HBS + 0.1% Tween-20 to remove any unbound phage. After which the bound phage would be eluted using 0.2 M glycine-HCl (9 min at r.t.). The resulting supernatant concentration can then be verified through qPCR and compared against the input to get the recovery percentage (Figure S18).

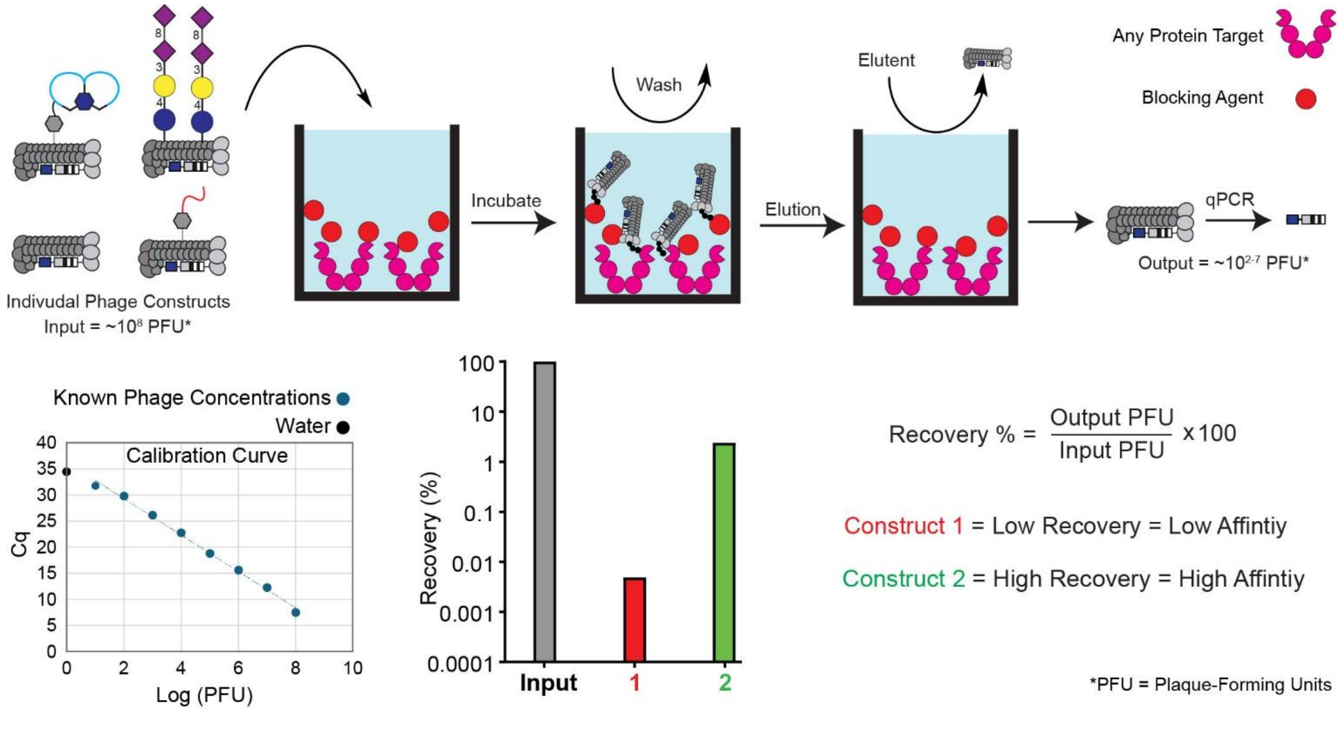

**Figure S18.** A general workflow of phage screening.

First incubate against the target then washing away unbound phage and eluting the bound phage, then through qPCR a recovery percentage can be found. A higher recovery showing a higher affinity of the construct to the target, by contrast a low recovery showing low affinity to the target.

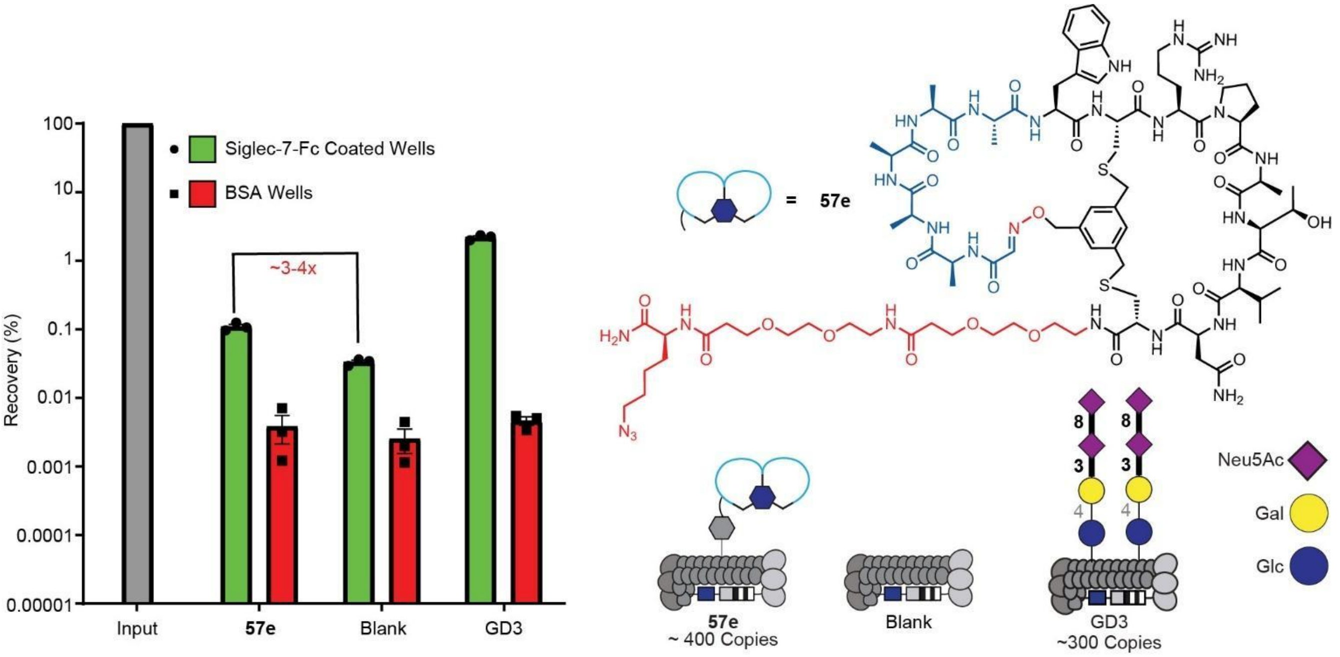

**Figure S19.** A preliminary test run with blank phage.

Preliminary test measuring binding affinity of **57e** conjugated on phage at an approximate density of 400 copies, and using blank phage as a negative control. Initial tests marked unexpected binding of blank wild-type M13 virion to purified Siglec-7 protein.

5.3.2 Phage recovery on cells

To cells cultured in T75 flasks (Thermo Fisher, Cat# 156800) that have reached confluency we remove the culture media (and if adherent detach cells by treating with 4 mL of TrypLE) and pellet the cells by centrifuge at 1000 rpm for 4 mins removing the supernatant, then resuspend the cell pellet in 5 mL of ice-cold washing buffer (1x HBS + 0.1% BSA), taking out 10 µL for cell counting. We then aliquot $10^6$ cells to each 5 mL Falcon tubes (Fisher Scientific, Cat# 14-959-2A), pelleting the cells by centrifuge at 300 kg. for 5 mins, removing the supernatant. Resuspend the cell pellet gently with 100 µL of the phage construct in incubation buffer (1x HBS + 1% BSA) at a concentration of ~$10^9$ PFU/mL (input), incubating the mixture for 1hr at 4ºC. Gently resuspend the cells, and to each tube add 3 mL of ice-cold washing buffer, pelleting the cell by centrifuge at 300 xg for 5 mins. Decant the supernatant to a waste container and orient the tube opening facing downward and press against a paper towel to remove residual liquid. Repeat the process by gently resuspending the cell pellet with 100 µL of ice-cold washing buffer and adding 3 mL to each tube. The process is then repeated by resuspending the cell pellet in 800 µL of 1x HBS buffer and transferred to 1.7 mL microcentrifuge tubes (Fisher Scientific, Cat# 14-222-168), pellet the cells by centrifuge at 300 x g for 5 mins, pipetting away the supernatant. To each tube, resuspend the cells in 50 µL of cell lysis buffer (0.1 mg/mL RNase A (Fisher Scientific, Cat#

FEREN0531) in 1x HF buffer). Vortex the cells at maximum speed for 1 min to release the bound phages. Centrifuge at 2000 xg for 3 min, transfer the supernatant to new tubes, then add 1 μL of 10 mg/mL Proteinase K (Thermo Fisher, Cat# 25530049) in nuclease free water (IDT, Cat# 11-04-02-01). Incubate the tubes at 55ºC for 10 mins, followed by heat-denaturing at 95ºC for 3 min. Centrifuge the tubes at 21000 xg for 3 mins, transfer 30 μL of the supernatant into new tubes (output) for qPCR quantification and PCR for NGS.

5.4 LiGA mixed with **57e** phage construct

Given the initial positive affinity observed of our phage construct (Figure 6b), we sought to observe what interactions our construct would have when mixed with our LiGA construct. Initial testing was done taking our peptide-conjugated phage with PEG8 on the surface, and taking PEGylated (PEG45-conjugated phage) and mixing it with our LiGA mixture. The resulting mixture was then screened against wells coated with Siglec-7-Fc and blank BSA wells as a control. Following deep sequencing getting the resulting enrichment we then forced the FC of PEGylated phage to be 1 and adjusted all the other constructs accordingly. While sialo-glycans should massive enrichment to Siglec, our peptide-conjugated construct showed a 56x fold enrichment (Figure S20). When checking enrichment to BSA coated wells, following same methodology, our peptide-conjugated phage showed a 1.3x fold enrichment over PEGylated phage (Figure S21). Highlighting the affinity and specificity of our construct to Siglec-7.

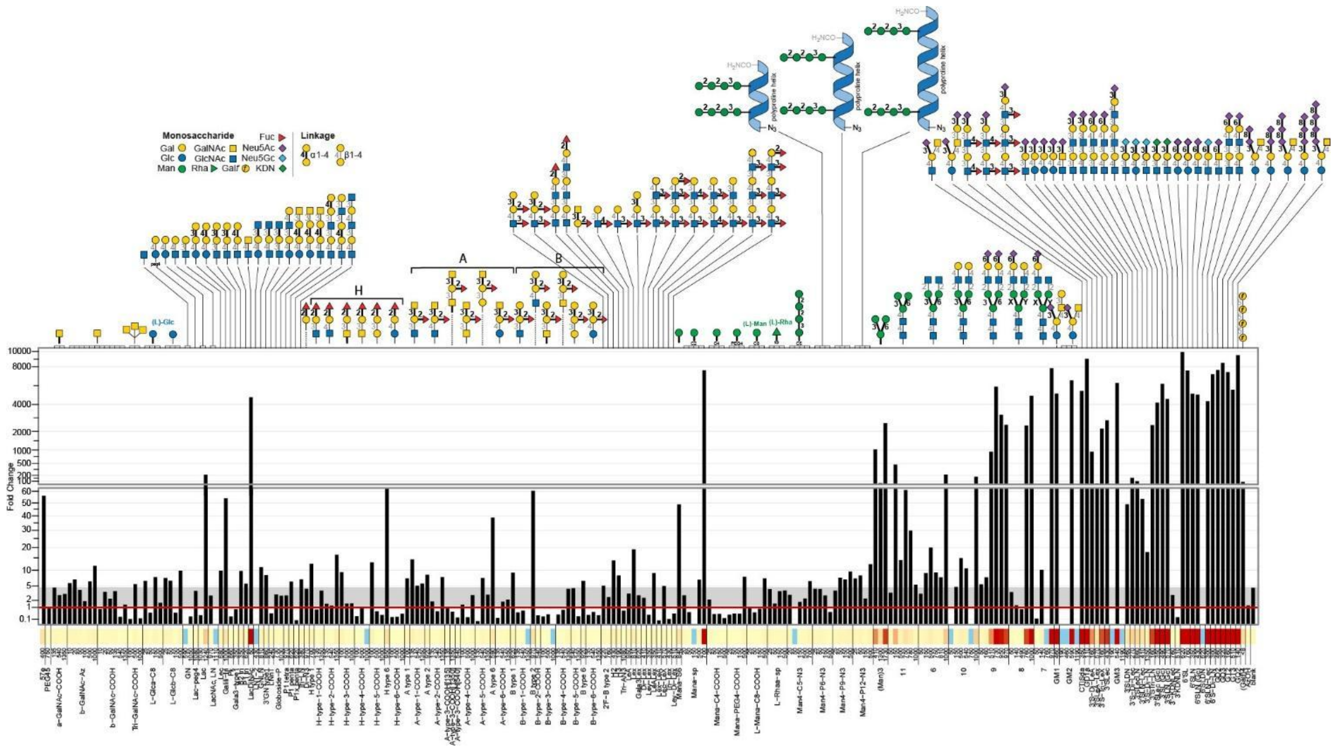

**Figure S20.** Bar chart showing affinity of LiGA mixture against wells coated with Siglec-7-Fc.

The left-most 2 columns being our peptide-conjugated phage construct and PEGylated phage (PEG45-conjugated phage) respectively.

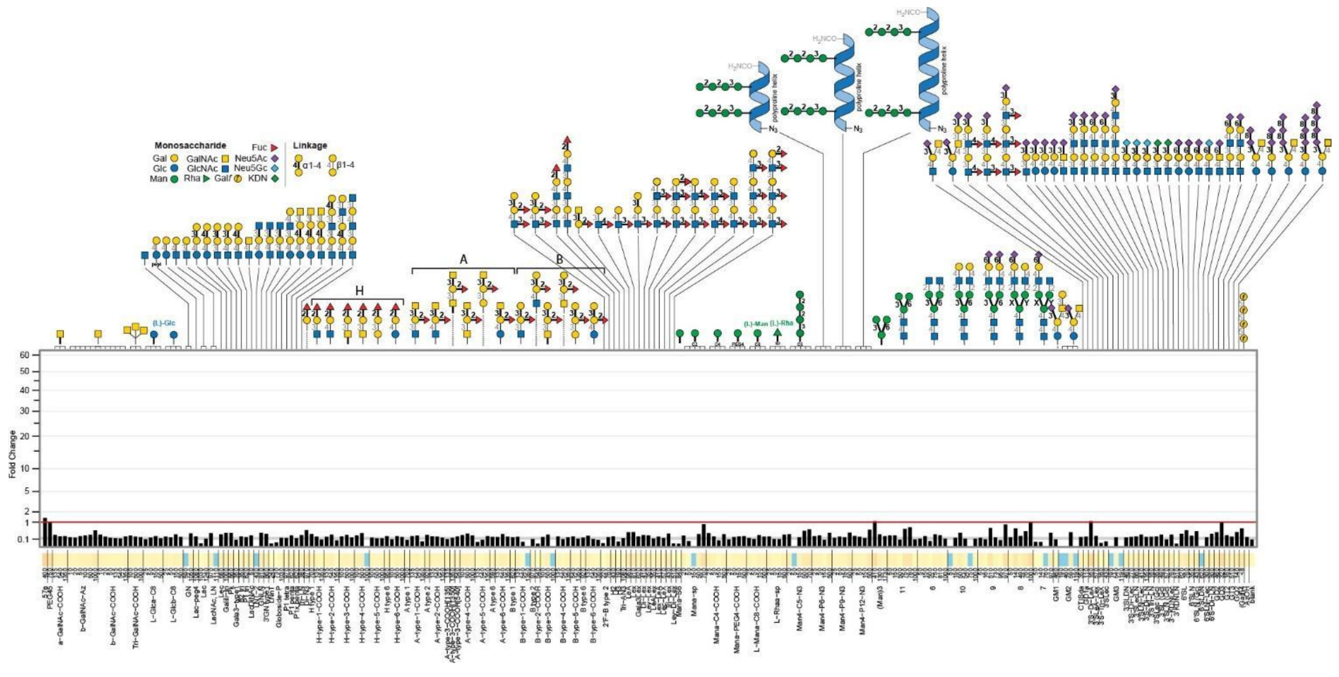

**Figure S21.** Bar chart showing affinity of LiGA mixture against wells coated with BSA.

The left-most 2 columns being our peptide-conjugated phage construct and PEGylated phage (PEG45-conjugated phage), respectively.

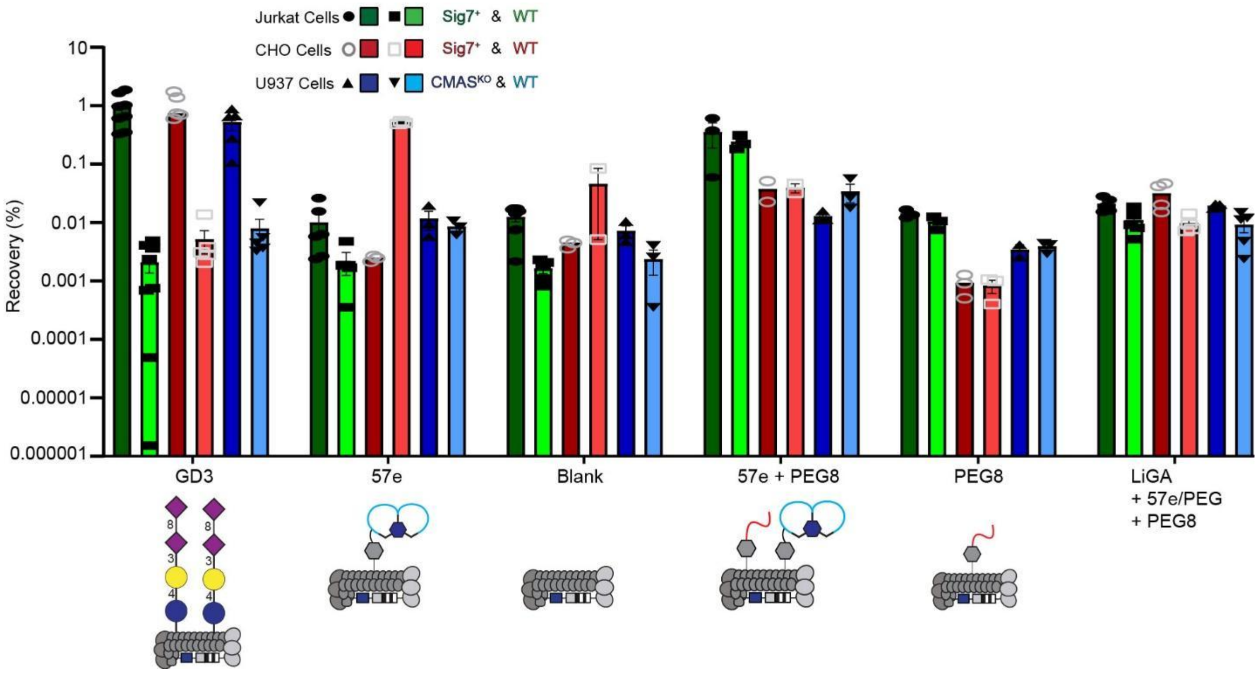

**Figure S22.** Phage recovery bar plot against cells.

Phage recovery percentage bar plot of phage constructs screened against positive cell lines and their wild-type counterparts. The plots highlight hSiglec-7$^{+}$ Jurkat cells and its wild-type (dark green and light green respectively), hSiglec-7$^{+}$ CHO cells and its wild-type (dark red and light red respectively), and CMAS$^{KO}$ U937 cells and its wild-type (dark blue and light blue respectively)

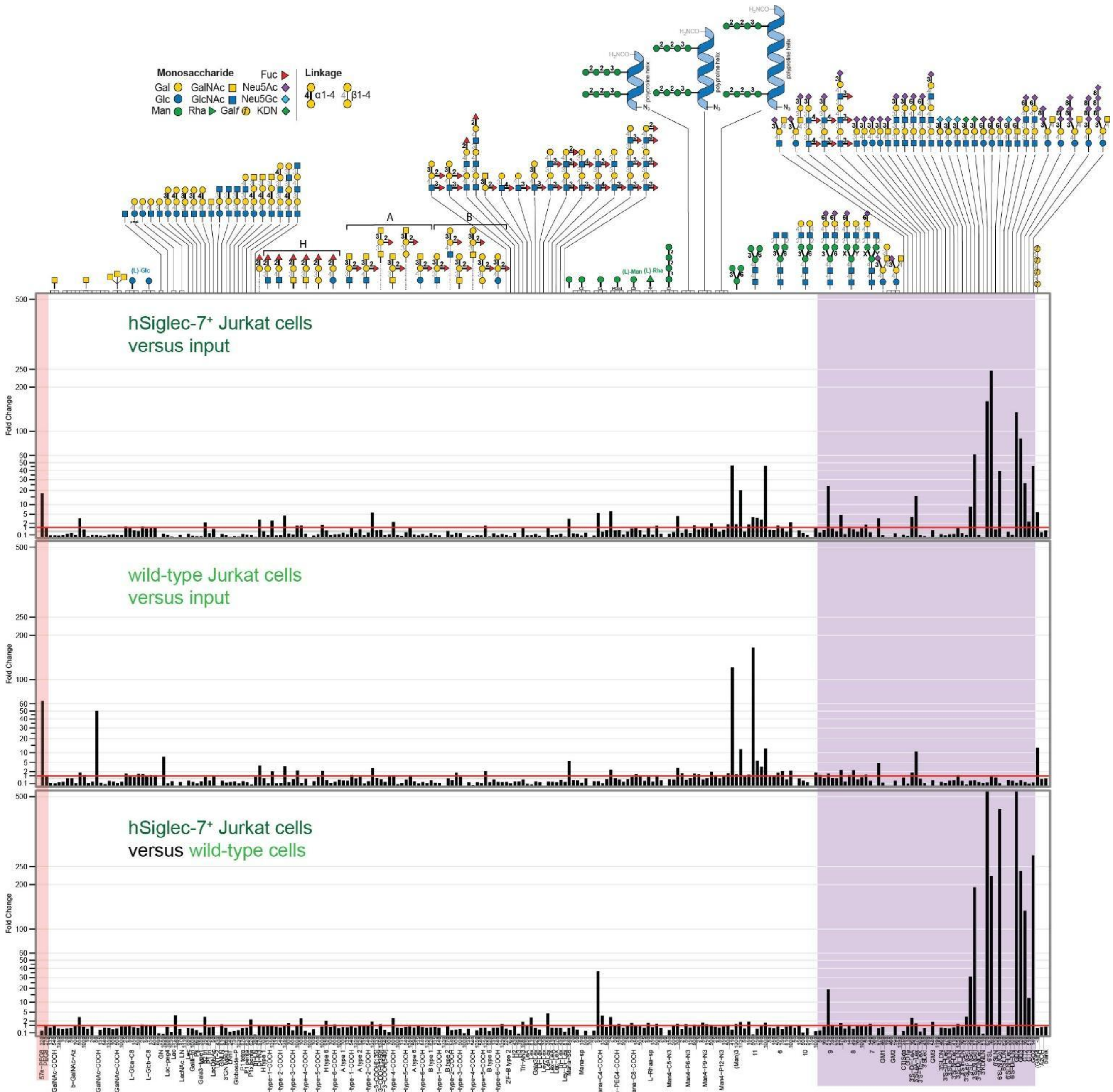

**Figure S23.** Bar chart showing affinity of LiGA mixture against Jurkat cells.

The left-most 2 columns being our peptide-conjugated phage construct and PEGylated phage respectively. Sialylated glycans are highlighted in purple on the right side.

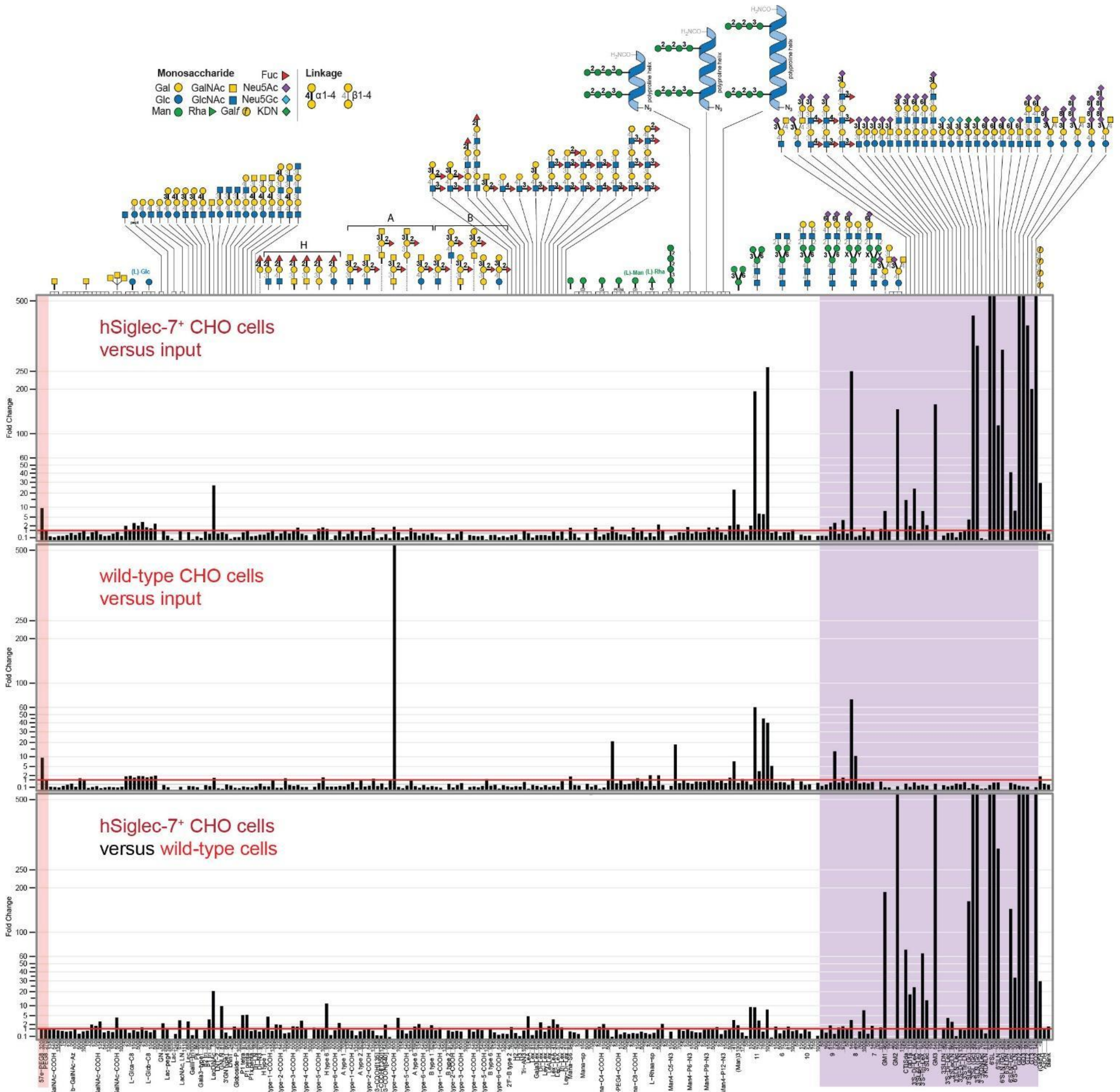

**Figure S24.** Bar chart showing affinity of LiGA mixture against CHO cells.

The left-most 2 columns being our peptide-conjugated phage construct and PEGylated phage respectively. Sialylated glycans are highlighted in purple on the right side.

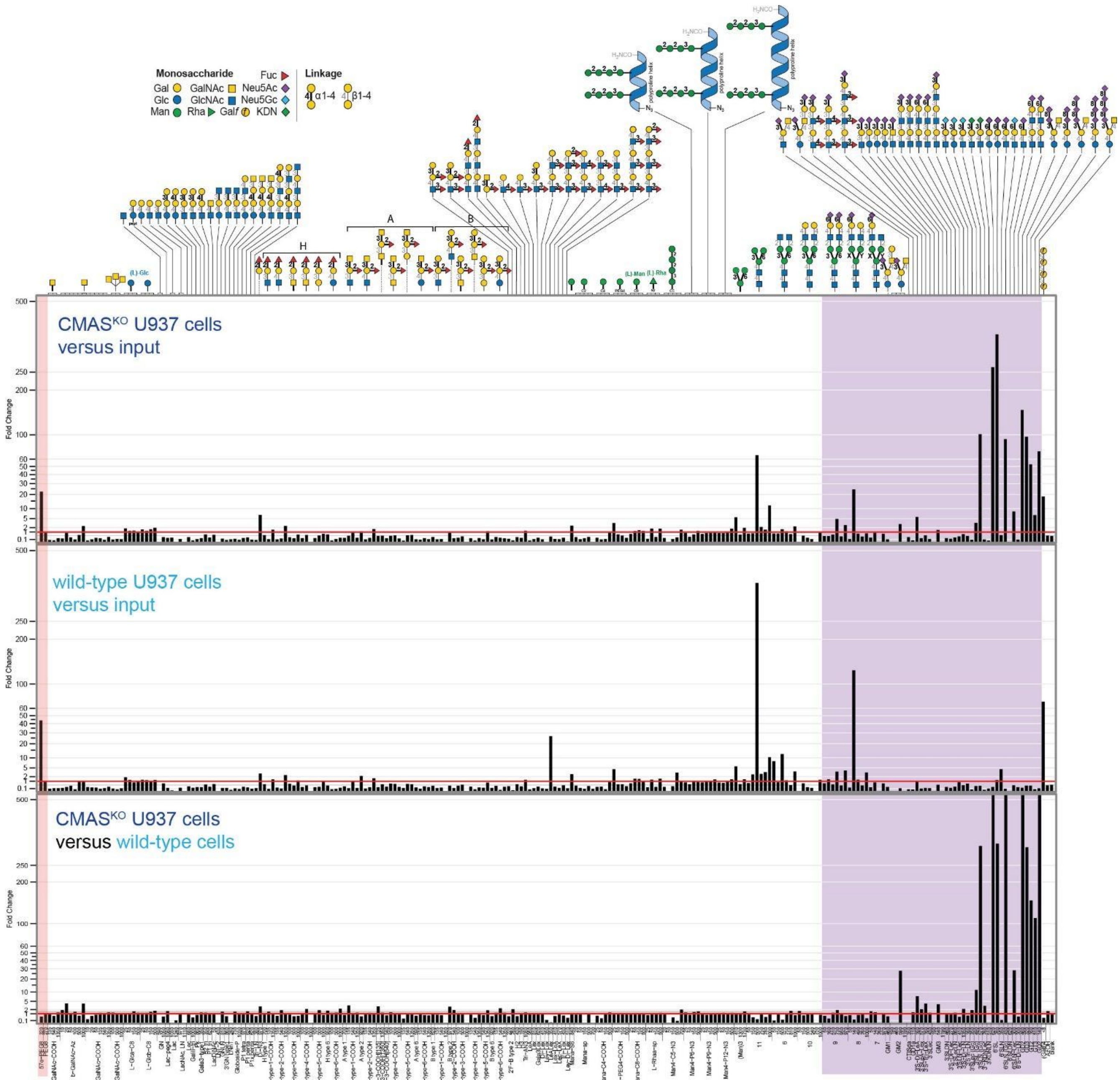

**Figure S25.** Bar chart showing affinity of LiGA mixture against U937 cells.

The left-most 2 columns being our peptide-conjugated phage construct and PEGylated phage respectively. Sialylated glycans are highlighted in purple on the right side.

### 6 Data summary pages

#### 6.1 SPR data curves

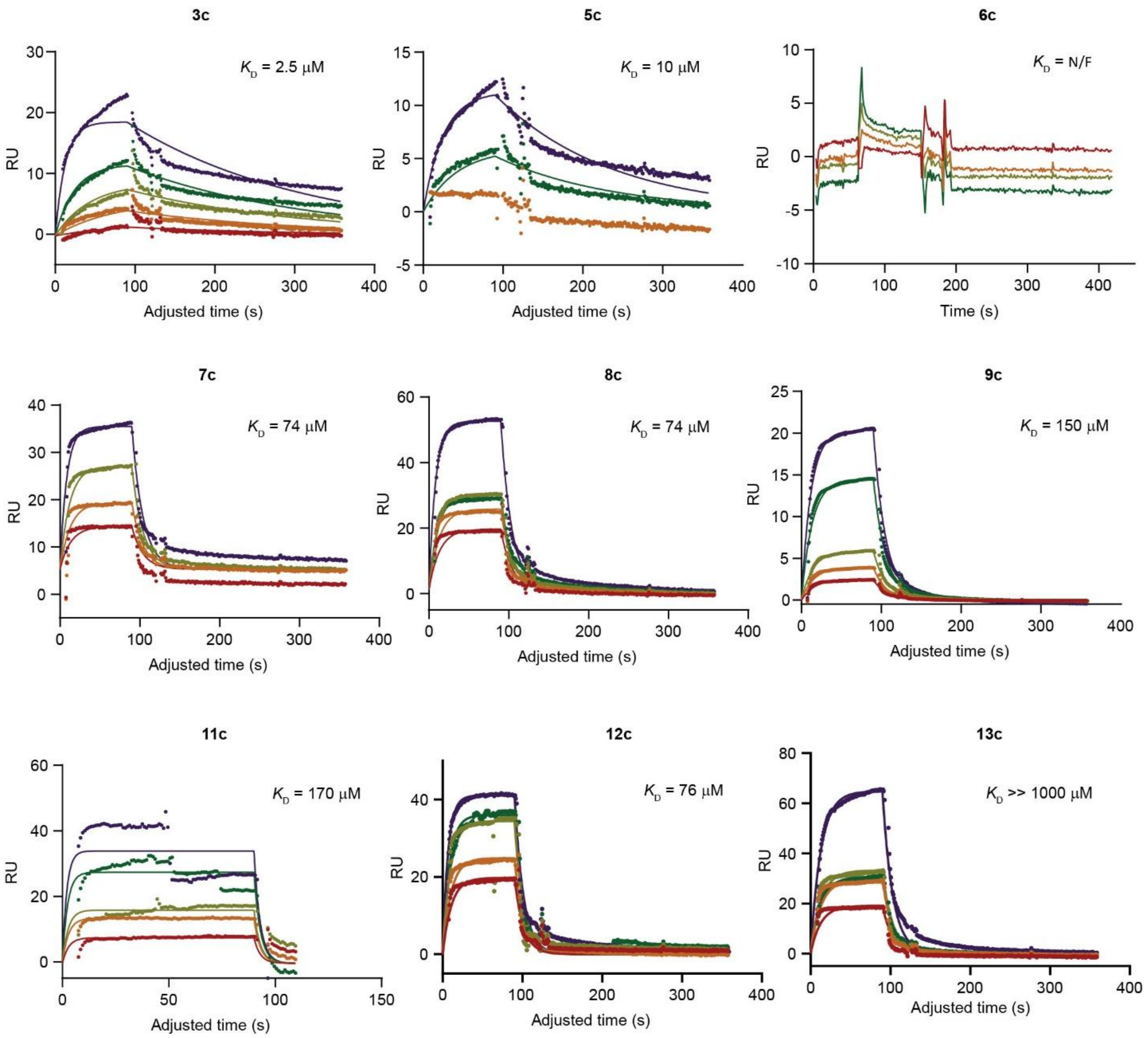

**Figure S26**. Fitted SPR curves for Siglec-7 (**Seq. 3-13**).

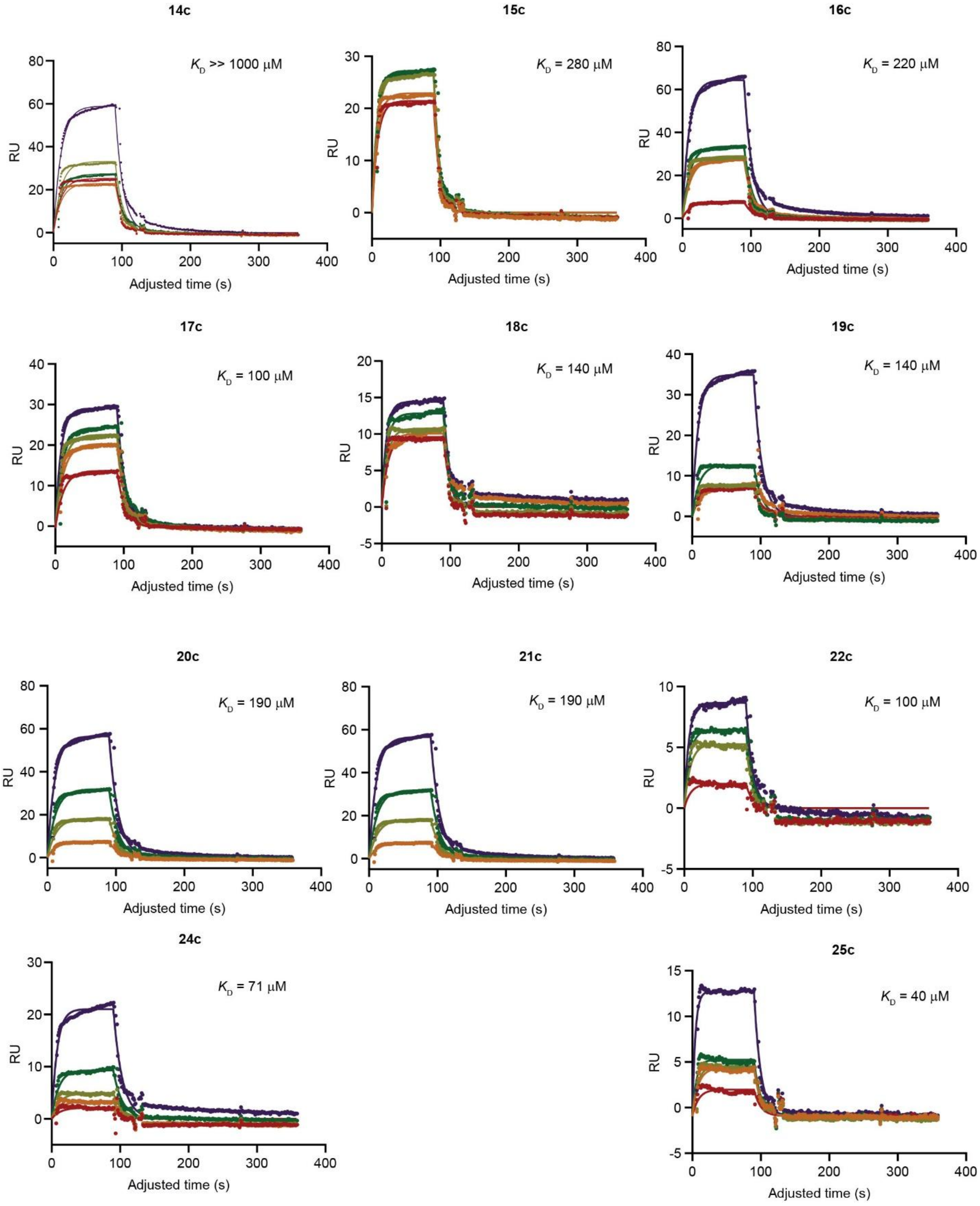

**Figure S27**. Fitted SPR curves for Siglec-7 (**Seq. 14-25**).

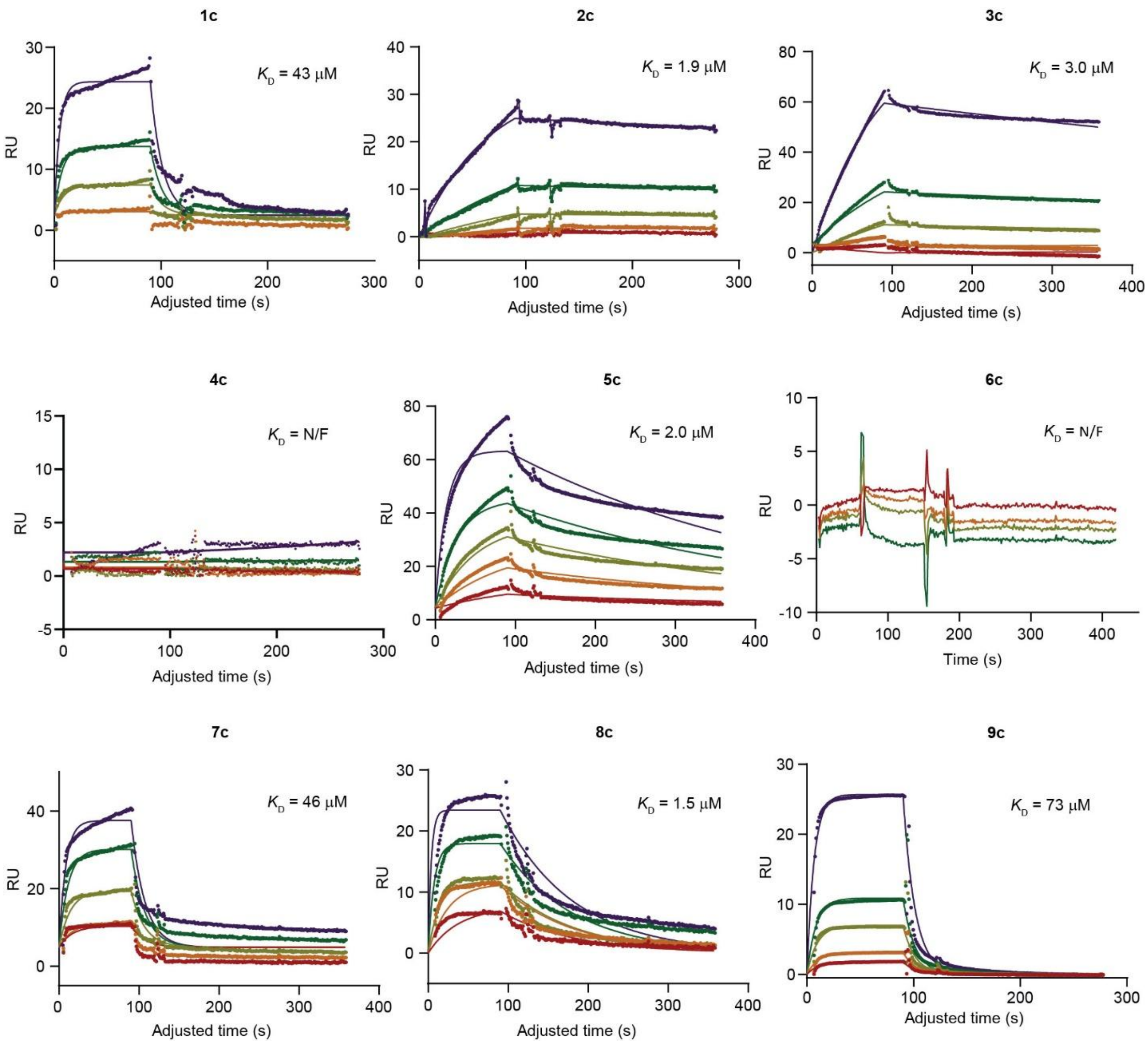

**Figure S28**. Fitted SPR curves for Siglec-7-FC (**Seq. 1-9**).

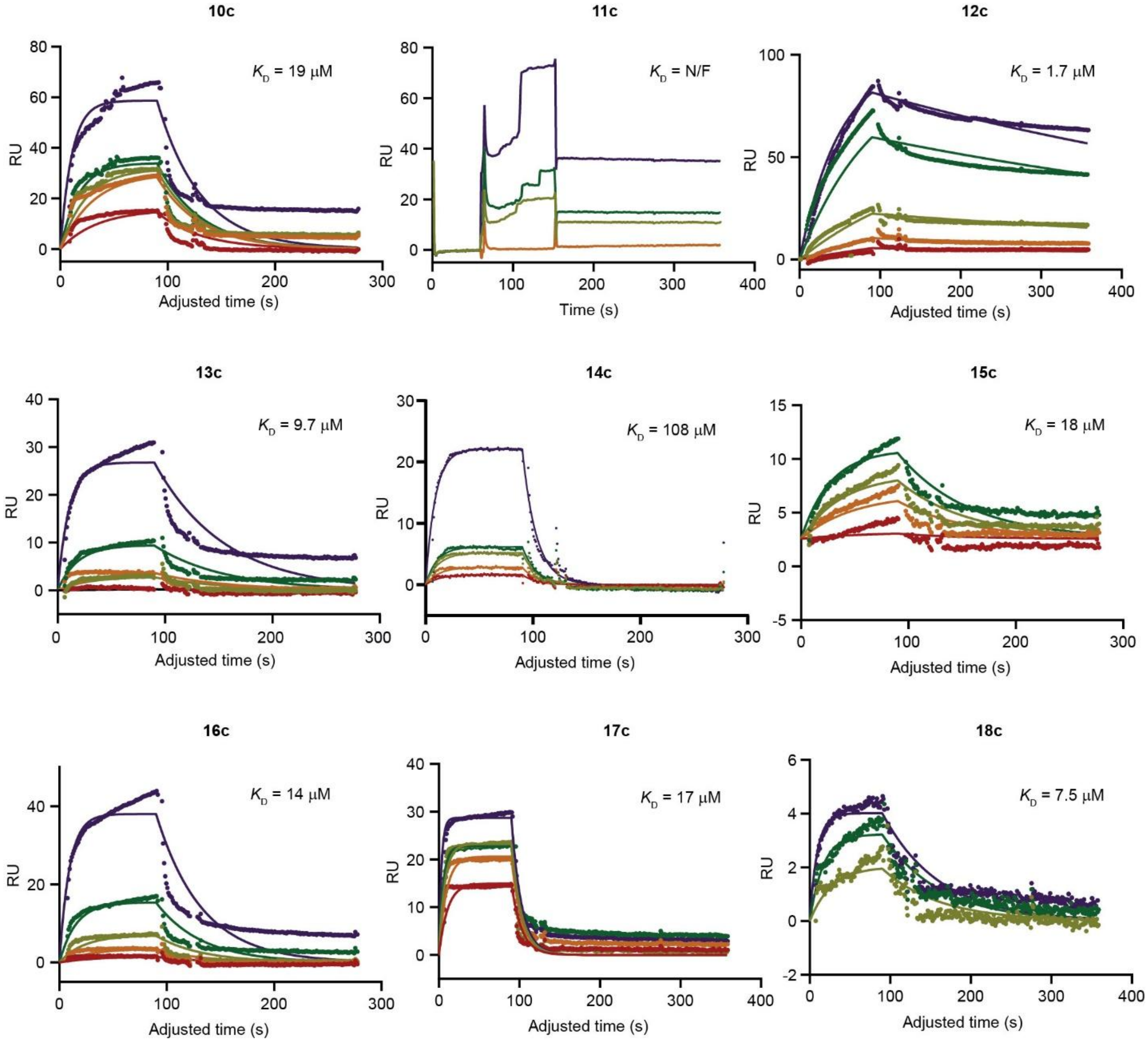

**Figure S29**. Fitted SPR curves for Siglec-7-FC (**Seq. 10-18**).

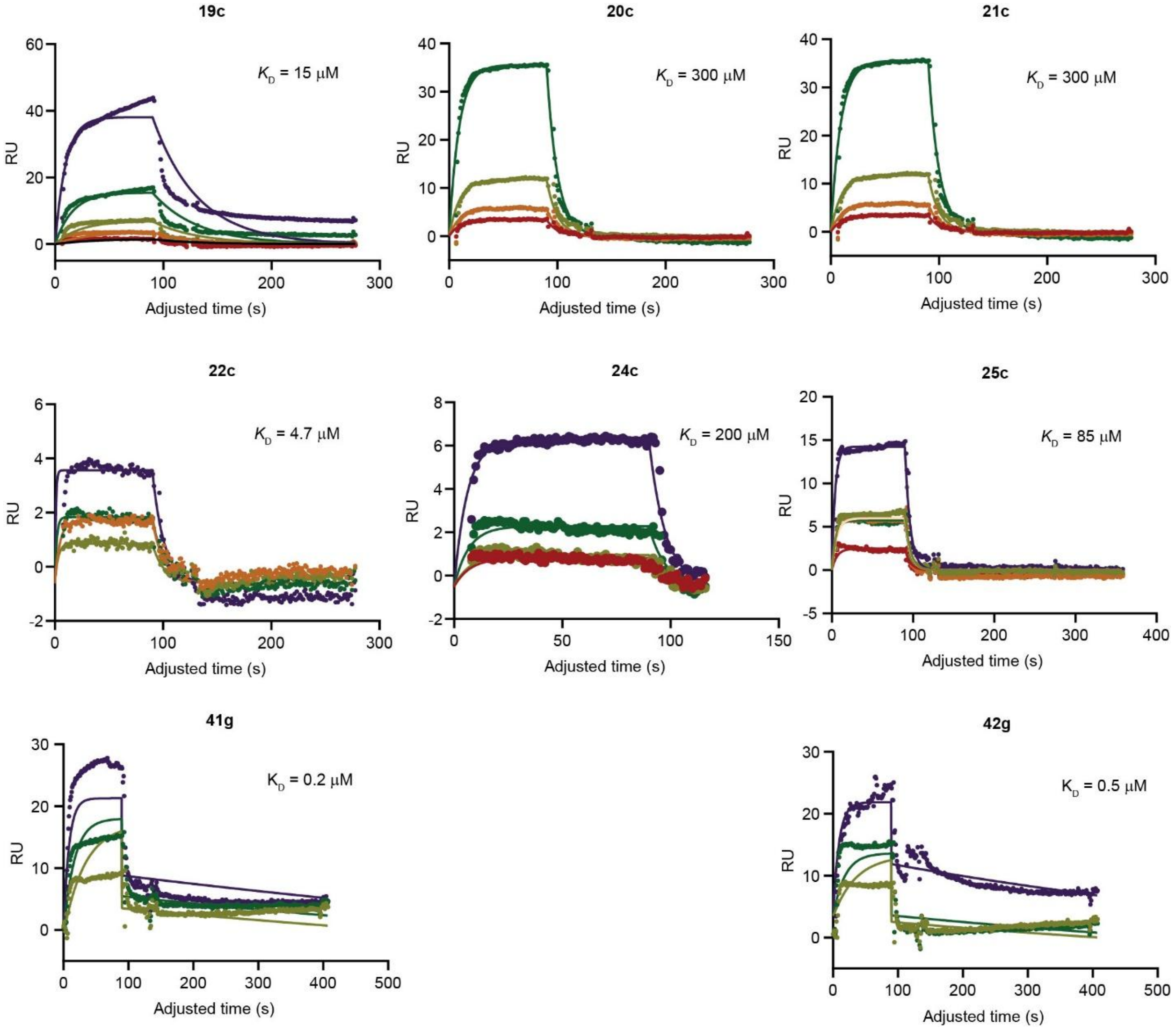

**Figure S30**. Fitted SPR curves for Siglec-7-FC (**Seq. 19-42**).

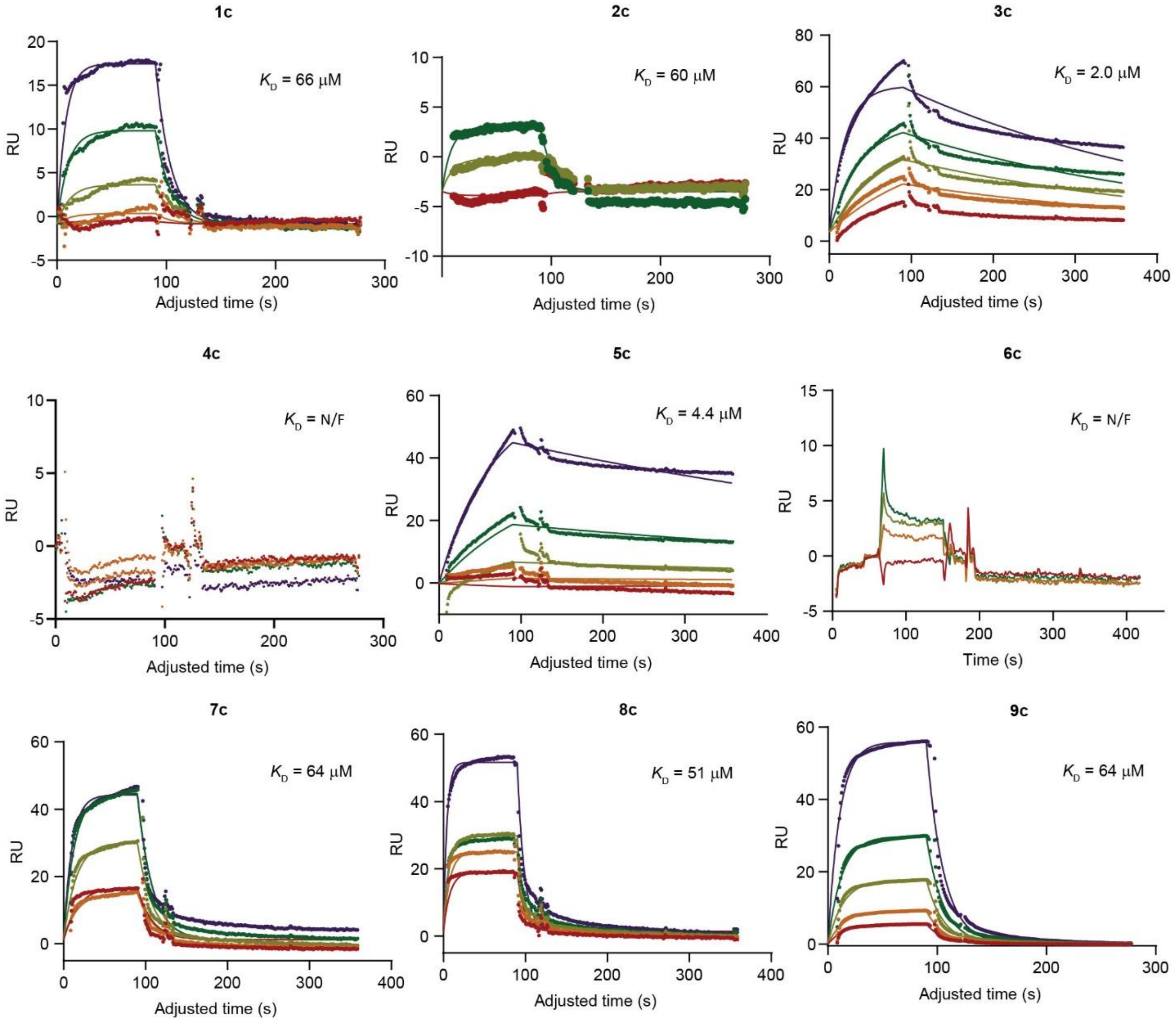

**Figure S31.** Fitted SPR curves for Siglec-9-FC (**Seq. 1-9).**

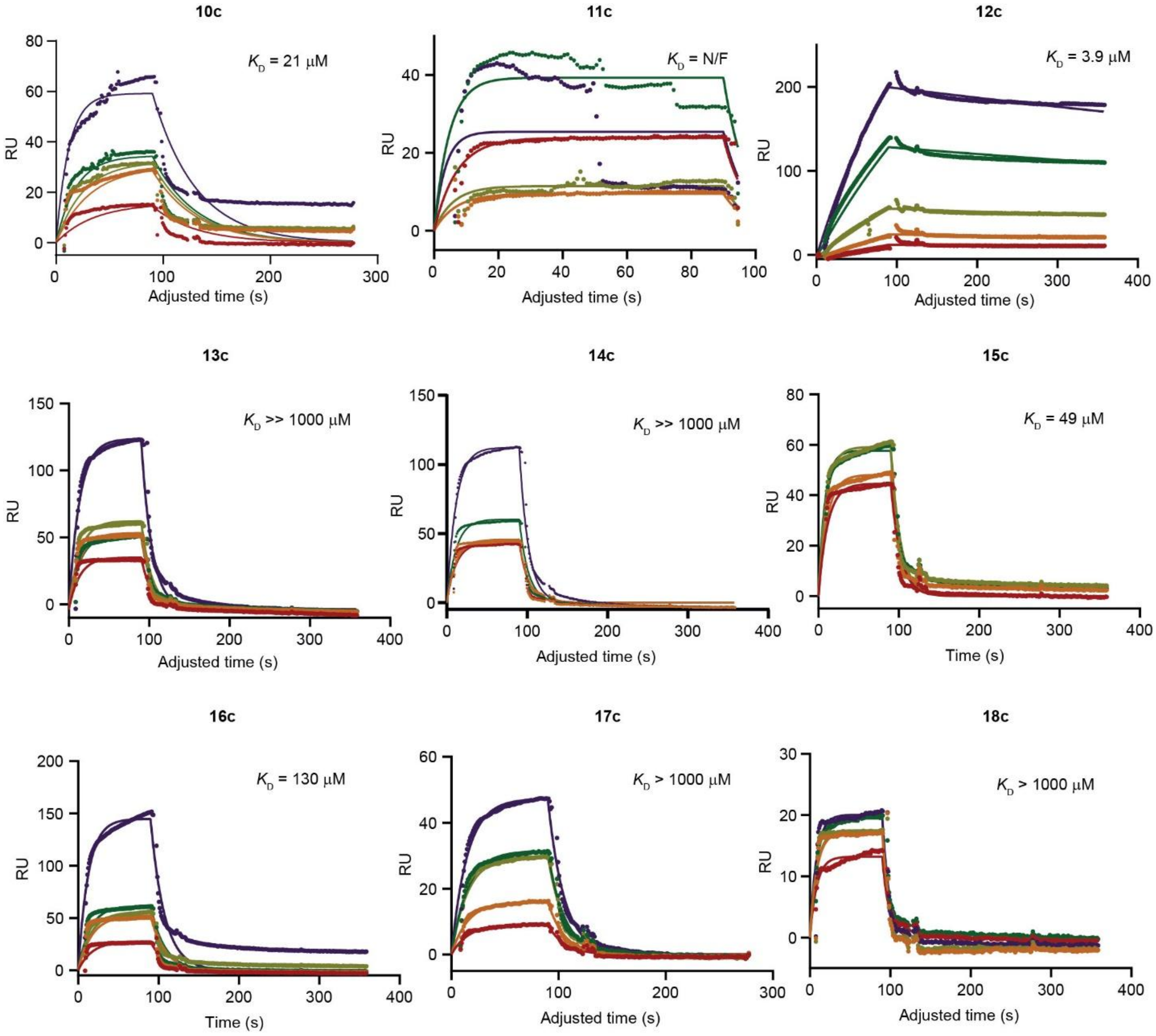

**Figure S32**. Fitted SPR curves for Siglec-9-FC (**Seq. 10-18**).

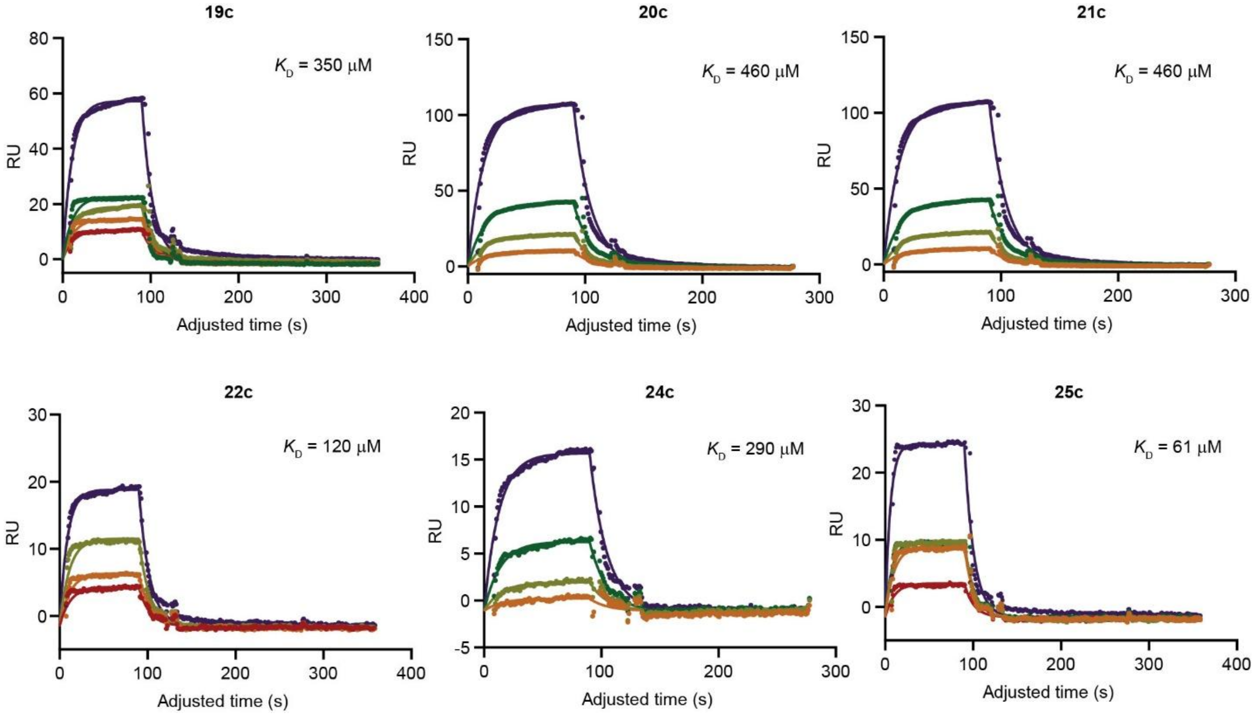

**Figure S33**. Fitted SPR curves for Siglec-9-FC (**Seq. 19-25**).

#### 6.2 ELISA plots

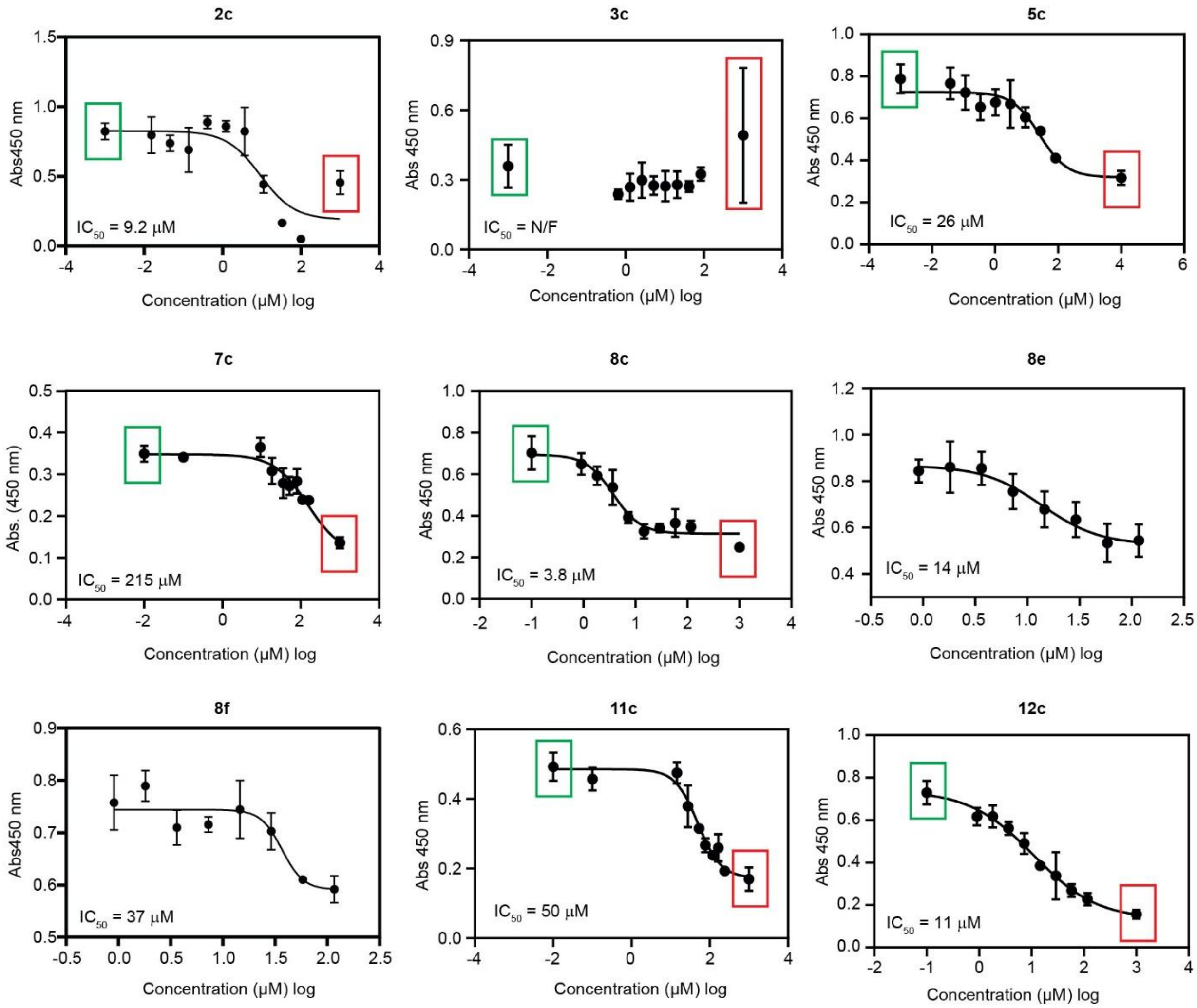

**Figure S34**. ELISA plots for Siglec-7-FC (**Seq. 2-12**)
[positive cont.; negative cont.].

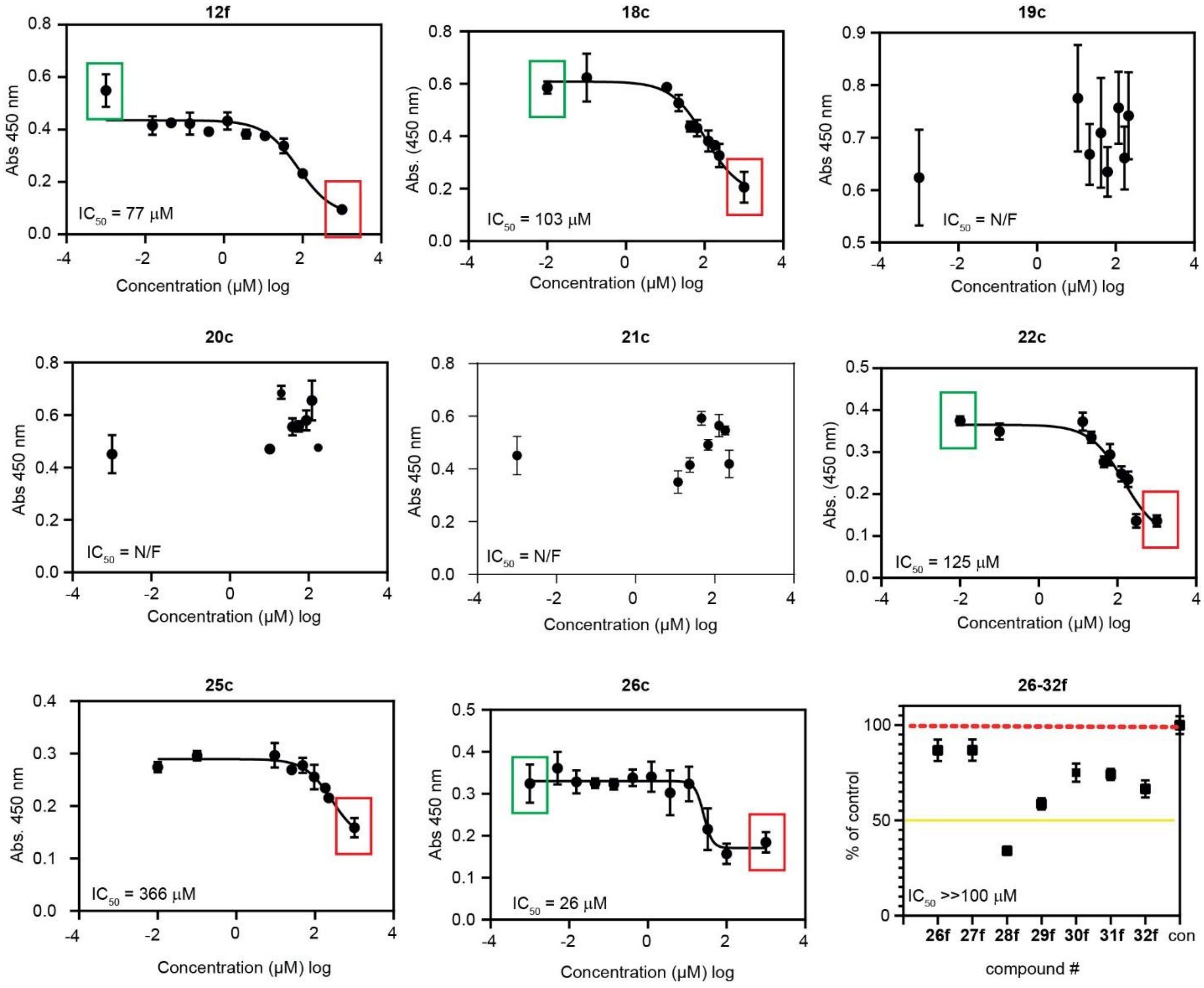

**Figure S35.** ELISA plots for Siglec-7-FC (**Seq. 12-32**)

[positive cont.; negative cont.].

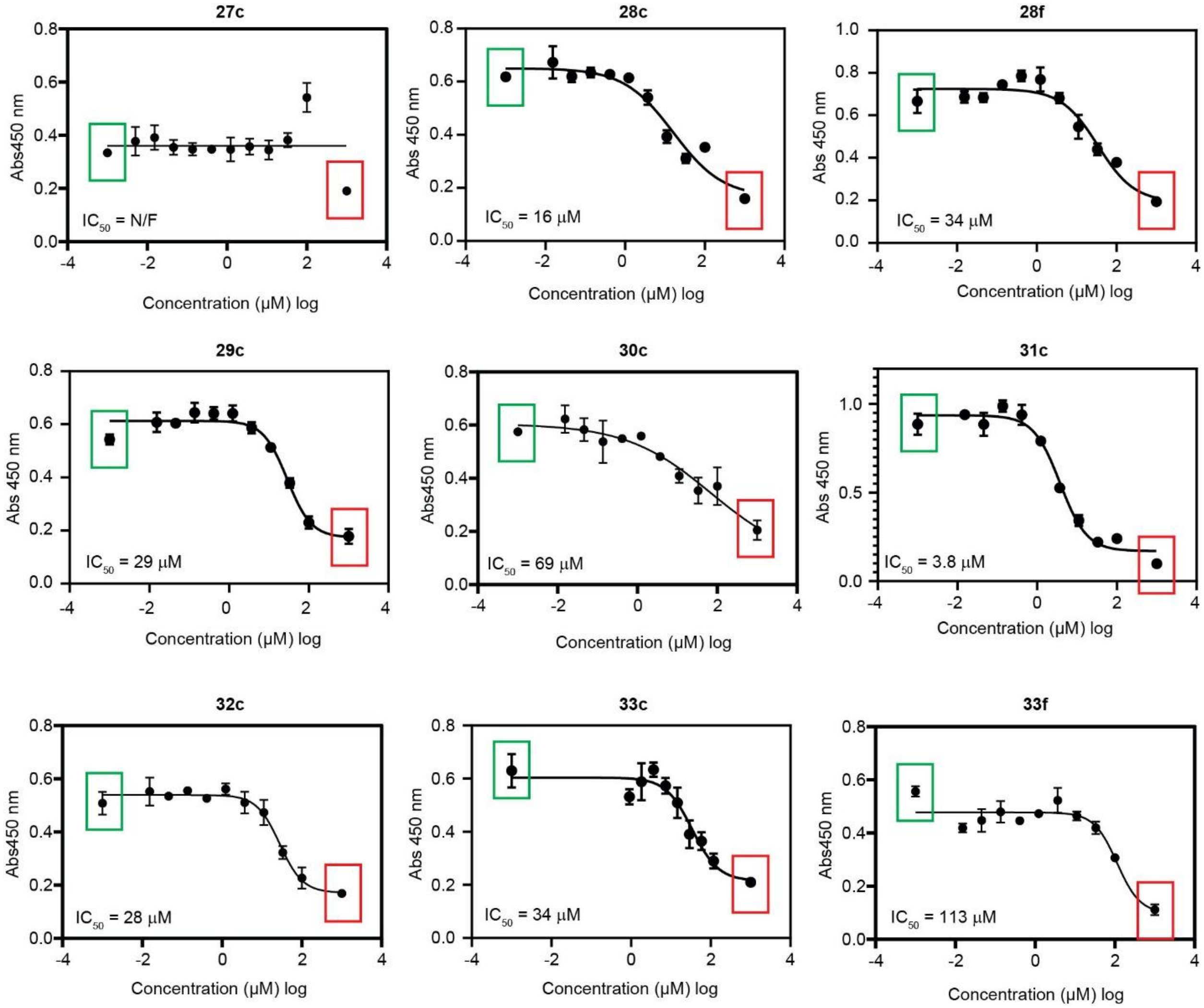

**Figure S36**. ELISA plots for Siglec-7-FC (**Seq. 27-33**)

[positive cont.; negative cont.].

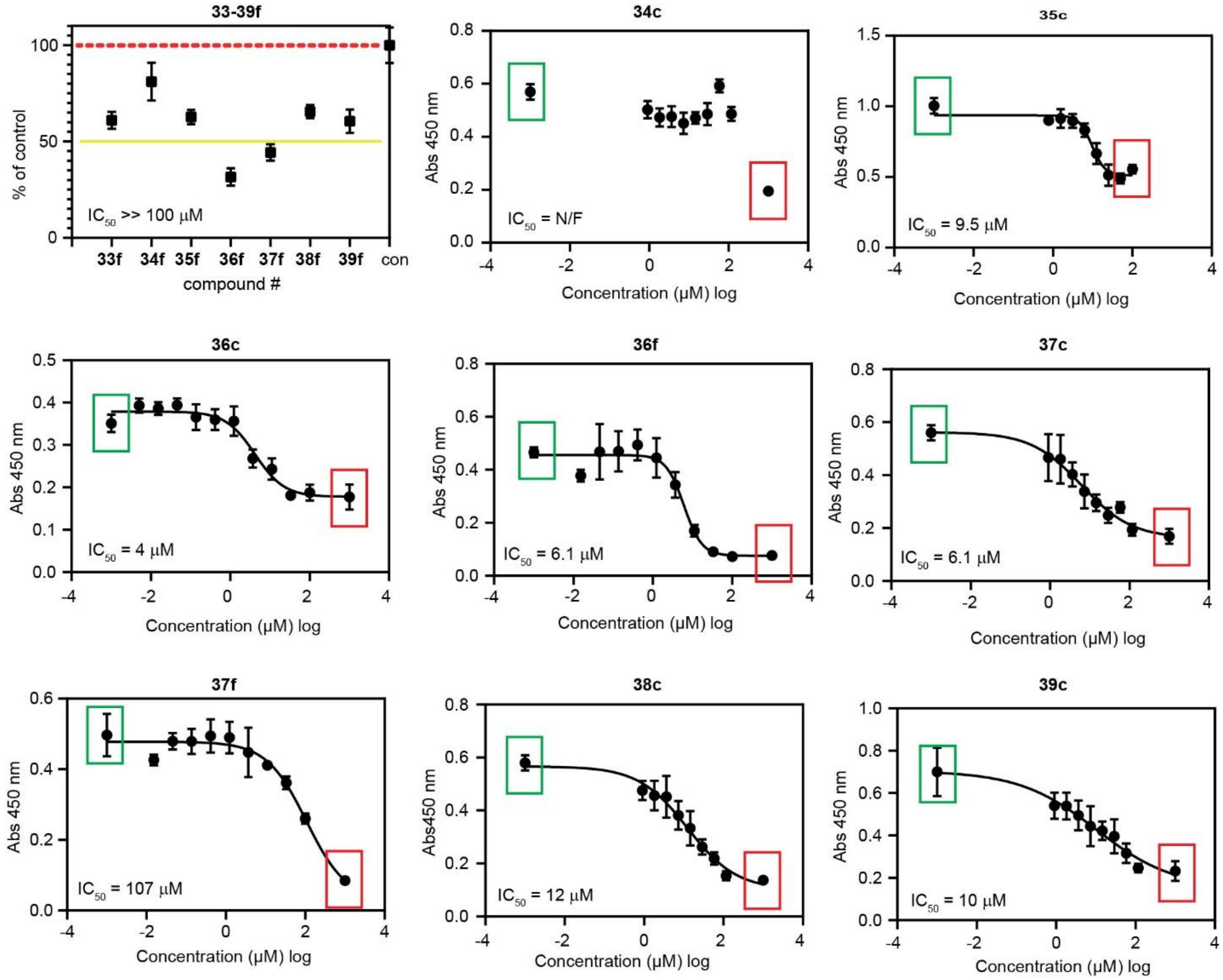

**Figure S37**. ELISA plots for Siglec-7-FC (**Seq. 33-39**)
[positive cont.; negative cont.].

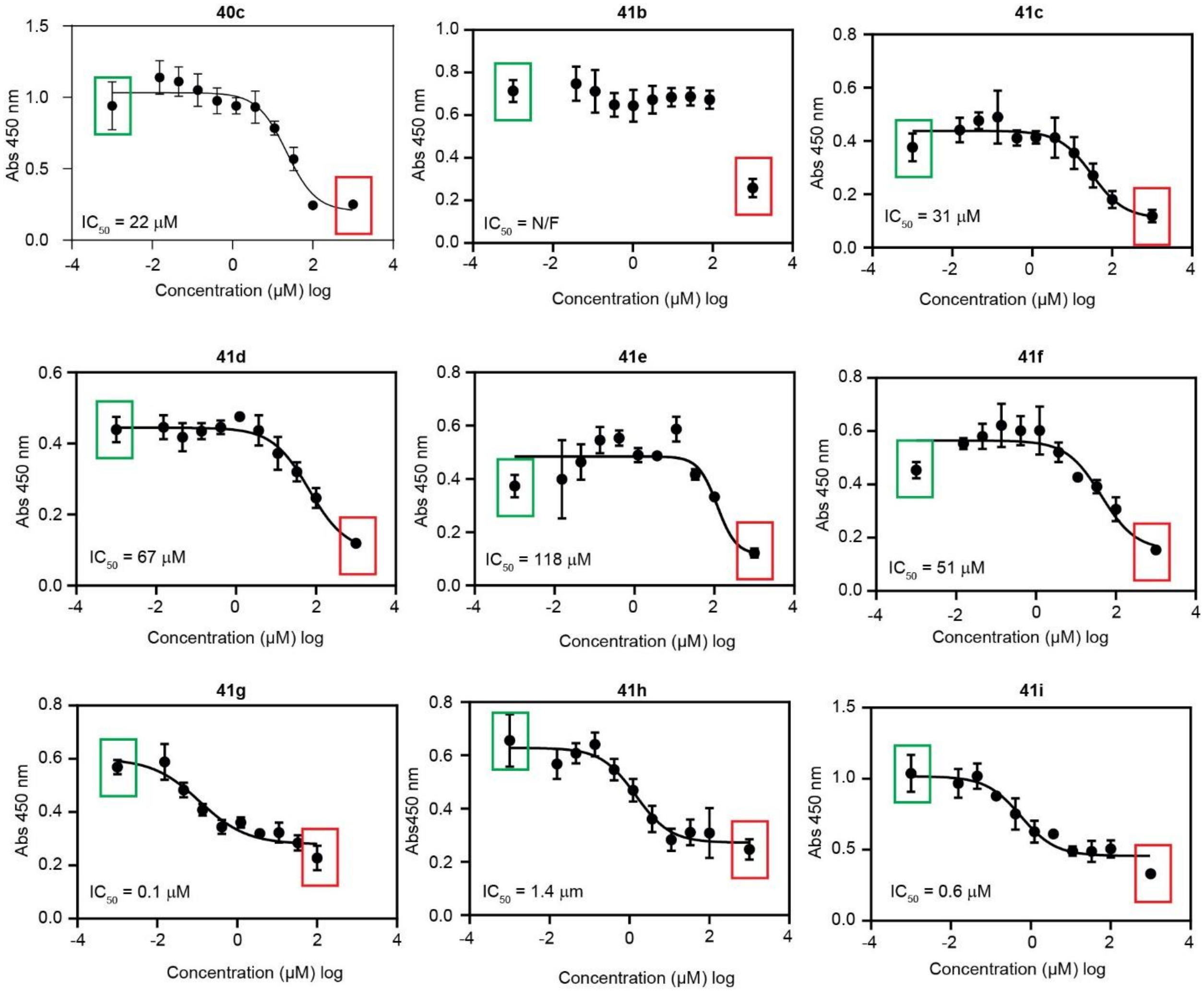

**Figure S38**. ELISA plots for Siglec-7-FC (**Seq. 40-41**)

[positive cont.; negative cont.].

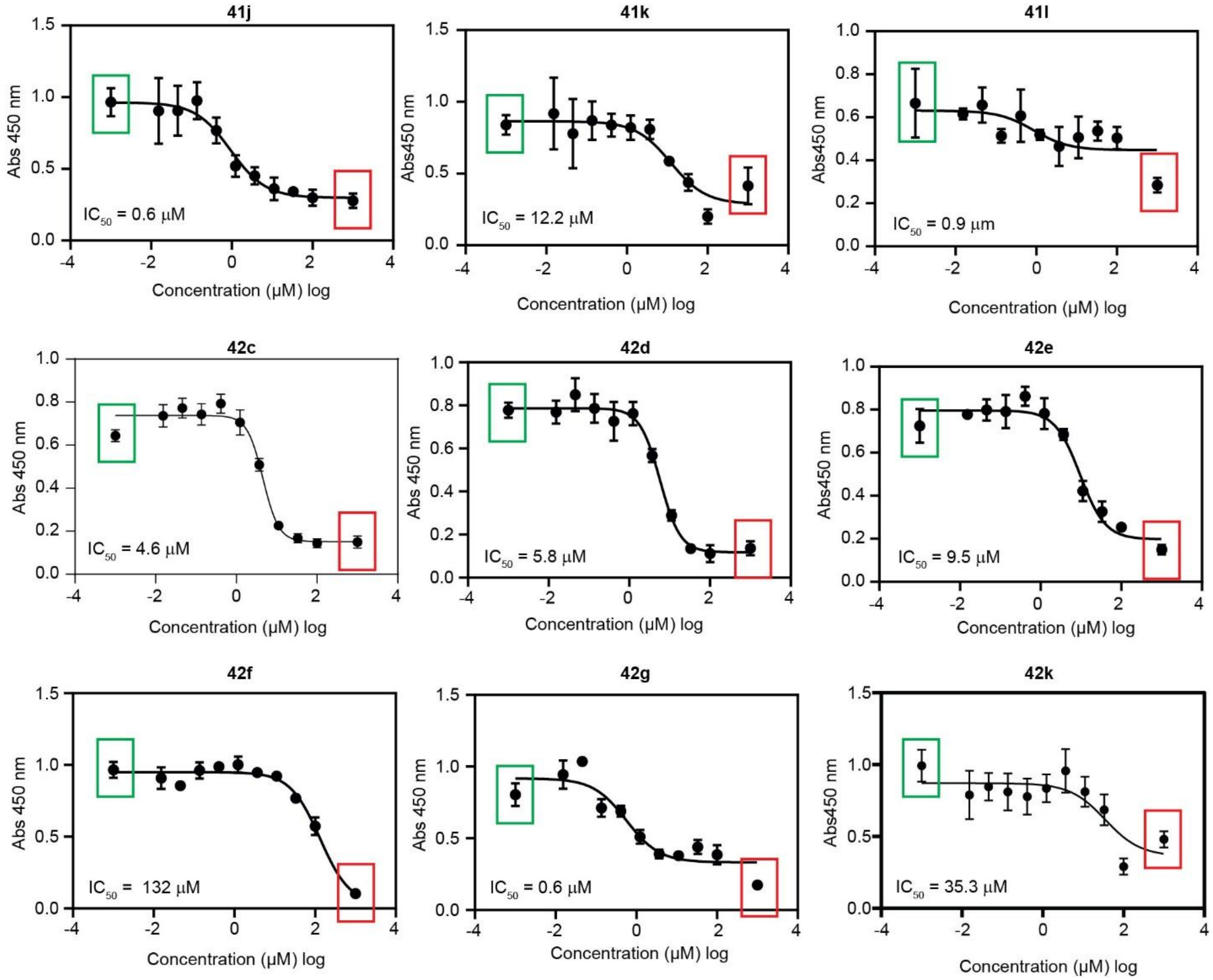

**Figure S39**. ELISA plots for Siglec-7-FC (**Seq. 41-42**)
[positive cont.; negative cont.].

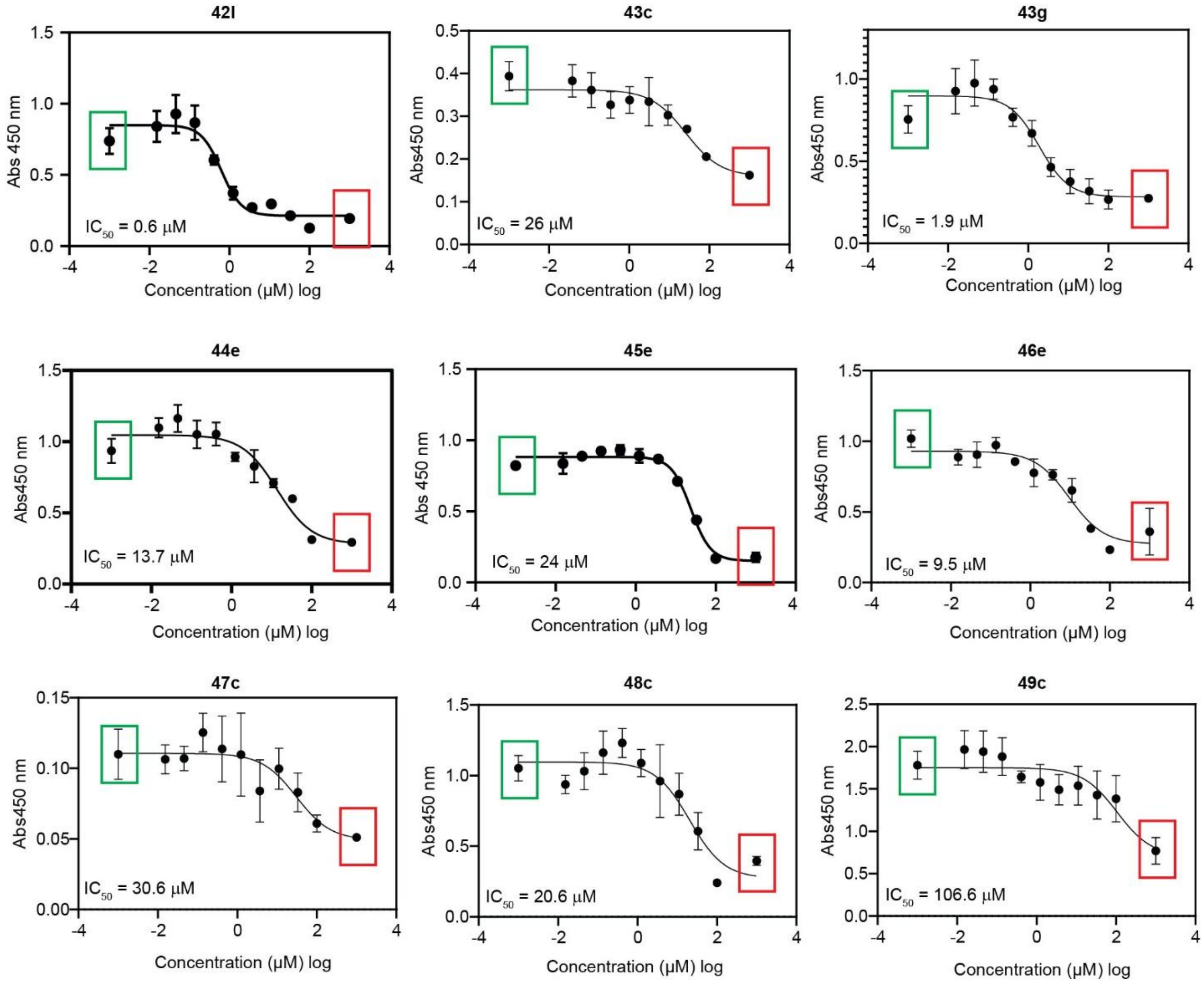

**Figure S40**. ELISA plots for Siglec-7-FC (**Seq. 42-49**)

[positive cont.; negative cont.].

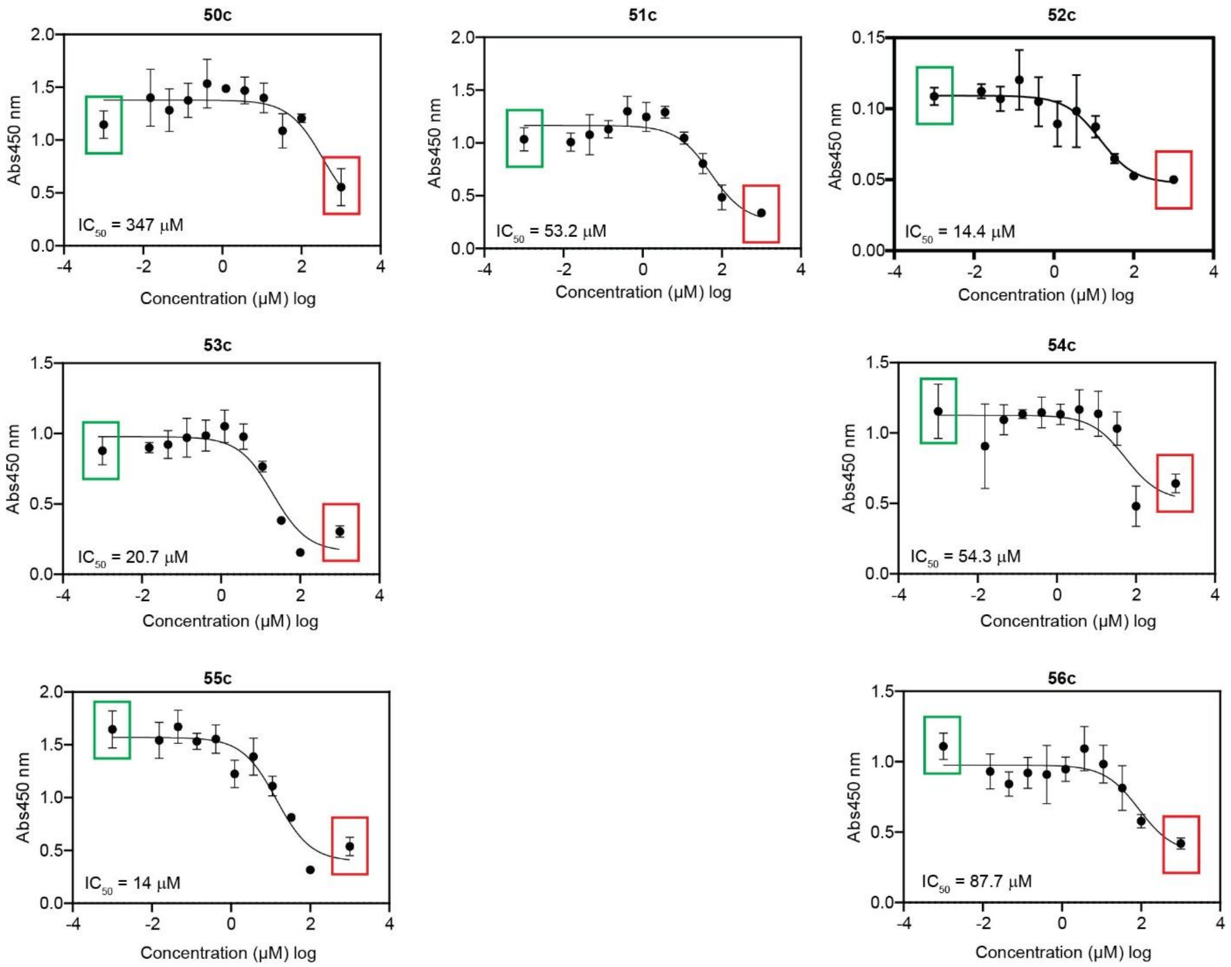

**Figure S41**. ELISA plots for Siglec-7-FC (**Seq. 50-56**)

[positive cont.; negative cont.].
