## Supporting Information 2 for "Non-Carbohydrate Inhibitors of Sialic Acid-binding Immunomodulatory-type Lectin-7 (Siglec-7) Discovered from Genetically Encoded Bicyclic Peptide Libraries"

### Peptide data pages

**TSL6-SHCAAYDQMC-OH**

Scale: 10.5 mg
Yield: 1.1 mg (9.0%)
Purity: ~80%

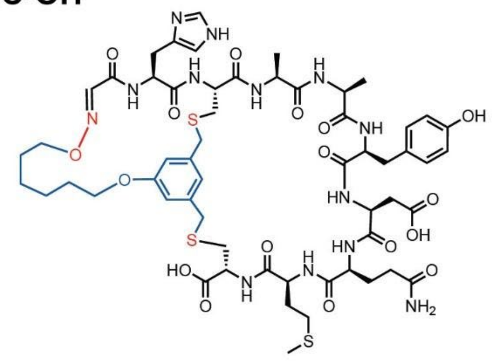

Chemical Formula: $C_{57}H_{77}N_{13}O_{17}S_2$
Exact Mass: 1311.47
Molecular Weight: 1312.50

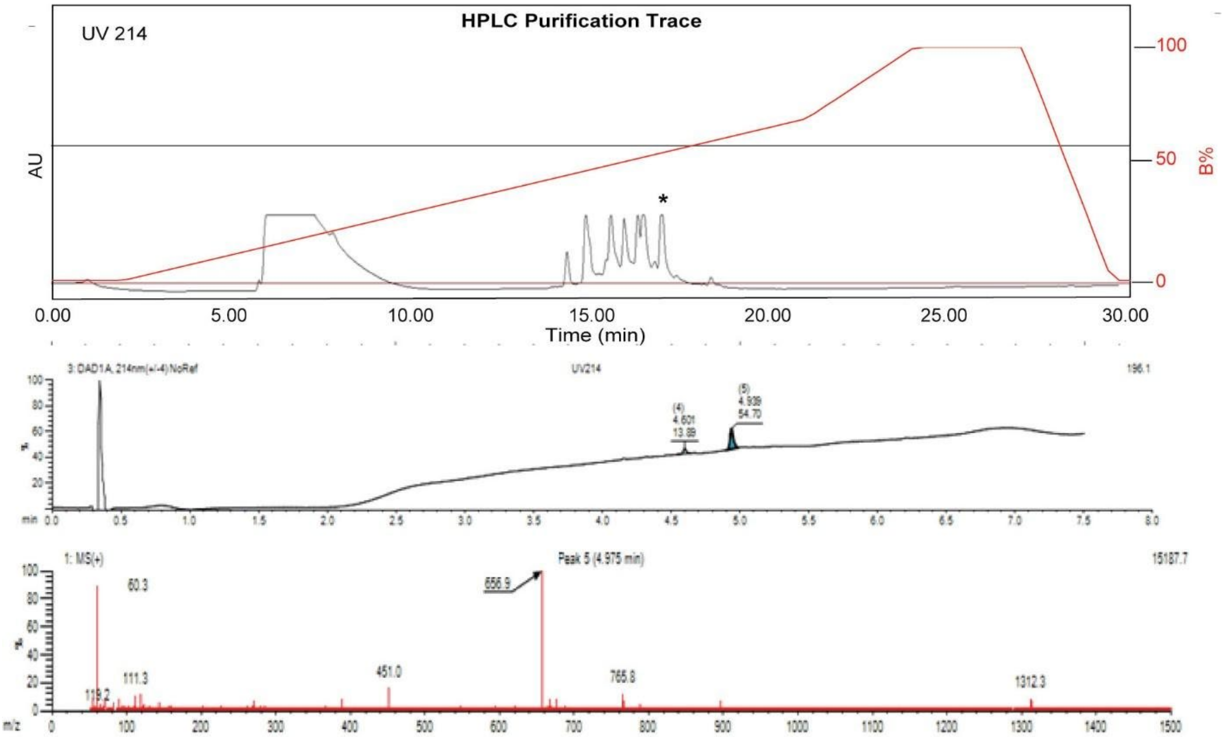

**Figure S42.** Data page for **1c**.

Expected: [M+1] = 1312.47, $[M+2]^{+2}$ = 656.735.

Observed: [M+1] = 1312.3, $[M+2]^{+2}$ = 656.9

**TSL6-SRCTGFHDIC-OH**
Scale: 10.0 mg
Yield: 1.1 mg (9.5%)
Purity: ~90%

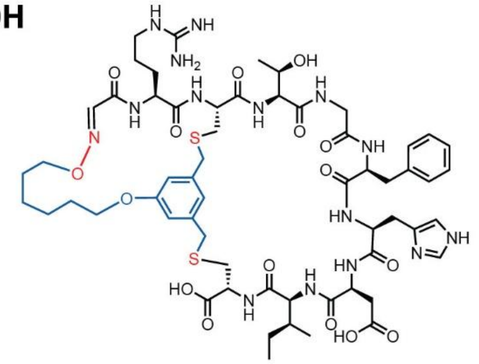

Chemical Formula: $C_{59}H_{83}N_{15}O_{16}S_2$
Exact Mass: 1321.56
Molecular Weight: 1322.52

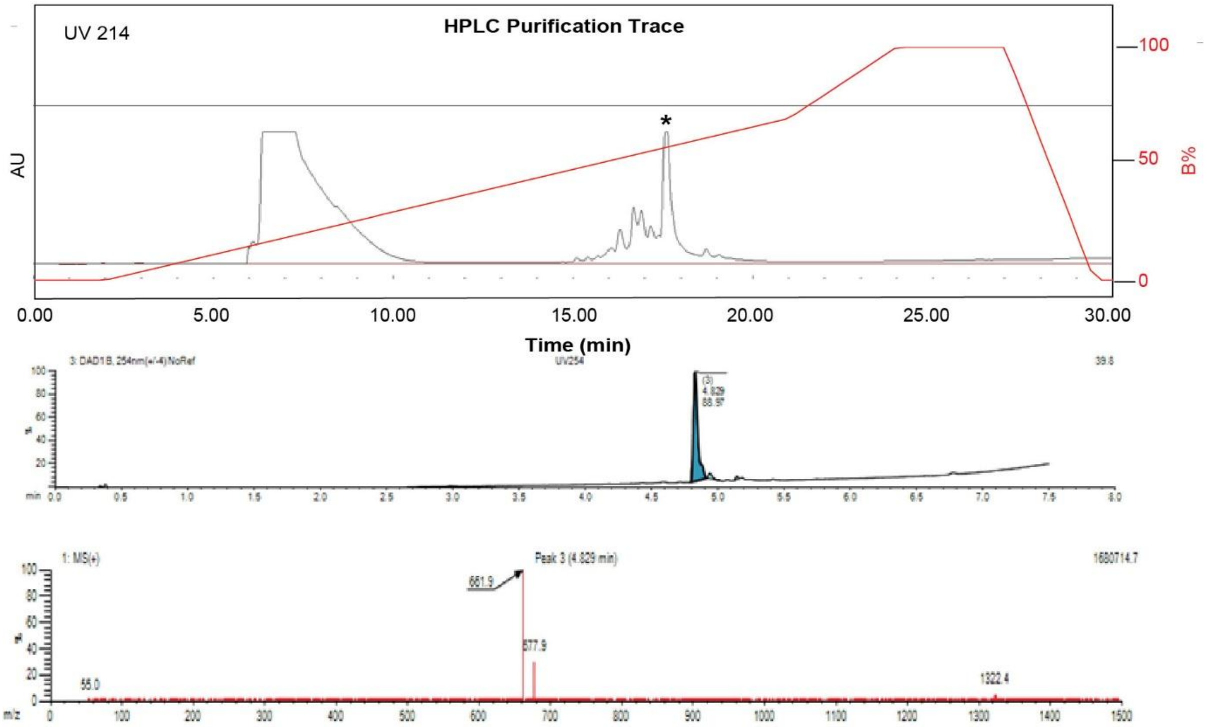

**Figure S43**. Data page for **2c**.

Expected: [M+1] = 1322.56, $[M+2]^{+2}$ = 661.78.

Observed: [M+1] = 1322.4, $[M+2]^{+2}$ = 661.9

### TSL6-SSCEQDVFYC-OH

Starting material mass: 14.0 mg
Final product mass: 3.1 mg (22%)
Purity: >95%

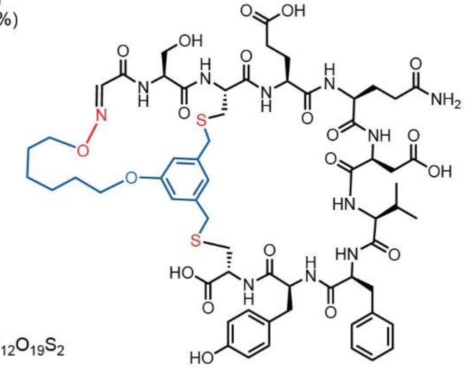

Chemical Formula: $C_{62}H_{82}N_{12}O_{19}S_2$
Exact Mass: 1363.51
Molecular Weight: 1364.51

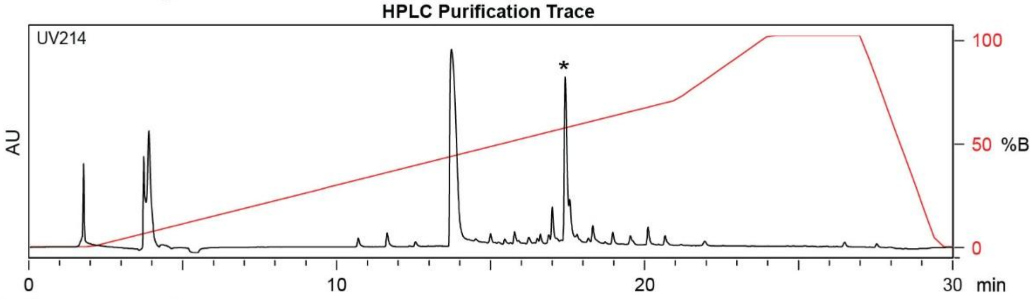

| Time (min) | Solvent B (%) |
|---|---|
| 0 | 0 |
| 2 | 0 |
| 22 | 70 |
| 24 | 100 |
| 27 | 100 |
| 29.5 | 0 |
| 30 | 0 |

Flow Rate: 13 mL / min
Phenomenex Kinetex EVO C18 Prep Column
(100 Å, 5 µm, 21.5 mm X 250 mm)

Solvent A: $H_2O$ + 0.1% (v/v) TFA
Solvent B: MeCN + 0.1% (v/v) TFA

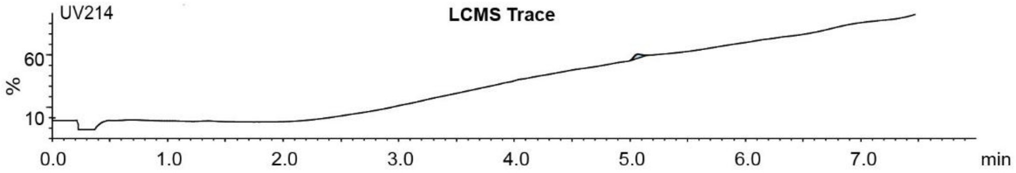

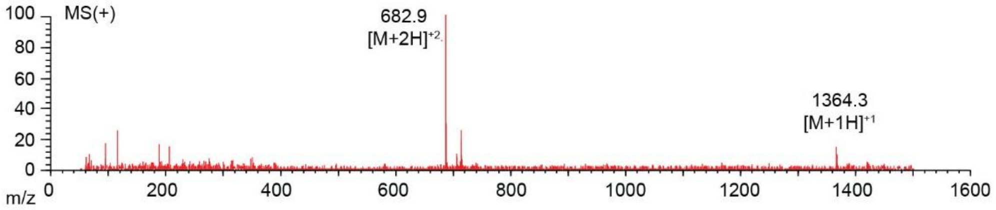

**Figure S44**. Data page for **3c**.

Expected: [M+1] = 1364.51, $[M+2]^{+2}$ = 682.755.

Observed: [M+1] = 1364.3, $[M+2]^{+2}$ = 682.9

**TSL6-SQCRPDQEIC-OH**

Scale: 10.3 mg
Yield: 1.1 mg (8.8%)
Purity: ~90%

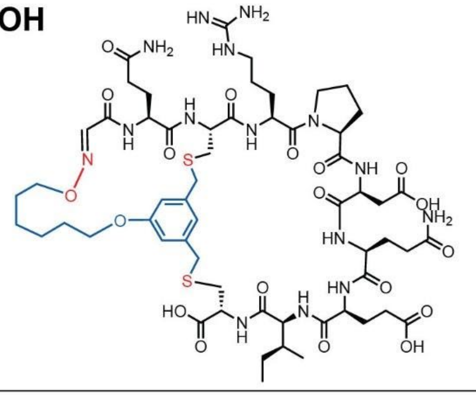

Chemical Formula: $C_{58}H_{87}N_{15}O_{19}S_2$
Exact Mass: 1361.57
Molecular Weight: 1362.54

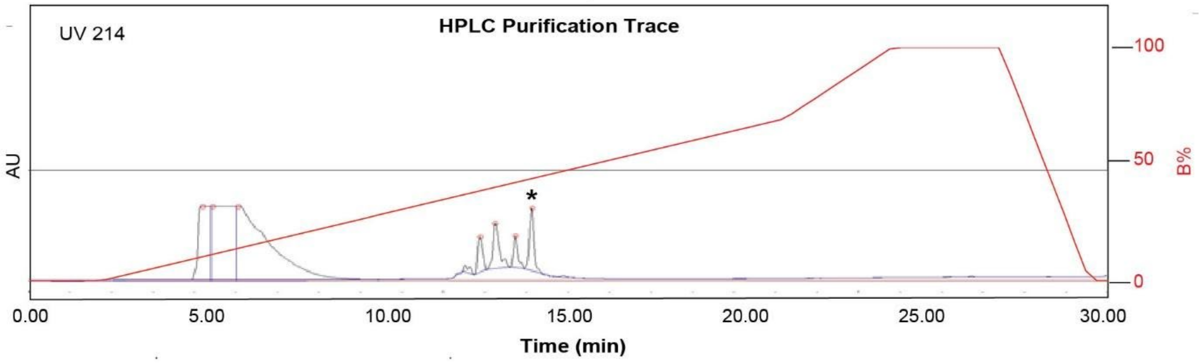

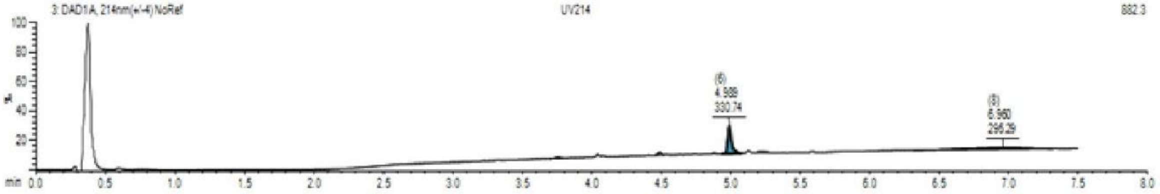

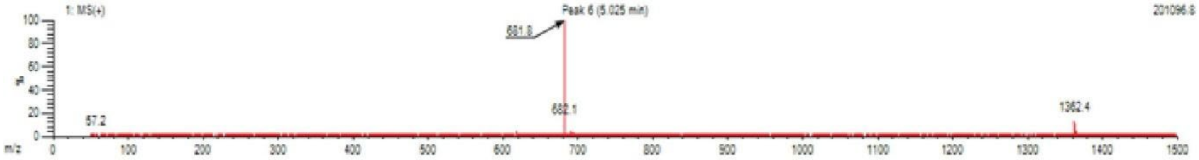

**Figure S45**. Data page for **4c**.

Expected: [M+1] = 1362.57, $[M+2]^{+2}$ = 681.785.

Observed: [M+1] = 1362.4, $[M+2]^{+2}$ = 681.8

### TSL6-SSCEIAIERC-OH

Starting material mass: 10.3 mg
Final product mass: 1.1 mg (8.8%)
Purity: >95%

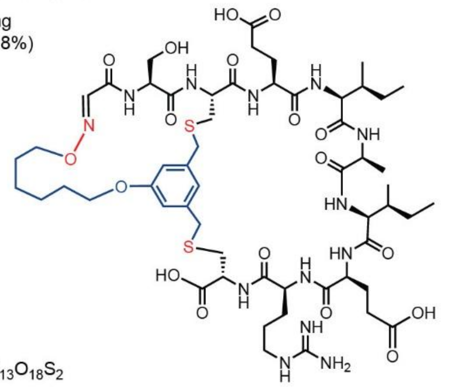

Chemical Formula: $C_{56}H_{87}N_{13}O_{18}S_2$
Exact Mass: 1293.57
Molecular Weight: 1294.51

**HPLC Purification Trace**

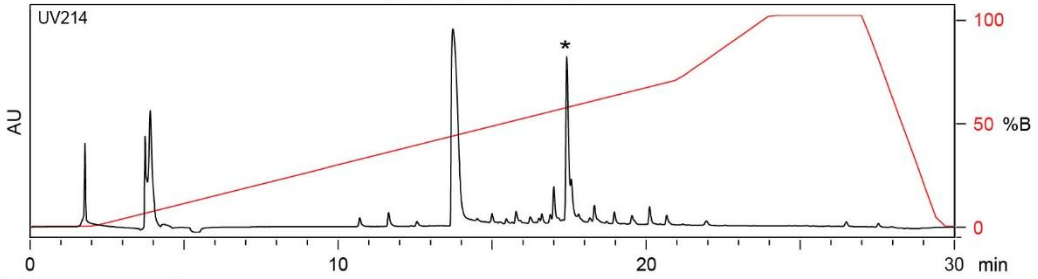

| Time (min) | Solvent B (%) |
|---|---|
| 0 | 0 |
| 2 | 0 |
| 22 | 70 |
| 24 | 100 |
| 27 | 100 |
| 29.5 | 0 |
| 30 | 0 |

Flow Rate: 13 mL / min
Phenomenex Kinetex EVO C18 Prep Column
(100 Å, 5 µm, 21.5 mm X 250 mm)

Solvent A: $H_2O$ + 0.1% (v/v) TFA
Solvent B: MeCN + 0.1% (v/v) TFA

**LCMS Trace**

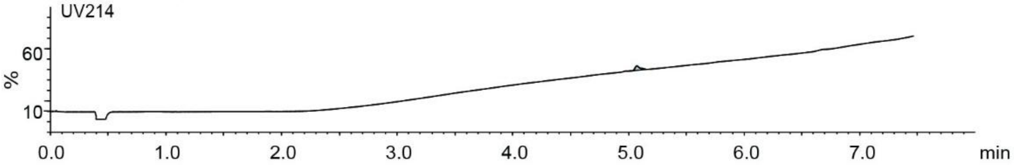

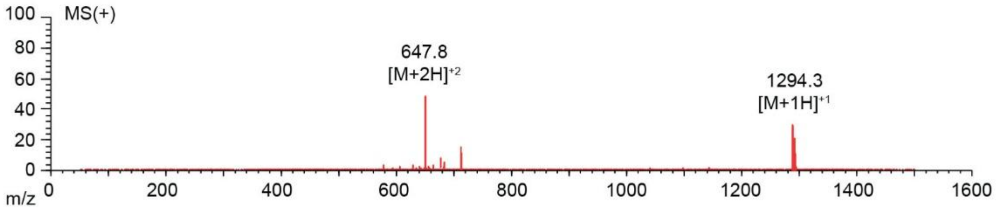

**Figure S46**. Data page for **5c**.

Expected: [M+1] = 1294.57, $[M+2]^{+2}$ = 647.785.

Observed: [M+1] = 1294.3, $[M+2]^{+2}$ = 647.8

### TSL6-SECTMEGTVC-OH

Scale: 10.9 mg
Yield: 0.1 mg (<1%)
Purity: ~80%

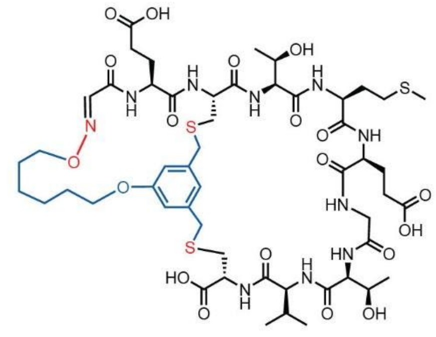

Chemical Formula: $C_{52}H_{78}N_{10}O_{19}S_3$
Exact Mass: 1242.46
Molecular Weight: 1243.43

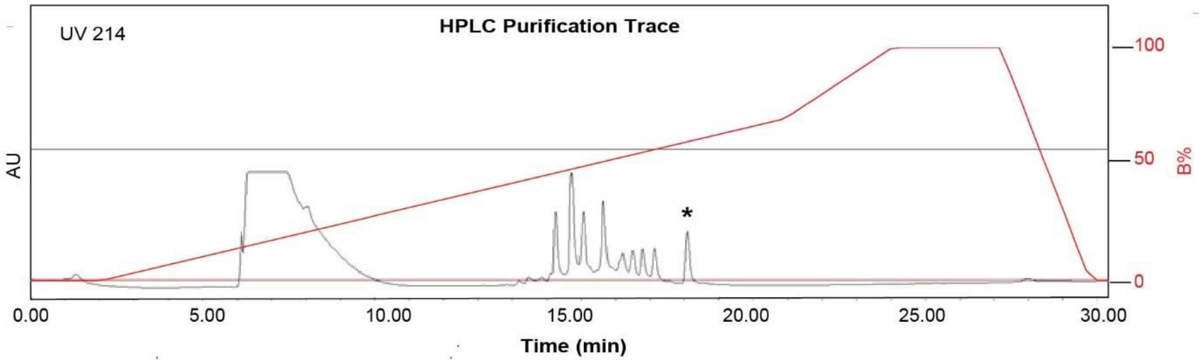

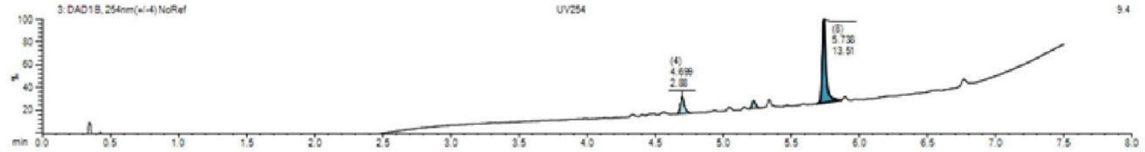

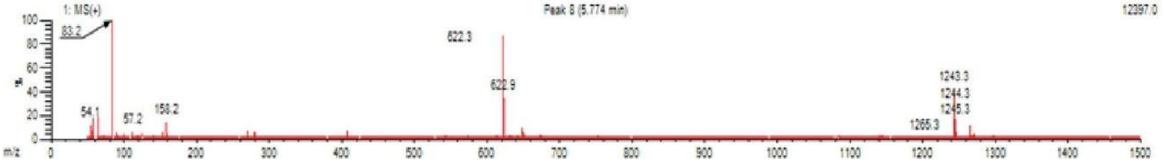

**Figure S47**. Data page for **6c**.

Expected: [M+1] = 1243.46, $[M+2]^{+2}$ = 622.23.

Observed: [M+1] = 1243.3, $[M+2]^{+2}$ = 622.3

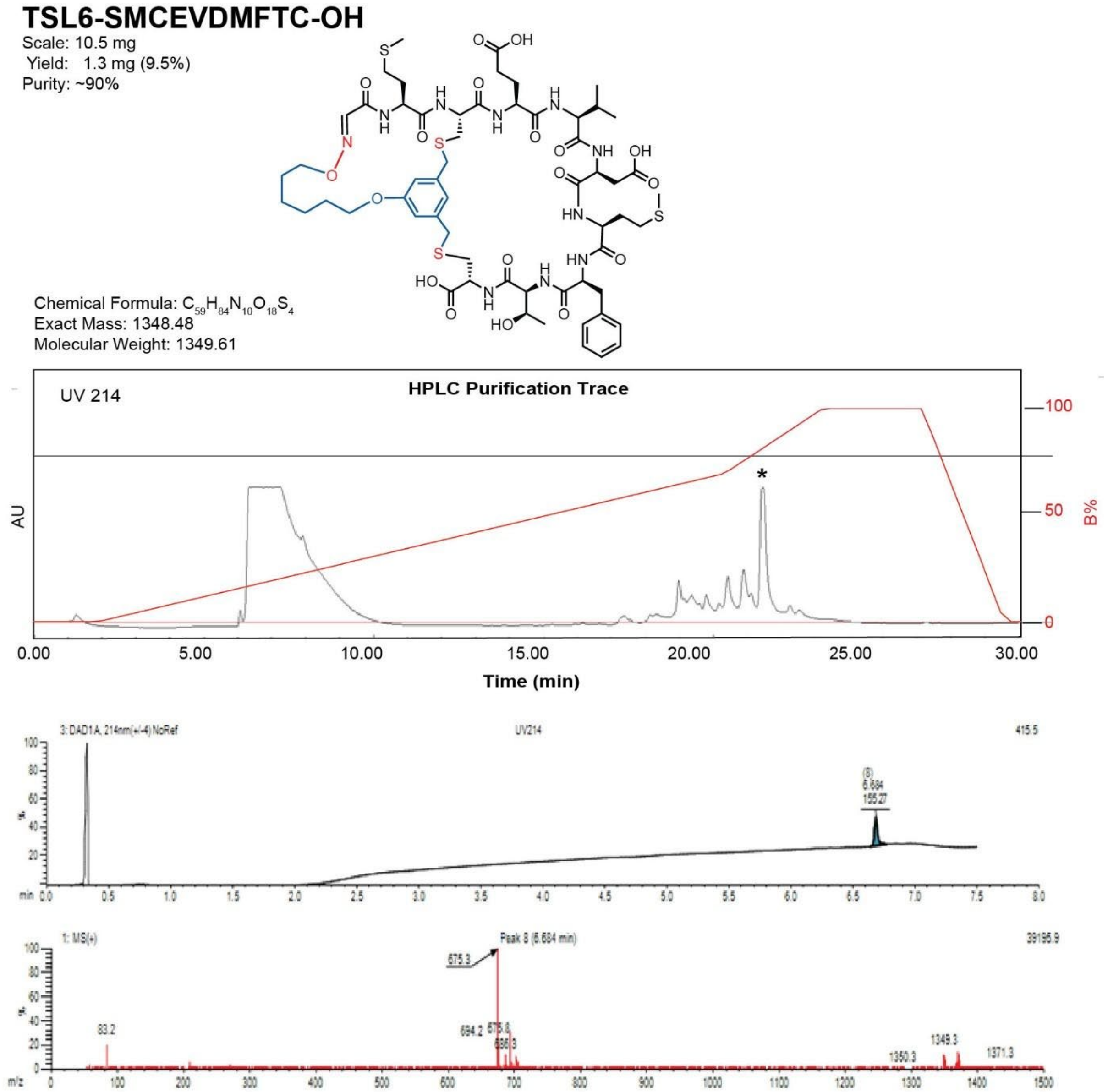

**Figure S48.** Data page for **7c**.

Expected: [M+1] = 1349.48, $[M+2]^{+2}$ = 675.24.

Observed: [M+1] = 1349.3, $[M+2]^{+2}$ = 675.3

**TSL6-SWCRPATVNC-OH**

Scale: 18.7 mg
Yield: 4.5 mg (23%)
Purity: >95%

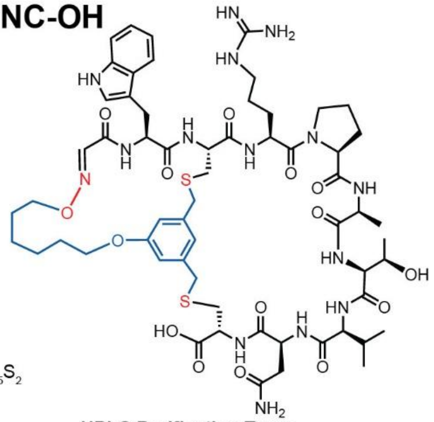

Chemical Formula: $C_{60}H_{85}N_{15}O_{15}S_{2}$
Exact Mass: 1319.58
Molecular Weight: 1320.55

**HPLC Purification Trace**

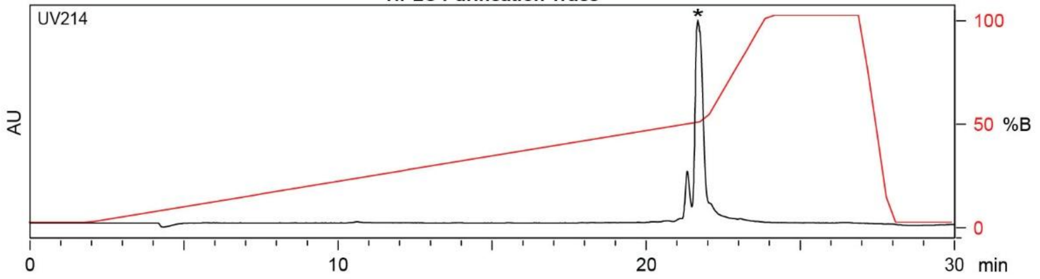

| Time (min) | Solvent B (%) |
|---|---|
| 0 | 2 |
| 2 | 2 |
| 22 | 50 |
| 24 | 100 |
| 27 | 100 |
| 28 | 2 |
| 30 | 2 |

Flow Rate: 13 mL / min
Phenomenex Kinetex EVO C18 Prep Column
(100 Å, 5 µm, 21.5 mm X 250 mm)

Solvent A: $H_2O$ + 0.1% (v/v) TFA
Solvent B: MeCN + 0.1% (v/v) TFA

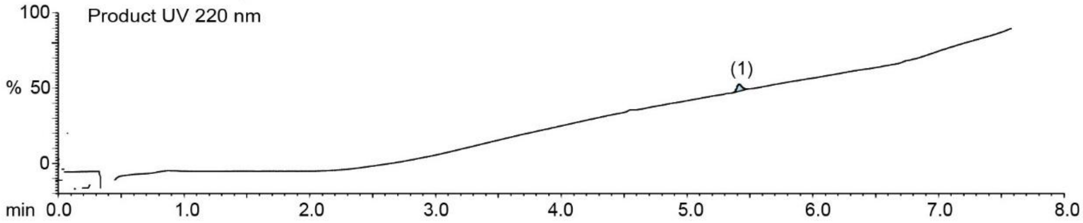

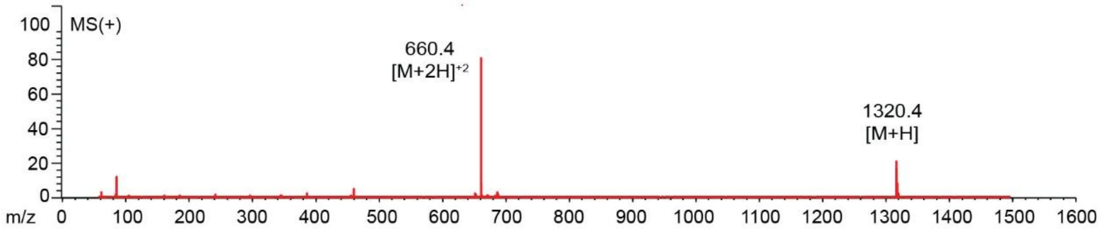

**Figure S49**. Data page for **8c**.

Expected: [M+1] = 1320.58, $[M+2]^{+2}$ = 660.79.

Observed: [M+1] = 1320.4, $[M+2]^{+2}$ = 660.4

**TSL1-SWCRPATVNC-$NH_2$**

Scale: 11.3 mg
Yield: 2.6 mg (23%)
Purity: >95%

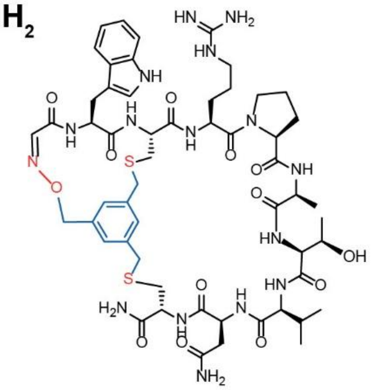

Chemical Formula: $C_{55}H_{76}N_{16}O_{13}S_2$
Exact Mass: 1232.52
Molecular Weight: 1233.43

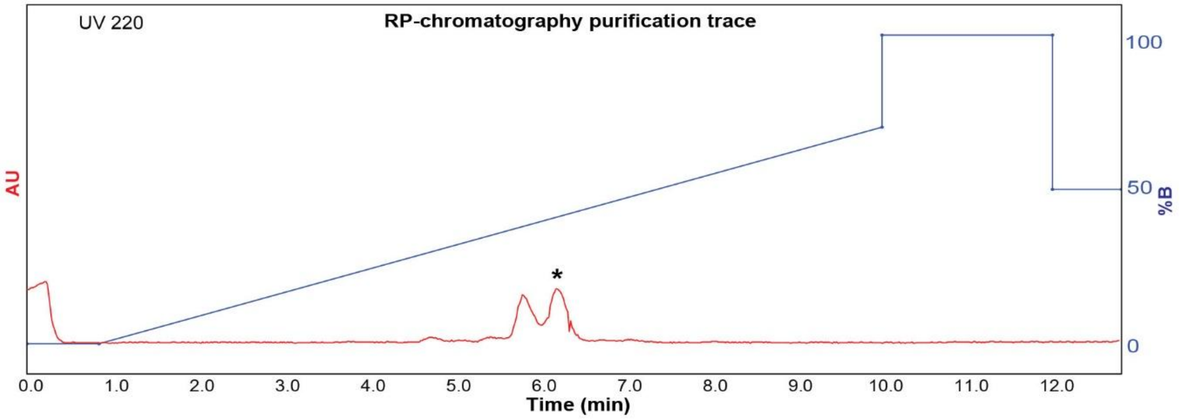

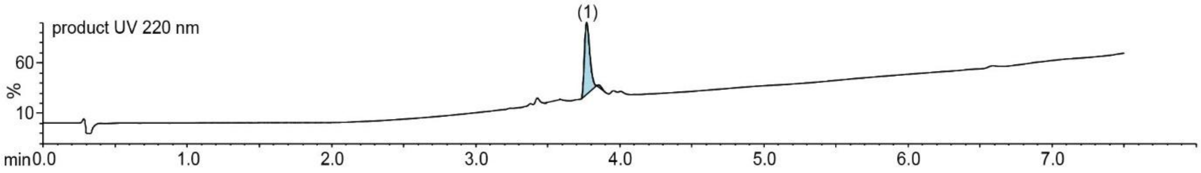

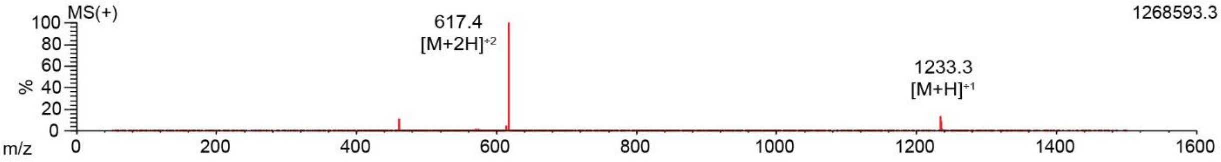

**Figure S50**. Data page for **8e**.

Expected: [M+1] = 1233.52, $[M+2]^{+2}$ = 617.26.

Observed: [M+1] = 1233.3, $[M+2]^{+2}$ = 617.4

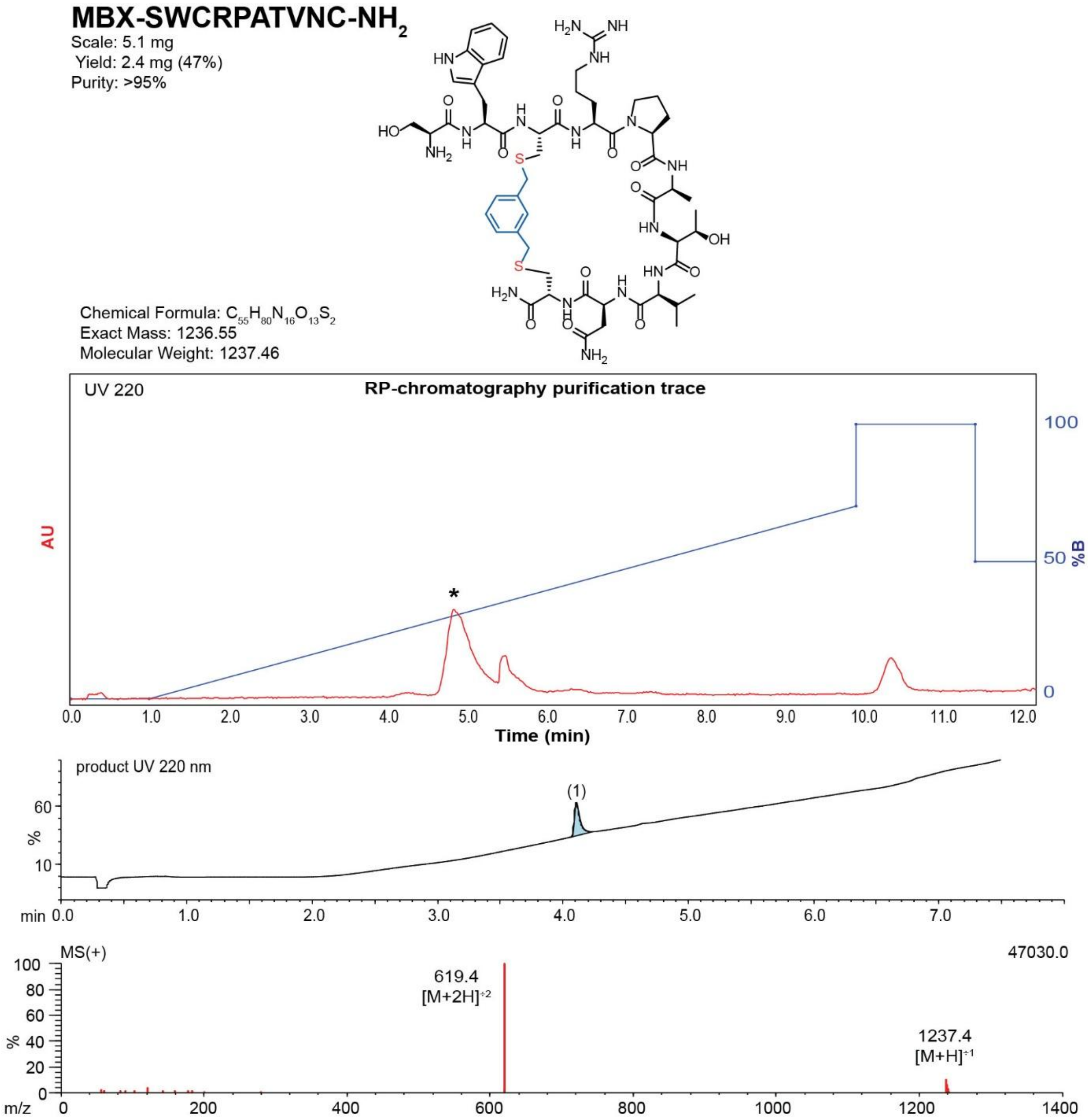

**Figure S51**. Data page for **8f**.

Expected: [M+1] = 1237.55, $[M+2]^{+2}$ = 619.275.

Observed: [M+1] = 1237.4, $[M+2]^{+2}$ = 619.4

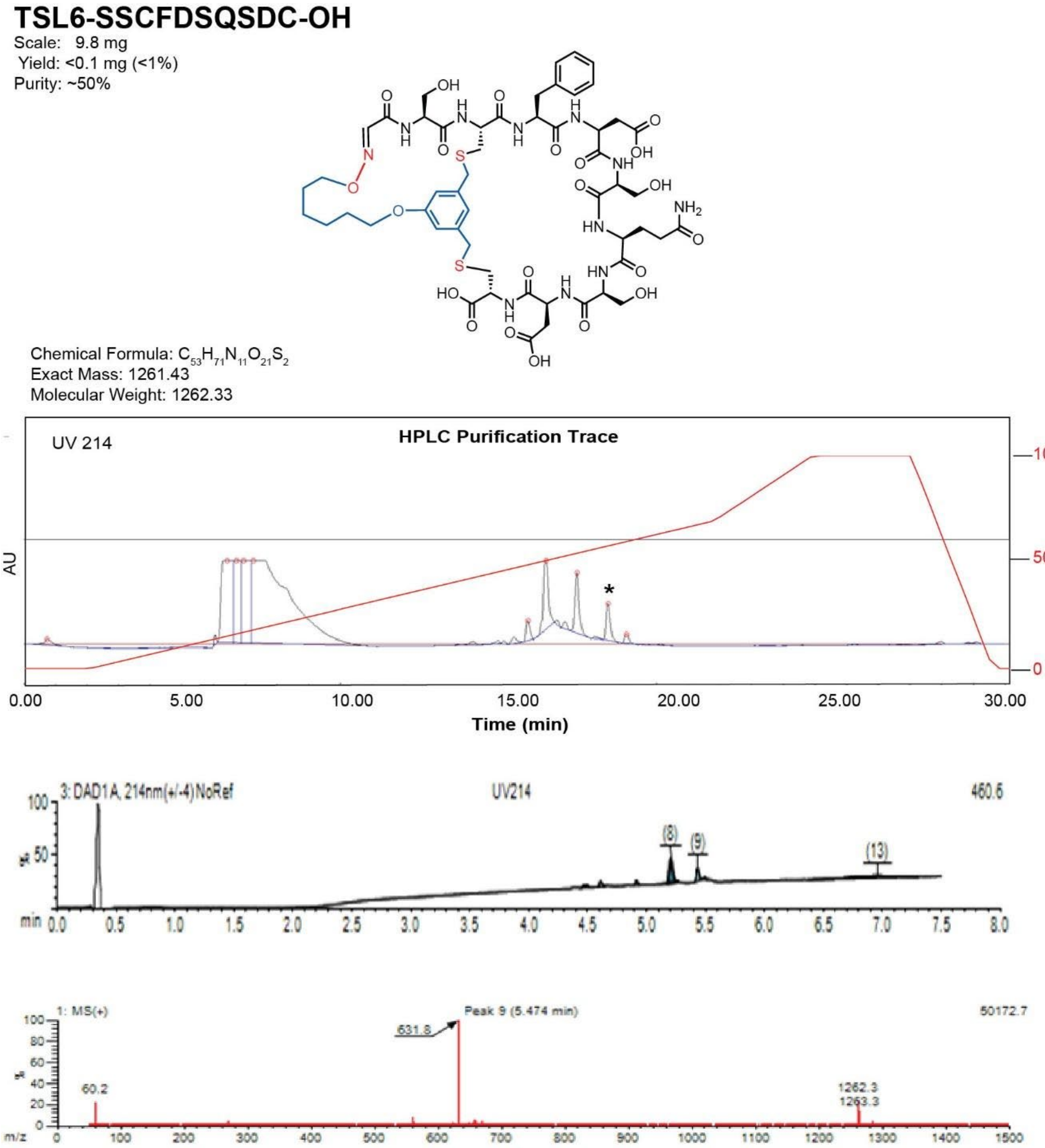

**Figure S52**. Data page for **9c**.

Expected: [M+1] = 1262.43, $[M+2]^{+2}$ = 631.72.

Observed: [M+1] = 1262.3, $[M+2]^{+2}$ = 631.8

**TSL6-SDCPQATIIC-OH**

Scale: 9.8 mg
Yield: 1.3 mg (11%)
Purity: ~80%

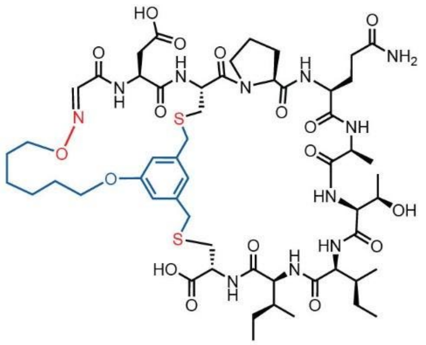

Chemical Formula: $C_{55}H_{83}N_{11}O_{17}S_2$
Exact Mass: 1233.54
Molecular Weight: 1234.45

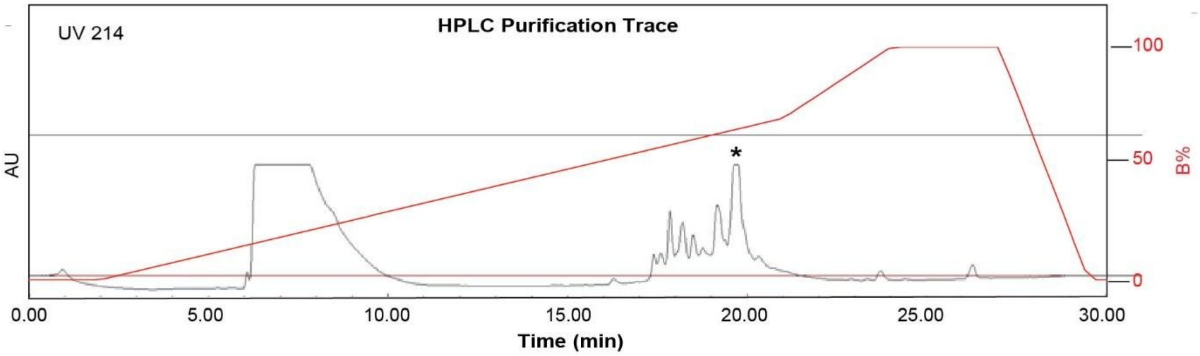

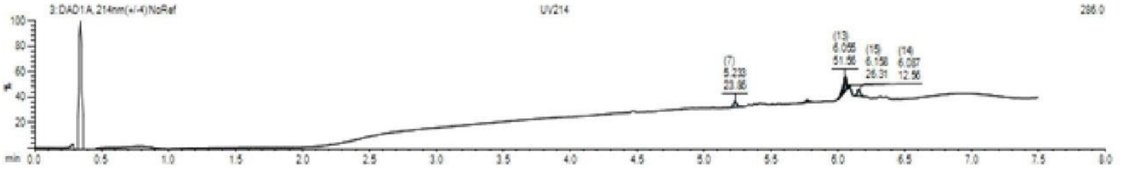

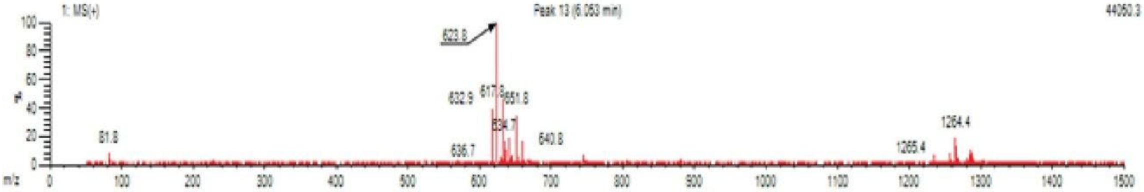

**Figure S53**. Data page for **10c**.

Expected: [M+1] = 1234.54, $[M+2]^{+2}$ = 617.77.

Observed: [M+1] = N/A, $[M+2]^{+2}$ = 617.9

**TSL6-SYCHDTAGRC-OH**

Scale: 9.6 mg

Yield: 1.0 mg (8.9%)

Purity: >95%

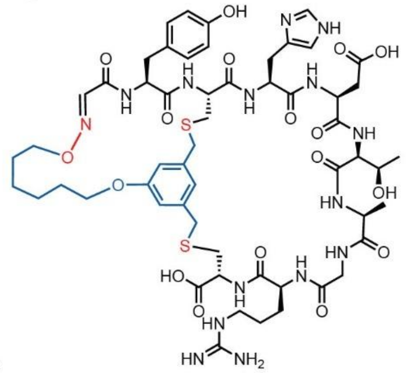

Chemical Formula: $C_{56}H_{77}N_{15}O_{17}S_2$

Exact Mass: 1295.51

Molecular Weight: 1296.44

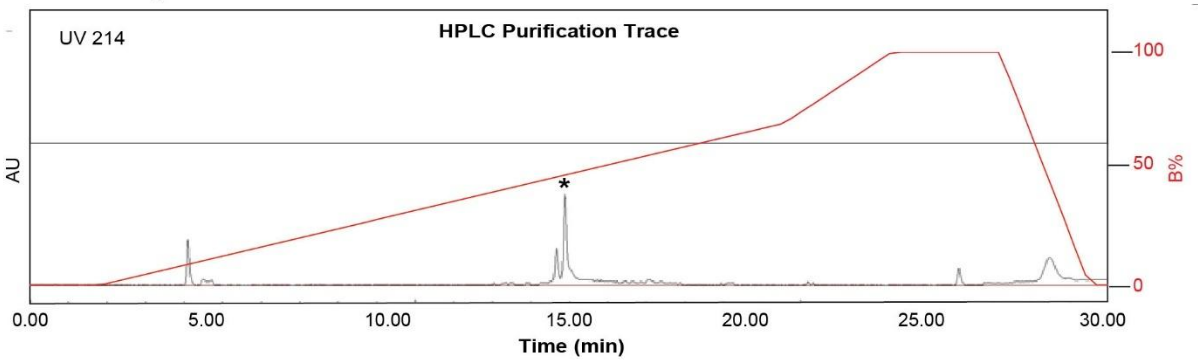

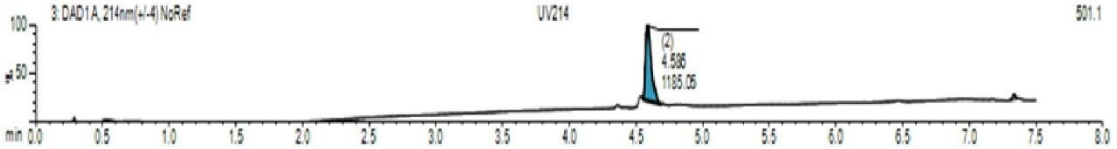

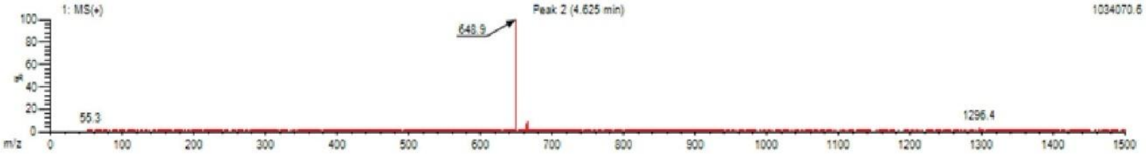

**Figure S54**. Data page for **11c**.

Expected: [M+1] = 1296.51, $[M+2]^{+2}$ = 648.76.

Observed: [M+1] = 1296.4, $[M+2]^{+2}$ = 648.9

**TSL6-SFCHYPTHVC-NH$_2$**

Scale: 5.3 mg
Yield: 1.8 mg (31%)
Purity: >95%

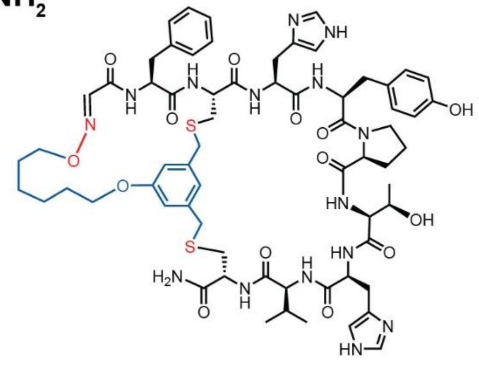

Chemical Formula: $C_{66}H_{85}N_{15}O_{14}S_2$
Exact Mass: 1375.58
Molecular Weight: 1376.62

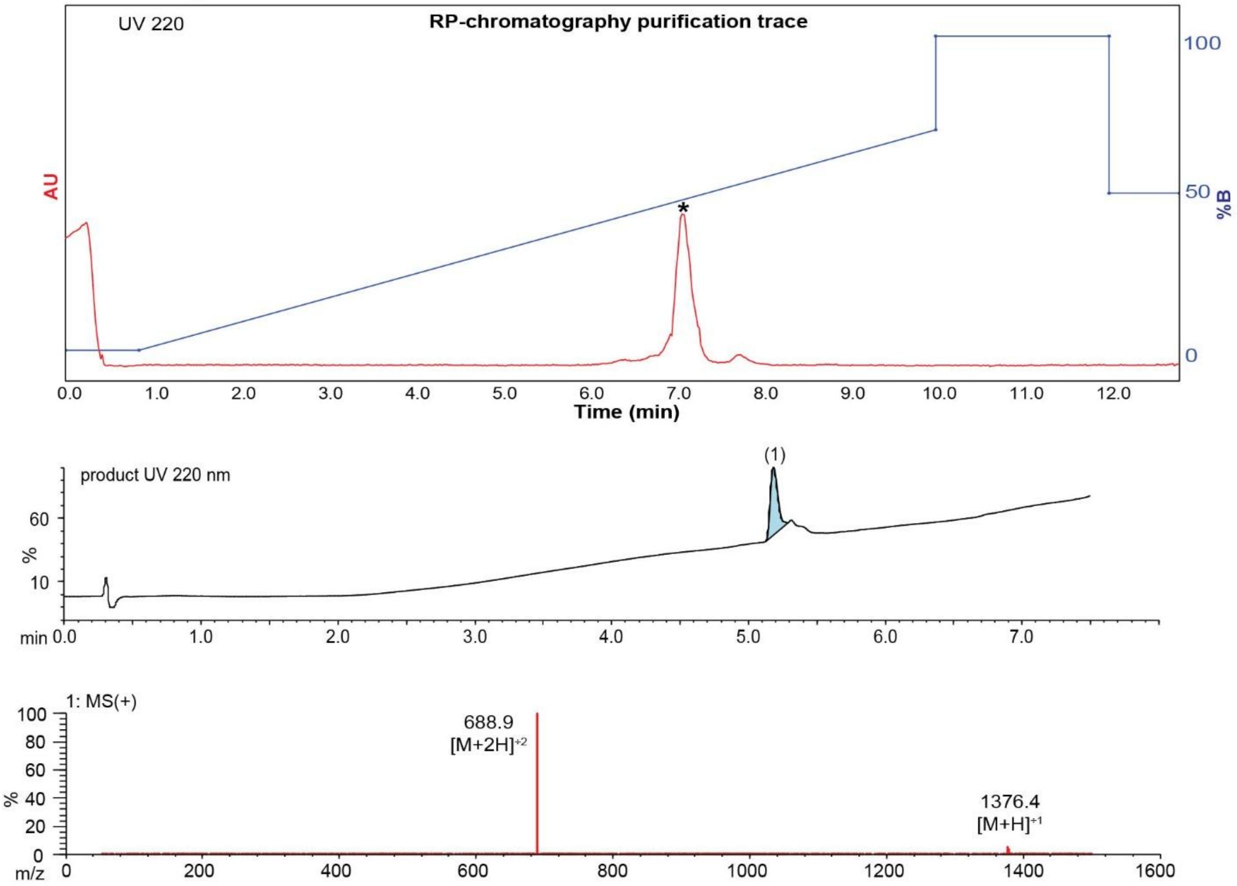

**Figure S55**. Data page for **12c**.

Expected: [M+1] = 1376.58, $[M+2]^{+2}$ = 688.79.

Observed: [M+1] = 1376.4, $[M+2]^{+2}$ = 688.9

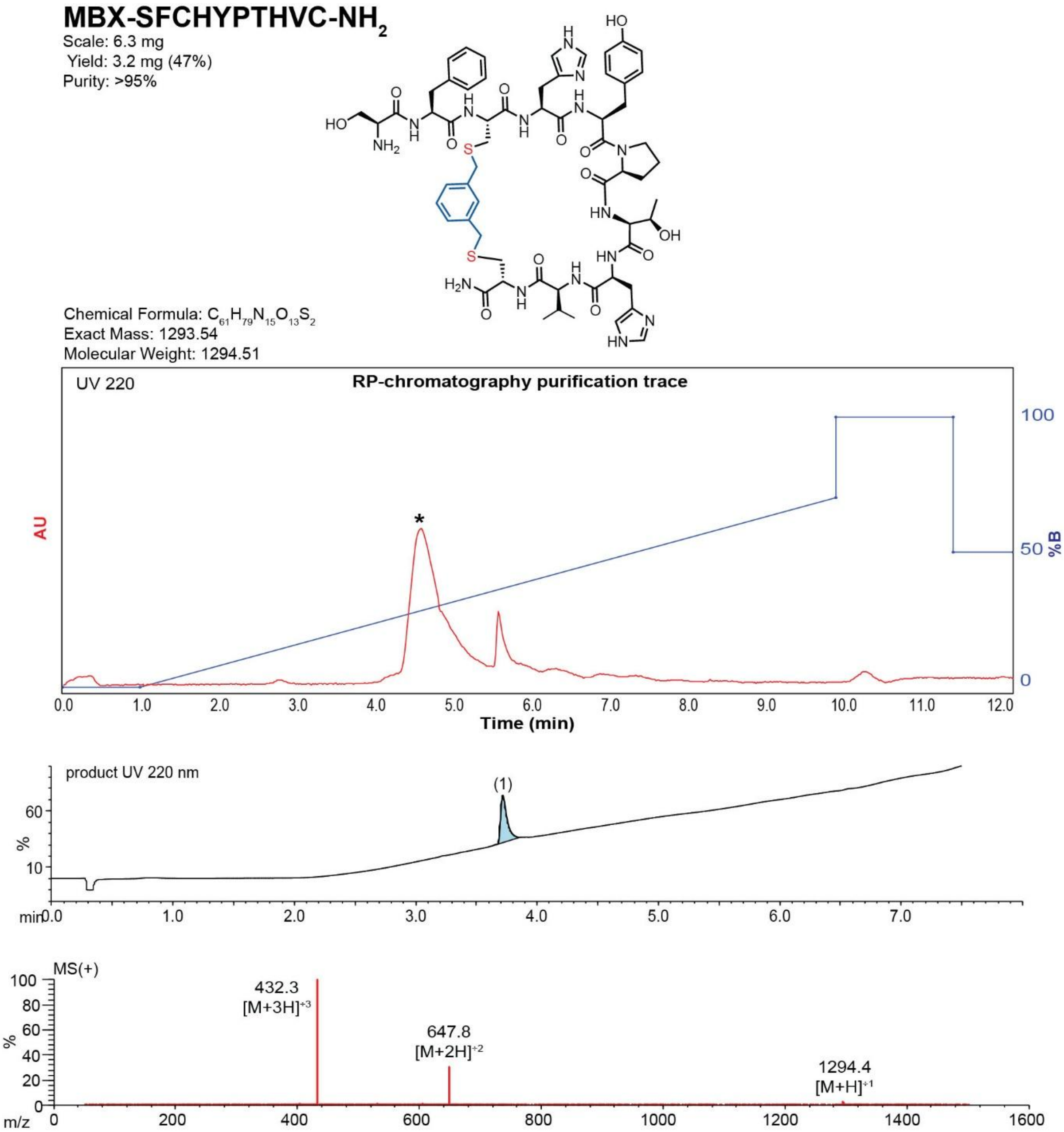

**Figure S56**. Data page for **12f.**

Expected: [M+1] = 1294.54, $[M+2]^{+2}$ = 647.77.

Observed: [M+1] = 1294.4, $[M+2]^{+2}$ = 647.8

### TSL6-SICTGEIEHC-OH

Scale: 9.7 mg
Yield: 0.9 mg (7.9%)
Purity: ~80%

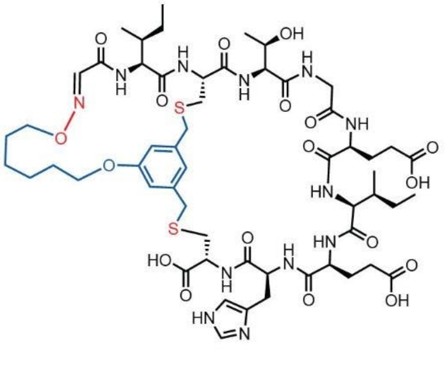

Chemical Formula: $C_{56}H_{82}N_{12}O_{18}S_2$
Exact Mass: 1274.53
Molecular Weight: 1275.46

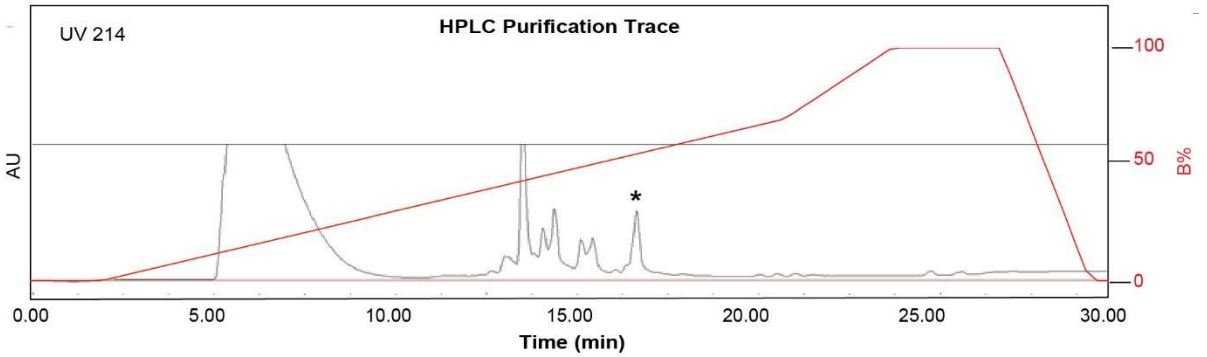

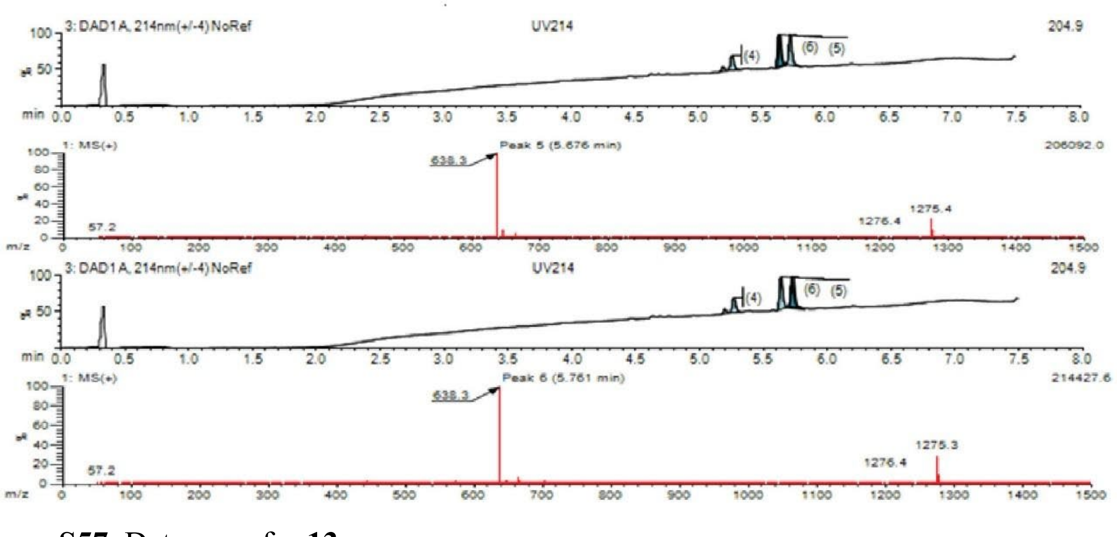

**Figure S57**. Data page for **13c**.

Expected: [M+1] = 1275.53, $[M+2]^{+2}$ = 638.265.

Observed: [M+1] = 1275.3, $[M+2]^{+2}$ = 638.3

### TSL6-SGCIIEGMQC-OH

Scale: 9.8 mg
Yield: 0.9 mg (7.8%)
Purity: N/A

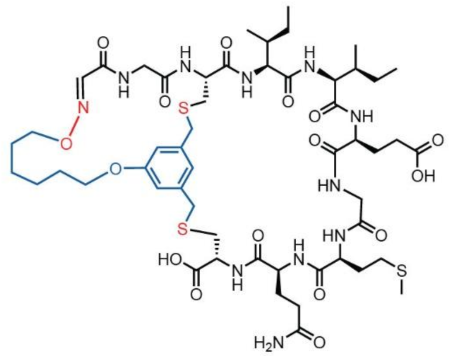

Chemical Formula: $C_{53}H_{81}N_{11}O_{16}S_3$
Exact Mass: 1223.50
Molecular Weight: 1224.47

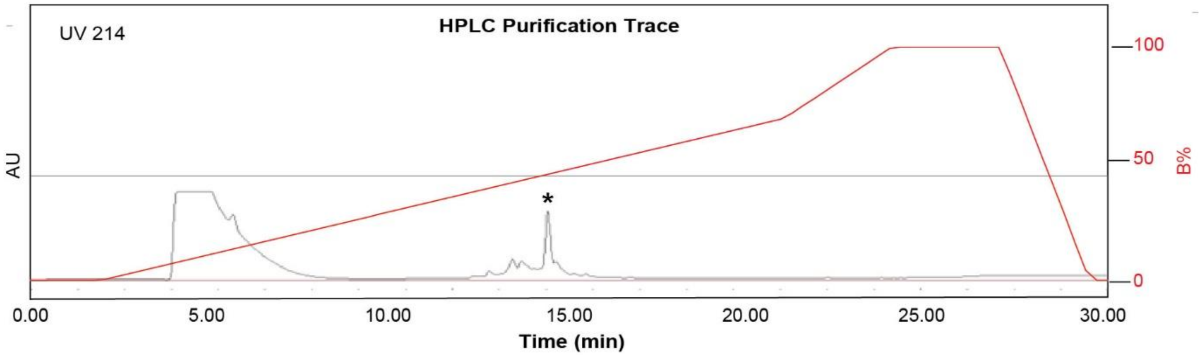

**Figure S58.** Data page for **14c**.

Expected: [M+1] = 1224.50, $[M+2]^{\div 2}$ = 612.75.

Observed: [M+1] = N/A, $[M+2]^{\div 2}$ = N/A.

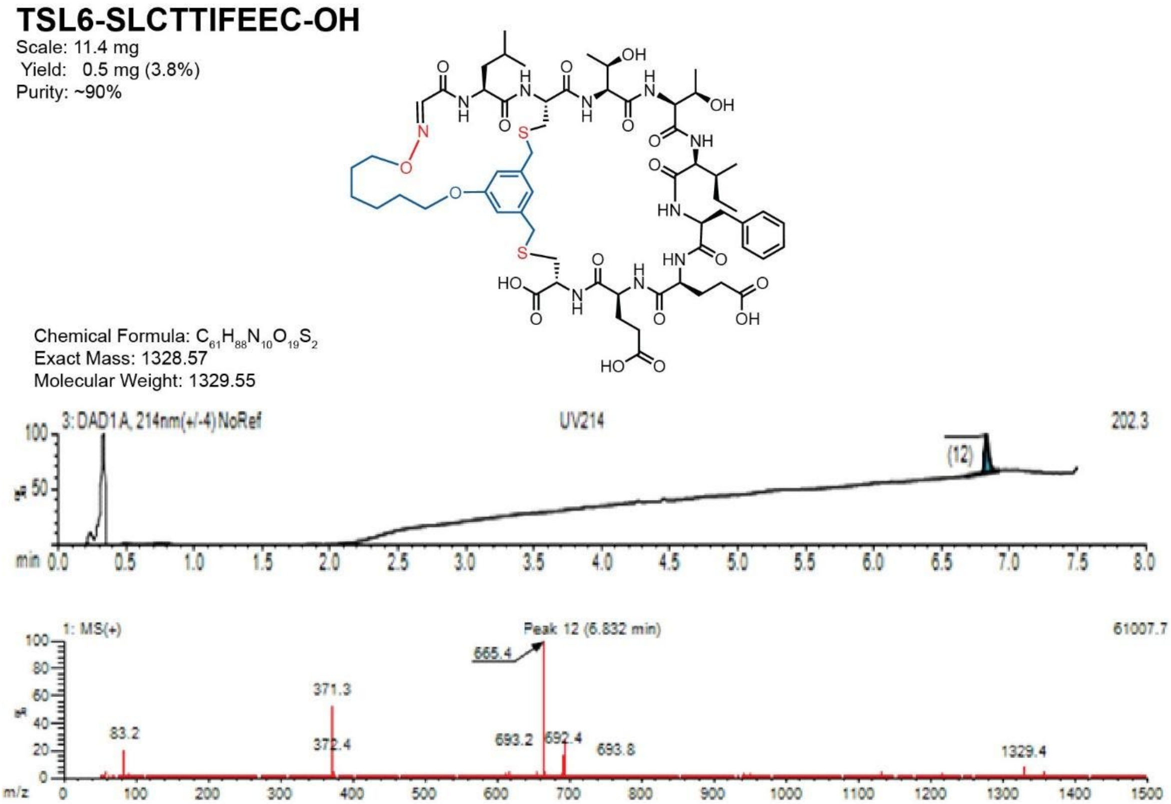

**Figure S59**. Data page for **16c**.

Expected: [M+1] = 1329.57, $[M+2]^{\div 2}$ = 665.285.

Observed: [M+1] = 1329.4, $[M+2]^{\div 2}$ = 665.4

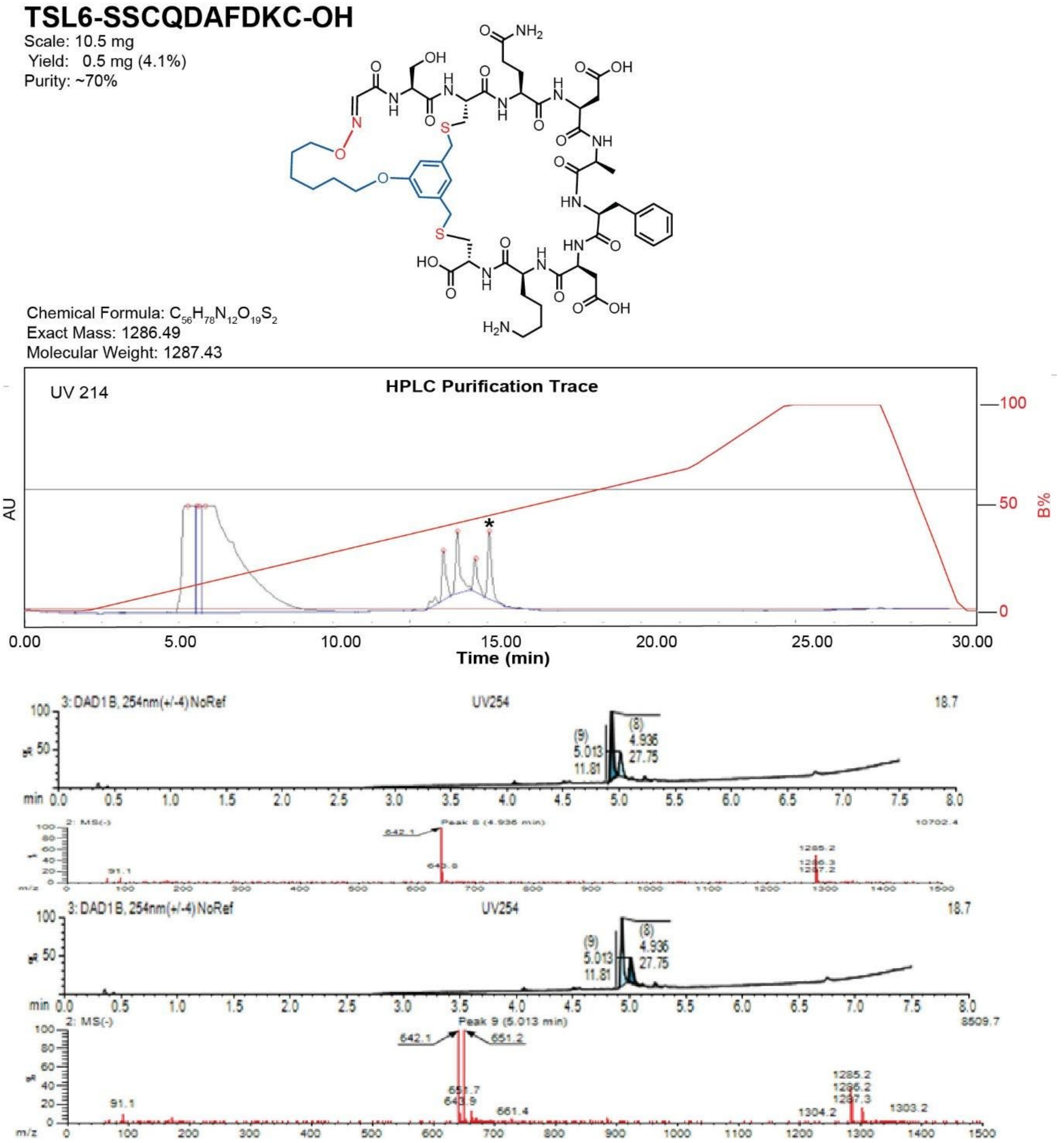

**Figure S60**. Data page for **17c**.

Expected: [M+1] = 1287.49, $[M+2]^{\div 2}$ = 644.245.

Observed: [M+1] = 1287.3, $[M+2]^{\div 2}$ = 643.9

**TSL6-SDCPWQSKTC-OH**

Scale: 9.2 mg
Yield: 3.4 mg (32%)
Purity: >95%

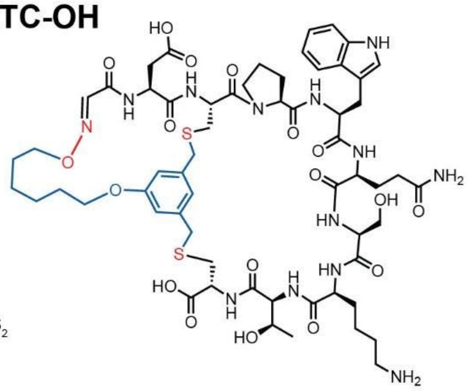

Chemical Formula: $C_{60}H_{83}N_{13}O_{18}S_2$
Exact Mass: 1337.54
Molecular Weight: 1338.52

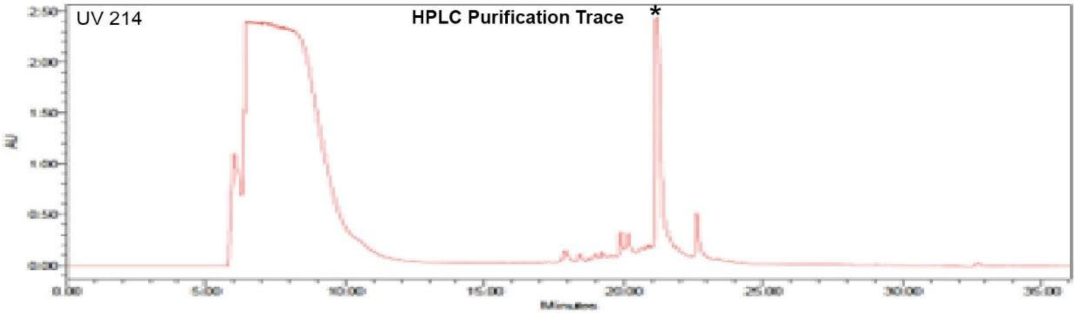

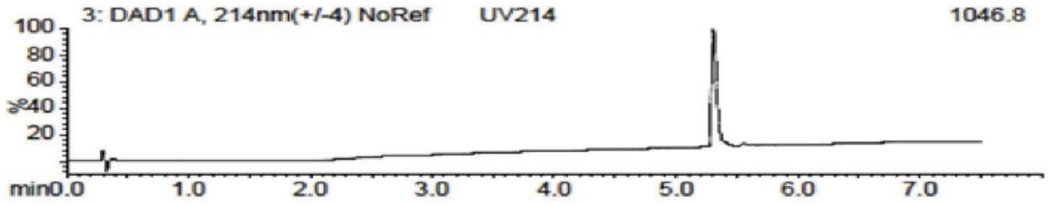

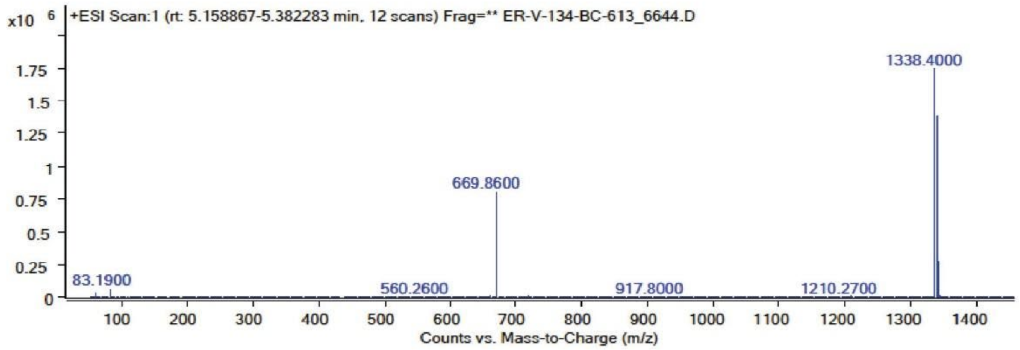

**Figure S61**. Data page for **18c**.

Expected: [M+1] = 1338.54, $[M+2]^{+2}$ = 669.77.

Observed: [M+1] = 1338.40, $[M+2]^{+2}$ = 669.86

### TSL6-SICPDGSNDC-OH

Scale: 9.8 mg
Yield: 4.2 mg (36%)
Purity: >95%

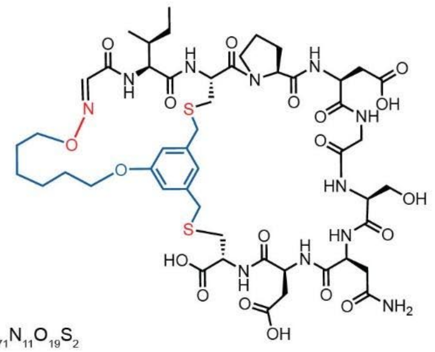

Chemical Formula: $C_{50}H_{71}N_{11}O_{19}S_2$
Exact Mass: 1193.44
Molecular Weight: 1194.30

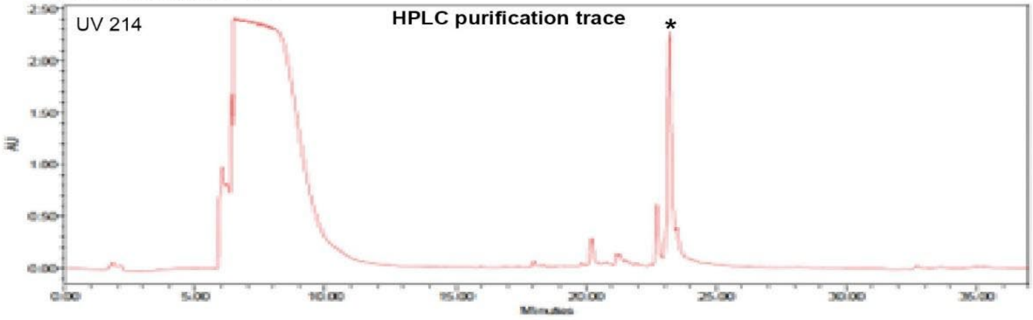

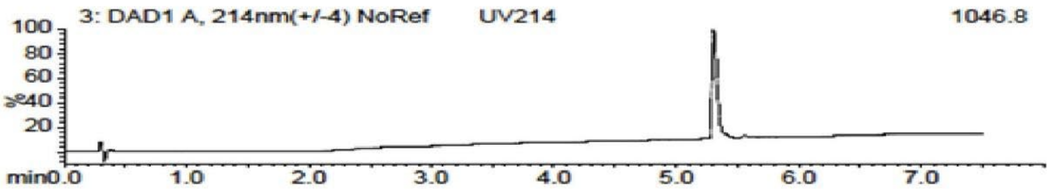

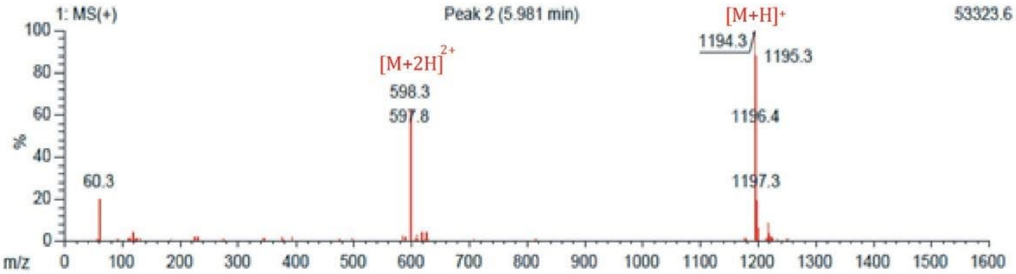

**Figure S62**. Data page for **19c**.

Expected: [M+1] = 1194.44, $[M+2]^{+2}$ = 597.72.

Observed: [M+1] = 1194.3, $[M+2]^{+2}$ = 597.8

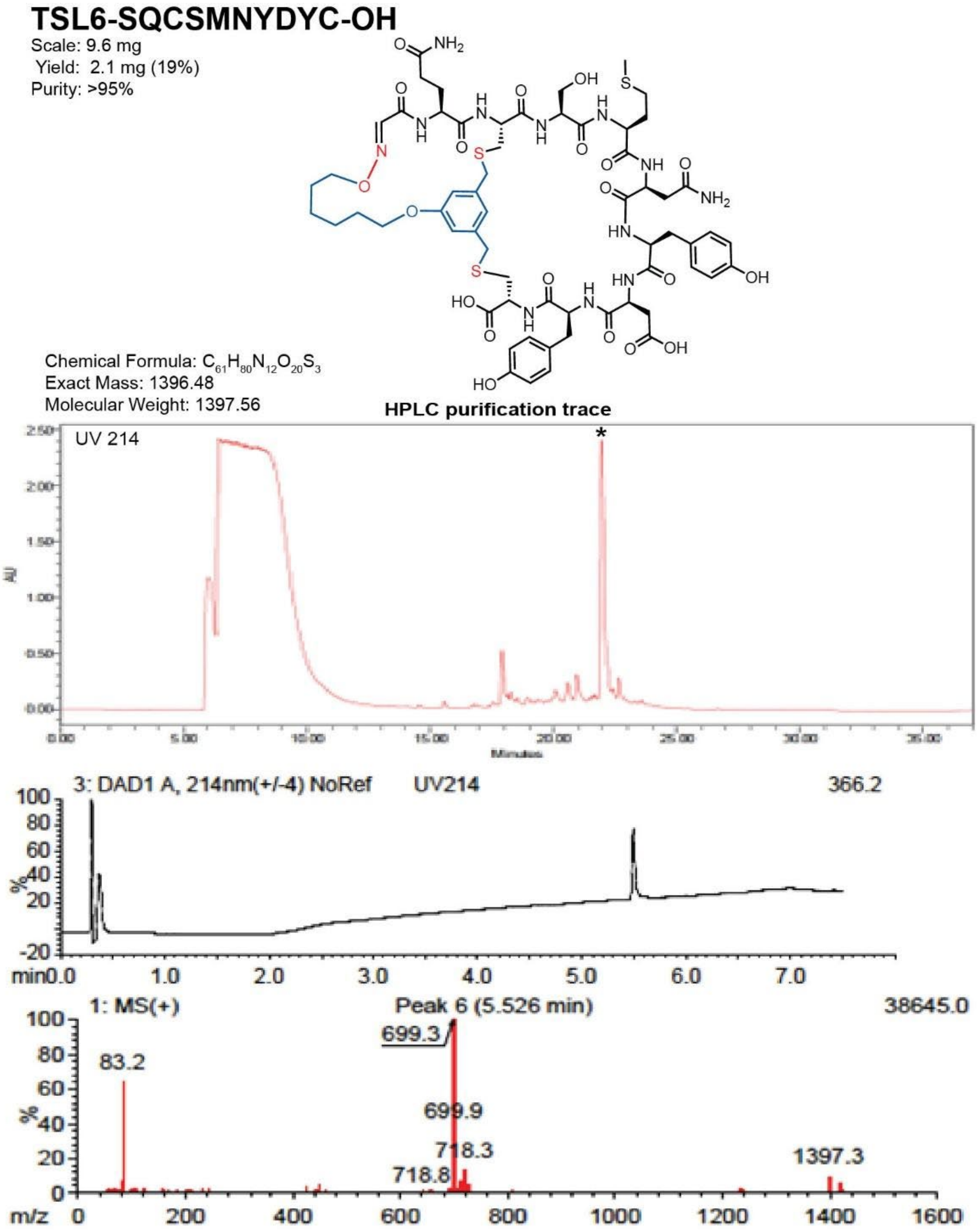

**Figure S63**. Data page for **20c**.

Expected: [M+1] = 1397.48, $[M+2]^{+2}$ = 699.24.

Observed: [M+1] = 1397.3, $[M+2]^{+2}$ = 699.3

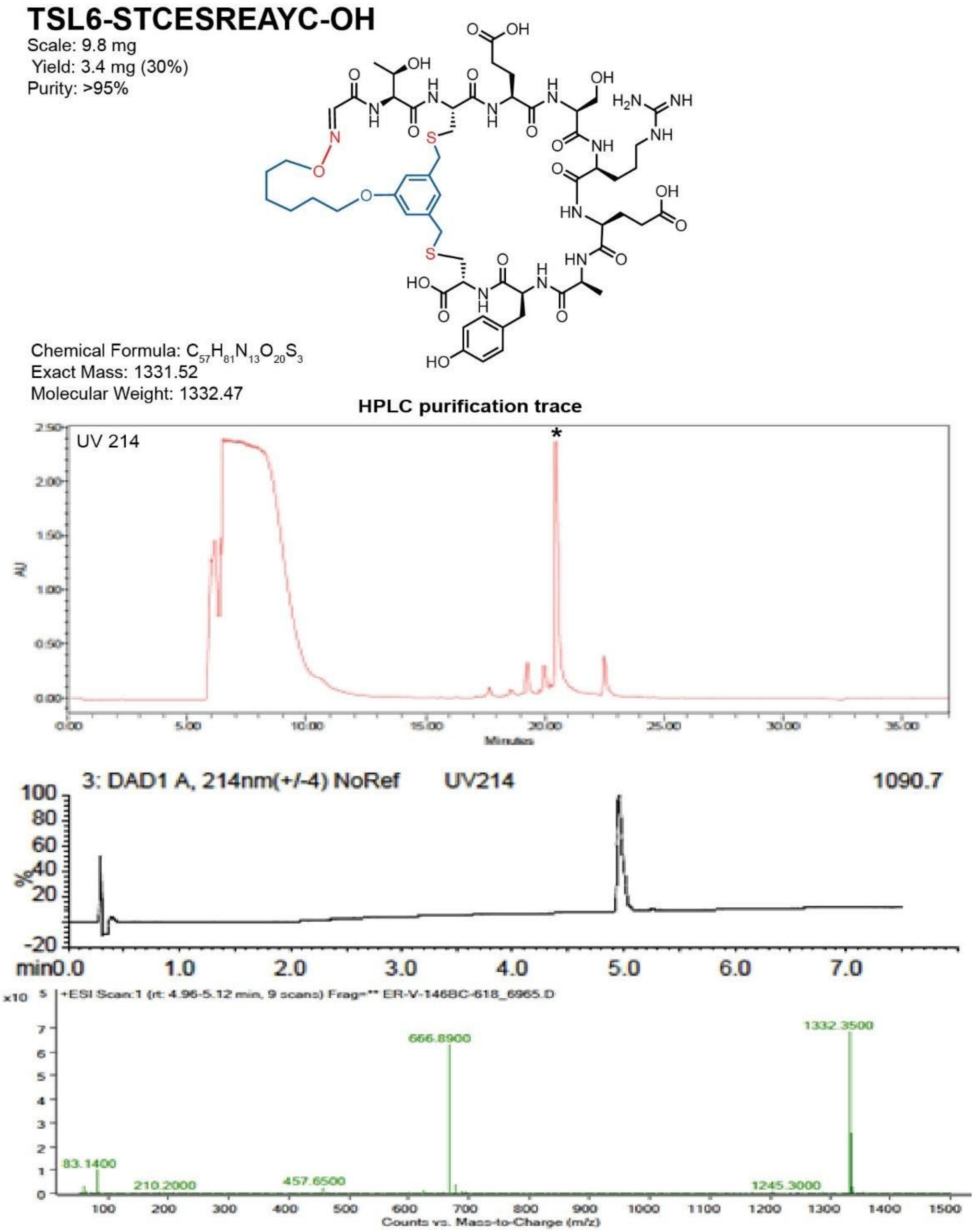

**Figure S64**. Data page for **21c**.

Expected: [M+1] = 1332.52, $[M+2]^{+2}$ = 666.76.

Observed: [M+1] = 1332.35, $[M+2]^{+2}$ = 666.89

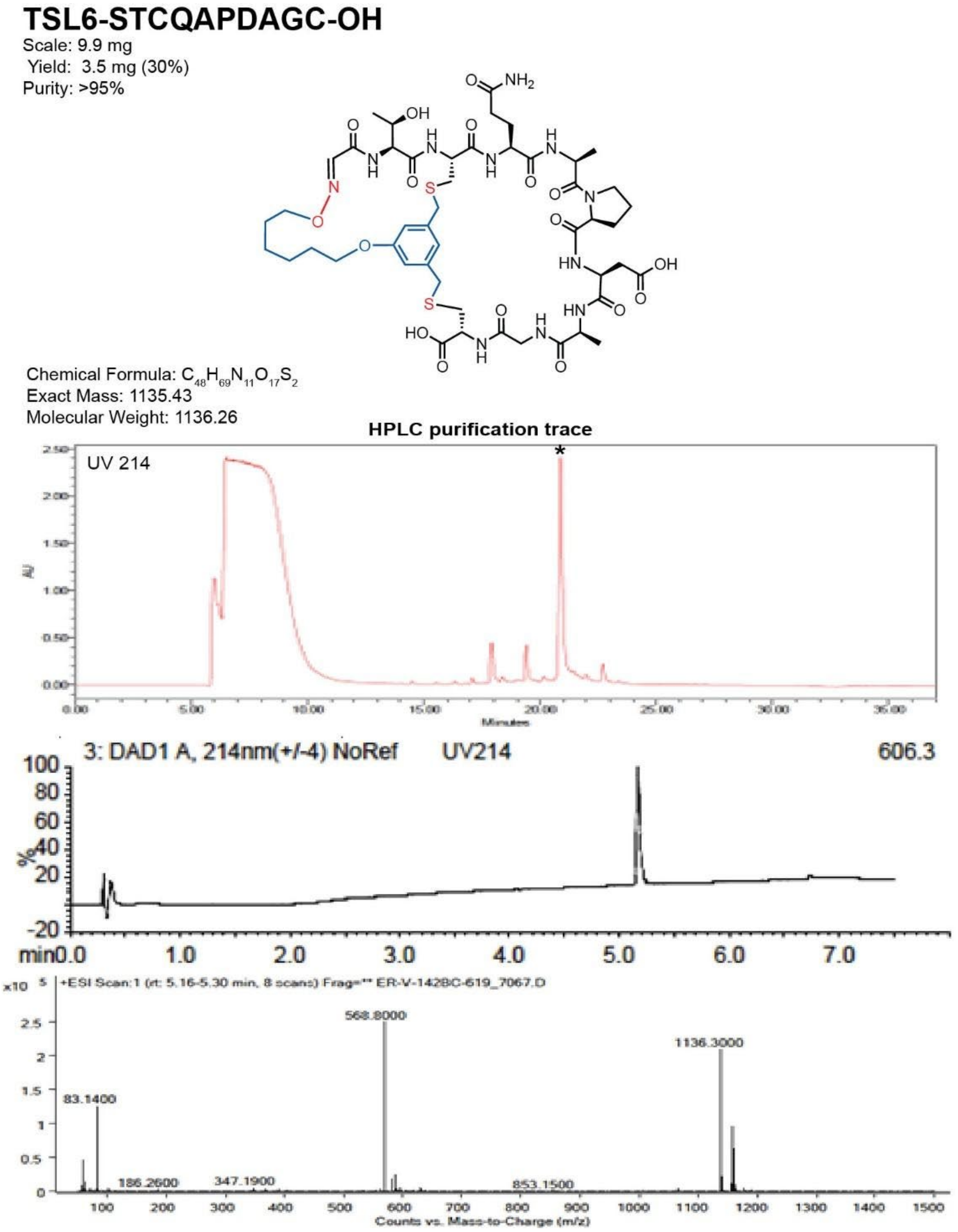

**Figure S65**. Data page for **22c**.

Expected: [M+1] = 1136.43, $[M+2]^{+2}$ = 568.715.

Observed: [M+1] = 1136.3, $[M+2]^{+2}$ = 568.8

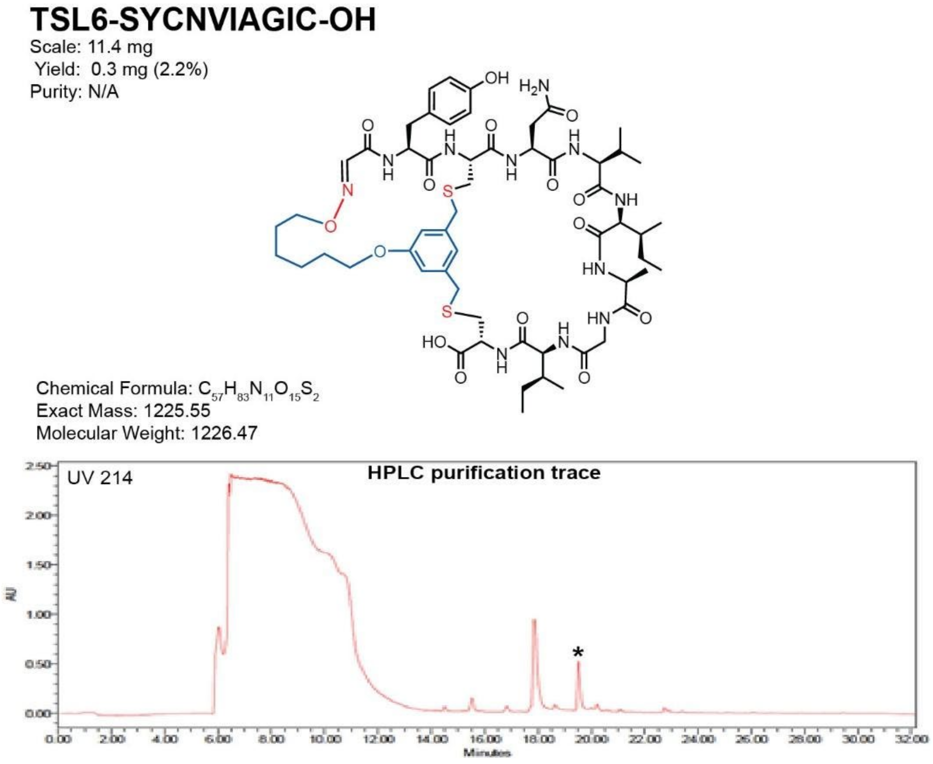

**Figure S66**. Data page for **23c**.

Expected: $[M+1] = 1226.54$, $[M+2]^{\div 2} = 613.77$

Observed: $[M+1]$ = N/A, $[M+2]^{\div 2}$ = N/A

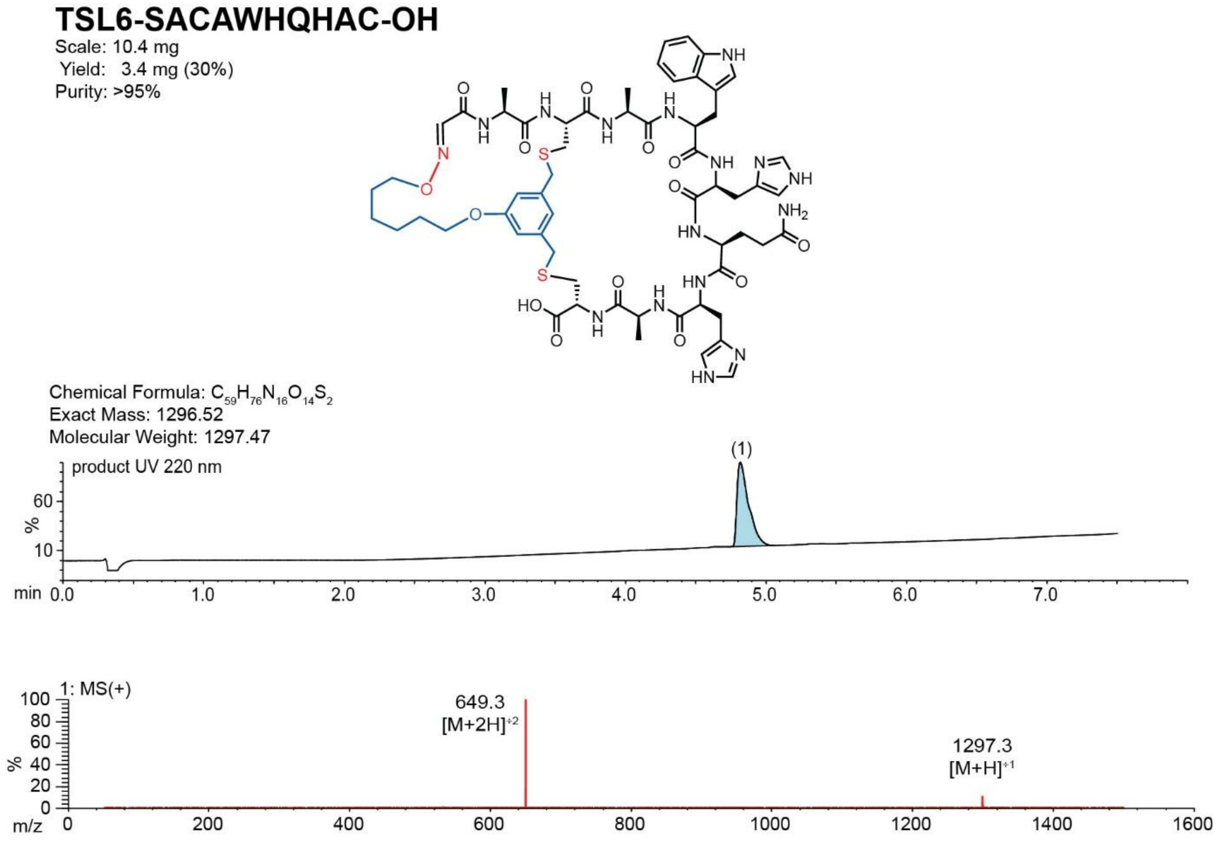

**Figure S67**. Data page for **24c**.

Expected: [M+1] = 1297.52, $[M+2]^{+2}$ = 649.26.

Observed: [M+1] = 1297.3, $[M+2]^{+2}$ = 649.3

**TSL6-SACSDTGLQC-OH**

Scale: 10.1 mg
Yield: 2.9 mg (28%)
Purity: >95%

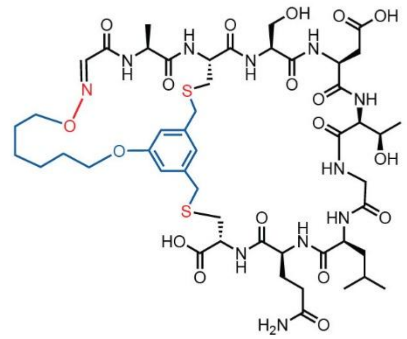

Chemical Formula: $C_{49}H_{73}N_{11}O_{18}S_2$
Exact Mass: 1167.46
Molecular Weight: 1168.3

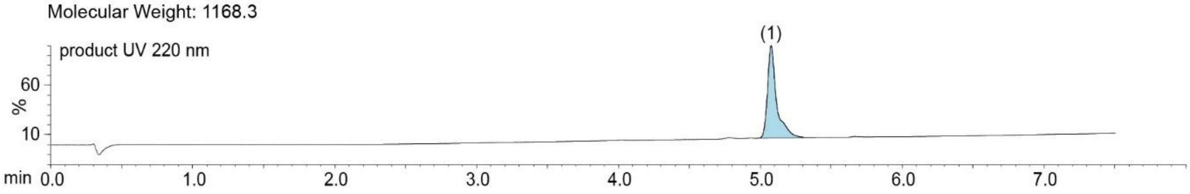

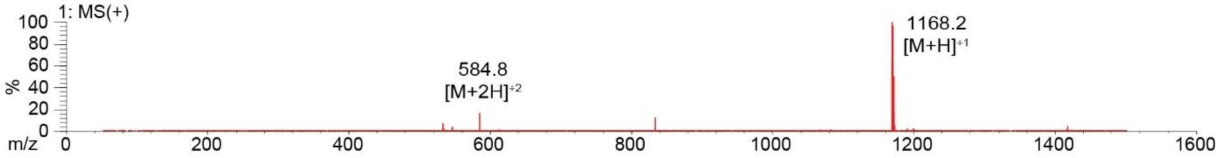

**Figure S68**. Data page for **25c**.

Expected: [M+1] = 1168.46, $[M+2]^{+2}$ = 584.73.

Observed: [M+1] = 1168.2, $[M+2]^{+2}$ = 584.8

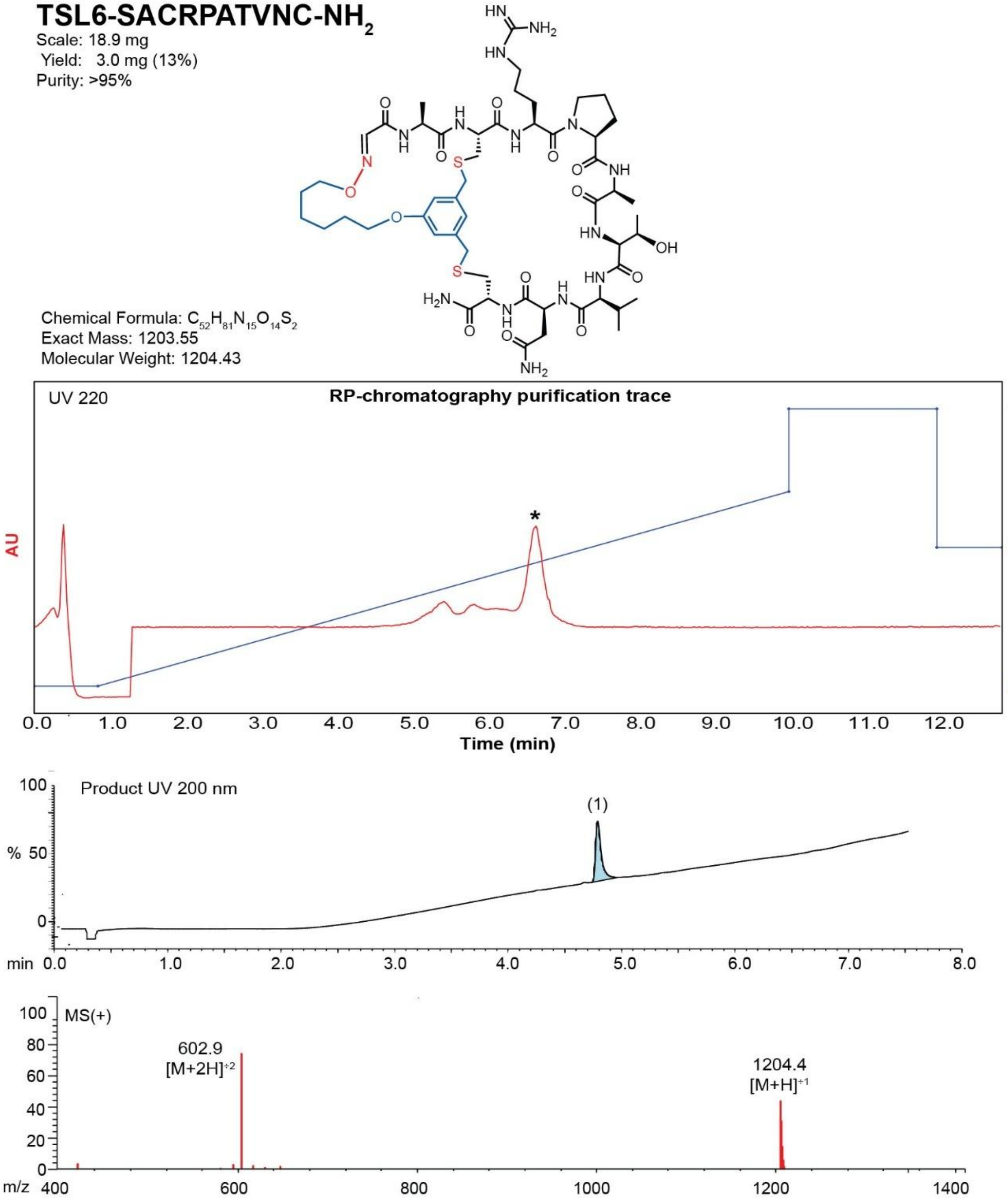

**Figure S69.** Data page for **26c**.

Expected: [M+1] = 1204.55, $[M+2]^{+2}$ = 602.775.

Observed: [M+1] = 1204.4, $[M+2]^{+2}$ = 602.9

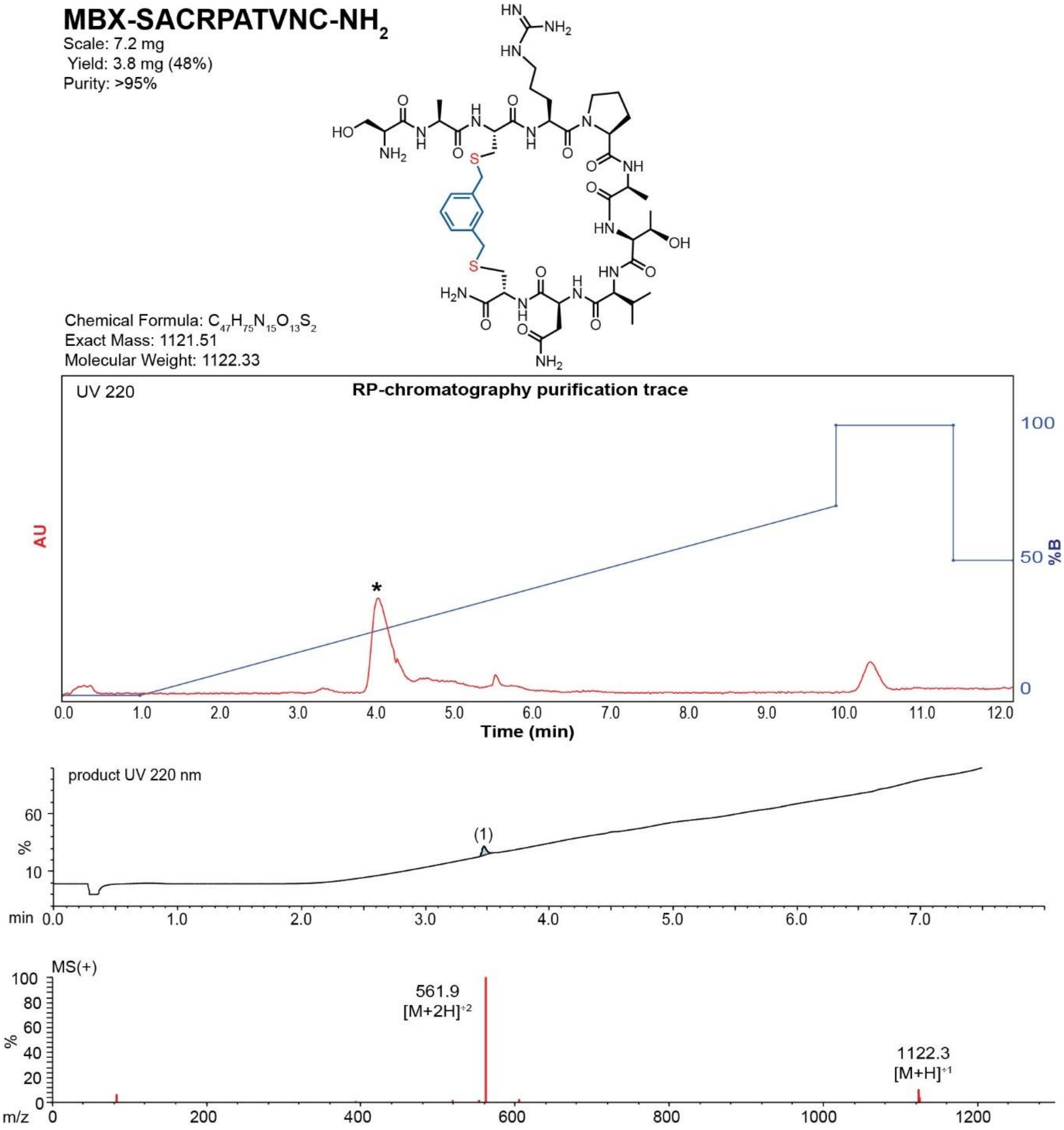

**Figure S70**. Data page for **26f**.

Expected: [M+1] = 1122.51, $[M+2]^{+2}$ = 561.755.

Observed: [M+1] = 1122.3, $[M+2]^{+2}$ = 561.9

**TSL6-SWCAPATVNC-NH$_2$**
Scale: 18.2 mg
Yield: 3.1 mg (15%)
Purity: >95%

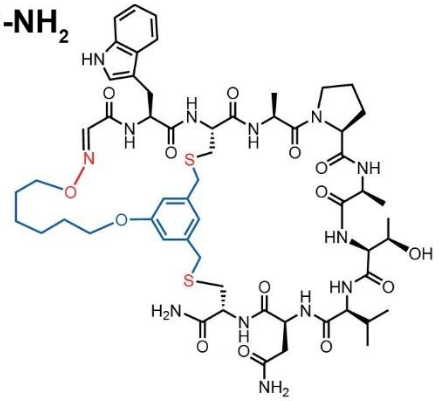

Chemical Formula: $C_{57}H_{79}N_{13}O_{14}S_2$
Exact Mass: 1233.53
Molecular Weight: 1234.46

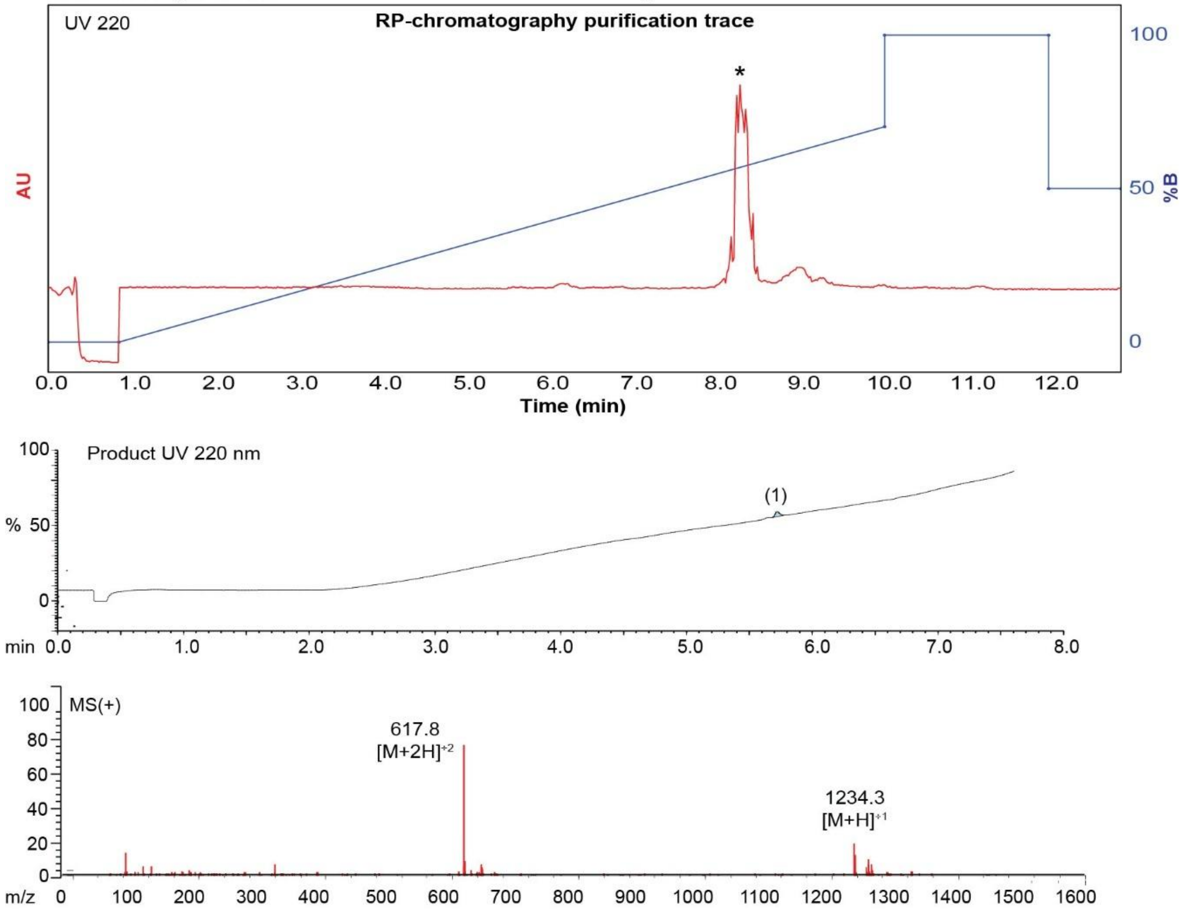

**Figure S71**. Data page for **27c**.

Expected: [M+1] = 1234.53, $[M+2]^{+2}$ = 617.765.

Observed: [M+1] = 1234.3, $[M+2]^{+2}$ = 617.8

**MBX-SWCAPATVNC-$NH_2$**

Scale: 4.8 mg

Yield: 2.5 mg (48%)

Purity: >95%

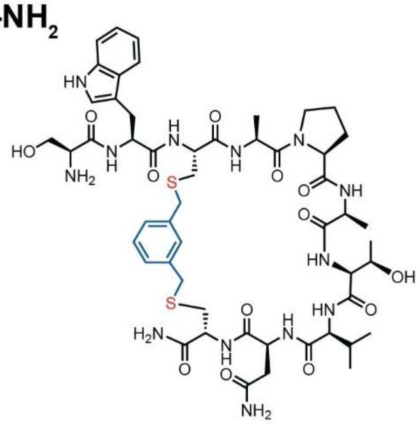

Chemical Formula: $C_{52}H_{73}N_{13}O_{13}S_2$

Exact Mass: 1151.49

Molecular Weight: 1152.35

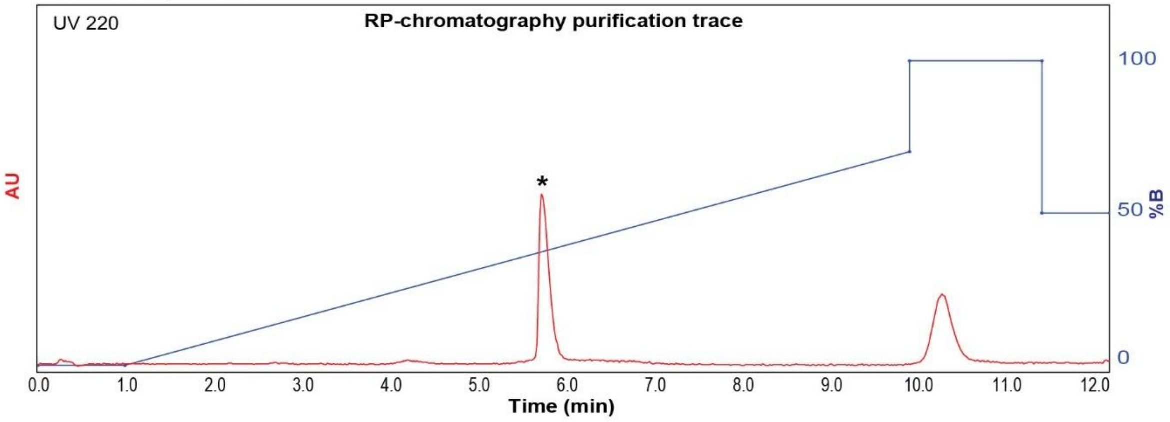

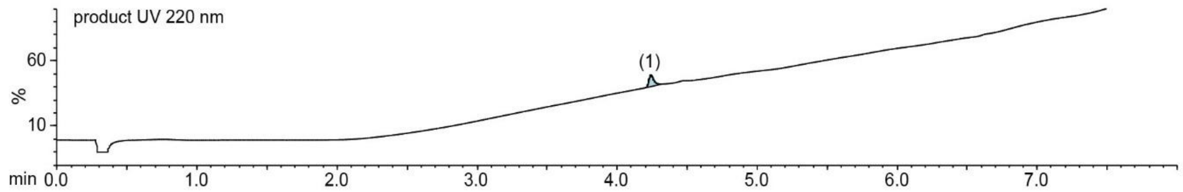

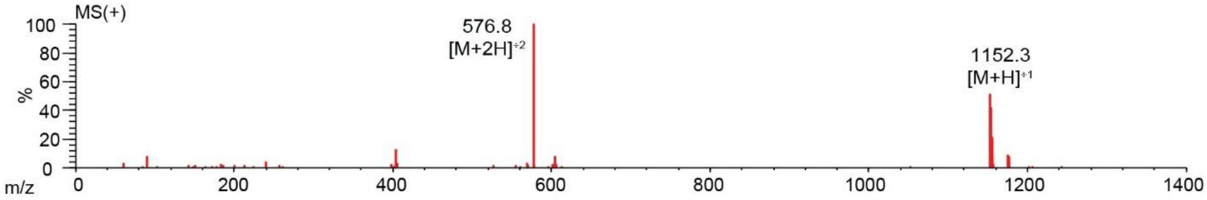

**Figure S72.** Data page for **27f**.

Expected: [M+1] = 1152.49, $[M+2]^{+2}$ = 576.745.

Observed: [M+1] = 1152.3, $[M+2]^{+2}$ = 576.8

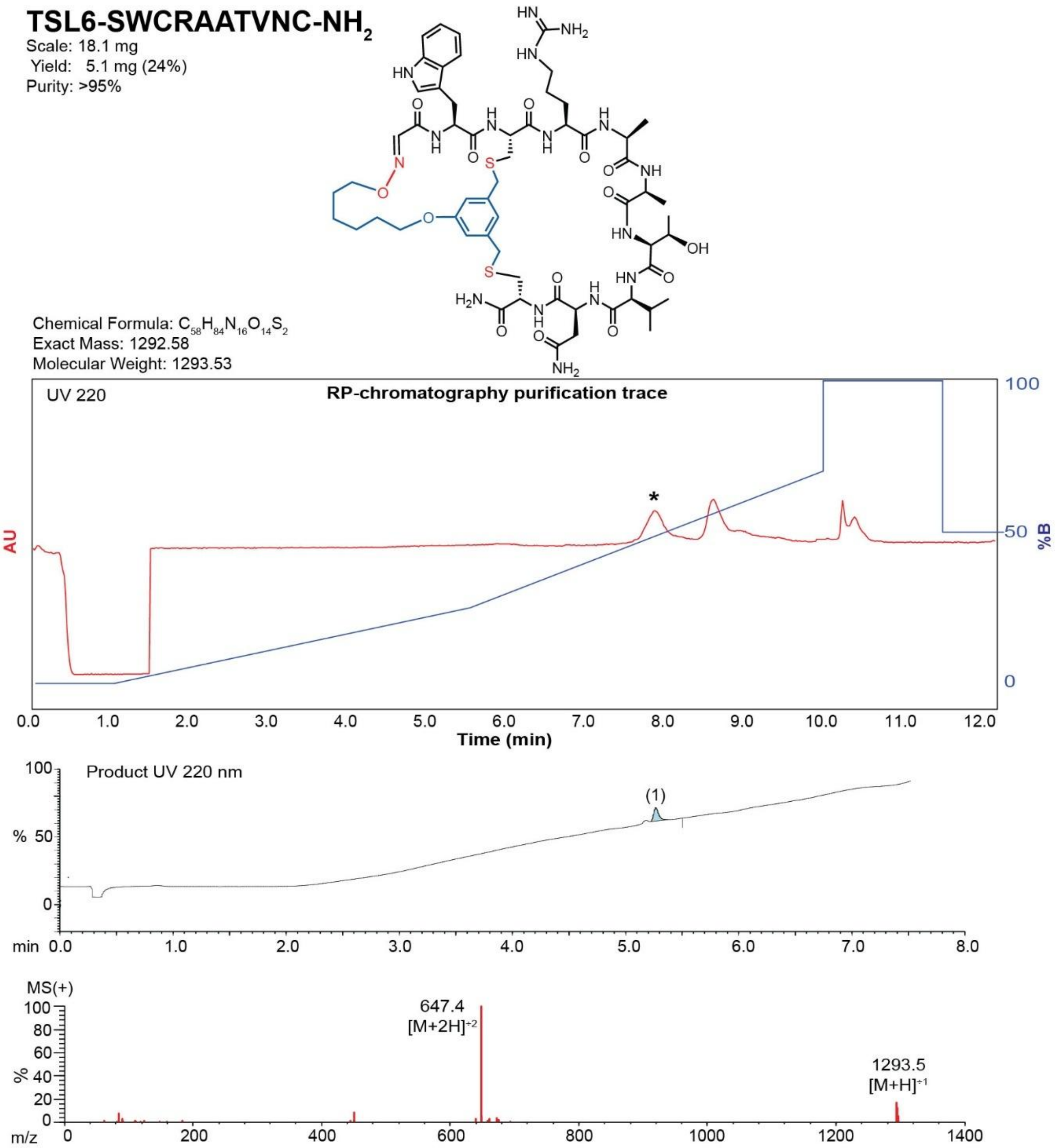

**Figure S73**. Data page for **28c**.

Expected: [M+1] = 1293.58, $[M+2]^{+2}$ = 647.29.

Observed: [M+1] = 1293.5, $[M+2]^{+2}$ = 647.4

**MBX-SWCRAATVNC-$NH_2$**

Scale: 4.0 mg
Yield: 2.1 mg (48%)
Purity: >95%

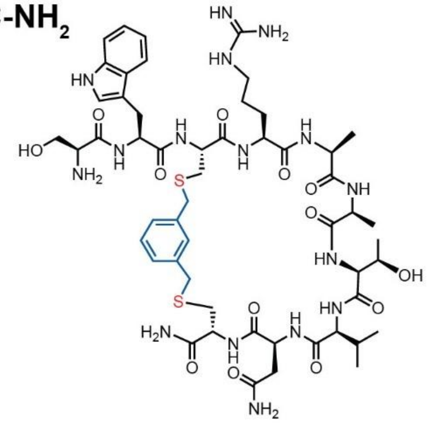

Chemical Formula: $C_{53}H_{78}N_{16}O_{13}S_2$
Exact Mass: 1210.54
Molecular Weight: 1211.43

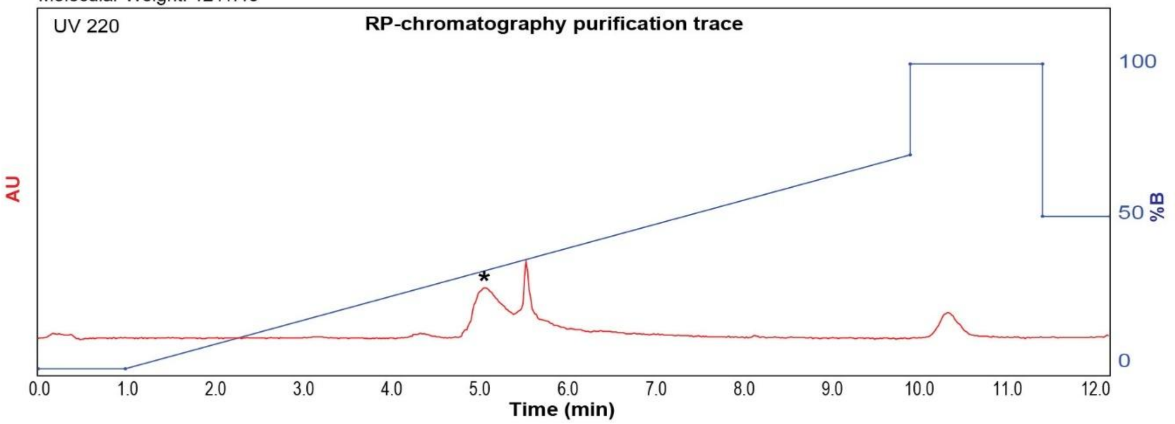

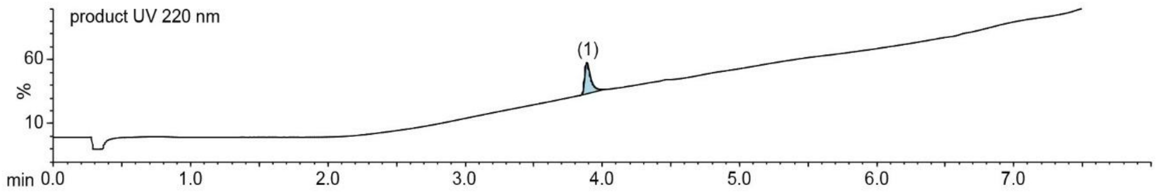

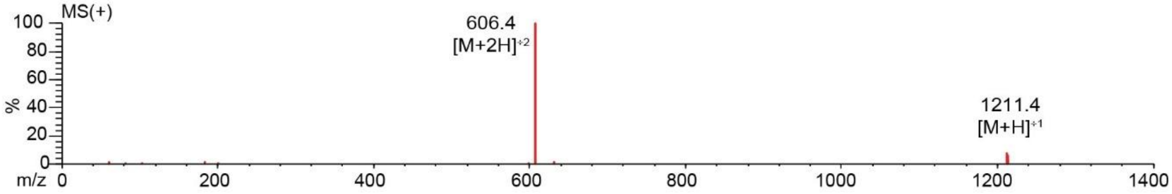

**Figure S74**. Data page for **28f**.

Expected: [M+1] = 1211.54, $[M+2]^{+2}$ = 606.27.

Observed: [M+1] = 1211.4, $[M+2]^{+2}$ = 606.4

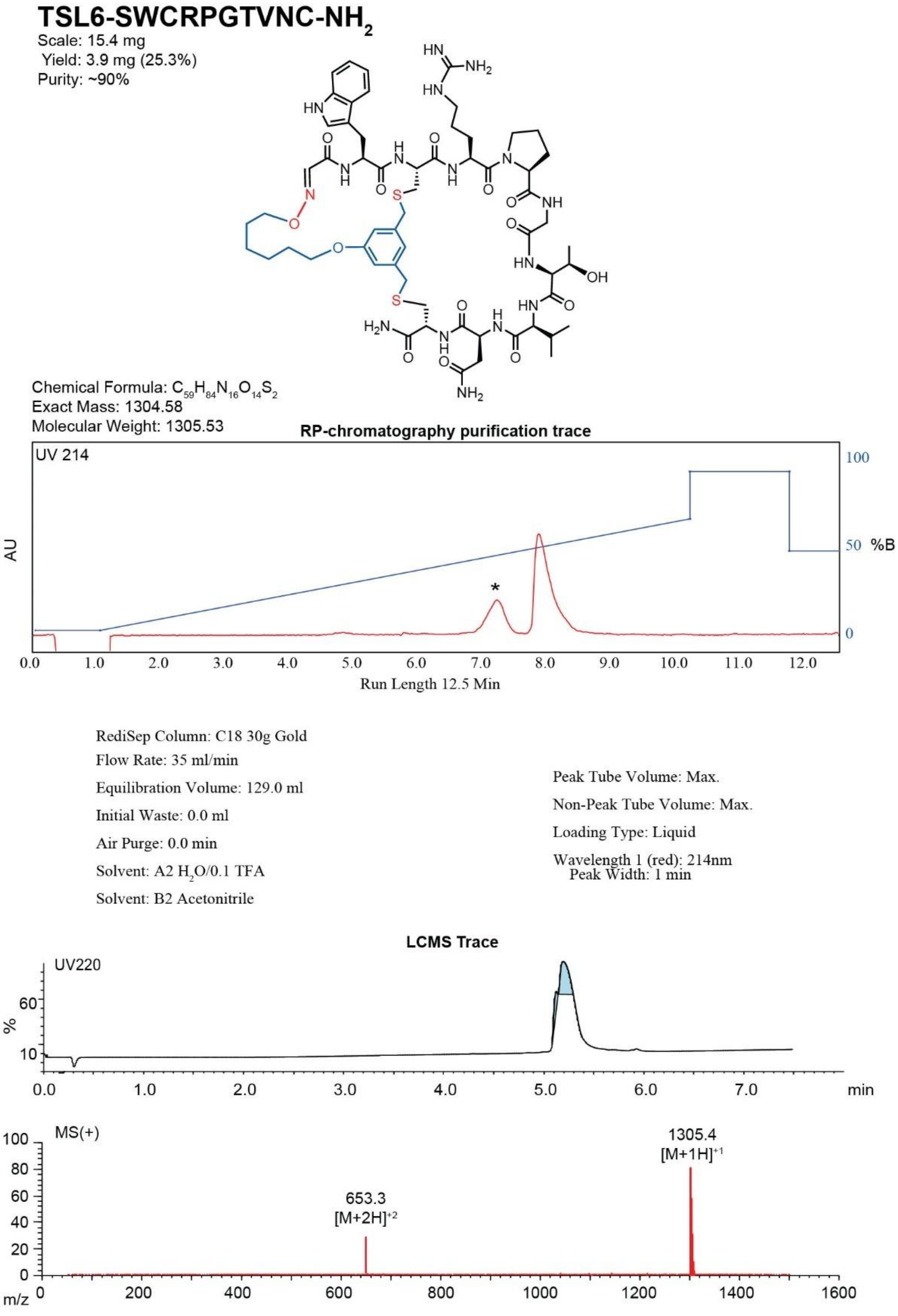

**Figure S75**. Data page for **29c**.

Expected: [M+1] = 1305.58, $[M+2]^{+2}$ = 653.29.

Observed: [M+1] = 1305.4, $[M+2]^{+2}$ = 653.3

**MBX-SWCRPGTVNC-NH$_2$**

Scale: 9.2 mg

Yield: 5.8 mg (63%)

Purity: >95%

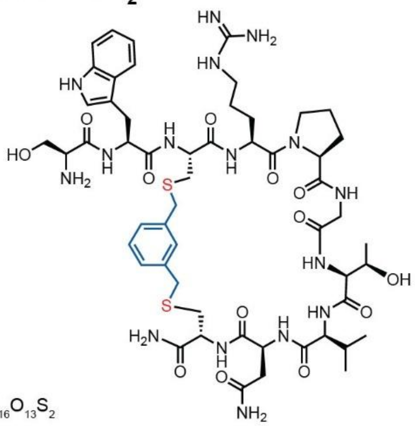

Chemical Formula: $C_{54}H_{78}N_{16}O_{13}S_2$

Exact Mass: 1222.54

Molecular Weight: 1223.43

**RP-chromatography purification trace**

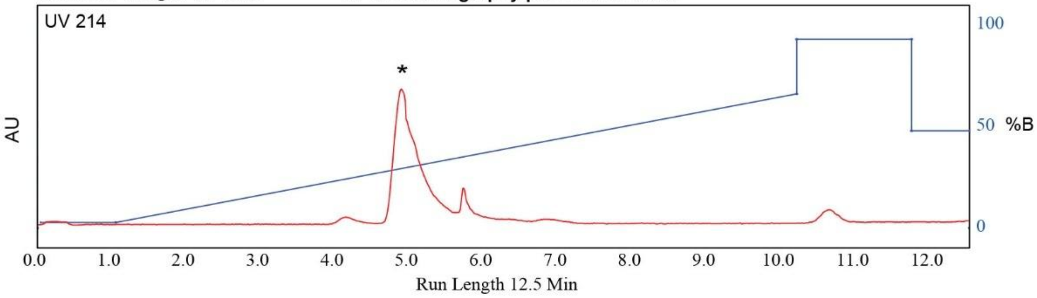

RediSep Column: C18 30g Gold

Flow Rate: 35 ml/min

Equilibration Volume: 129.0 ml

Initial Waste: 0.0 ml

Air Purge: 0.0 min

Solvent: A2 $H_2O$/0.1 TFA

Solvent: B2 Acetonitrile

Peak Tube Volume: Max.

Non-Peak Tube Volume: Max.

Loading Type: Liquid

Wavelength 1 (red): 214nm

Peak Width: 1 min

**LCMS Trace**

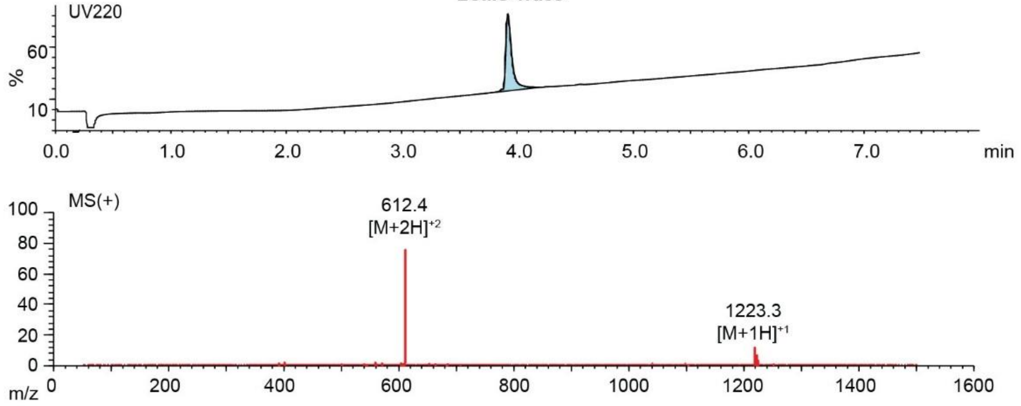

**Figure S76**. Data page for **29f**.

Expected: $[M+1] = 1223.54$, $[M+2]^{+2} = 612.27$.

Observed: $[M+1] = 1223.3$, $[M+2]^{+2} = 612.4$

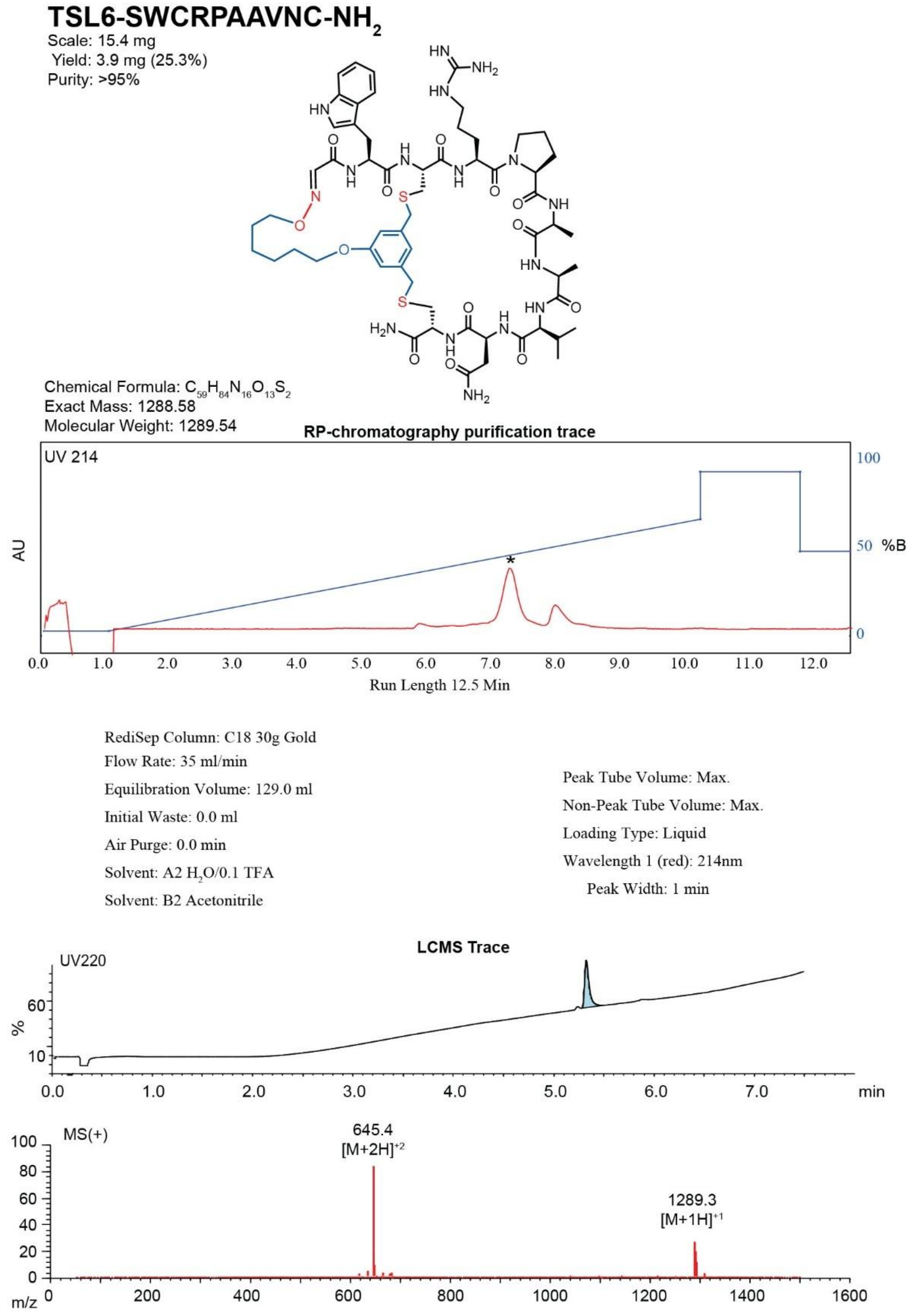

**Figure S77**. Data page for **30c**.

Expected: [M+1] = 1289.58, $[M+2]^{\div 2}$ = 645.29.

Observed: [M+1] = 1289.3, $[M+2]^{\div 2}$ = 645.4

### MBX-SWCRPAAVNC-$NH_2$

Scale: 6.4 mg
Yield: 3.1 mg (48.4%)
Purity: >95%

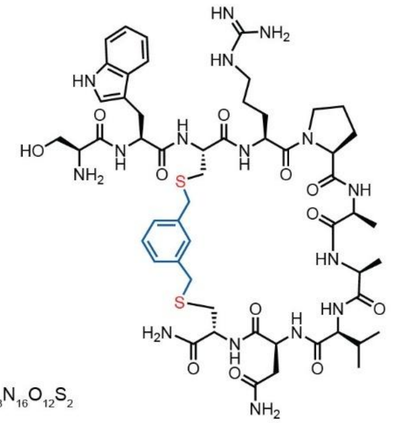

Chemical Formula: $C_{54}H_{78}N_{16}O_{12}S_2$
Exact Mass: 1206.54
Molecular Weight: 1207.44

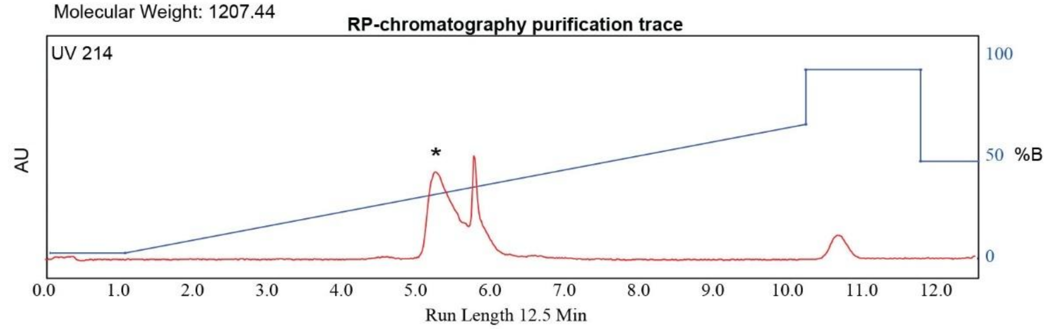

RediSep Column: C18 30g Gold
Flow Rate: 35 ml/min
Equilibration Volume: 129.0 ml
Initial Waste: 0.0 ml
Air Purge: 0.0 min
Solvent: A2 $H_2O$/0.1 TFA
Solvent: B2 Acetonitrile

Peak Tube Volume: Max.
Non-Peak Tube Volume: Max.
Loading Type: Liquid
Wavelength 1 (red): 214nm
Peak Width: 1 min

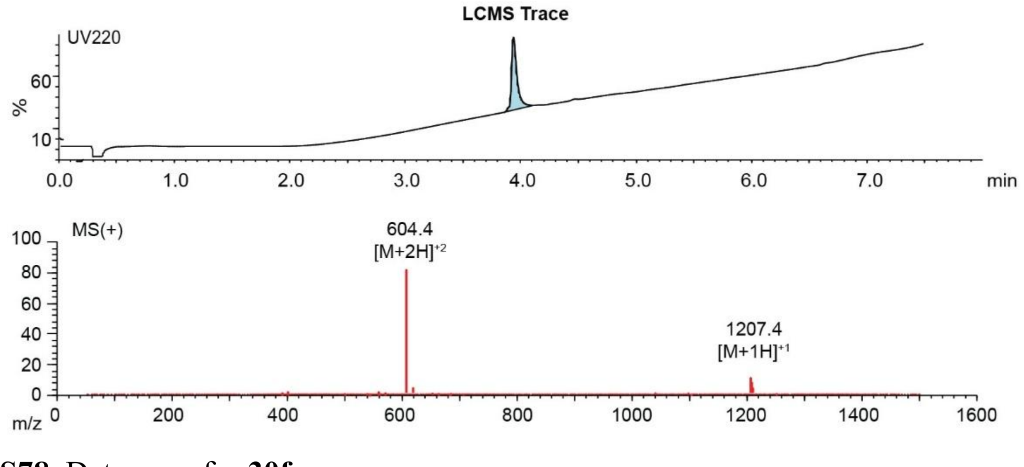

**Figure S78**. Data page for **30f**.

Expected: $[M+1] = 1207.54$, $[M+2]^{+2} = 604.27$.

Observed: $[M+1] = 1207.4$, $[M+2]^{+2} = 604.4$

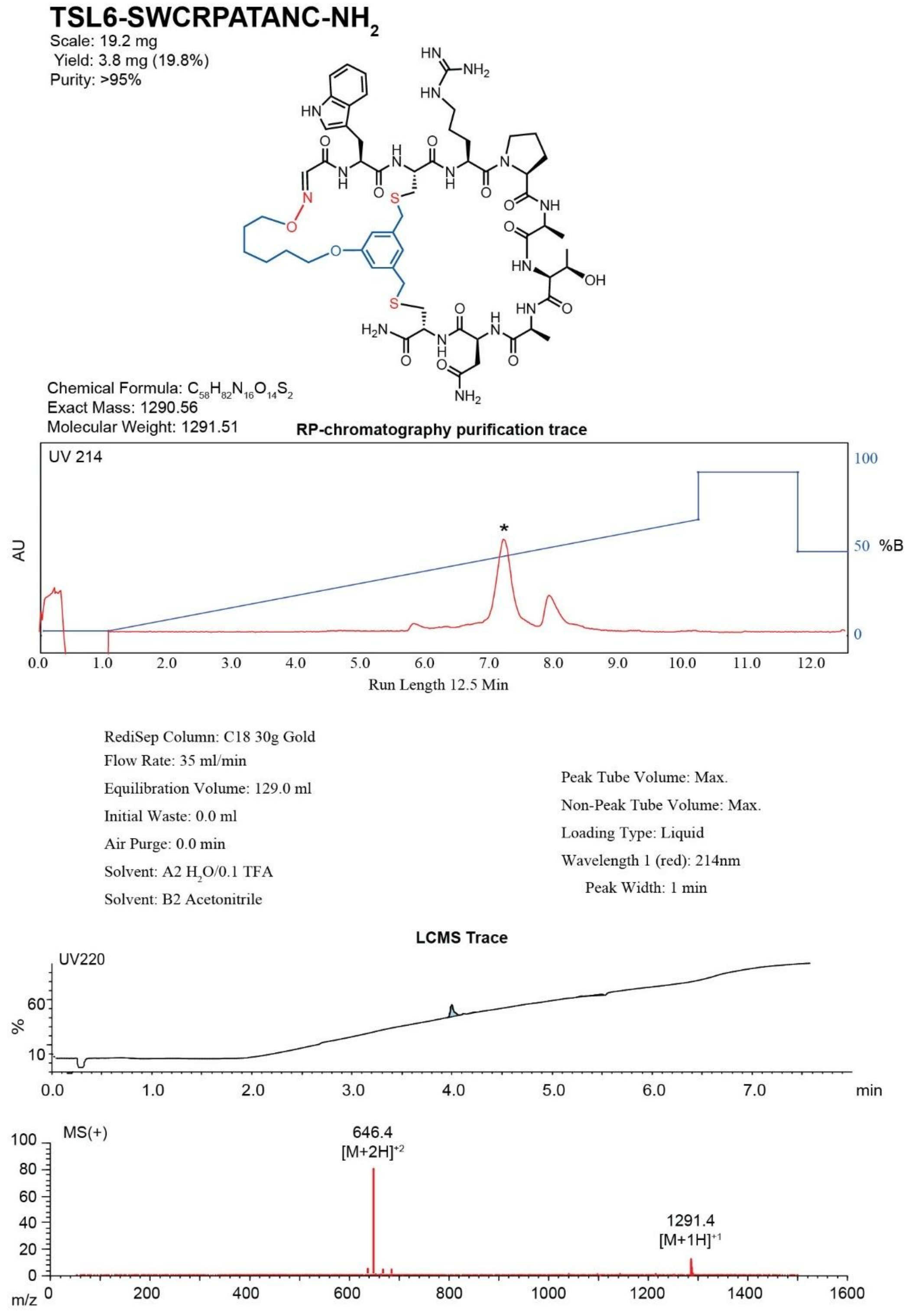

**Figure S79**. Data page for **31c**.

Expected: [M+1] = 1291.56, $[M+2]^{+2}$ = 646.28.

Observed: [M+1] = 1291.4, $[M+2]^{+2}$ = 646.4

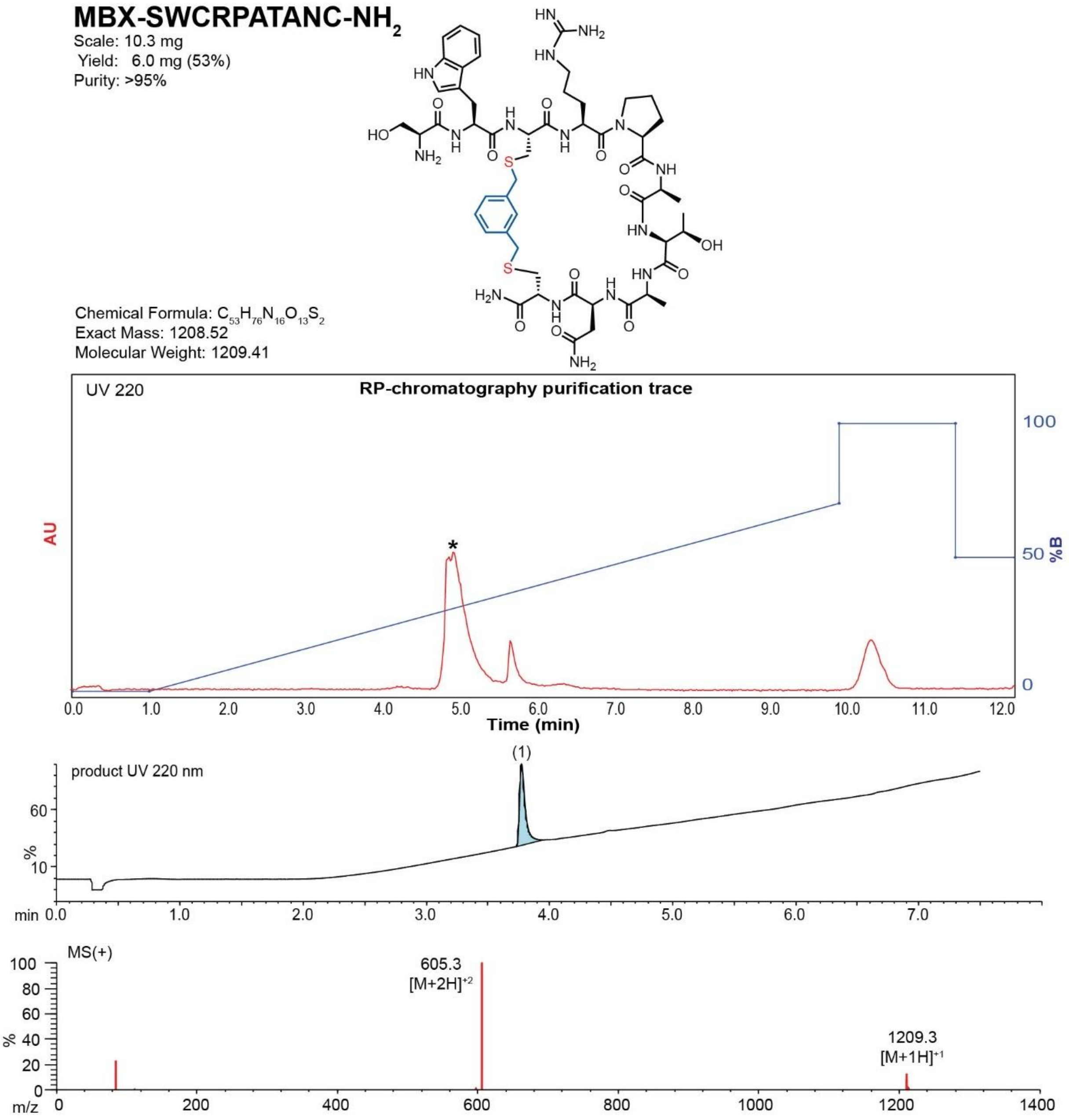

**Figure S80**. Data page for **31f**.

Expected: [M+1] = 1209.52, $[M+2]^{+2}$ = 605.26.

Observed: [M+1] = 1209.3, $[M+2]^{+2}$ = 605.3

**TSL6-SWCRPATVAC-$NH_2$**

Scale: 18.5 mg

Yield: 3.9 mg (21.1%)

Purity: >95%

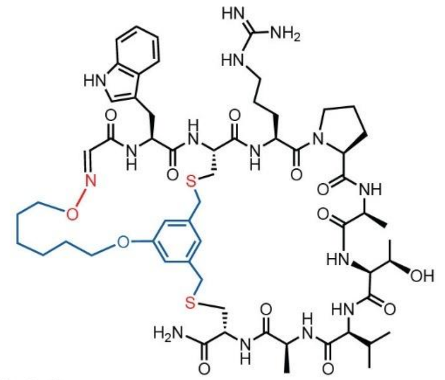

Chemical Formula: $C_{59}H_{85}N_{15}O_{13}S_2$

Exact Mass: 1275.59

Molecular Weight: 1276.54

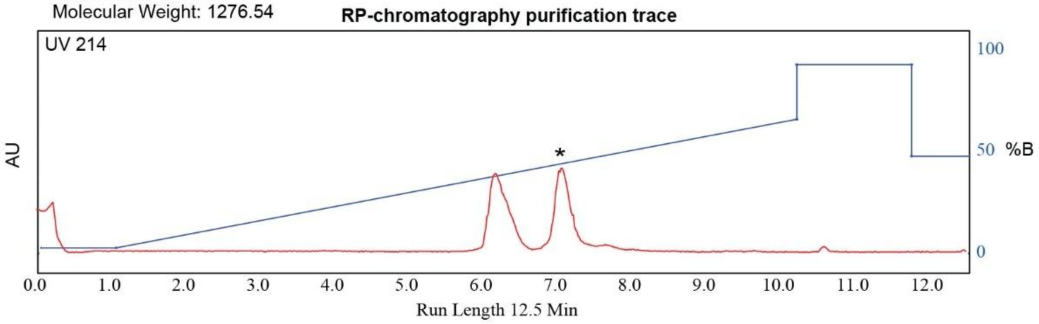

RediSep Column: C18 30g Gold

Flow Rate: 35 ml/min

Equilibration Volume: 129.0 ml

Initial Waste: 0.0 ml

Air Purge: 0.0 min

Solvent: A2 $H_2O$/0.1 TFA

Solvent: B2 Acetonitrile

Peak Tube Volume: Max.

Non-Peak Tube Volume: Max.

Loading Type: Liquid

Wavelength 1 (red): 214nm

Peak Width: 1 min

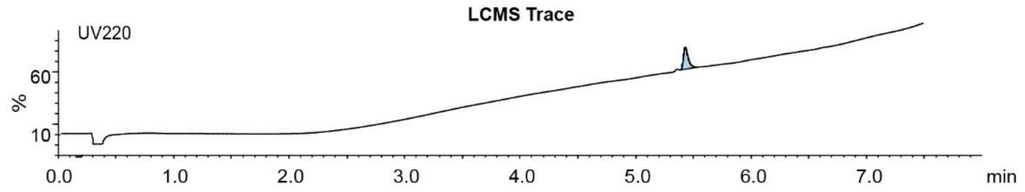

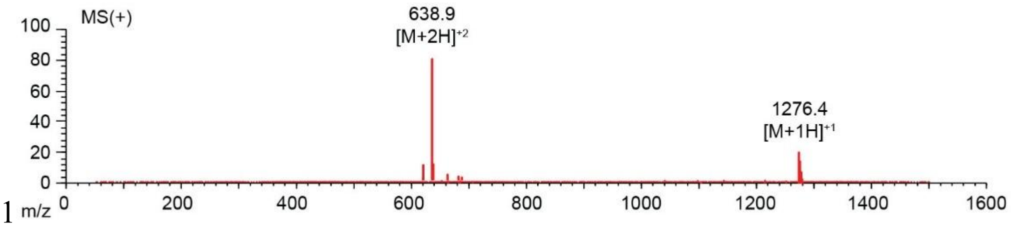

**Figure S81**. Data page for **32c**.

Expected: [M+1] = 1276.59, $[M+2]^{\div 2}$ = 638.795.

Observed: [M+1] = 1276.4, $[M+2]^{\div 2}$ = 638.9

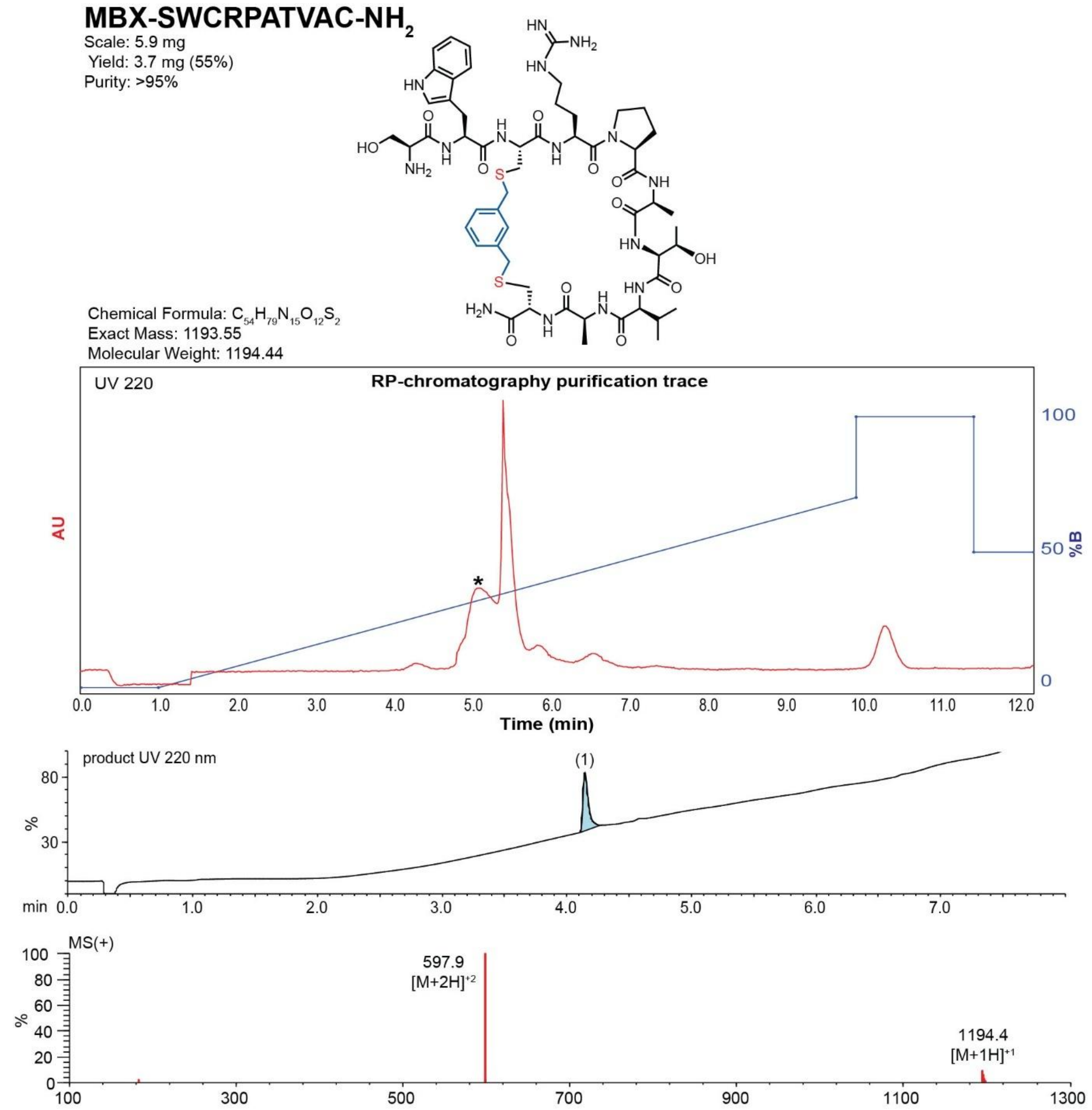

**Figure S82**. Data page for **32f**.

Expected: [M+1] = 1194.55, $[M+2]^{+2}$ = 597.775.

Observed: [M+1] = 1194.4, $[M+2]^{+2}$ = 597.9

**TSL6-SACHYPTHVC-$NH_2$**

Scale: 4.7 mg
Yield: 1.7 mg (31%)
Purity: >95%

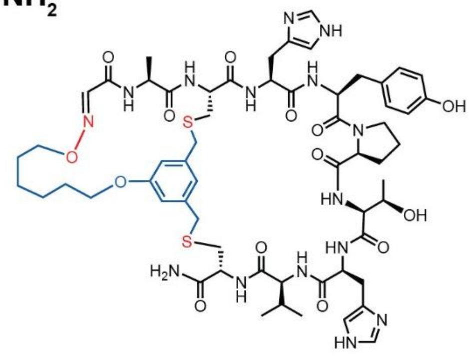

Chemical Formula: $C_{60}H_{81}N_{15}O_{14}S_2$
Exact Mass: 1299.55
Molecular Weight: 1300.52

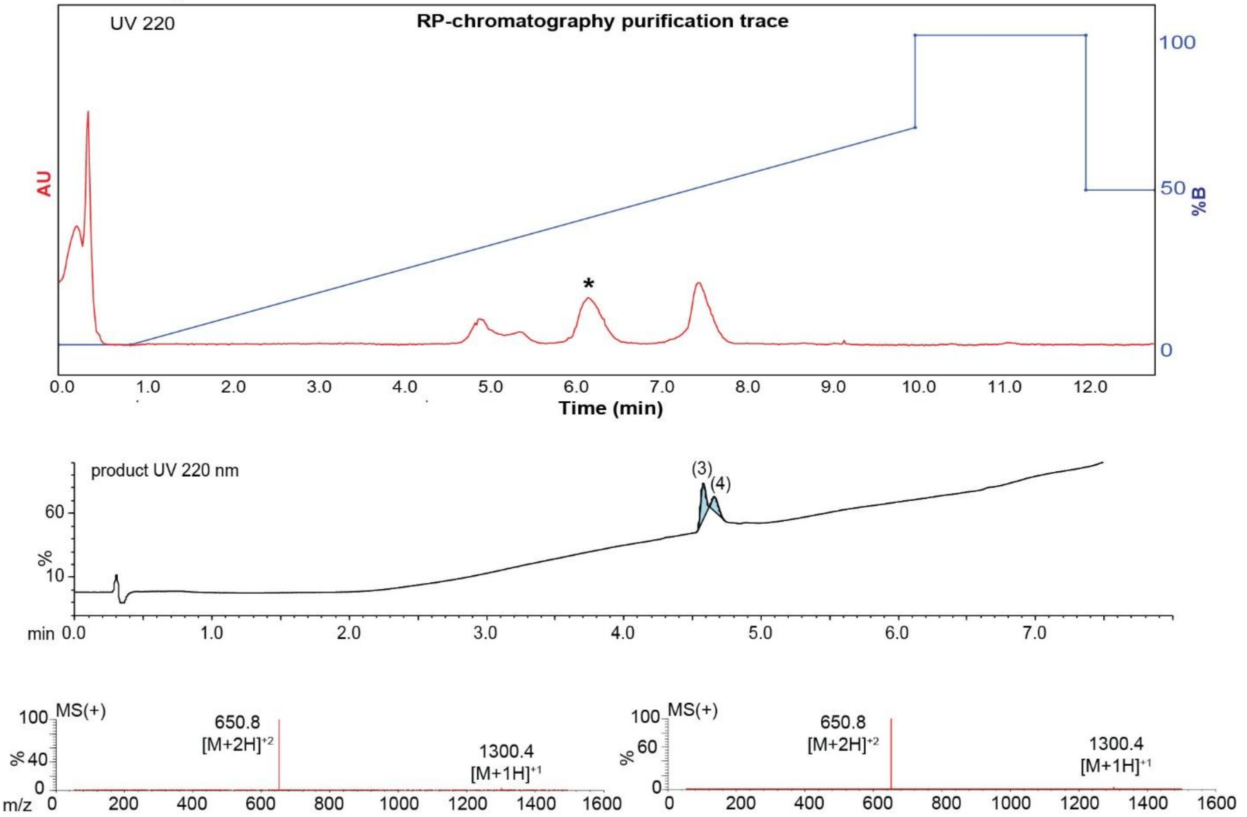

**Figure S83**. Data page for **33c**.

Expected: [M+1] = 1300.55, $[M+2]^{+2}$ = 650.775.

Observed: [M+1] = 1300.4, $[M+2]^{+2}$ = 650.8

### MBX-SACHYPTHVC-$NH_2$

Scale: 6.1 mg
Yield: 3.5 mg (53%)
Purity: >95%

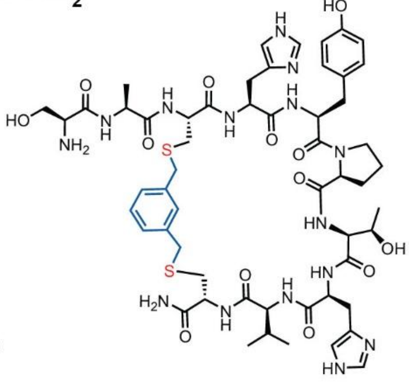

Chemical Formula: $C_{55}H_{75}N_{15}O_{13}S_2$
Exact Mass: 1217.51
Molecular Weight: 1218.42

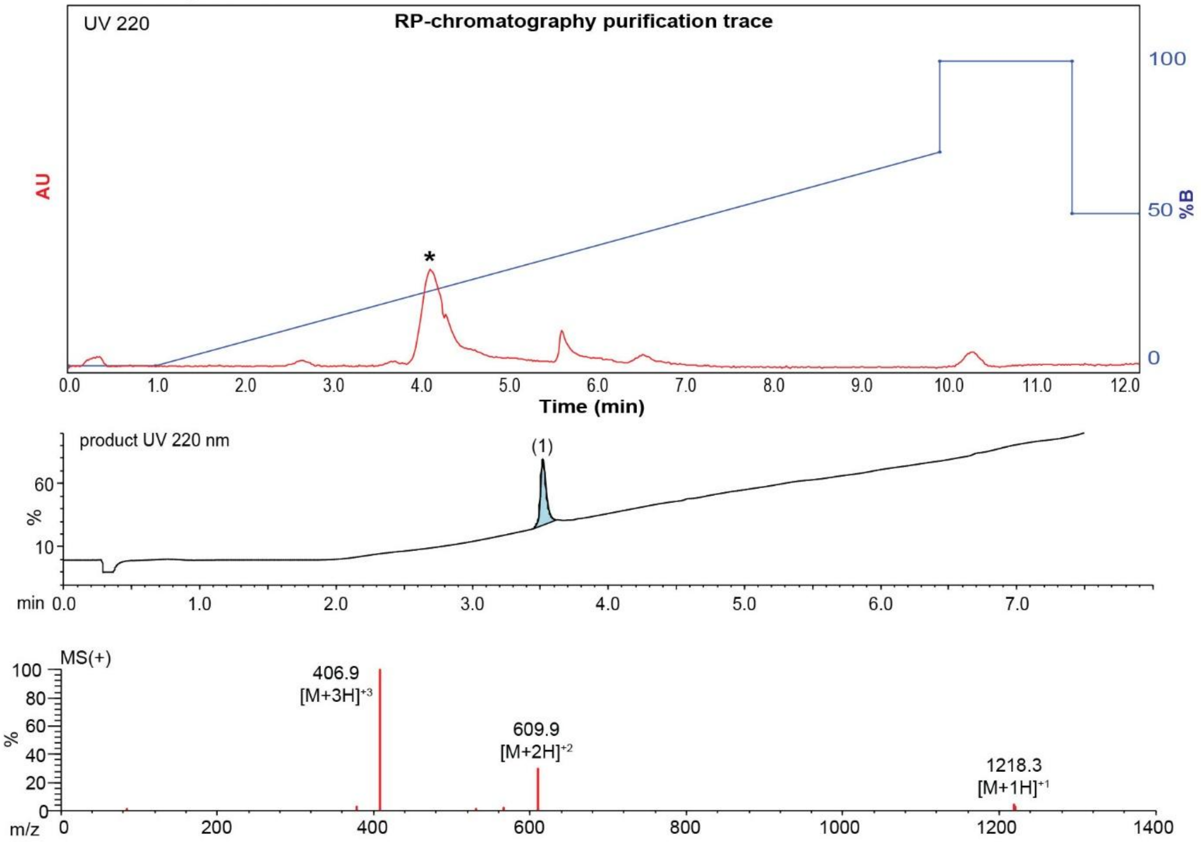

**Figure S84**. Data page for **33f**.

Expected: [M+1] = 1218.51, $[M+2]^{+2}$ = 609.755.

Observed: [M+1] = 1218.3, $[M+2]^{+2}$ = 609.9

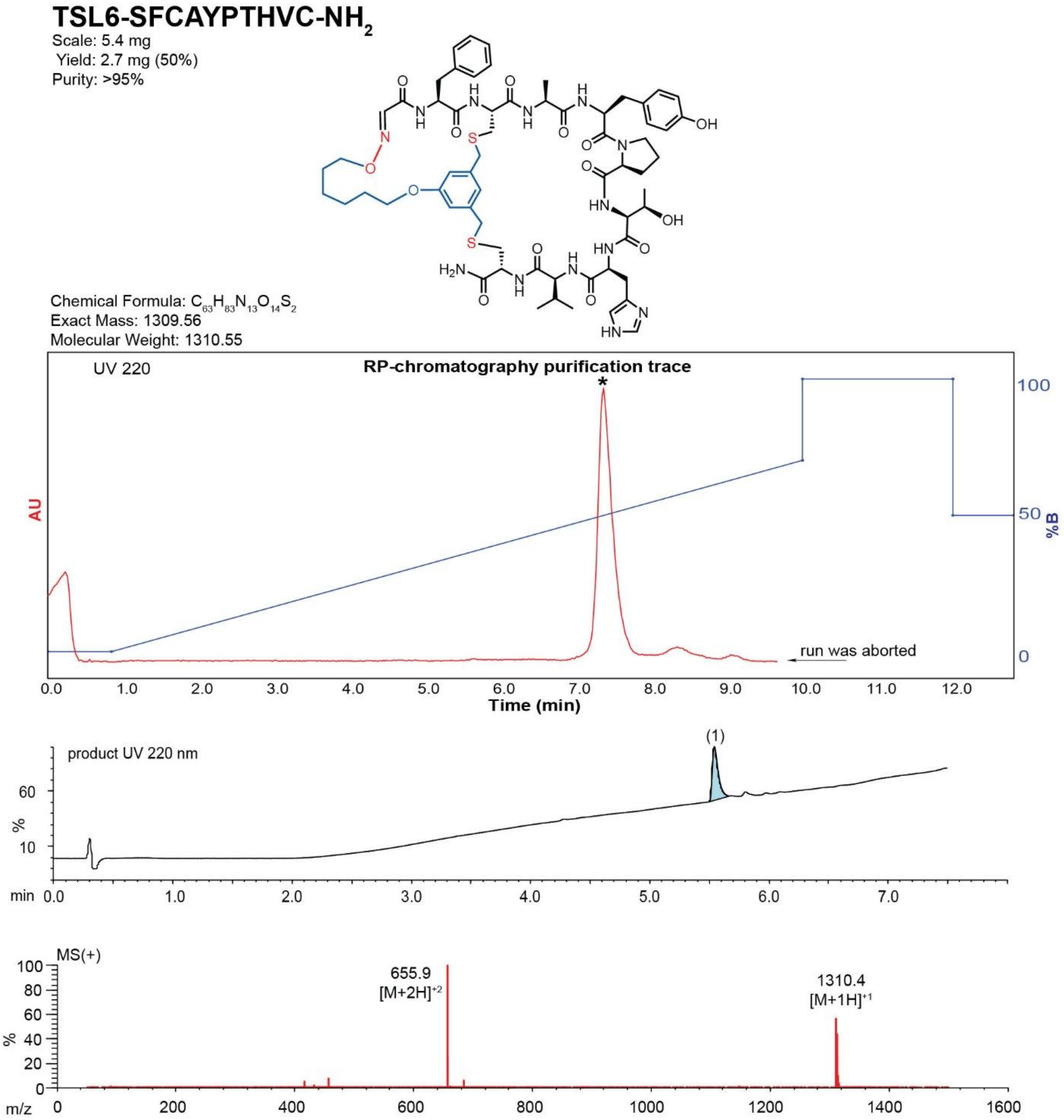

**Figure S85**. Data page for **34c**.

Expected: [M+1] = 1310.56, $[M+2]^{+2}$ = 655.78.

Observed: [M+1] = 1310.4, $[M+2]^{+2}$ = 655.9

**MBX-SFCAYPTHVC-NH$_2$**

Scale: 6.9 mg
Yield: 5.7 mg (76%)
Purity: >95%

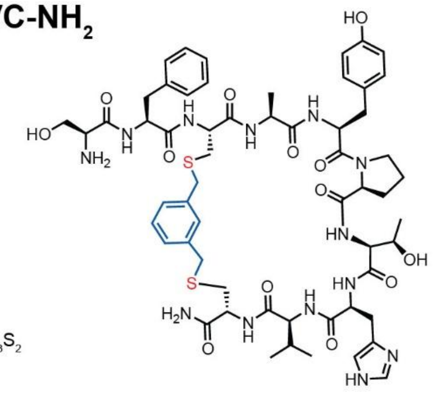

Chemical Formula: $C_{58}H_{77}N_{13}O_{13}S_2$
Exact Mass: 1227.52
Molecular Weight: 1228.45

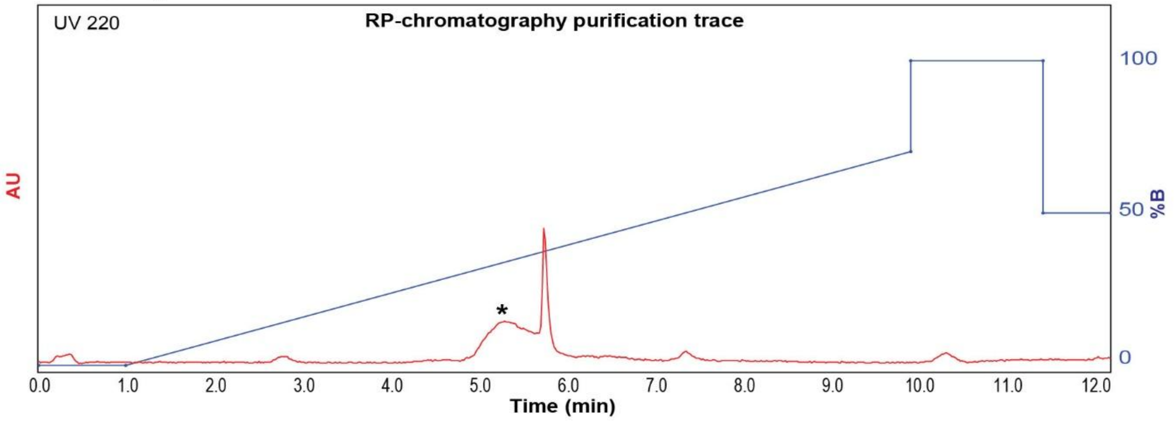

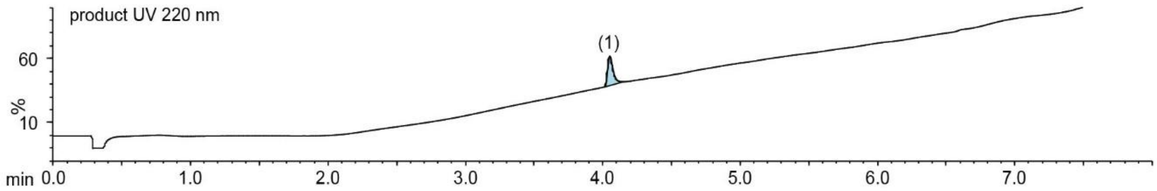

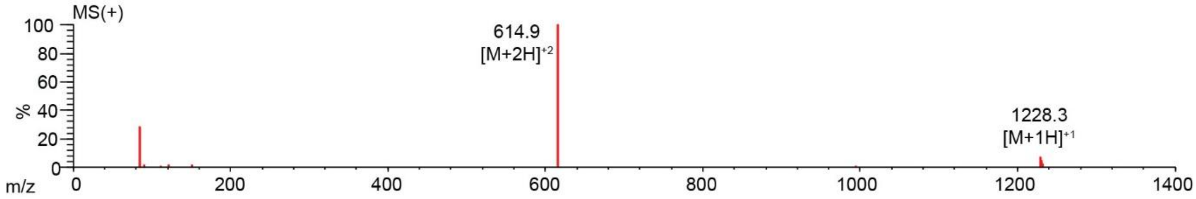

**Figure S86**. Data page for **34f**.

Expected: [M+1] = 1228.52, $[M+2]^{+2}$ = 614.76.

Observed: [M+1] = 1228.3, $[M+2]^{+2}$ = 614.9

### TSL6-SFCHAPTHVC-$NH_2$

Scale: 4.9 mg
Yield: 1.8 mg (30%)
Purity: >80%

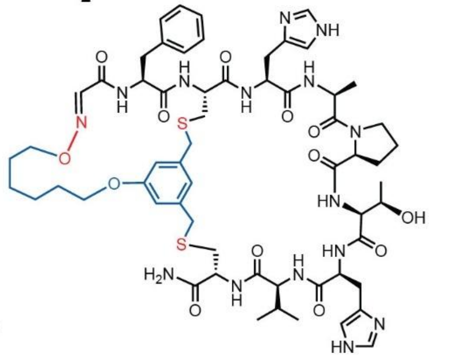

Chemical Formula: $C_{60}H_{81}N_{15}O_{13}S_2$
Exact Mass: 1283.56
Molecular Weight: 1284.52

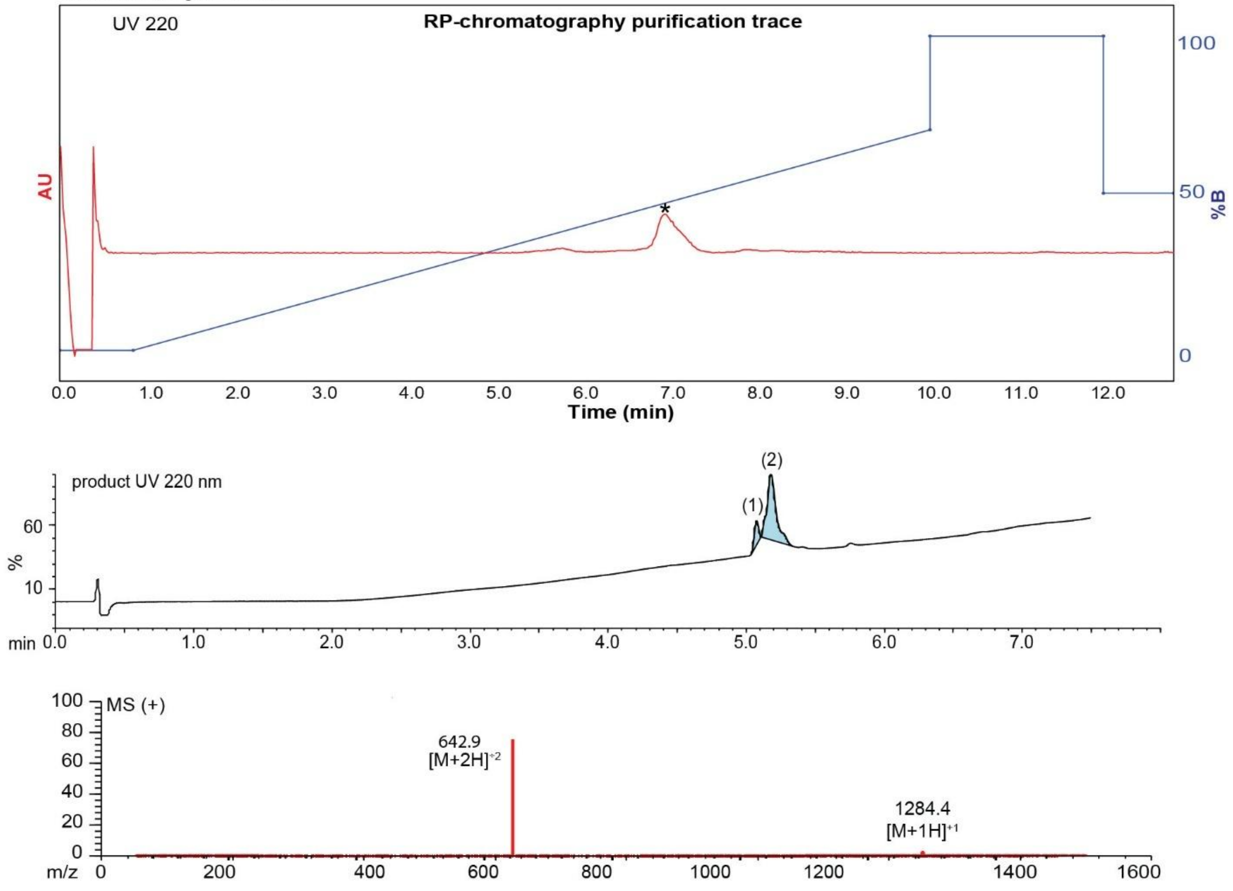

**Figure S87**. Data page for **35c**.

Expected: [M+1] = 1284.56, $[M+2]^{+2}$ = 642.78.

Observed: [M+1] = 1284.4, $[M+2]^{+2}$ = 642.9

**MBX-SFCHAPTHVC-$NH_2$**

Scale: 7.3 mg
Yield: 4.1 mg (51%)
Purity: >95%

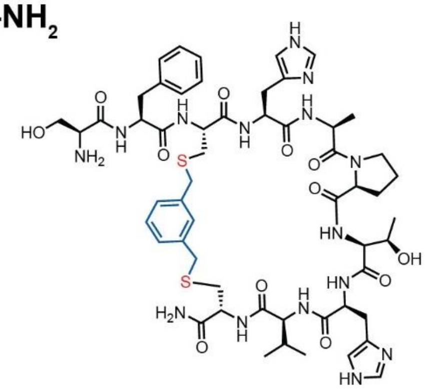

Chemical Formula: $C_{55}H_{75}N_{15}O_{12}S_2$
Exact Mass: 1201.52
Molecular Weight: 1202.42

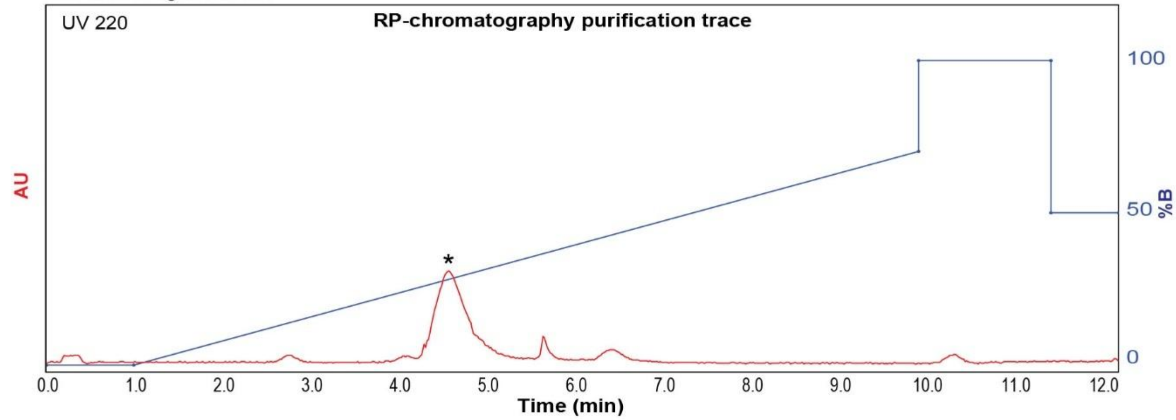

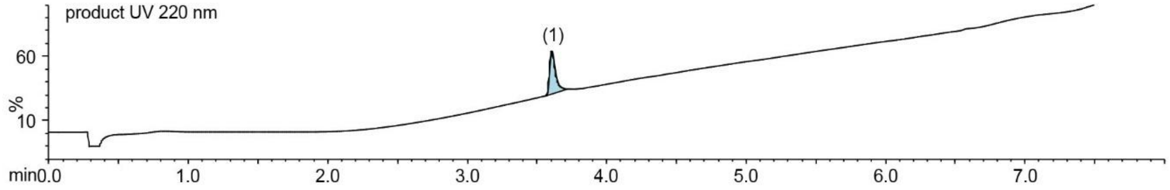

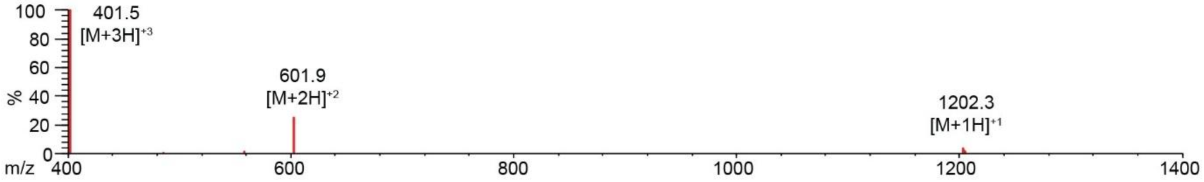

**Figure S88**. Data page for **35f**.

Expected: [M+1] = 1202.52, $[M+2]^{+2}$ = 601.76.

Observed: [M+1] = 1202.3, $[M+2]^{+2}$ = 601.9

**TSL6-SFCHYATHVC-$NH_2$**

Scale: 5.0 mg
Yield: 1.8 mg (32%)
Purity: >95%

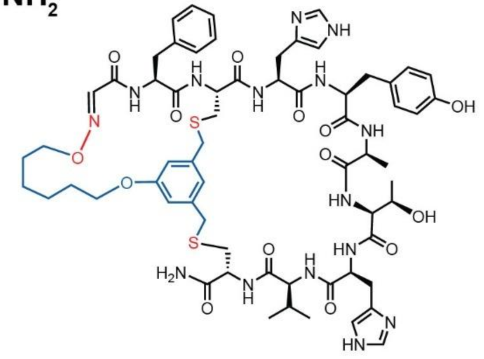

Chemical Formula: $C_{64}H_{83}N_{15}O_{14}S_2$
Exact Mass: 1349.57
Molecular Weight: 1350.58

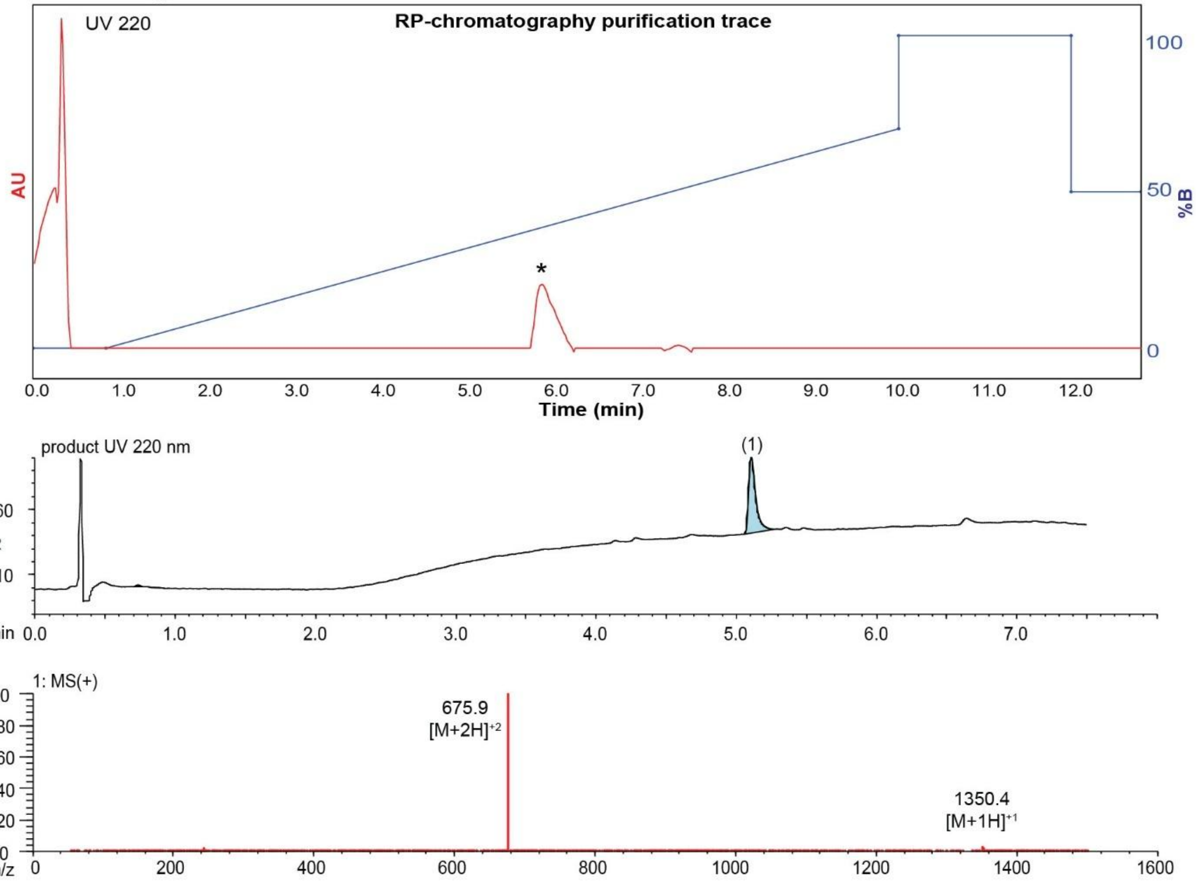

**Figure S89**. Data page for **36c**.

Expected: [M+1] = 1350.57, $[M+2]^{+2}$ = 675.785.

Observed: [M+1] = 1350.4, $[M+2]^{+2}$ = 675.9

**MBX-SFCHYATHVC-$NH_2$**

Scale: 6.3 mg
Yield: 3.9 mg (57%)
Purity: >95%

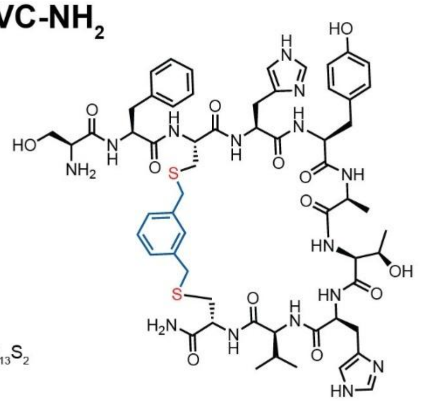

Chemical Formula: $C_{59}H_{77}N_{15}O_{13}S_2$
Exact Mass: 1267.53
Molecular Weight: 1268.48

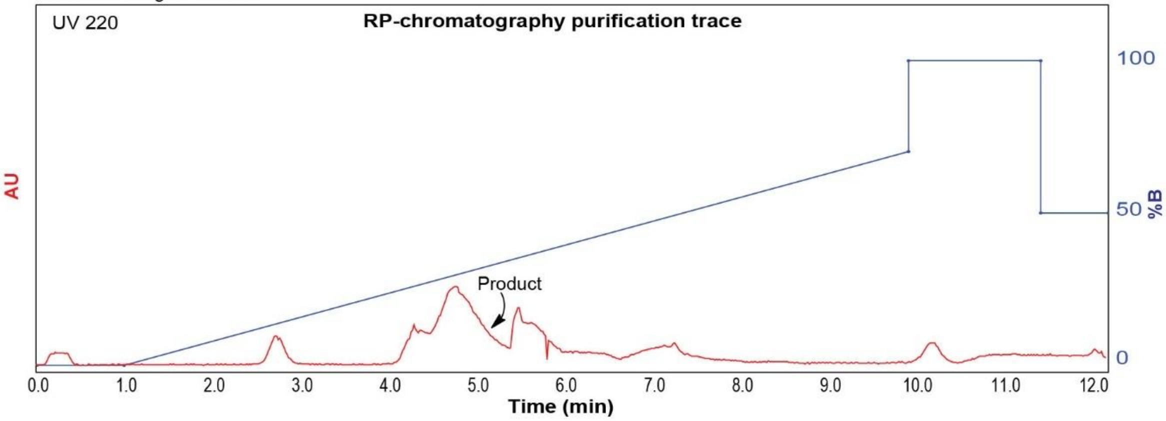

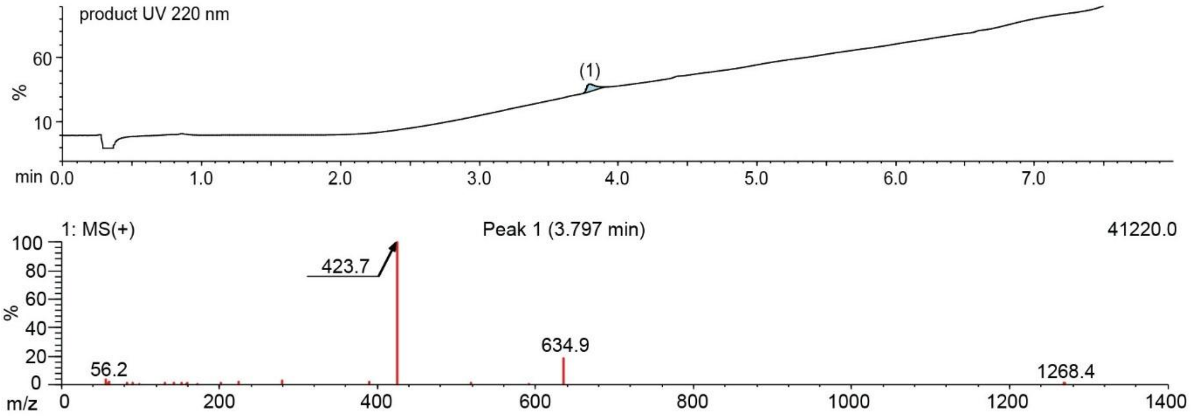

**Figure S90**. Data page for **36f**.

Expected: [M+1] = 1268.53, $[M+2]^{+2}$ = 634.765.

Observed: [M+1] = 1268.4, $[M+2]^{+2}$ = 634.9

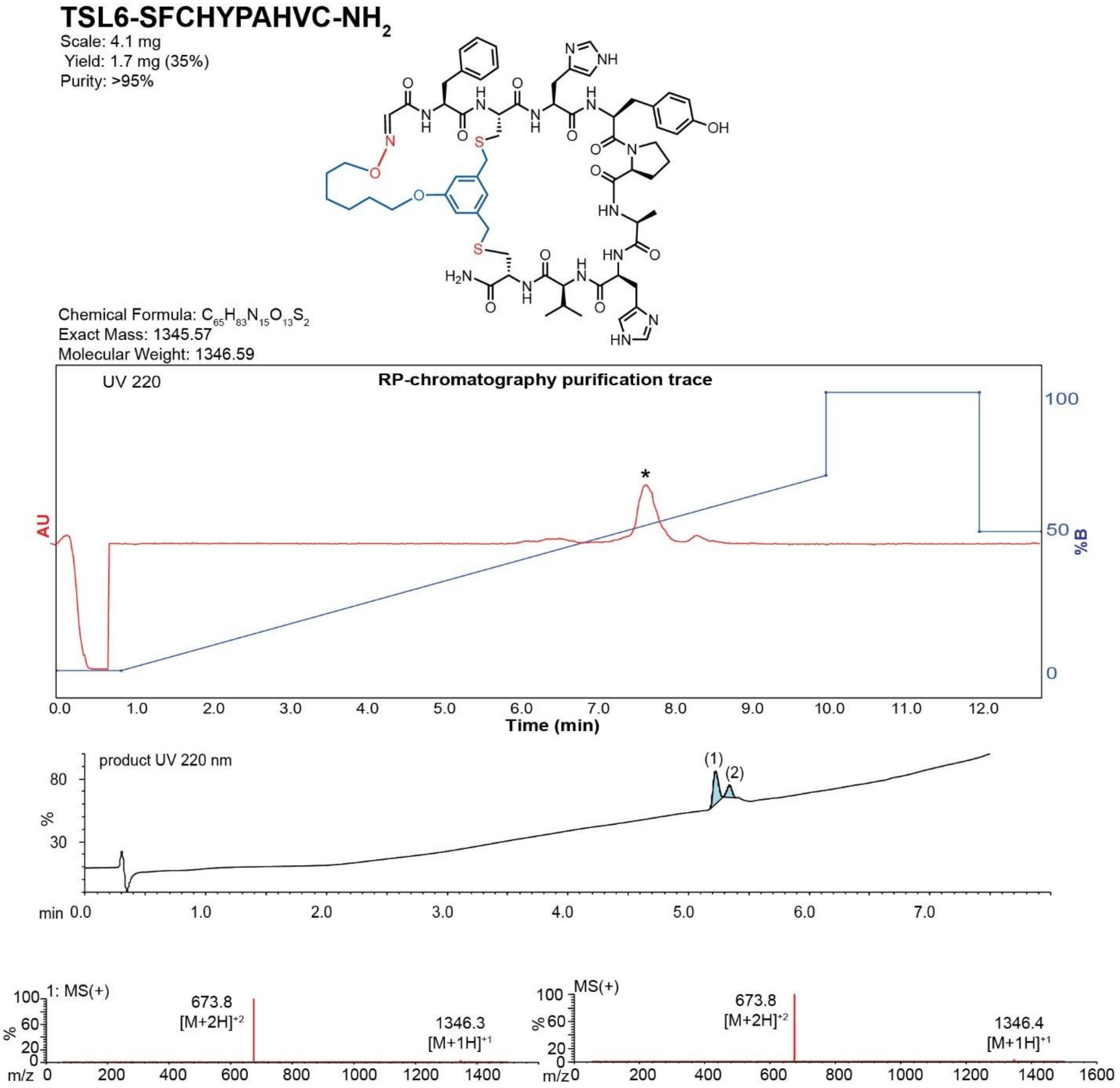

**Figure S91**. Data page for **37c**.

Expected: [M+1] = 1346.57, $[M+2]^{\div 2}$ = 673.785.

Observed: [M+1] = 1346.4, $[M+2]^{\div 2}$ = 673.8

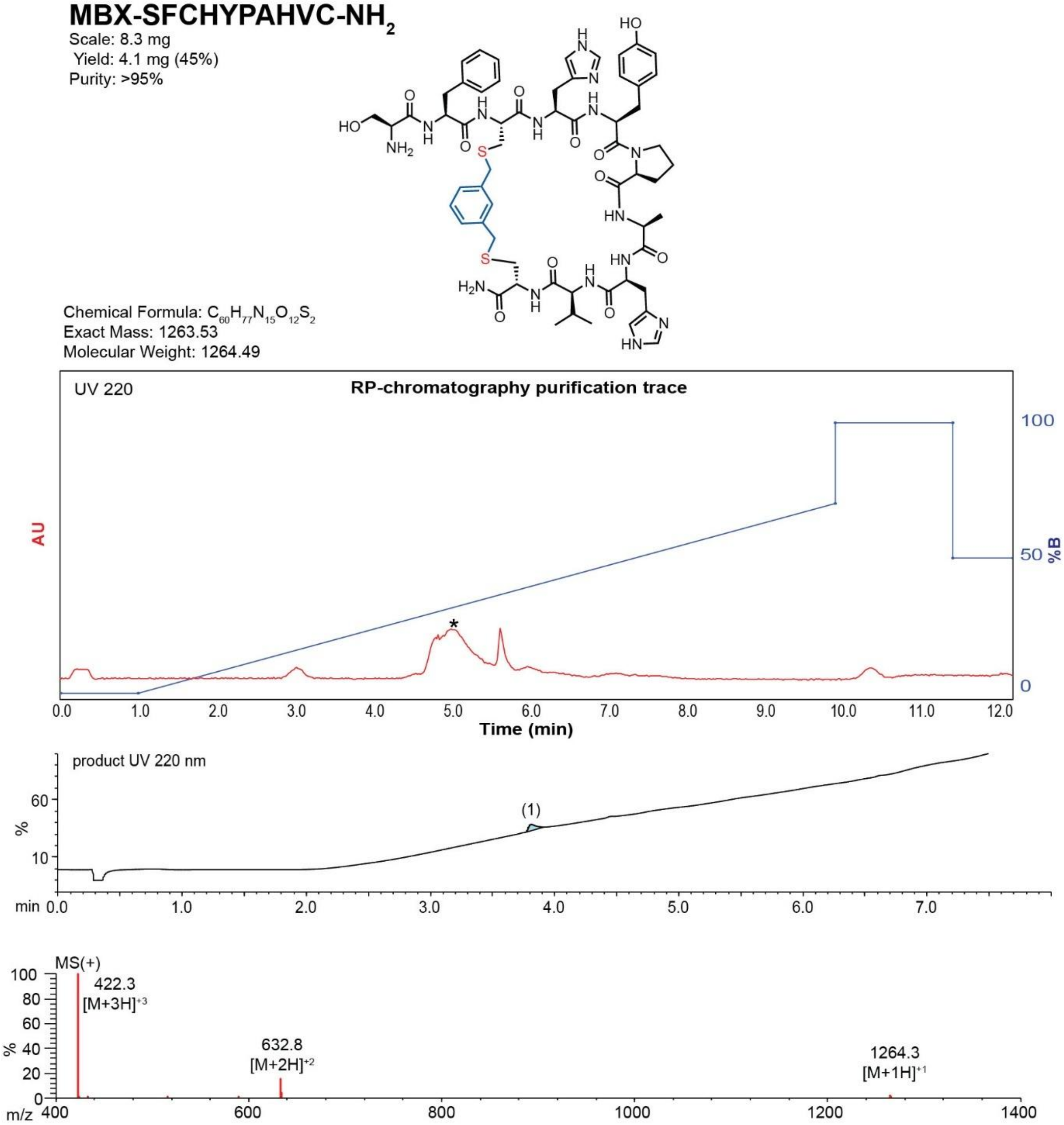

**Figure S92**. Data page for **37f**.

Expected: [M+1] = 1264.53, $[M+2]^{+2}$ = 632.765.

Observed: [M+1] = 1264.3, $[M+2]^{+2}$ = 632.8

**TSL6-SFCHYPTAVC-NH$_2$**

Scale: 5.0 mg
Yield: 1.7 mg (29%)
Purity: >95%

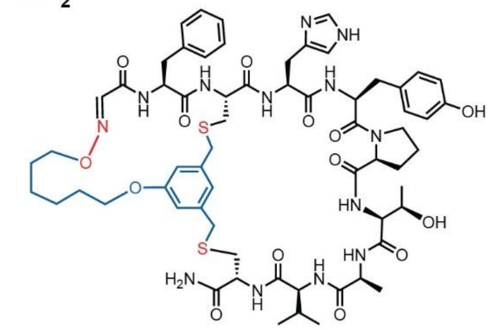

Chemical Formula: $C_{63}H_{83}N_{13}O_{14}S_2$
Exact Mass: 1309.56
Molecular Weight: 1310.55

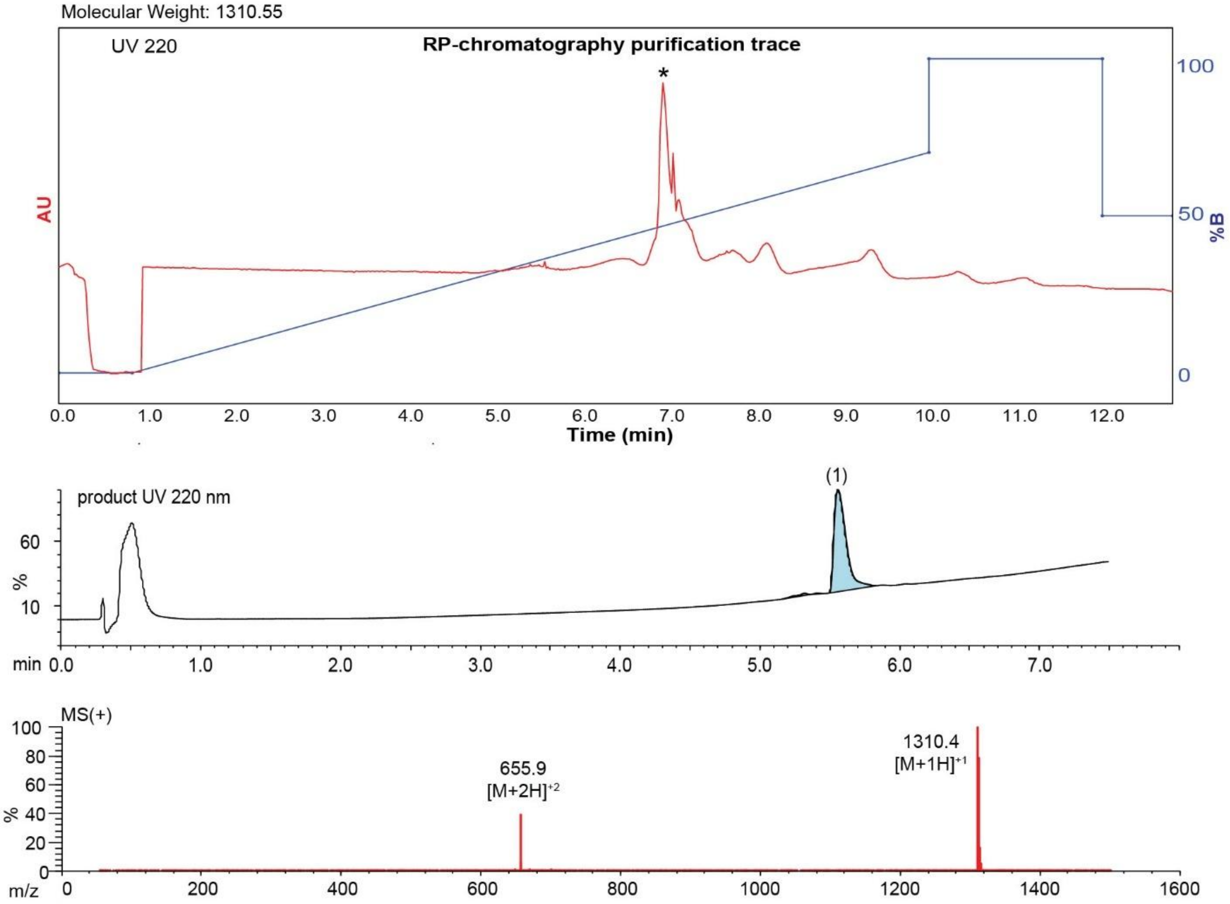

**Figure S93**. Data page for **38c**.

Expected: [M+1] = 1310.56, $[M+2]^{+2}$ = 655.78.

Observed: [M+1] = 1310.4, $[M+2]^{+2}$ = 655.9

### MBX-SFCHYPTAVC-$NH_2$

Scale: 5.0 mg
Yield: 6.3 mg (48%)
Purity: >95%

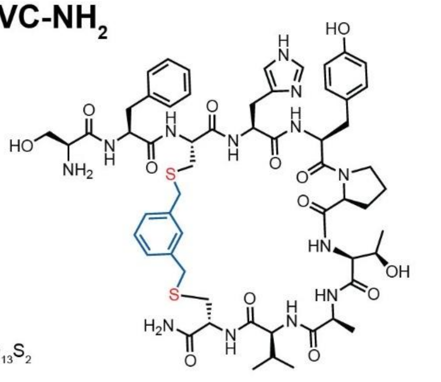

Chemical Formula: $C_{58}H_{77}N_{13}O_{13}S_2$
Exact Mass: 1227.52
Molecular Weight: 1228.45

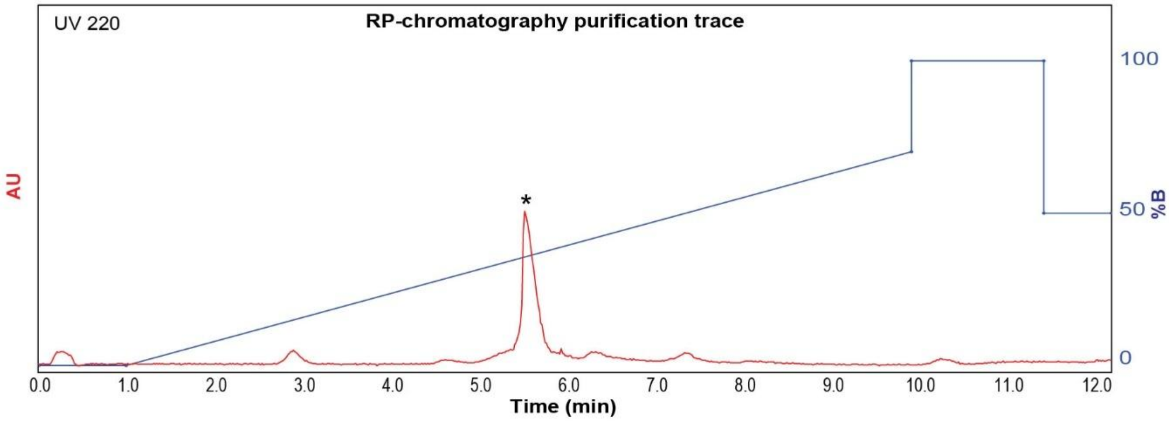

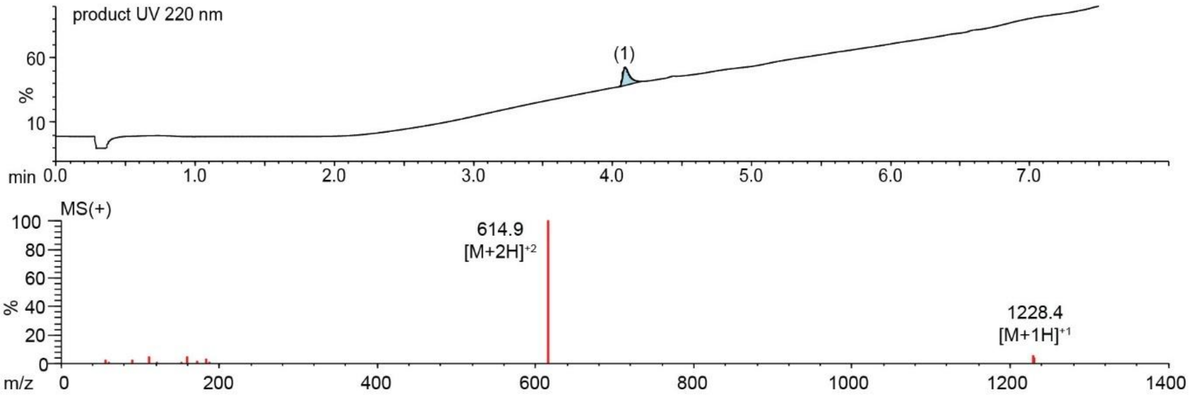

**Figure S94**. Data page for **38f**.

Expected: [M+1] = 1228.52, $[M+2]^{+2}$ = 614.76.

Observed: [M+1] = 1228.4, $[M+2]^{+2}$ = 614.9

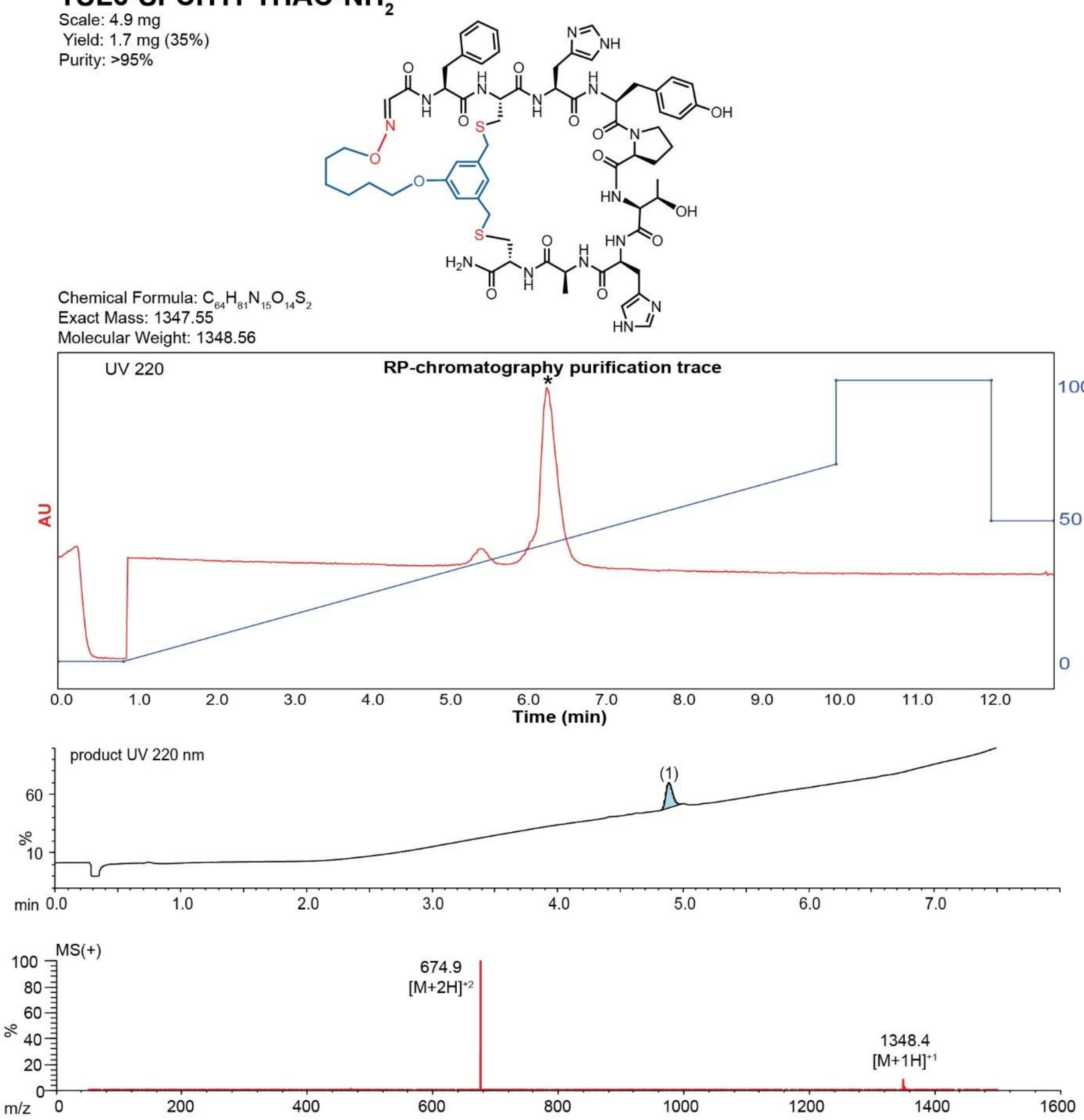

**Figure S95**. Data page for **39c**.

Expected: [M+1] = 1348.55, $[M+2]^{+2}$ = 674.775.

Observed: [M+1] = 1348.4, $[M+2]^{+2}$ = 674.9

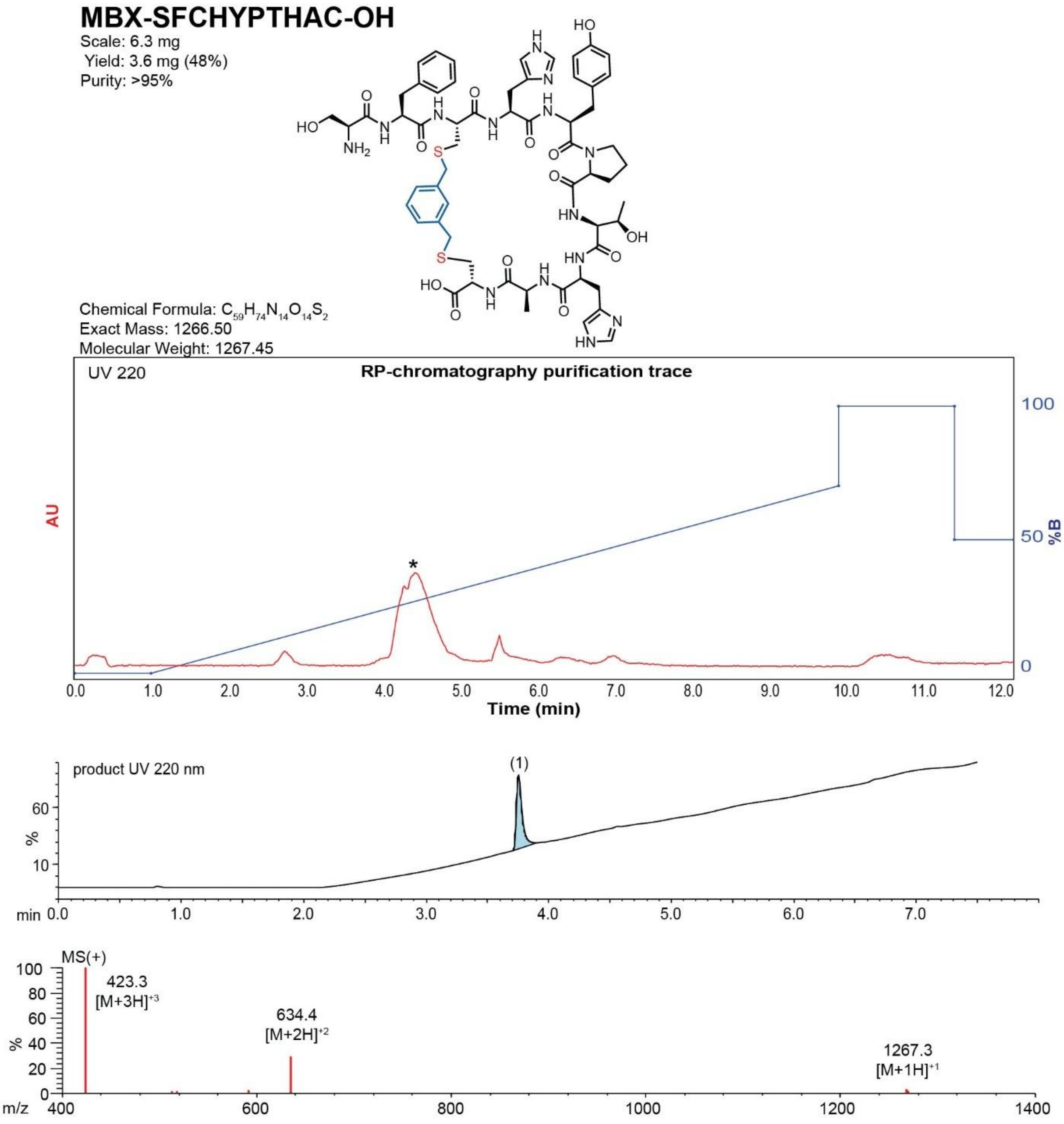

**Figure S96**. Data page for **39f**.

Expected: [M+1] = 1267.50, $[M+2]^{+2}$ = 634.25.

Observed: [M+1] = 1267.3, $[M+2]^{+2}$ = 634.4

**TSL6-SFCHAAAHVC-$NH_2$**

Scale: 6.0 mg
Yield: 1.8 mg (26%)
Purity: >95%

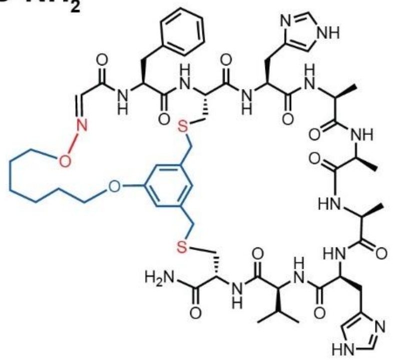

Chemical Formula: $C_{57}H_{77}N_{15}O_{12}S_2$
Exact Mass: 1227.53
Molecular Weight: 1228.46

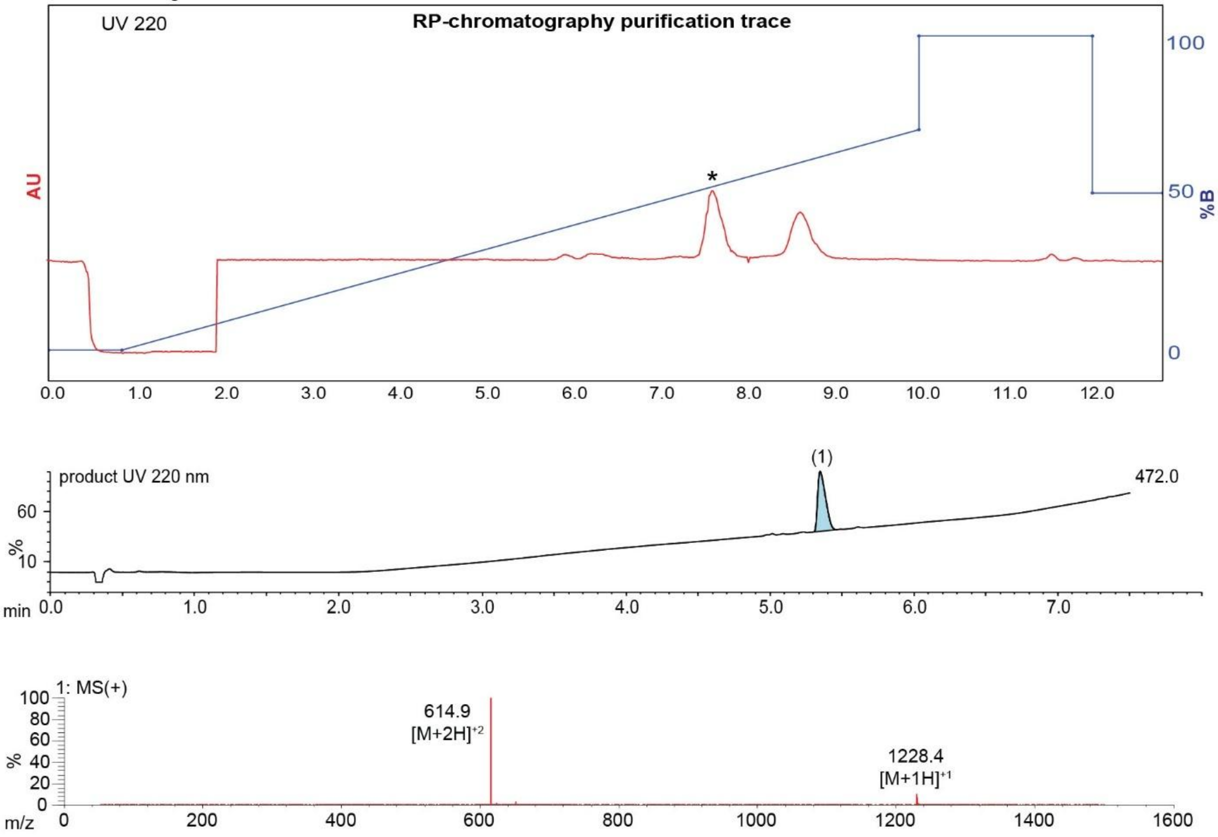

**Figure S97**. Data page for **40c**.

Expected: [M+1] = 1228.53, $[M+2]^{+2}$ = 614.765.

Observed: [M+1] = 1228.4, $[M+2]^{+2}$ = 614.9

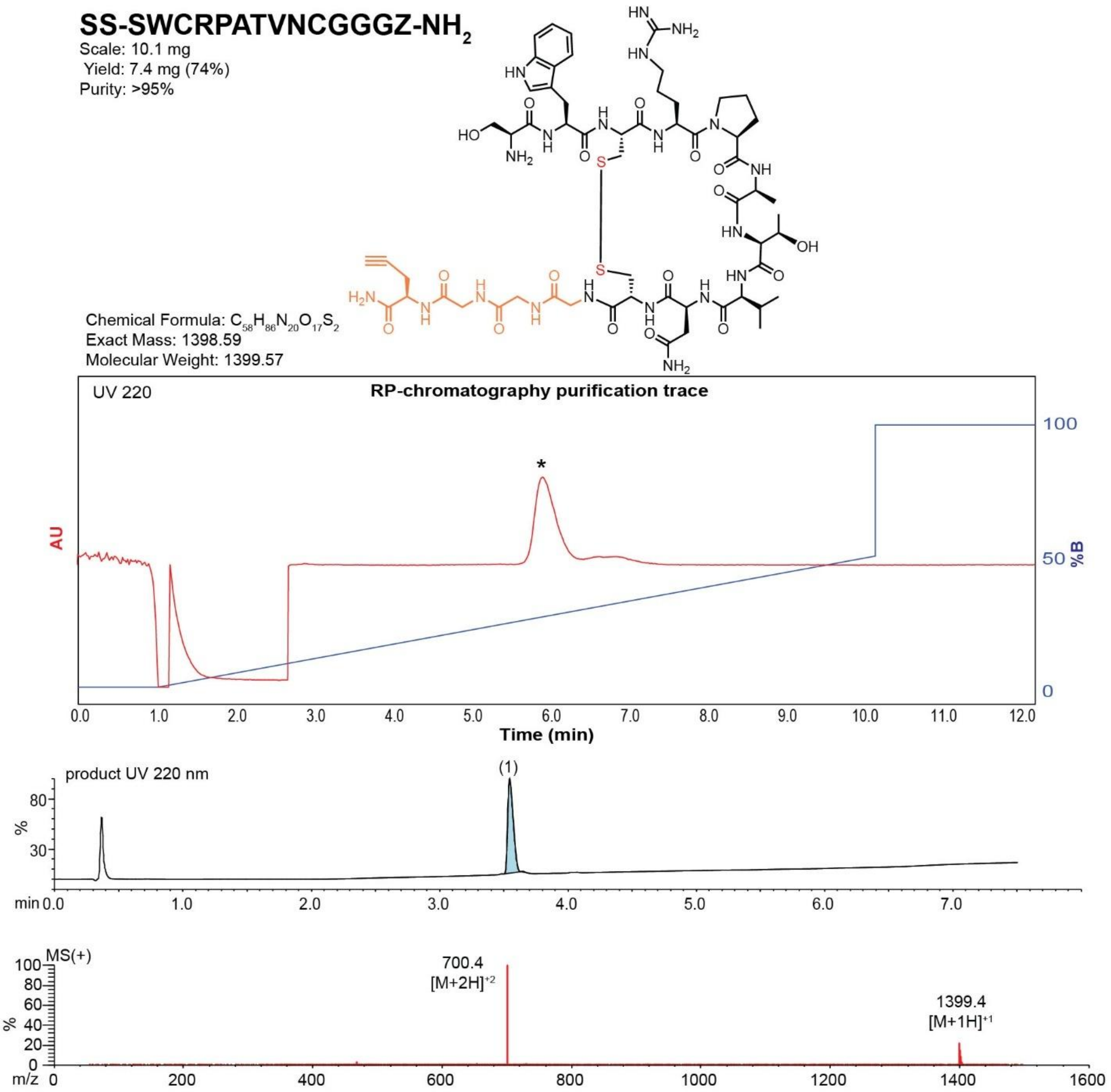

**Figure S98**. Data page for **41b**.

Expected: [M+1] = 1399.59, $[M+2]^{+2}$ = 700.295.

Observed: [M+1] = 1399.4, $[M+2]^{+2}$ = 700.4

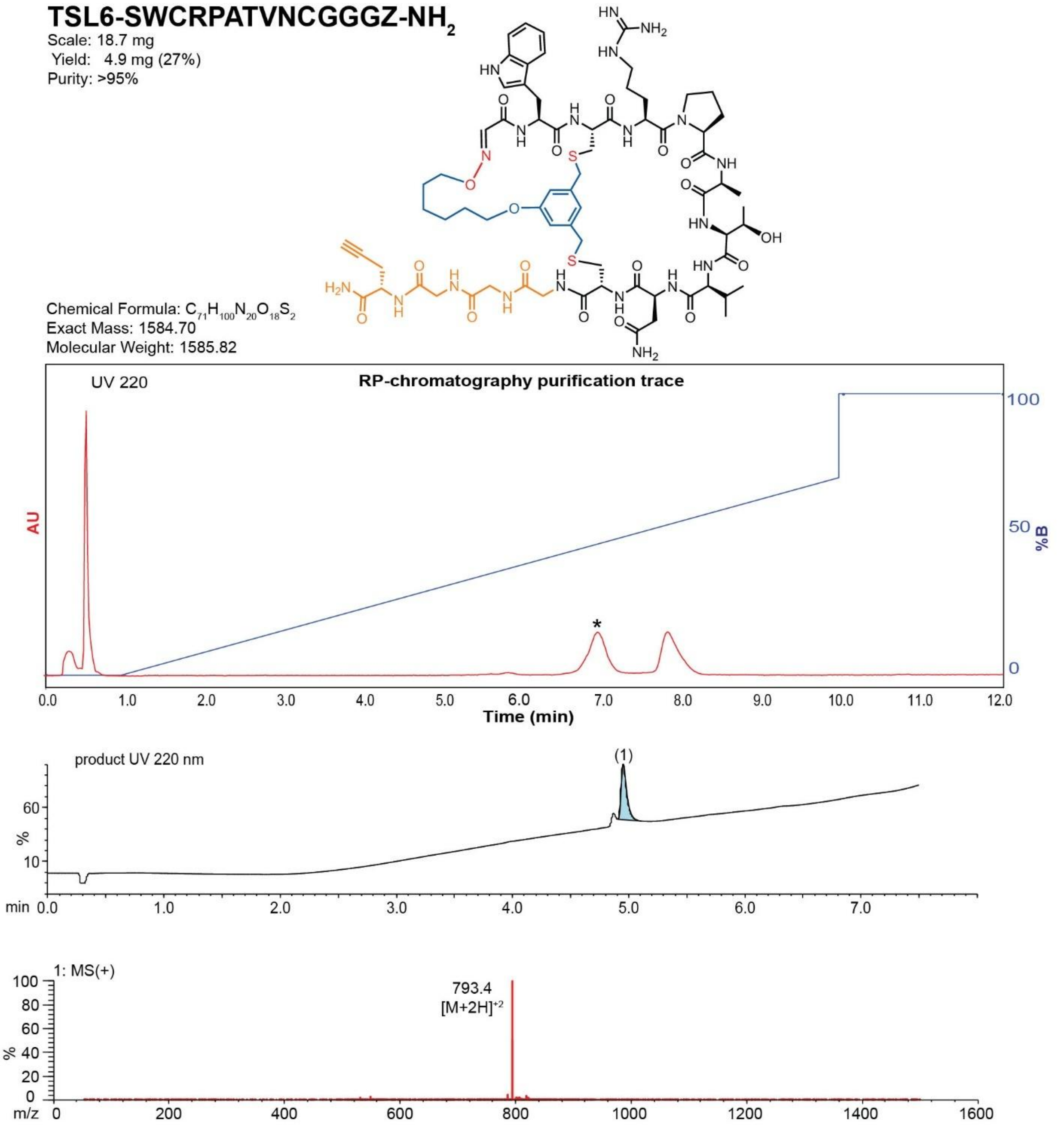

**Figure S99.** Data page for **41c**.

Expected: [M+1] = 1585.70, $[M+2]^{\div 2}$ = 793.45.

Observed: [M+1] = N/A, $[M+2]^{\div 2}$ = 793.4

Note: Detection limit is 1500

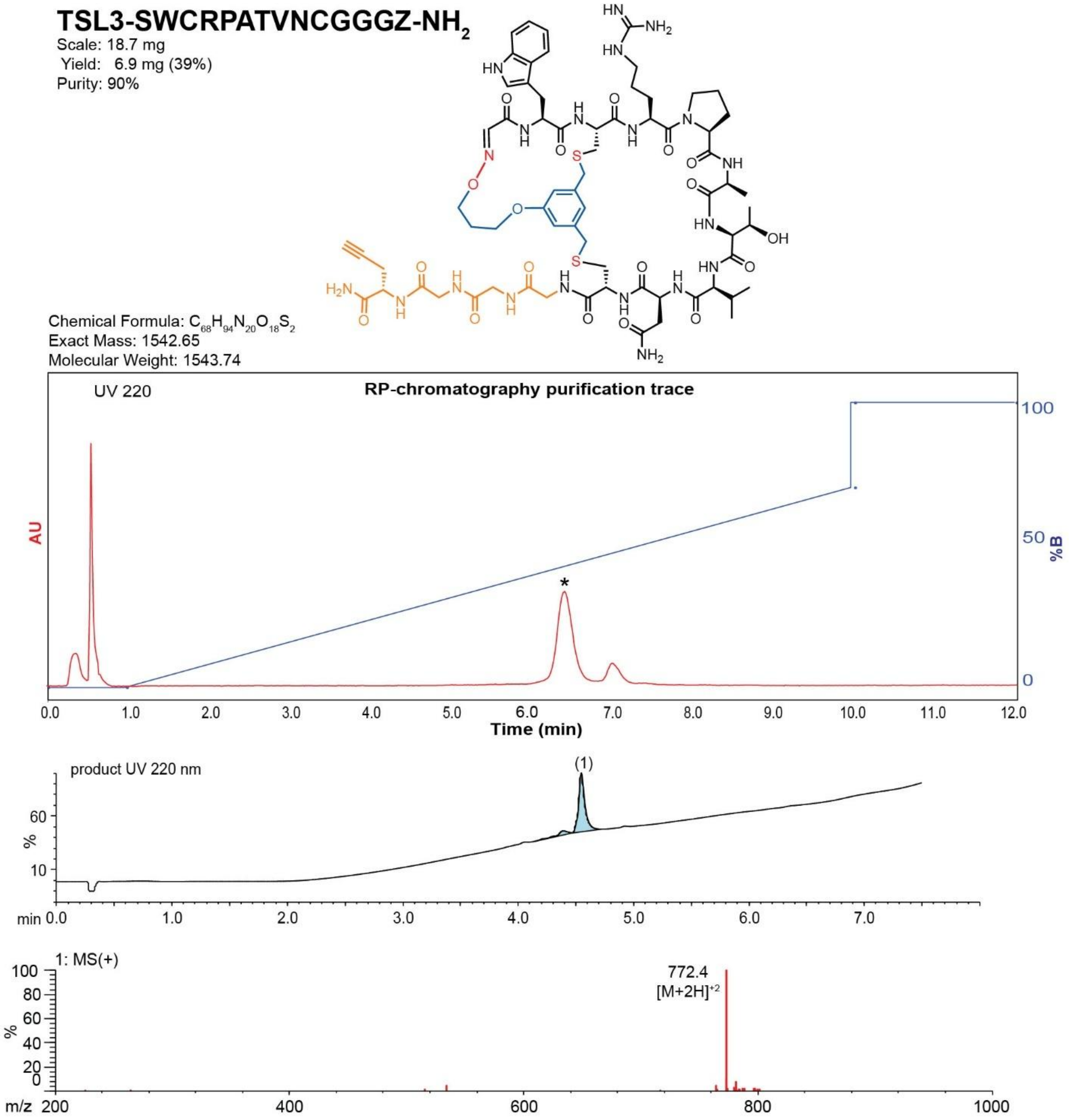

**Figure S100.** Data page for **41d**.

Expected: [M+1] = 1543.65, $[M+2]^{+2}$ = 772.325.

Observed: [M+1] = N/A, $[M+2]^{+2}$ = 772.4

Note: Detection limit is 1500

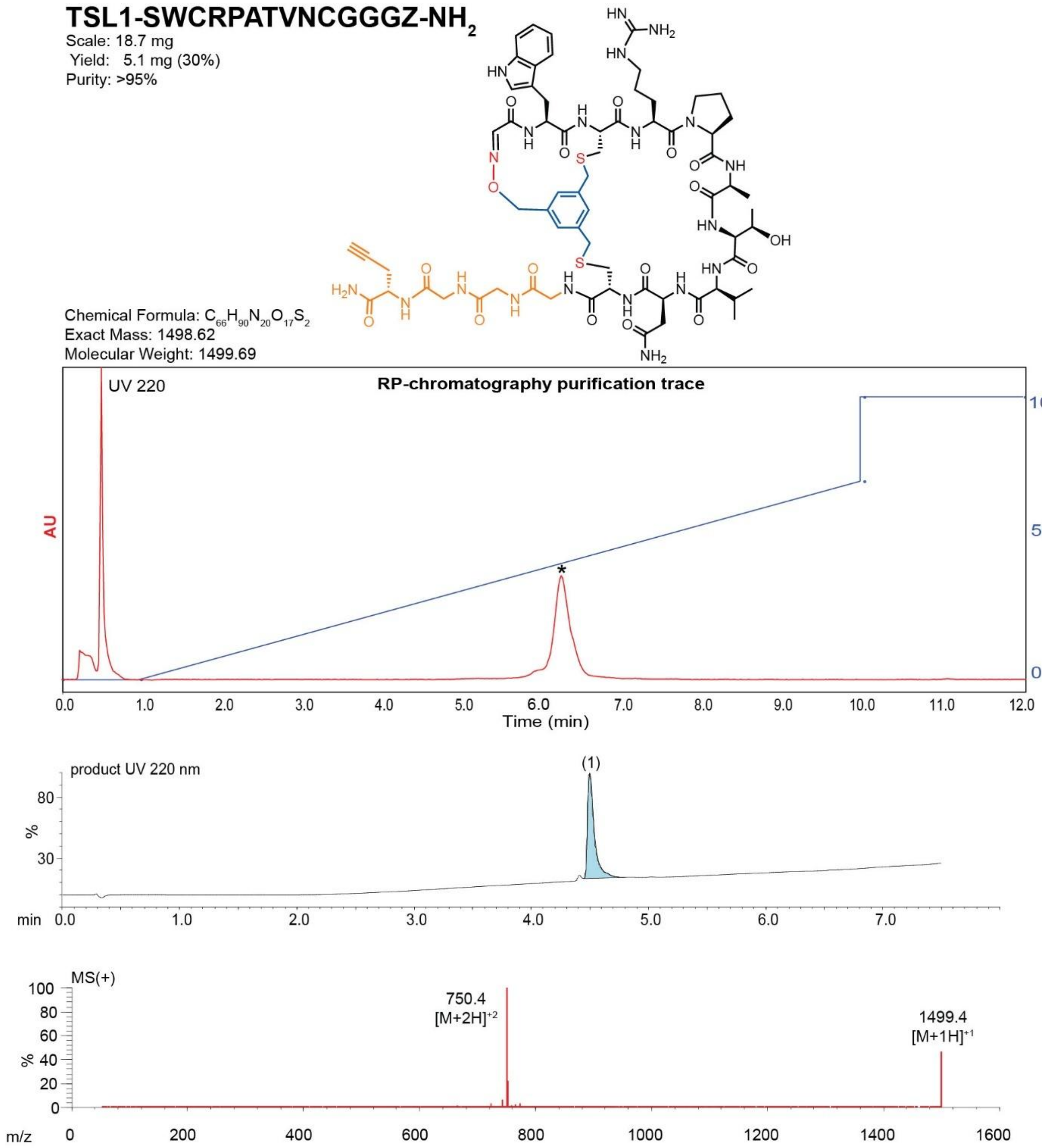

**Figure S101**. Data page for **41e**.

Expected: [M+1] = 1499.62, $[M+2]^{+2}$ = 750.31.

Observed: [M+1] = 1499.4, $[M+2]^{+2}$ = 750.4

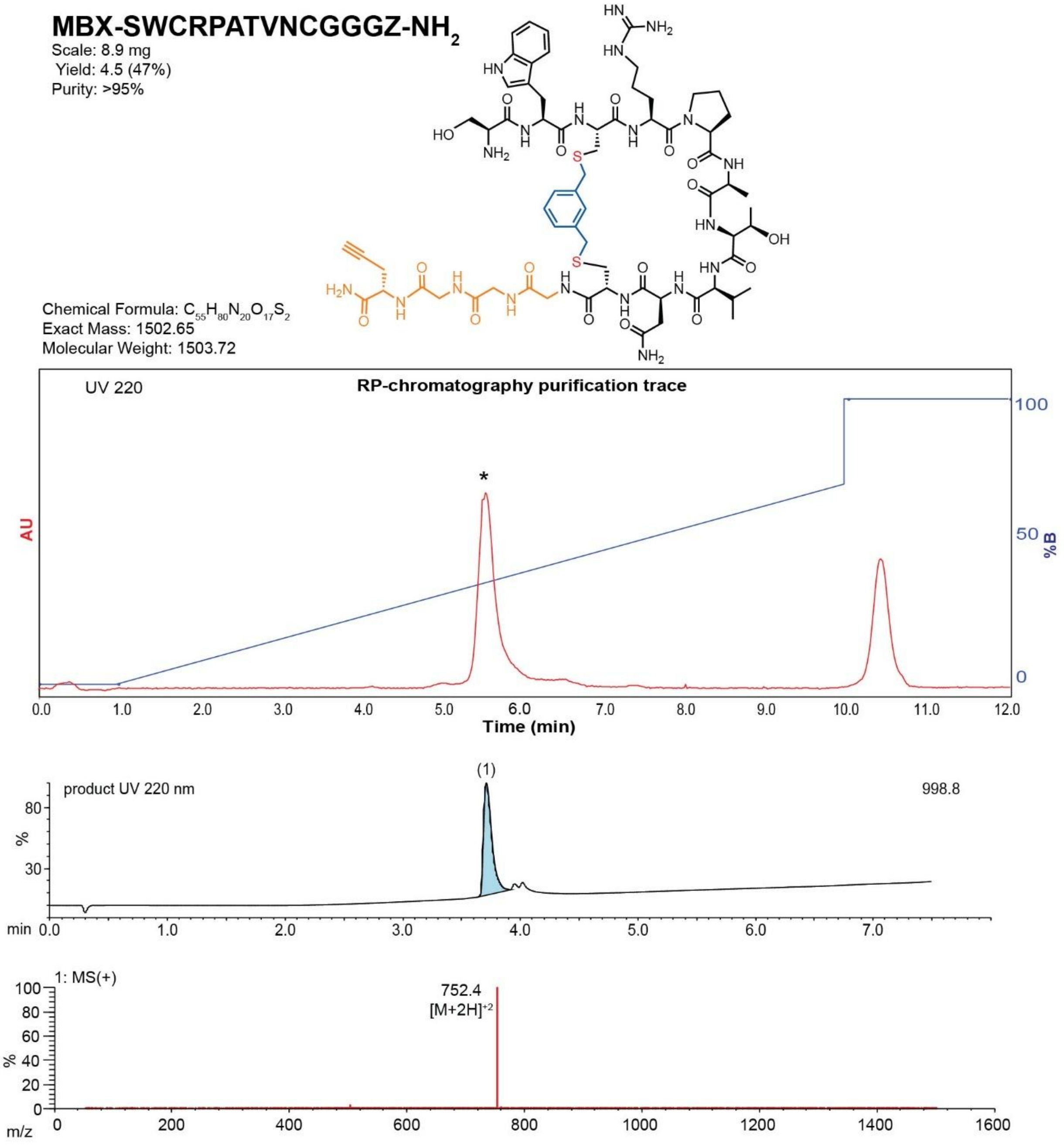

**Figure S102**. Data page for **41f**.

Expected: $[M+1] = 1503.65$, $[M+2]^{\div 2} = 752.325$.

Observed: $[M+1] = N/A$, $[M+2]^{\div 2} = 752.4$

Note: Detection limit is 1500

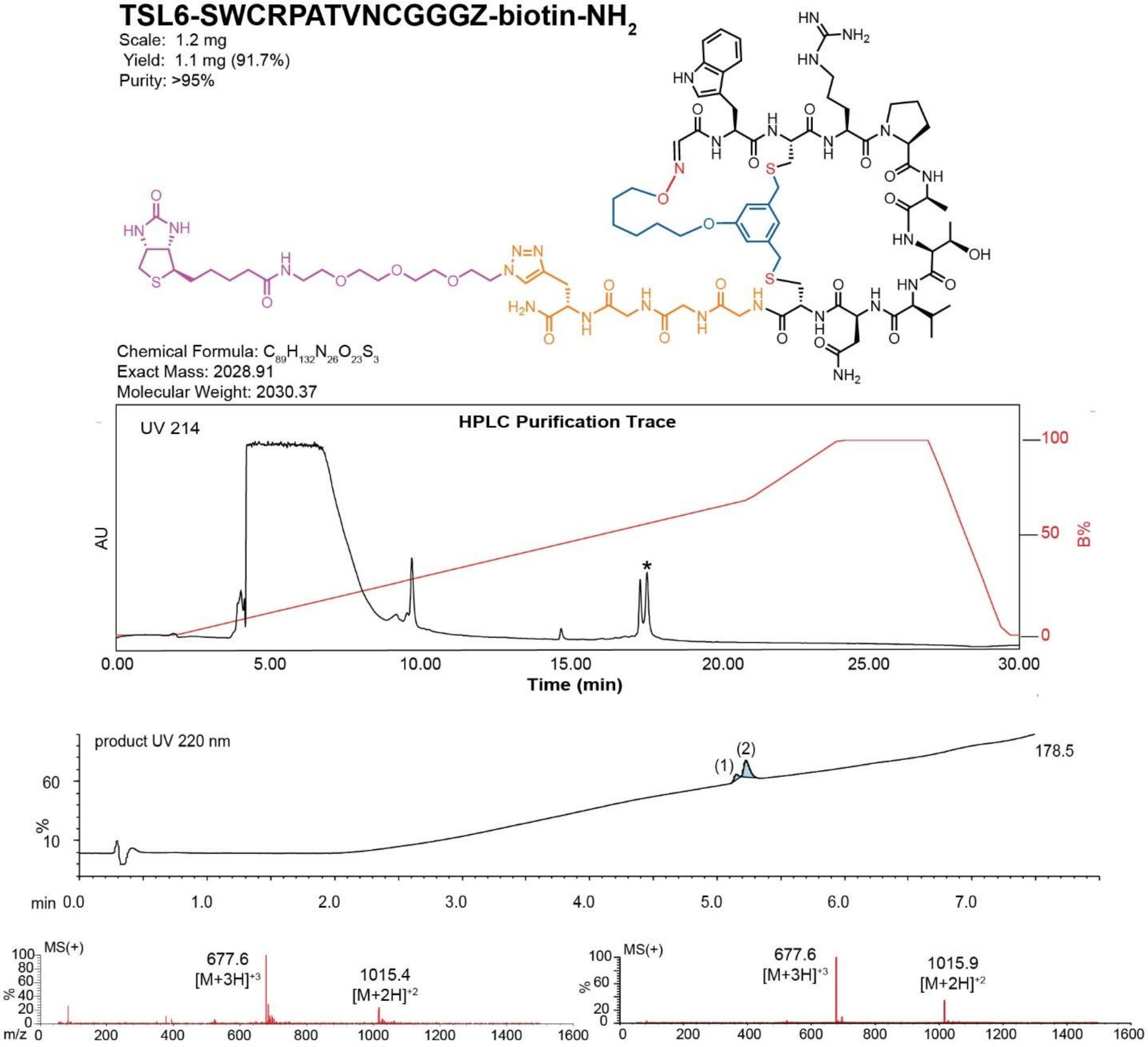

**Figure S103**. Data page for **41g**.

Expected: [M+1] = 2029.91, $[M+2]^{\div 2}$ = 1015.455.

Observed: [M+1] = N/A, $[M+2]^{\div 2}$ = 1015.4

Note: Detection limit is 1500

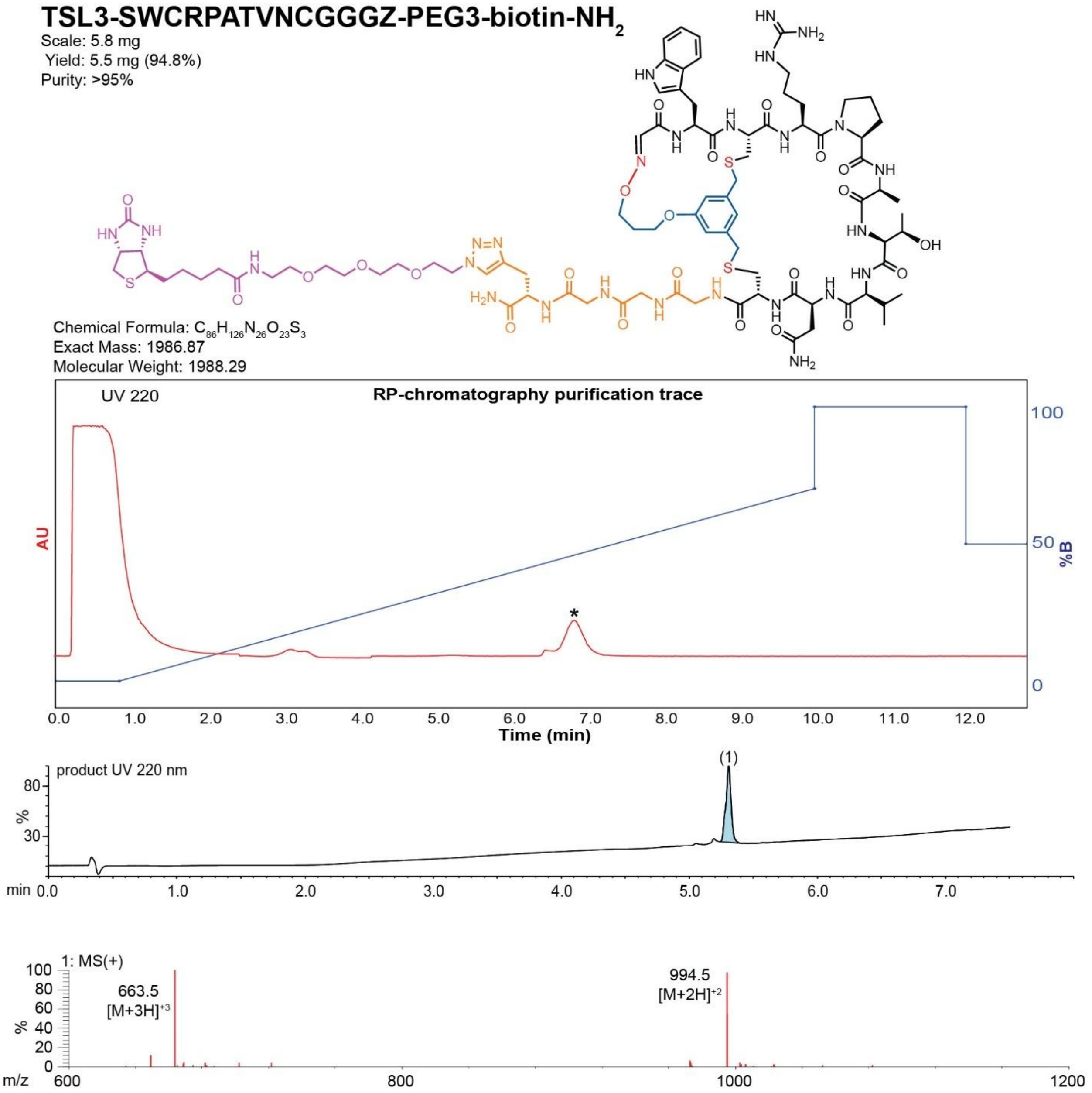

**Figure S104**. Data page for **41h**.

Expected: [M+1] = 1987.87, $[M+2]^{\div 2}$ = 994.435.

Observed: [M+1] = N/A, $[M+2]^{\div 2}$ = 994.5

Note: Detection limit is 1500

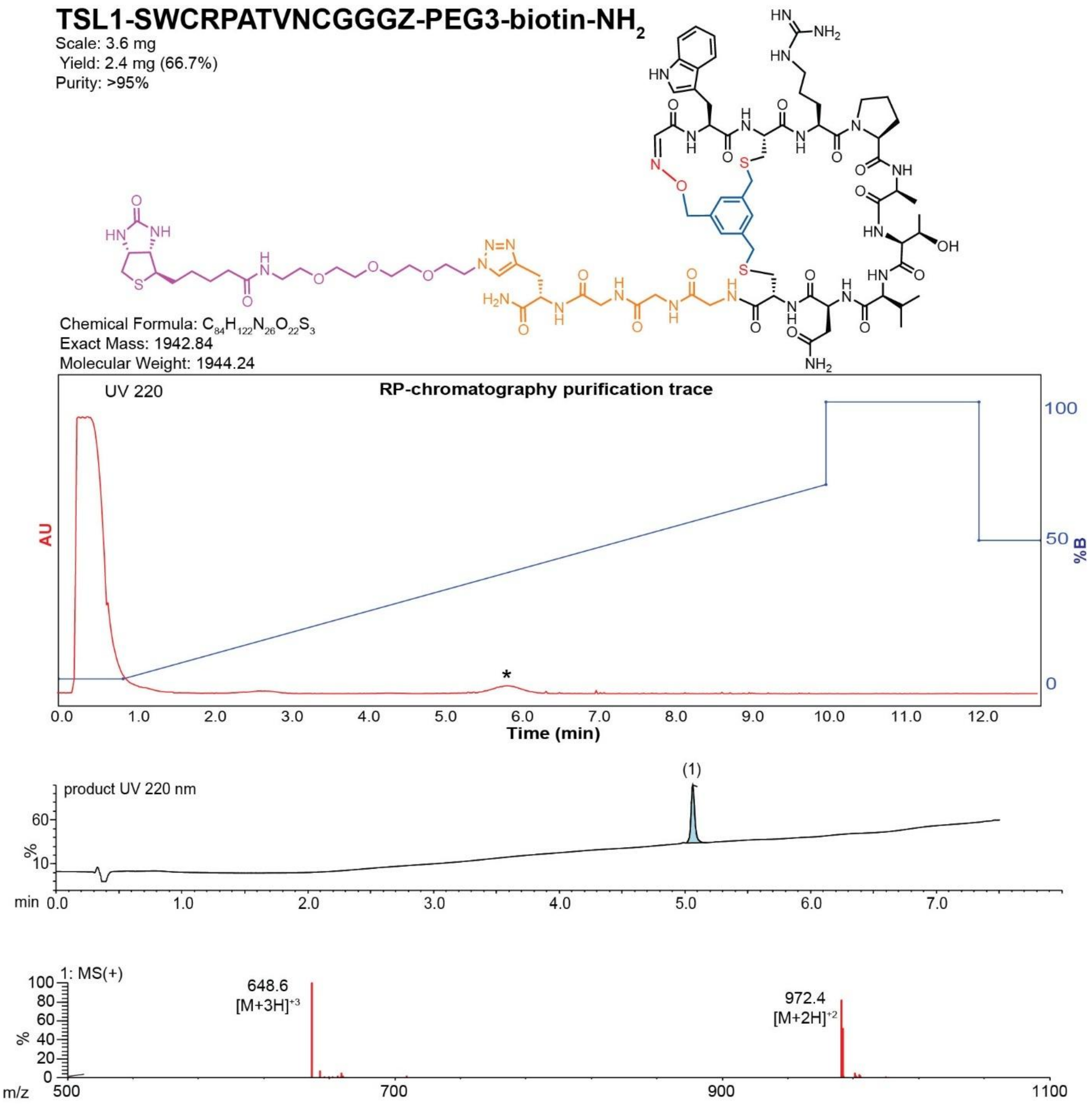

**Figure S105**. Data page for **41i**.

Expected: [M+1] = 1943.84, $[M+2]^{\div 2}$ = 972.42.

Observed: [M+1] = N/A, $[M+2]^{\div 2}$ = 972.4

Note: Detection limit is 1500

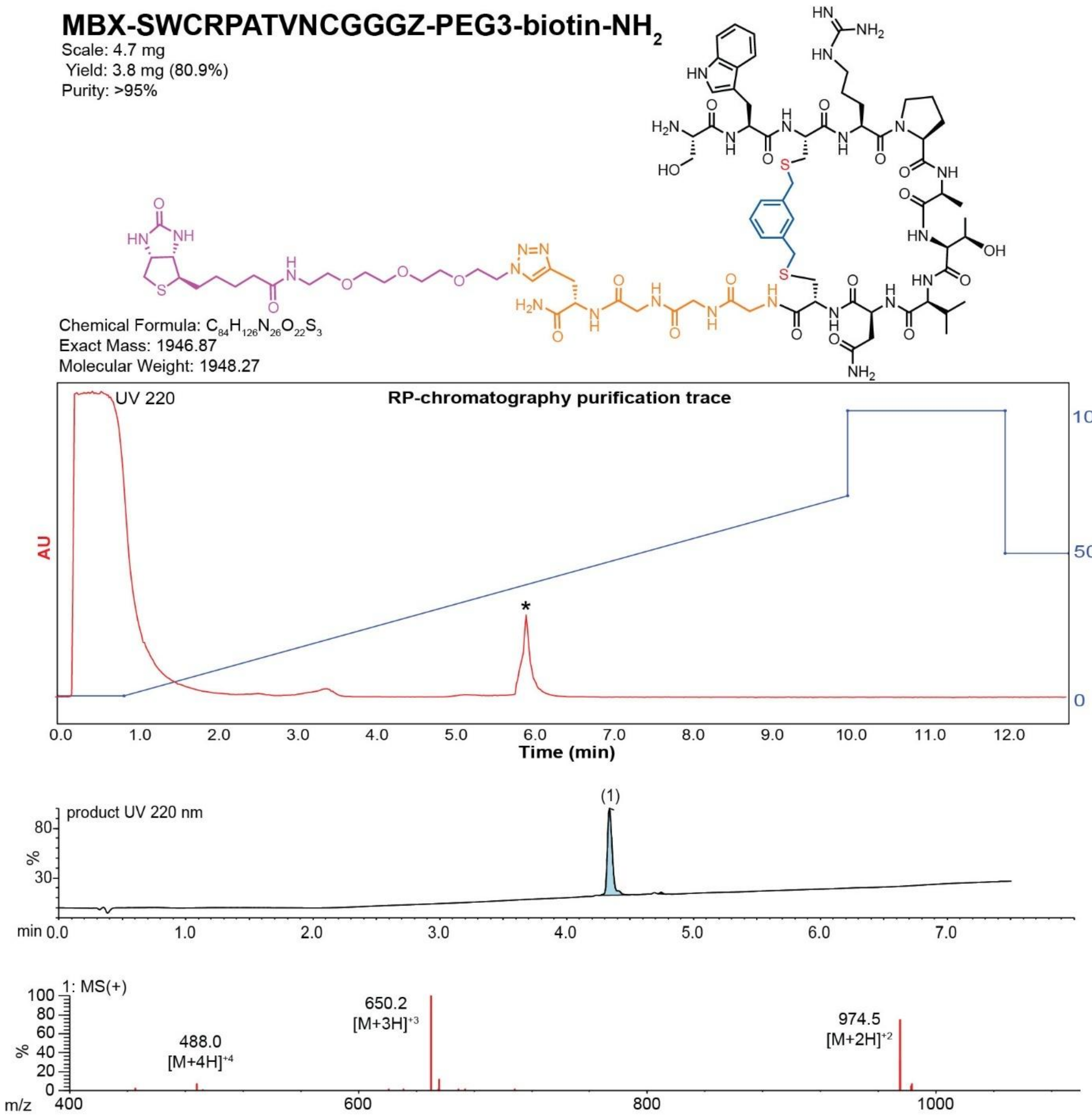

**Figure S106**. Data page for **41j**.

Expected: [M+1] = 1947.87, $[M+2]^{+2}$ = 974.435.

Observed: [M+1] = N/A, $[M+2]^{+2}$ = 974.5

Note: Detection limit is 1500

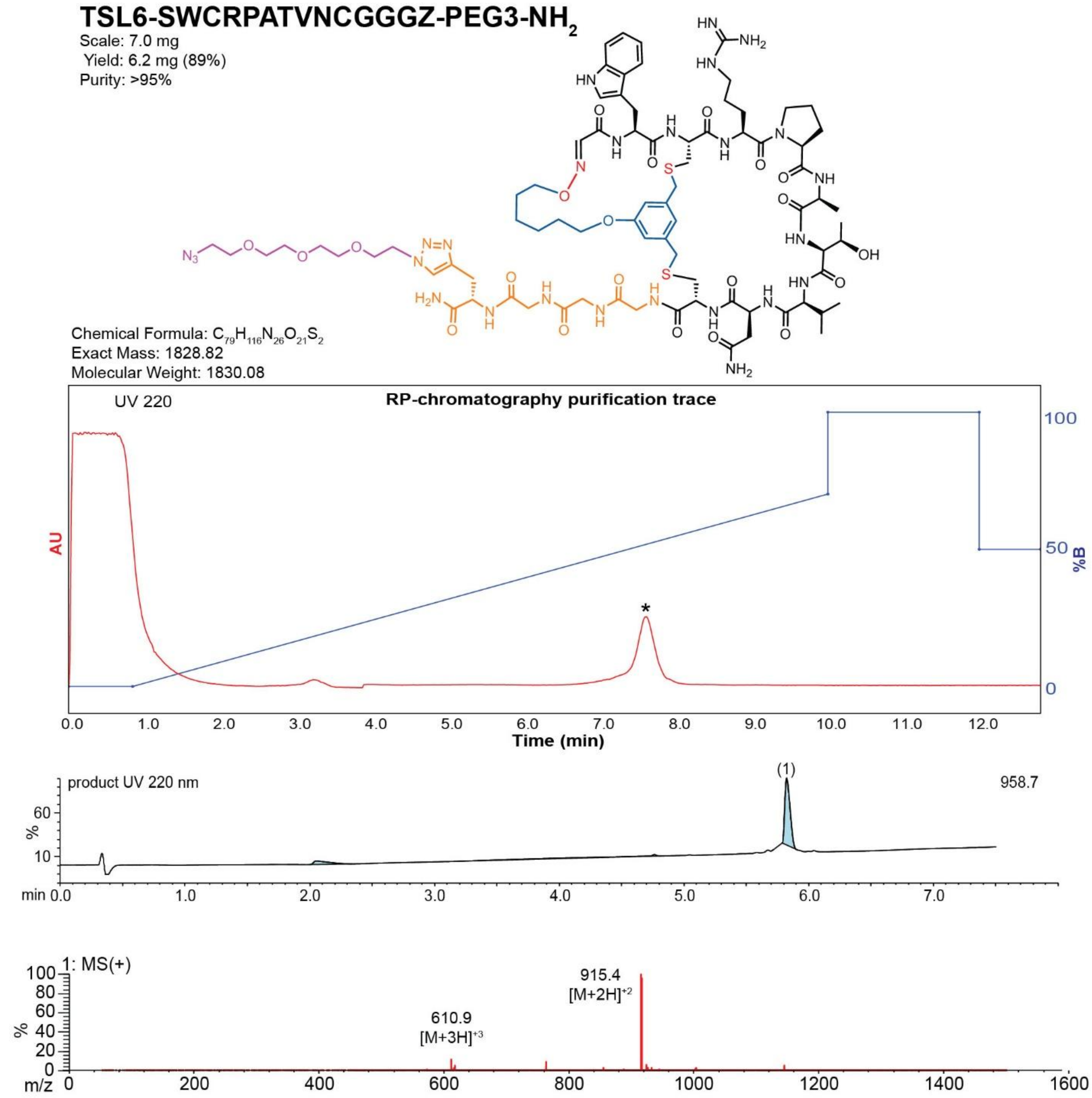

**Figure S107**. Data page for **41k**.

Expected: [M+1] = 1829.82, $[M+2]^{+2}$ = 915.41.

Observed: [M+1] = N/A, $[M+2]^{+2}$ = 915.4

Note: Detection limit is 1500

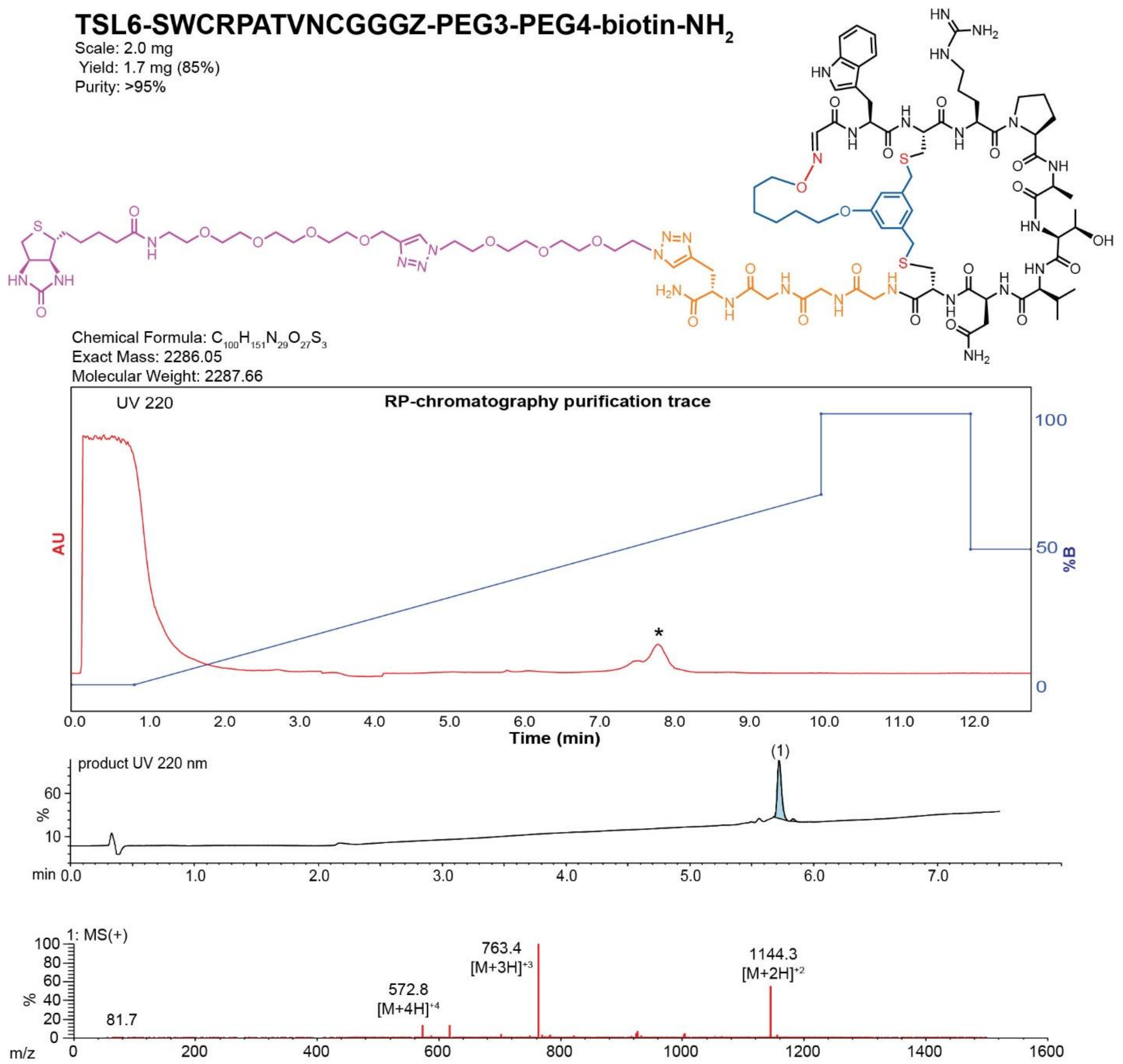

**Figure S108**. Data page for **41l**.

Expected: [M+1] = 2287.05, $[M+2]^{\div 2}$ = 1144.025.

Observed: [M+1] = N/A, $[M+2]^{\div 2}$ = 1144.3

Note: Detection limit is 1500

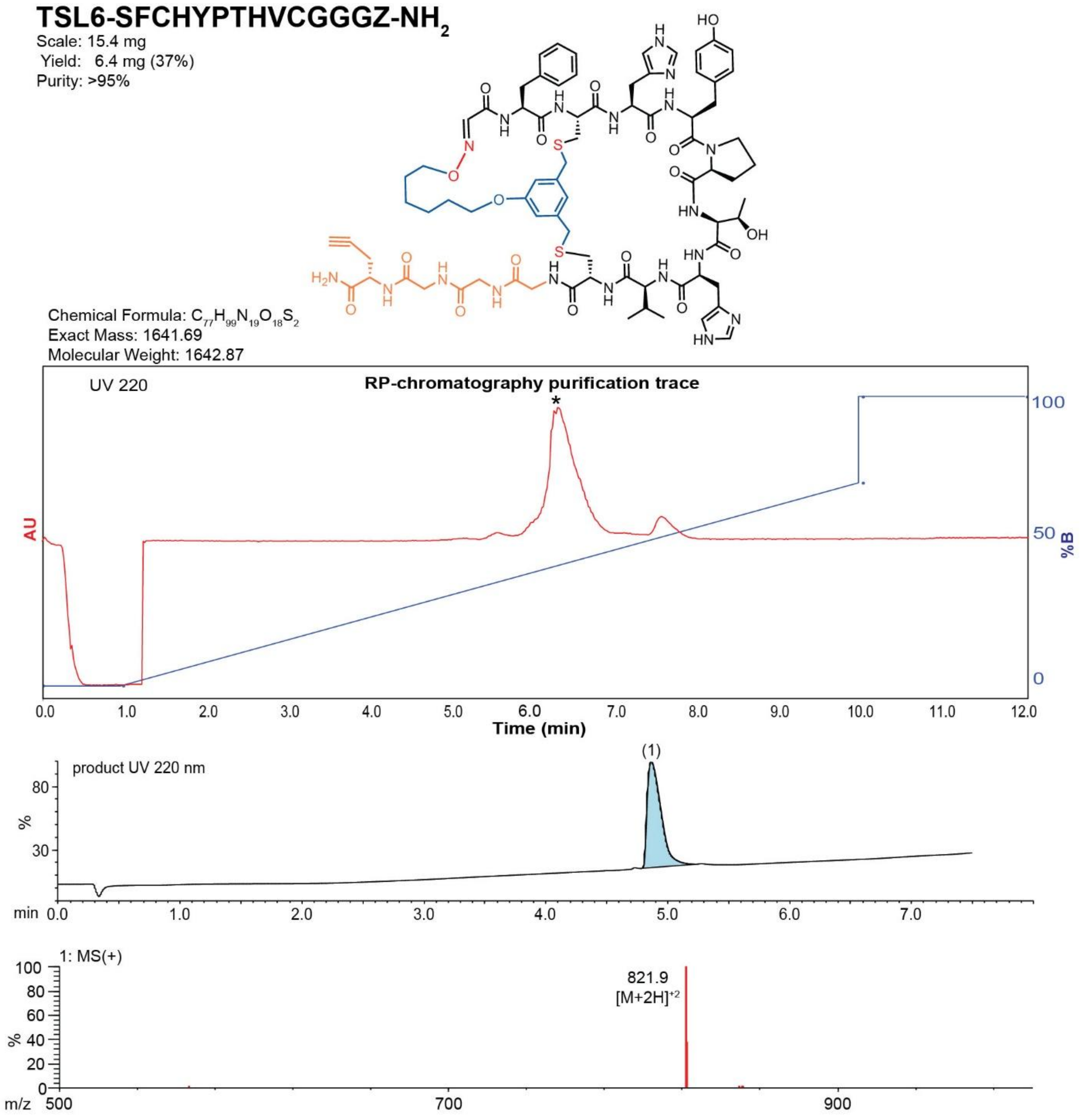

**Figure S109**. Data page for **42c**.

Expected: [M+1] = 1642.69, $[M+2]^{+2}$ = 821.845.

Observed: [M+1] = N/A, $[M+2]^{+2}$ = 821.9

Note: Detection limit is 1500

### TSL3-SFCHYPTHVCGGGZ-$NH_2$

Scale: 15.2 mg
Yield: 6.8 mg (40%)
Purity: >95%

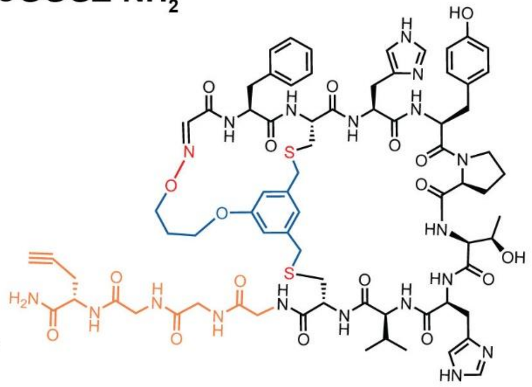

Chemical Formula: $C_{74}H_{93}N_{19}O_{18}S_2$
Exact Mass: 1599.64
Molecular Weight: 1600.79

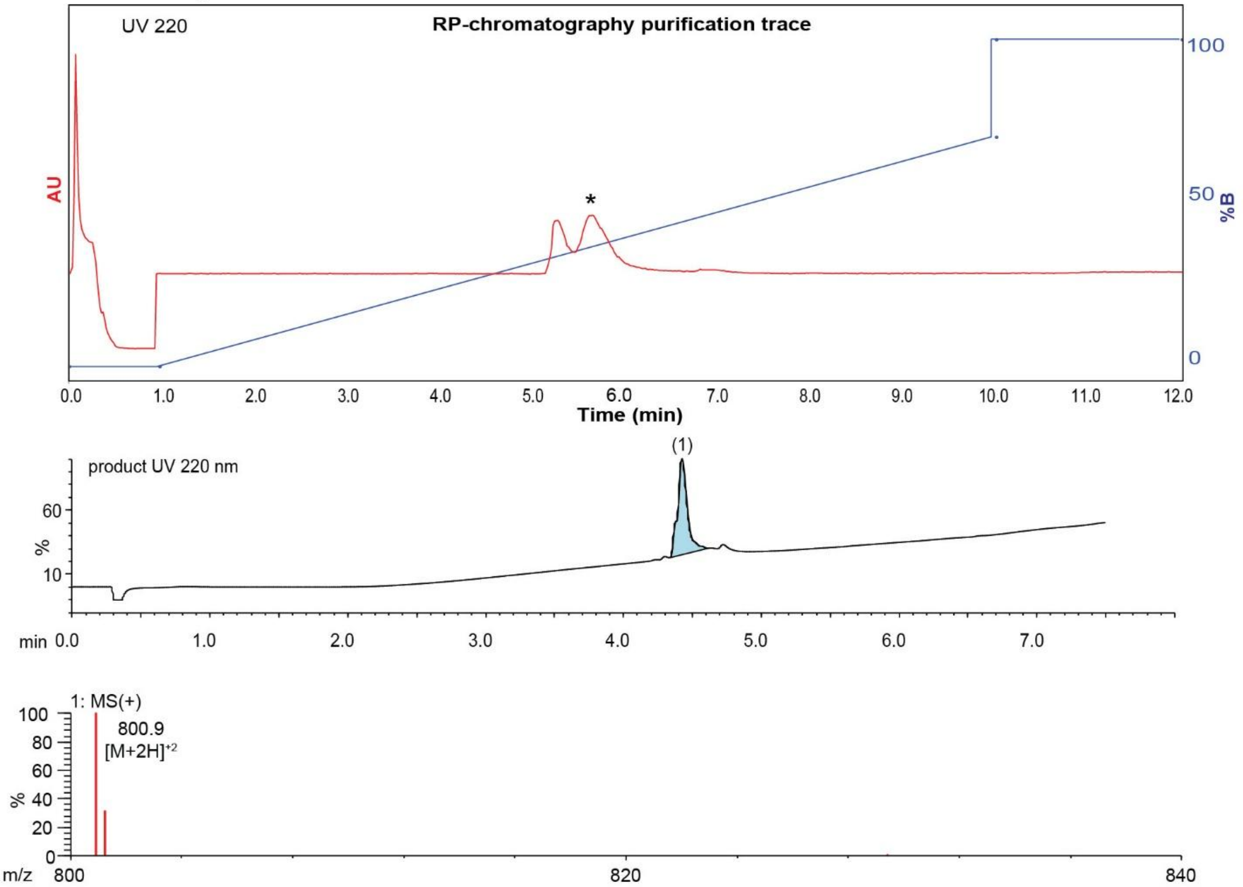

**Figure S110**. Data page for **42d**.

Expected: [M+1] = 1600.64, $[M+2]^{\div 2}$ = 800.82.

Observed: [M+1] = N/A, $[M+2]^{\div 2}$ = 800.9

Note: Detection limit is 1500

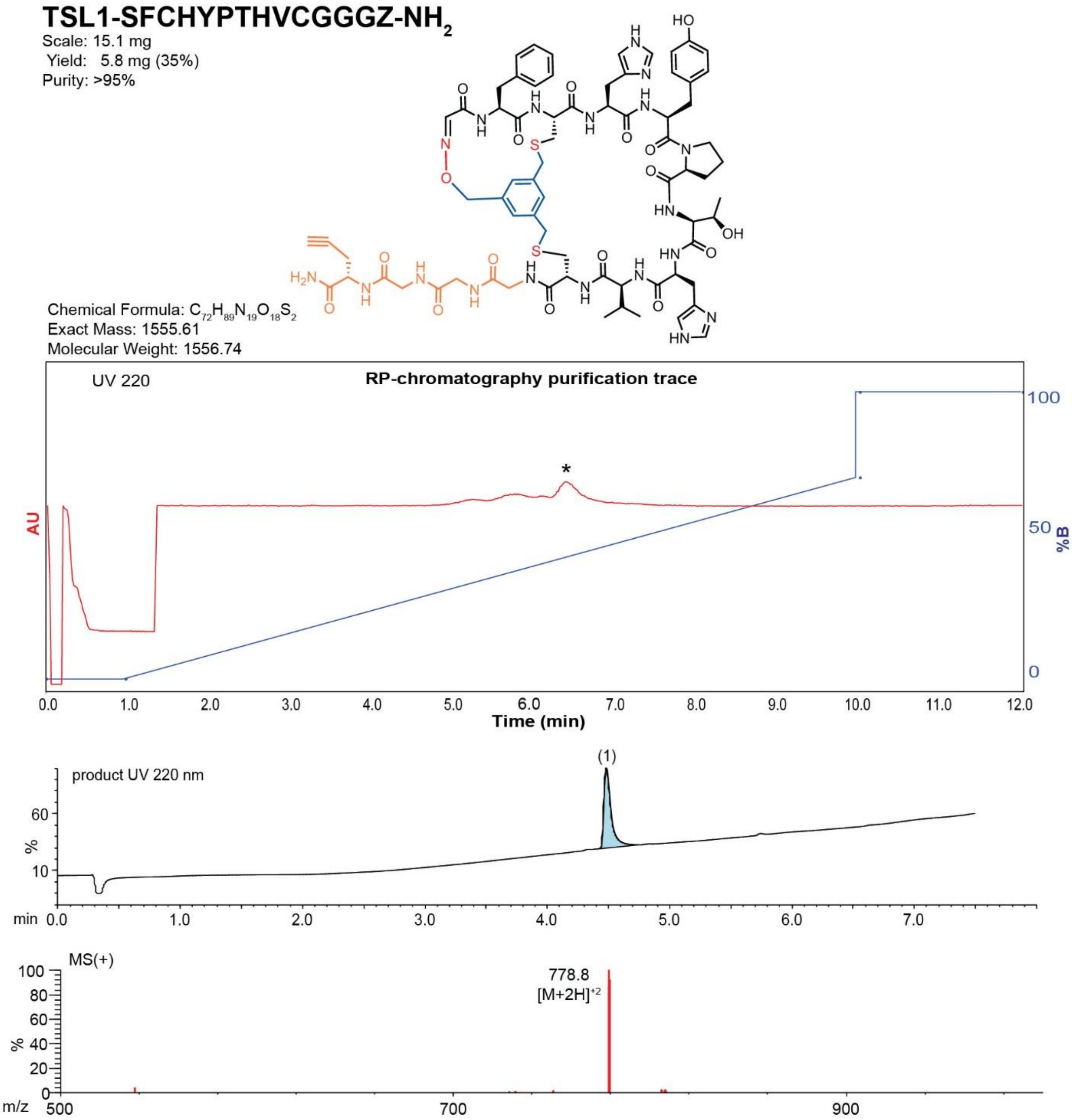

**Figure S111**. Data page for **42e**.

Expected: [M+1] = 1556.61, $[M+2]^{+2}$ = 778.805.

Observed: [M+1] = N/A, $[M+2]^{+2}$ = 778.8

Note: Detection limit is 1500

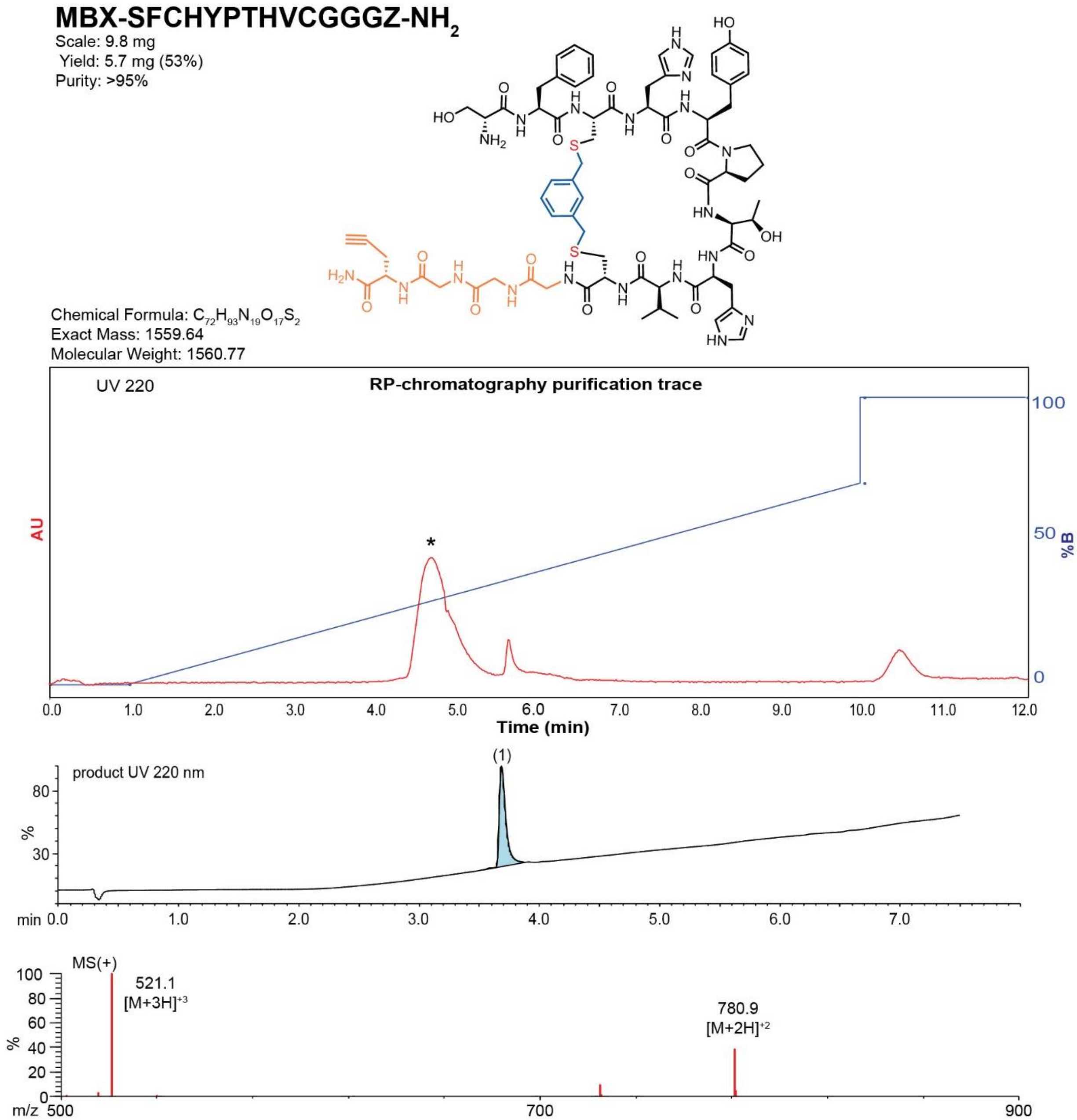

**Figure S112**. Data page for **42f**.

Expected: [M+1] = 1559.64, $[M+2]^{+2}$ = 780.82.

Observed: [M+1] = N/A, $[M+2]^{+2}$ = 780.9

Note: Detection limit is 1500

**TSL6-SYCHPTHVNCGGGZ-PEG3-biotin-$NH_2$**

Scale: 1.5 mg

Yield: 1.5 mg (79%)

Purity: >95%

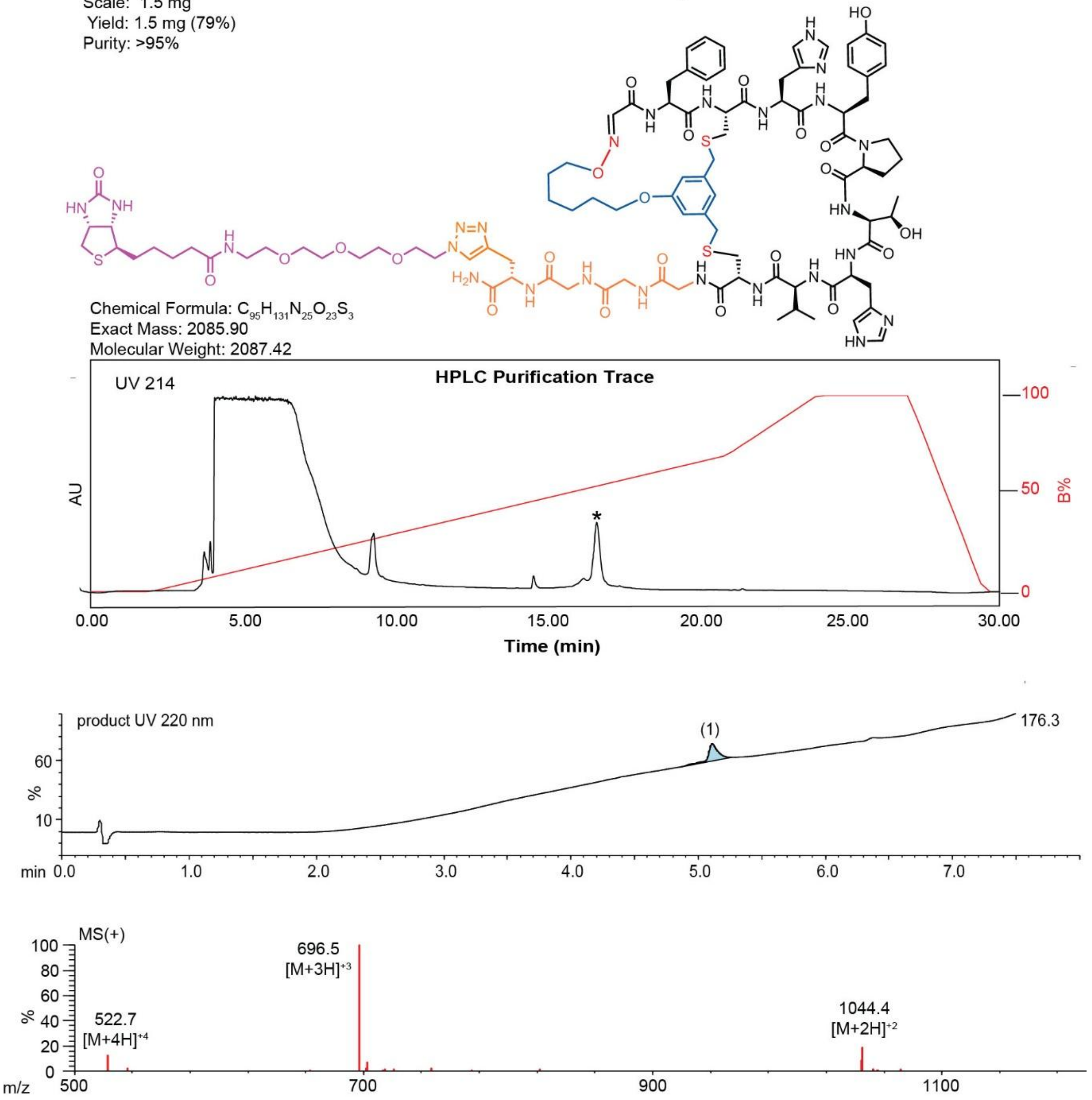

**Figure S113**. Data page for **42g**.

Expected: [M+1] = 2086.90, $[M+2]^{+2}$ = 1043.95.

Observed: [M+1] = N/A, $[M+2]^{+2}$ = 1044.4

Note: Detection limit is 1500

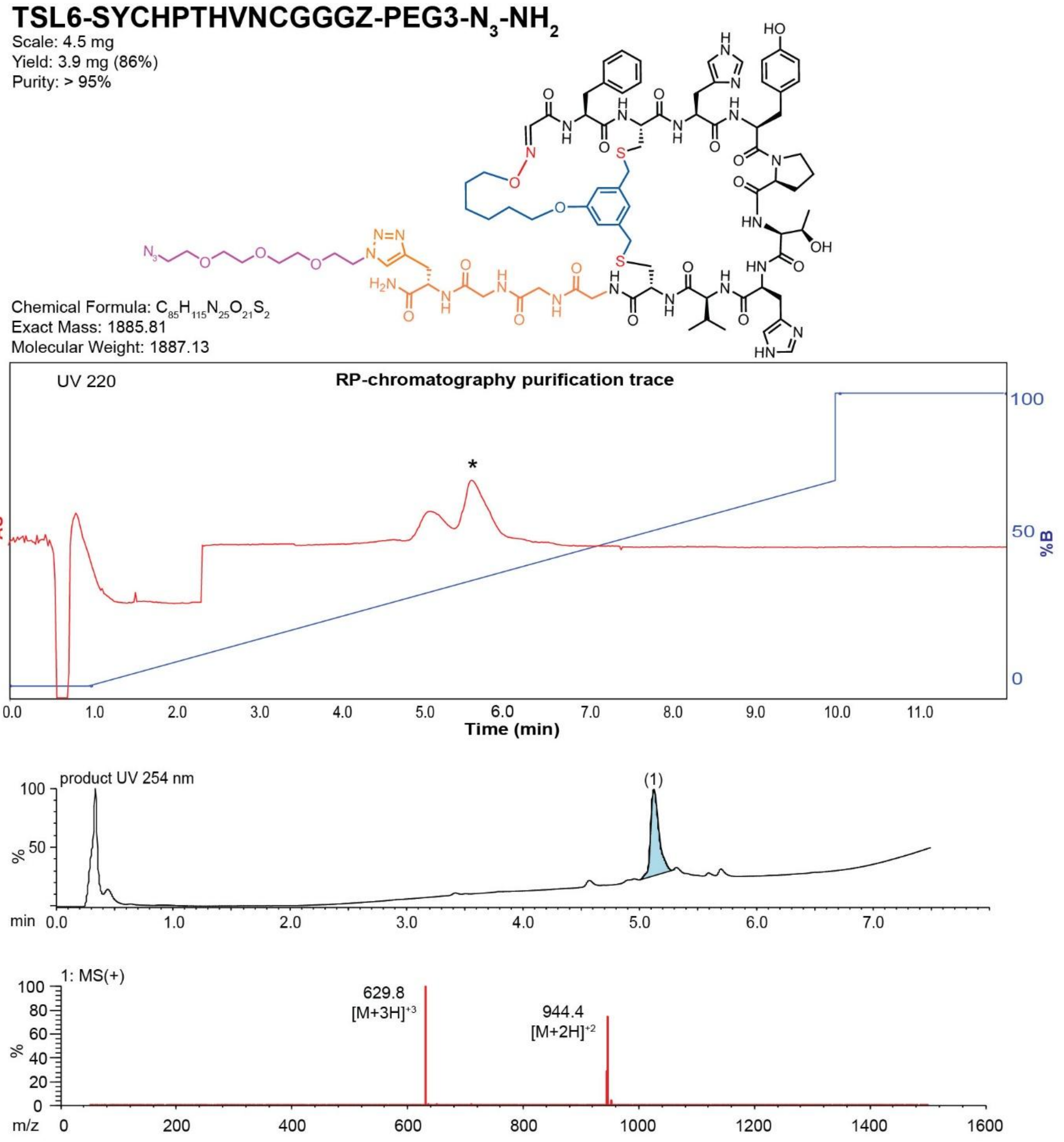

**Figure S114**. Data page for **42k**.

Expected: [M+1] = 1886.81, $[M+2]^{+2}$ = 943.905.

Observed: [M+1] = N/A, $[M+2]^{+2}$ = 944.4

Note: Detection limit is 1500

**TSL6-SYCHPTHVNCGGGZ-PEG3-PEG4-biotin-$NH_2$**

Scale: 2.5 mg
Yield: 3.9 mg (48%)
Purity: >95%

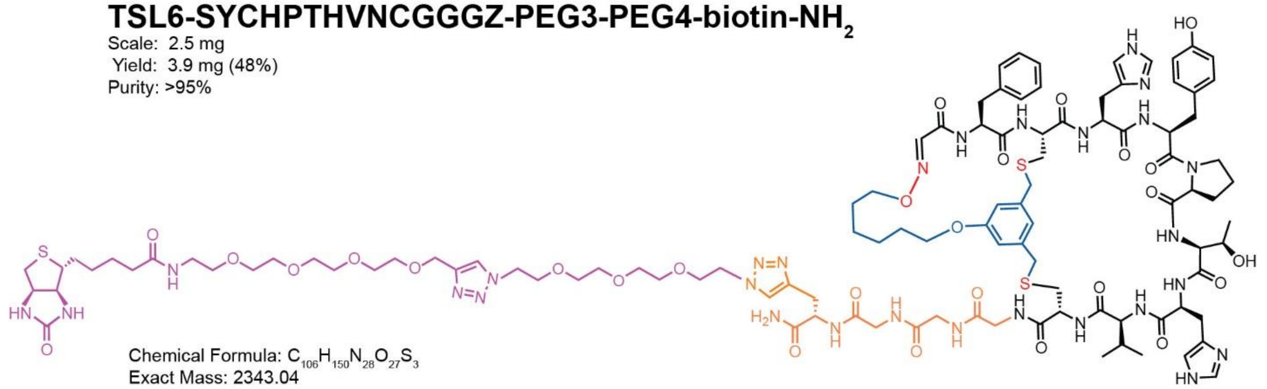

Chemical Formula: $C_{106}H_{150}N_{28}O_{27}S_3$
Exact Mass: 2343.04
Molecular Weight: 2344.71

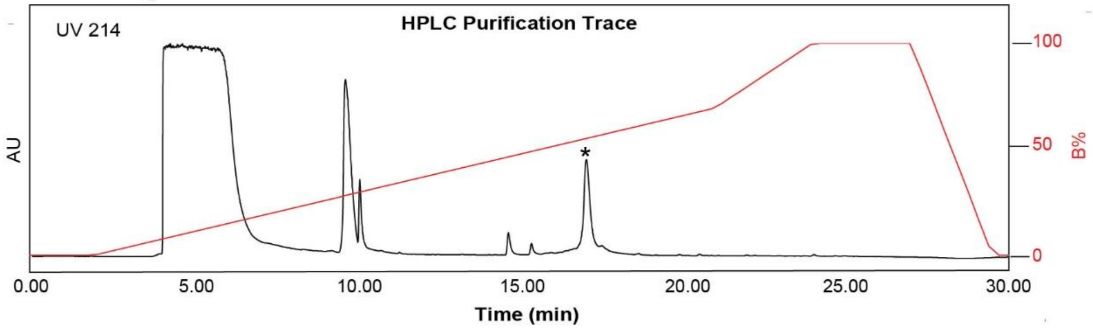

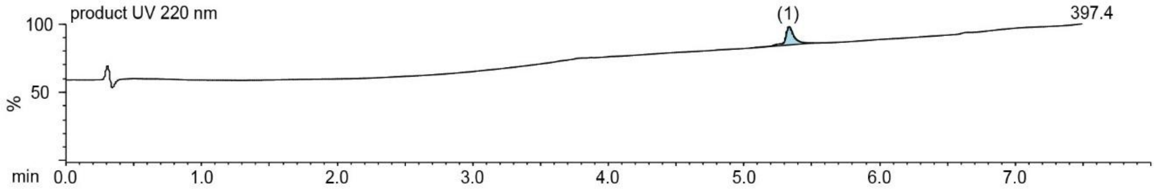

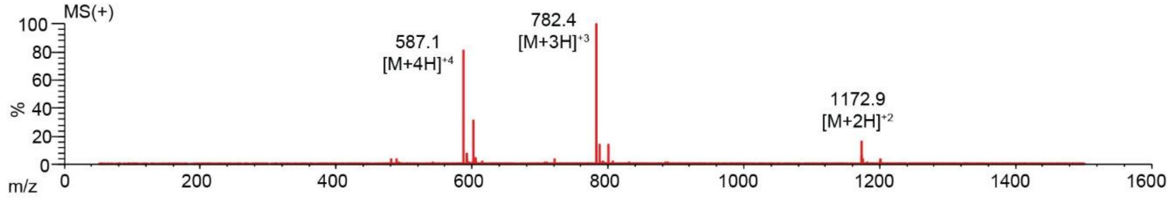

**Figure S115**. Data page for **42l**.

Expected: [M+1] = 2344.04, $[M+2]^{\div 2}$ = 1172.52.

Observed: [M+1] = N/A, $[M+2]^{\div 2}$ = 1172.9

Note: Detection limit is 1500

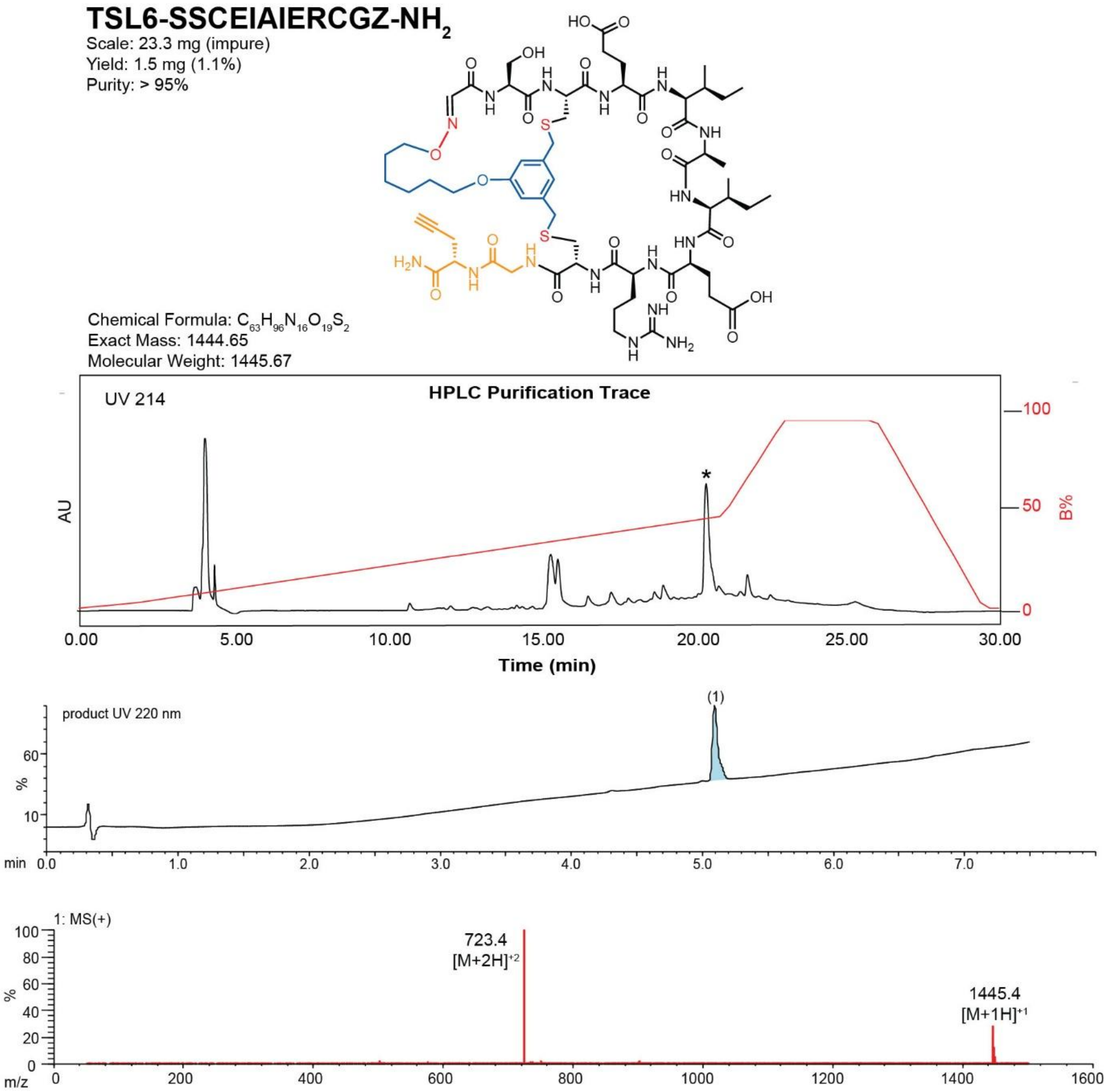

**Figure S116**. Data page for **43c**.

Expected: [M+1] = 1445.65, $[M+2]^{+2}$ = 723.325.

Observed: [M+1] = 1445.4, $[M+2]^{+2}$ = 723.4

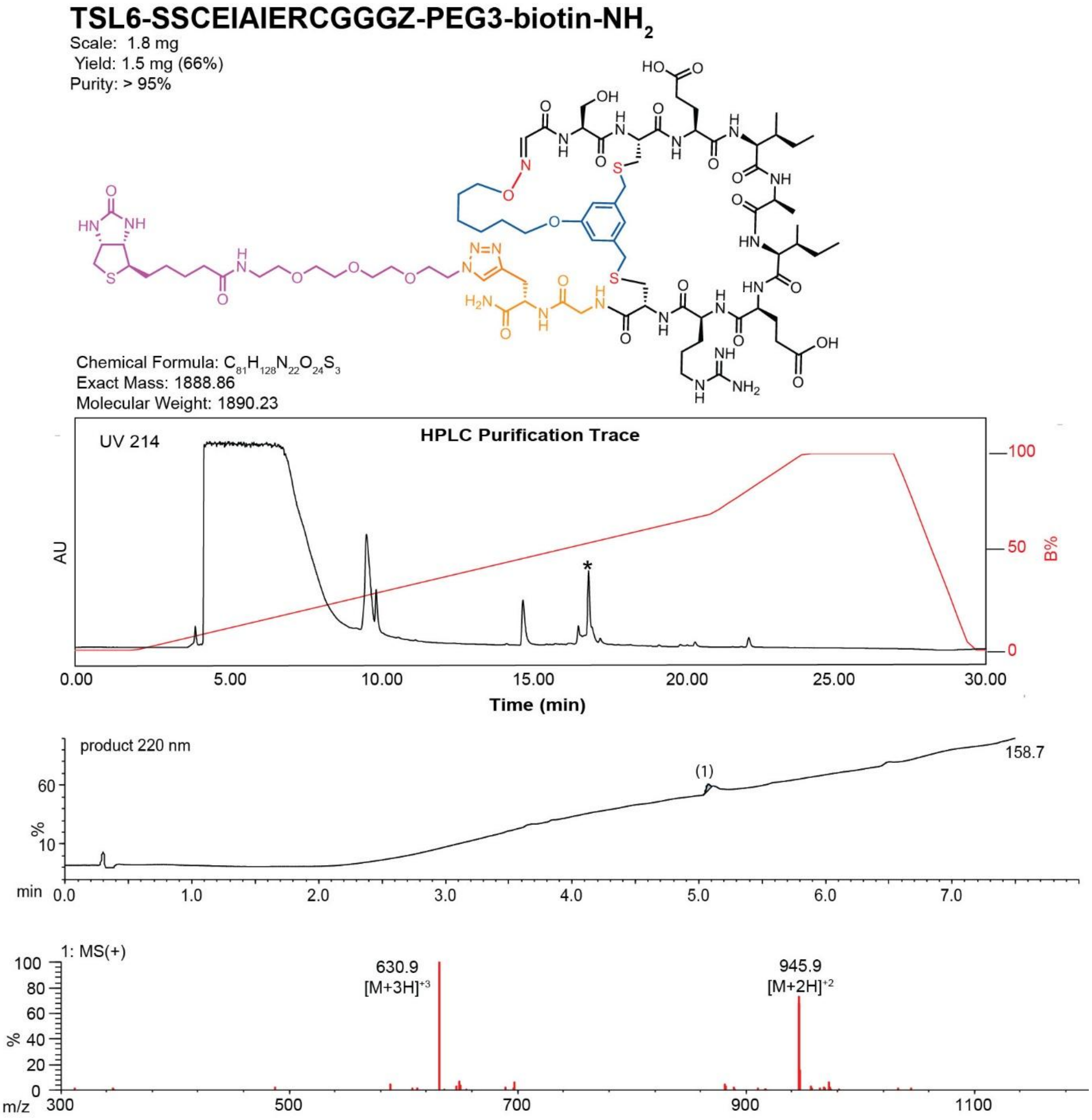

**Figure S117**. Data page for **43g**.

Expected: [M+1] = 1889.86, $[M+2]^{\div 2}$ = 945.43.

Observed: [M+1] = N/A, $[M+2]^{\div 2}$ = 945.9

Note: Detection limit is 1500

### TSL1-SAAAAAWCRPATVNCGGGZ-$NH_2$

Scale: 20.3 mg
Yield: 6.9 mg (32%)
Purity: >95%

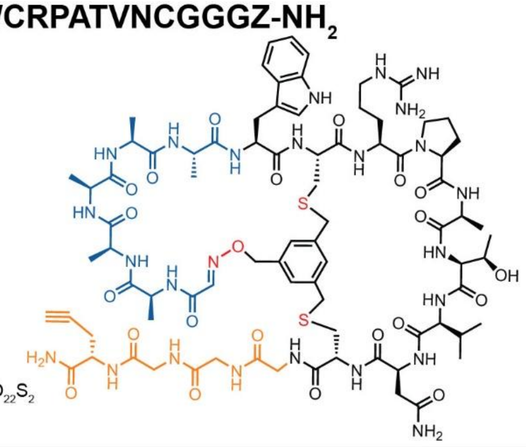

Chemical Formula: $C_{81}H_{115}N_{25}O_{22}S_2$
Exact Mass: 1853.81
Molecular Weight: 1855.08

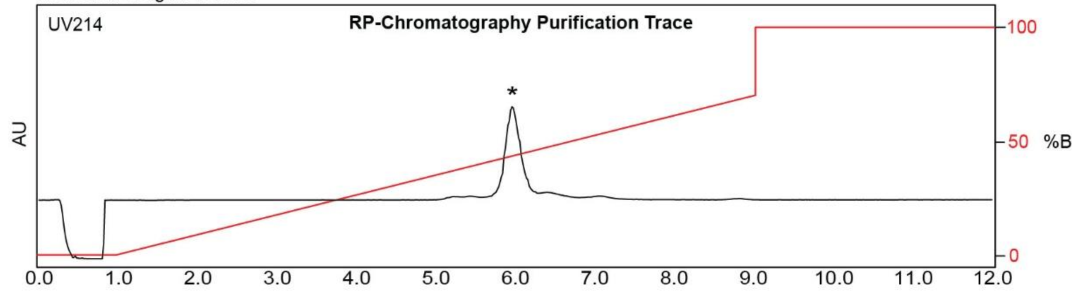

| Time (min) | Solvent B (%) |
|---|---|
| 0 | 0 |
| 1 | 0 |
| 9 | 70 |
| 9 | 100 |
| 12 | 100 |

Flow Rate: 18 mL / min
Teledyne ISCO, Inc RediSep Gold C18 Reversed-Phase column
(100 Å, 20-40 µm, 5.5 g)

Solvent A: $H_2O$ + 0.1% (v/v) TFA
Solvent B: MeCN + 0.1% (v/v) TFA

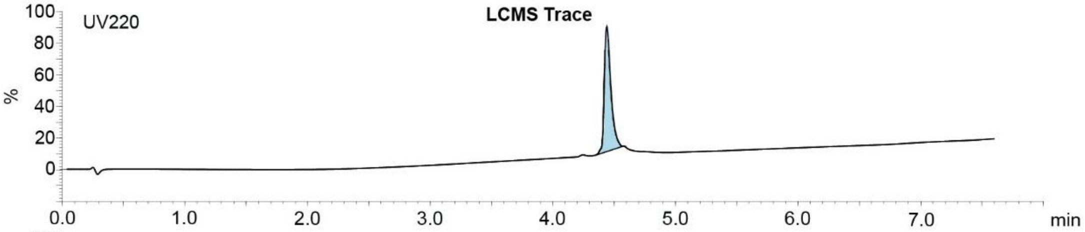

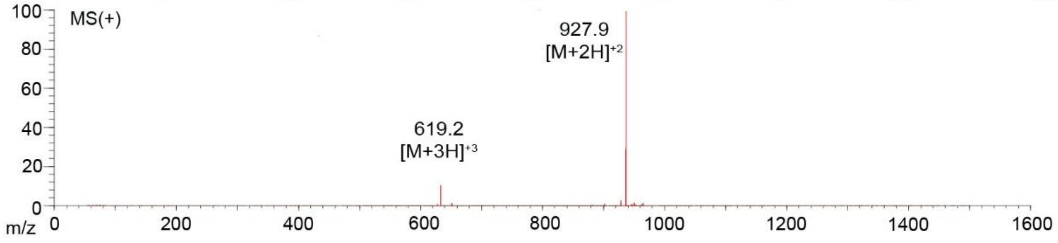

**Figure S118**. Data page for **44e**.

Expected: [M+1] = 1854.81, $[M+2]^{+2}$ = 927.905.

Observed: [M+1] = N/A, $[M+2]^{+2}$ = 927.9

Note: Detection limit is 1500

### TSL1-SAAAAAFCHYPTHVCGGGZ-$NH_2$

Scale: 20.6 mg
Yield: 5.4 mg (26%)
Purity: >95%

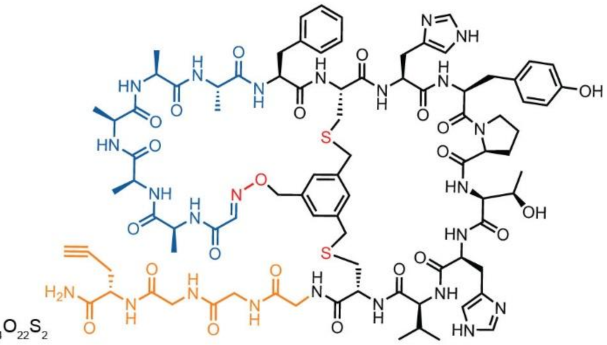

Chemical Formula: $C_{87}H_{114}N_{24}O_{22}S_2$
Exact Mass: 1910.80
Molecular Weight: 1912.13

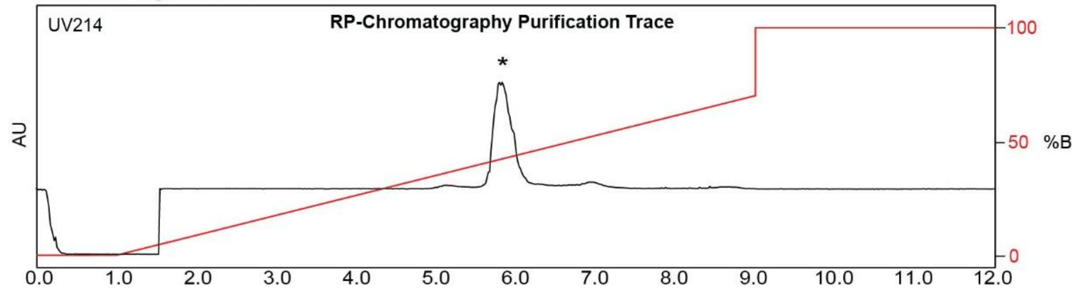

| Time (min) | Solvent B (%) |
|---|---|
| 0 | 0 |
| 1 | 0 |
| 9 | 70 |
| 9 | 100 |
| 12 | 100 |

Flow Rate: 18 mL / min
Teledyne ISCO, Inc RediSep Gold C18 Reversed-Phase column
(100 Å, 20-40 μm, 5.5 g)

Solvent A: $H_2O$ + 0.1% (v/v) TFA
Solvent B: MeCN + 0.1% (v/v) TFA

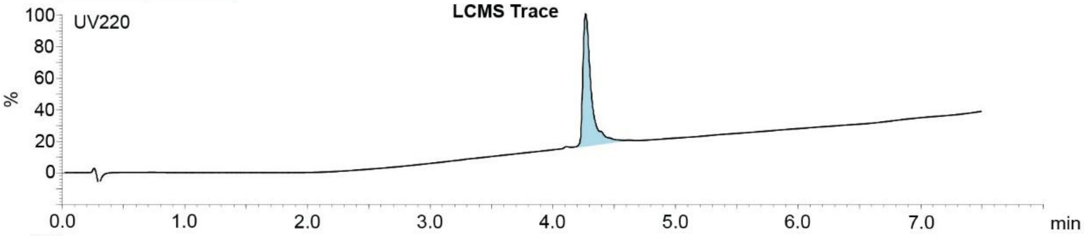

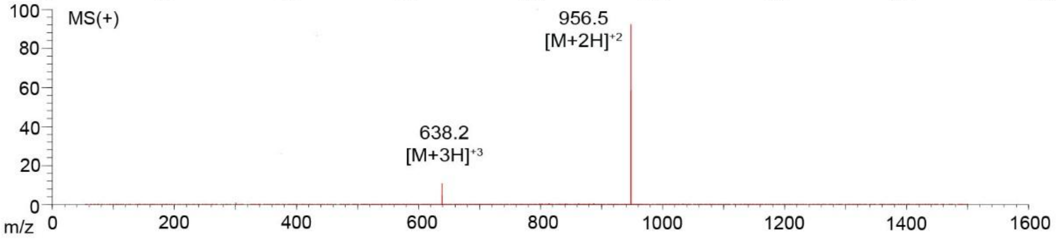

**Figure S119**. Data page for **45e**.

Expected: [M+1] = 1911.80, $[M+2]^{+2}$ = 956.40.

Observed: [M+1] = N/A, $[M+2]^{+2}$ = 956.5

Note: Detection limit is 1500

### TSL1-SAAAAAWCRPATVNC-$NH_2$

Scale: 21.3 mg
Yield: 1.6 mg (7.5%)
Purity: >95%

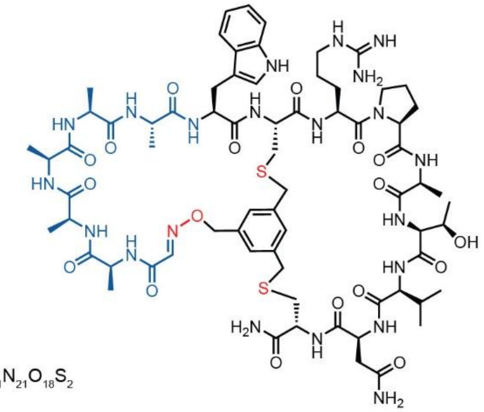

Chemical Formula: $C_{70}H_{101}N_{21}O_{18}S_2$
Exact Mass: 1587.71
Molecular Weight: 1588.83

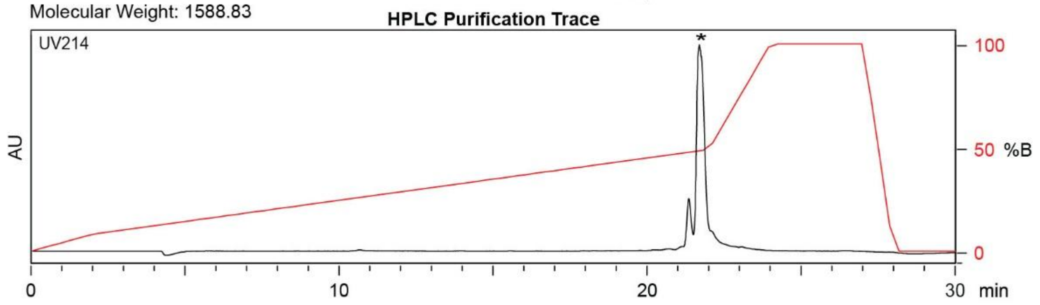

| Time (min) | Solvent B (%) |
|---|---|
| 0 | 2 |
| 2 | 10 |
| 22 | 50 |
| 24 | 100 |
| 27 | 100 |
| 28 | 2 |
| 30 | 2 |

Flow Rate: 13 mL / min
Phenomenex Kinetex EVO C18 Prep Column
(100 Å, 5 µm, 21.5 mm X 250 mm)

Solvent A: $H_2O$ + 0.1% (v/v) TFA
Solvent B: MeCN + 0.1% (v/v) TFA

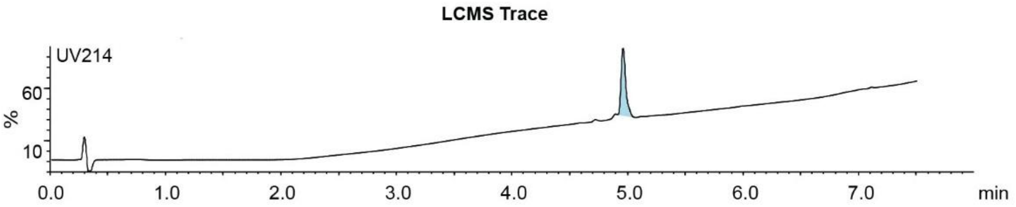

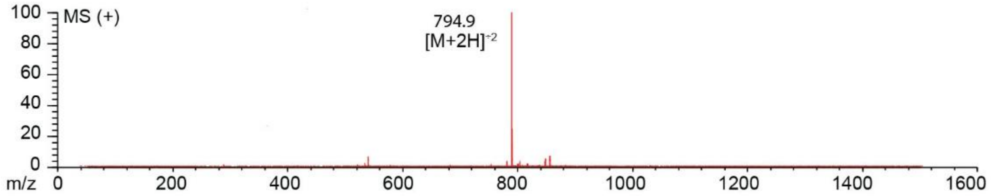

**Figure S120**. Data page for **46e**.

Expected: [M+1] = 1588.71, $[M+2]^{+2}$ = 794.855.

Observed: [M+1] = N/A, $[M+2]^{+2}$ = 794.9

Note: Detection limit is 1500

**TSL6-SECITAAGTC-$NH_2$**

Scale: 19.3 mg
Yield: 2.4 mg (12.4%)
Purity: > 95%

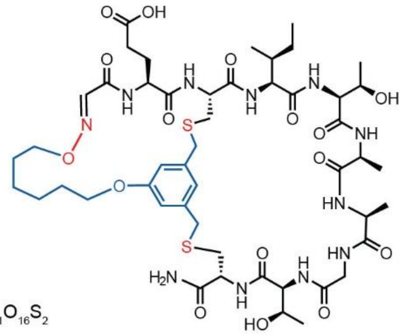

Chemical Formula: $C_{49}H_{75}N_{11}O_{16}S_2$
Exact Mass: 1137.48
Molecular Weight: 1138.32

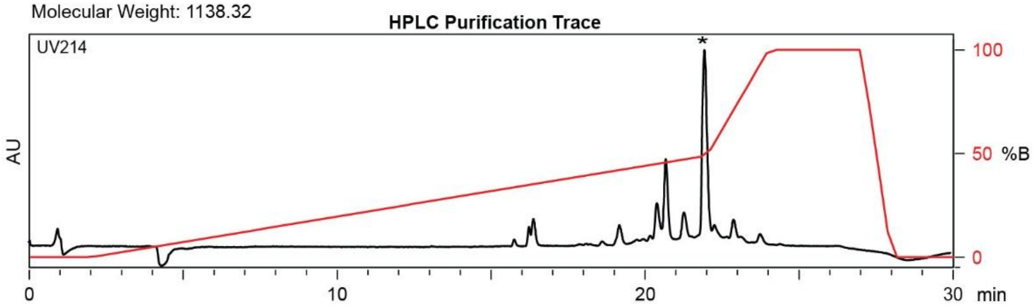

| Time (min) | Solvent B (%) |
|---|---|
| 0 | 2 |
| 2 | 2 |
| 22 | 50 |
| 24 | 100 |
| 27 | 100 |
| 28 | 2 |
| 30 | 2 |

Flow Rate: 13 mL / min
Phenomenex Kinetex EVO C18 Prep Column
(100 Å, 5 μm, 21.5 mm X 250 mm)

Solvent A: $H_2O$ + 0.1% (v/v) TFA
Solvent B: MeCN + 0.1% (v/v) TFA

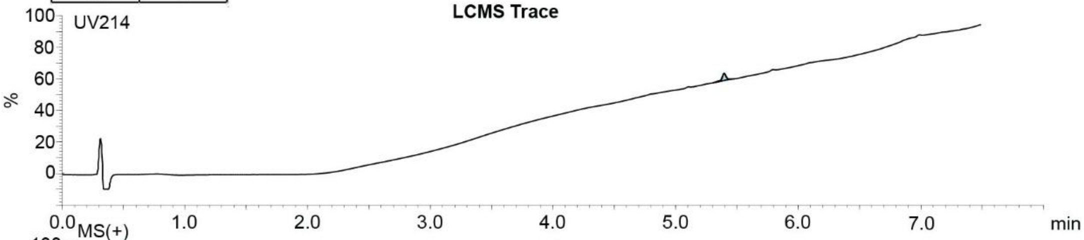

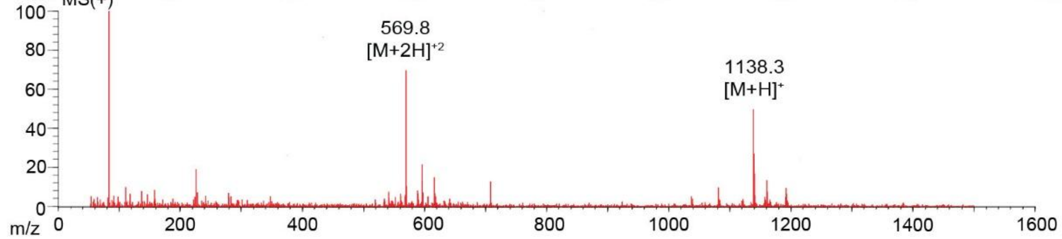

**Figure S121**. Data page for **47c**.

Expected: [M+1] = 1138.48, $[M+2]^{+2}$ = 569.74.

Observed: [M+1] = 1138.3, $[M+2]^{+2}$ = 569.8

### TSL6-SPCTQGVKIC-$NH_2$

Scale: 10.1 mg
Yield: 1.1 mg (10.9%)
Purity: > 95%

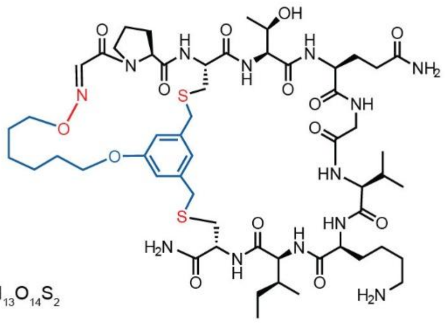

Chemical Formula: $C_{55}H_{87}N_{13}O_{14}S_2$
Exact Mass: 1217.59
Molecular Weight: 1218.50

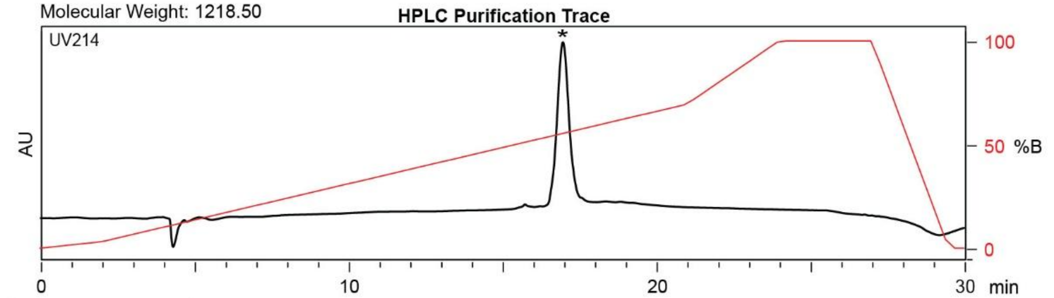

| Time (min) | Solvent B (%) |
|---|---|
| 0 | 2 |
| 2 | 2 |
| 22 | 70 |
| 24 | 100 |
| 27 | 100 |
| 28 | 2 |
| 30 | 2 |

Flow Rate: 13 mL / min
Phenomenex Kinetex EVO C18 Prep Column
(100 Å, 5 µm, 21.5 mm X 250 mm)

Solvent A: $H_2O$ + 0.1% (v/v) TFA
Solvent B: MeCN + 0.1% (v/v) TFA

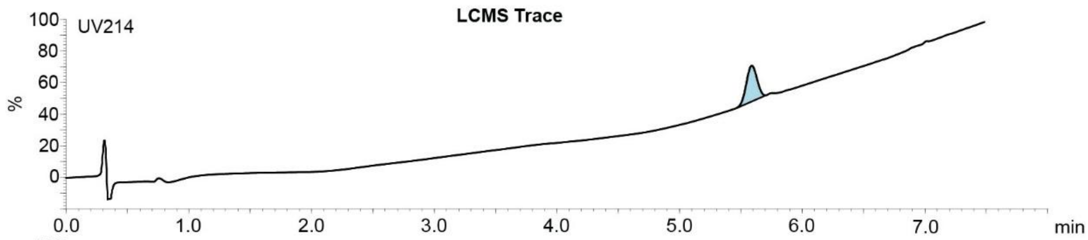

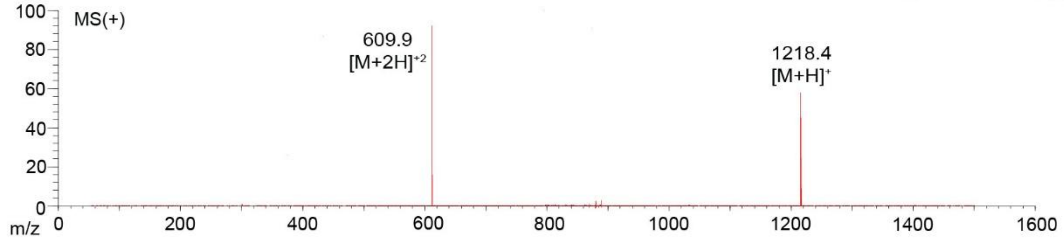

**Figure S122**. Data page for **48c**.

Expected: [M+1] = 1218.59, $[M+2]^{+2}$ = 609.795.

Observed: [M+1] = 1218.4, $[M+2]^{+2}$ = 609.9

### TSL6-STCPVRARNC-$NH_2$

Scale: 10.0 mg
Yield: 2.7 mg (27%)
Purity: >95%

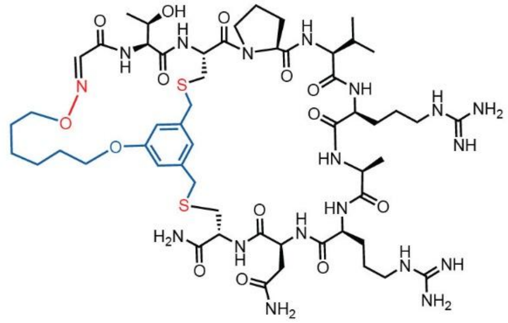

Chemical Formula: $C_{55}H_{88}N_{18}O_{14}S_2$
Exact Mass: 1288.62
Molecular Weight: 1289.53

**HPLC Purification Trace**

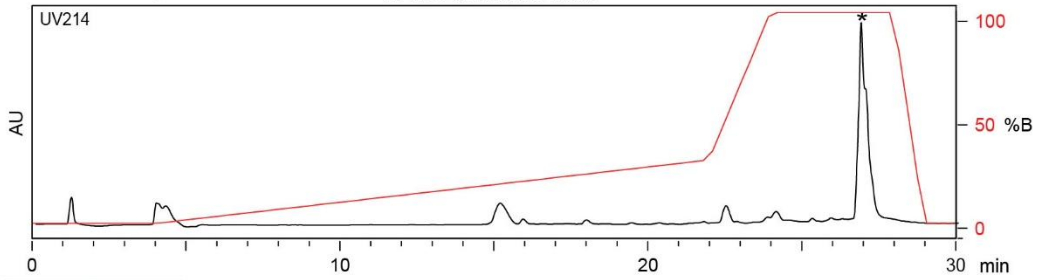

| Time (min) | Solvent B (%) |
|---|---|
| 0 | 0 |
| 4 | 0 |
| 22 | 30 |
| 24 | 100 |
| 27 | 100 |
| 29.5 | 0 |
| 30 | 0 |

Flow Rate: 13 mL / min
Phenomenex Kinetex EVO C18 Prep Column
(100 Å, 5 µm, 21.5 mm X 250 mm)

Solvent A: $H_2O$ + 0.1% (v/v) TFA
Solvent B: MeCN + 0.1% (v/v) TFA

**LCMS Trace**

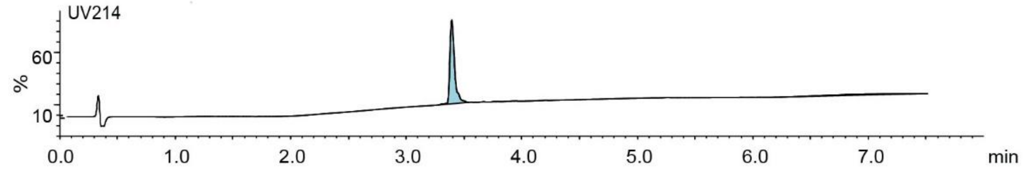

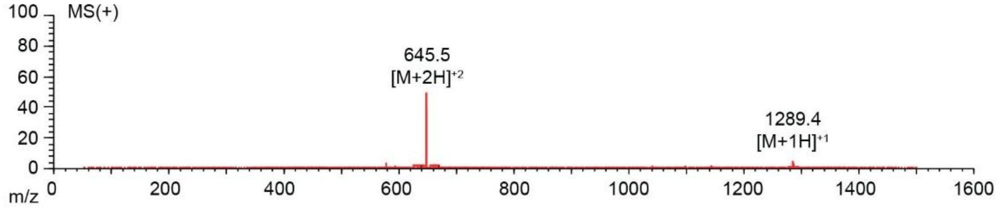

**Figure S123**. Data page for **49c**.

Expected: [M+1] = 1289.62, $[M+2]^{+2}$ = 645.31.

Observed: [M+1] = 1289.4, $[M+2]^{+2}$ = 645.5

### TSL6-SECEEPSIKC-$NH_2$

Scale: 10.0 mg
Yield: 5.0 mg (50%)
Purity: >95%

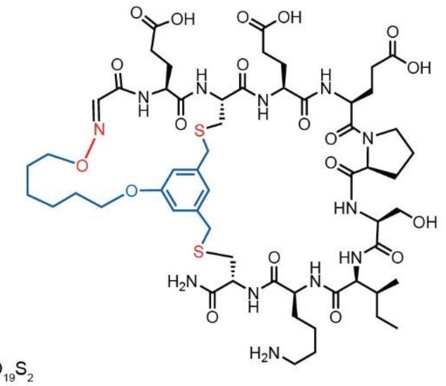

Chemical Formula: $C_{57}H_{86}N_{12}O_{19}S_2$
Exact Mass: 1306.56
Molecular Weight: 1307.49

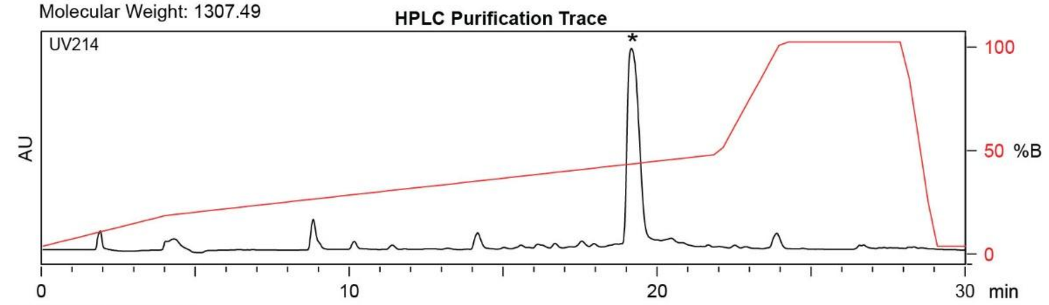

| Time (min) | Solvent B (%) |
|---|---|
| 0 | 0 |
| 4 | 15 |
| 22 | 45 |
| 24 | 100 |
| 28 | 100 |
| 29 | 0 |
| 30 | 0 |

Flow Rate: 13 mL / min
Phenomenex Kinetex EVO C18 Prep Column
(100 Å, 5 µm, 21.5 mm X 250 mm)

Solvent A: $H_2O$ + 0.1% (v/v) TFA
Solvent B: MeCN + 0.1% (v/v) TFA

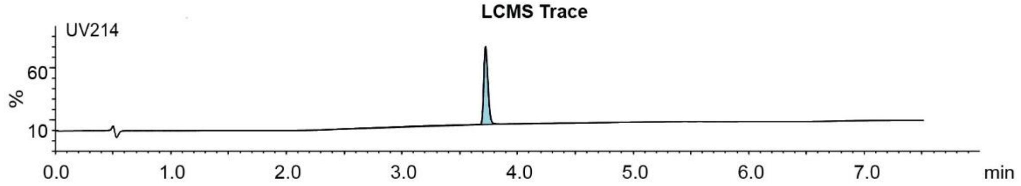

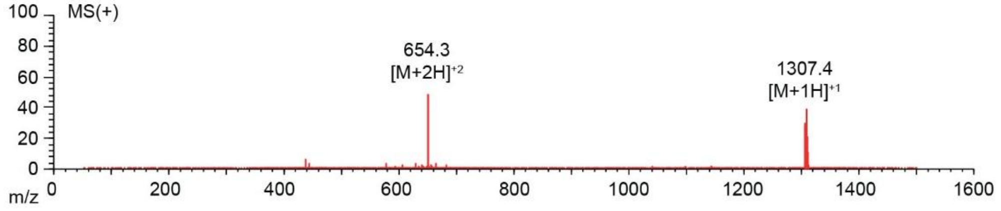

**Figure S124**. Data page for **50c**.

Expected: [M+1] = 1307.56, $[M+2]^{+2}$ = 654.28.

Observed: [M+1] = 1307.4, $[M+2]^{+2}$ = 654.3

### TSL6-SSCDITREHC-$NH_2$

Scale: 10.0 mg
Yield: 2.2 mg (22%)
Purity: >95%

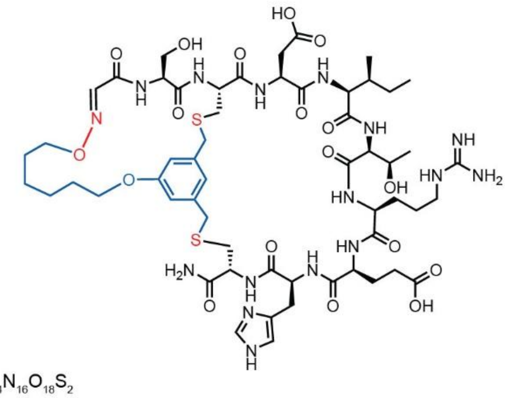

Chemical Formula: $C_{56}H_{84}N_{16}O_{18}S_2$
Exact Mass: 1332.56
Molecular Weight: 1333.49

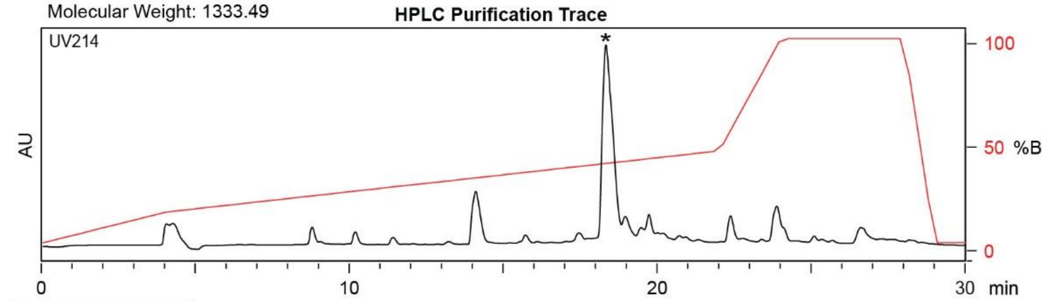

| Time (min) | Solvent B (%) |
|---|---|
| 0 | 0 |
| 4 | 15 |
| 22 | 45 |
| 24 | 100 |
| 28 | 100 |
| 29 | 0 |
| 30 | 0 |

Flow Rate: 13 mL / min
Phenomenex Kinetex EVO C18 Prep Column
(100 Å, 5 µm, 21.5 mm X 250 mm)

Solvent A: $H_2O$ + 0.1% (v/v) TFA
Solvent B: MeCN + 0.1% (v/v) TFA

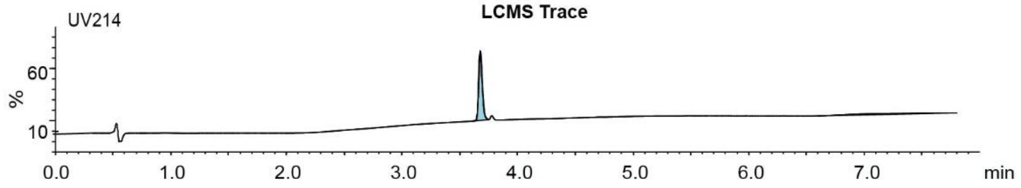

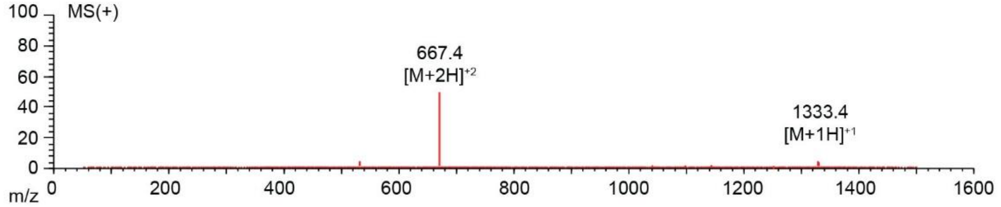

**Figure S125**. Data page for **51c**.

Expected: [M+1] = 1333.56, $[M+2]^{+2}$ = 667.28.

Observed: [M+1] = 1333.4, $[M+2]^{+2}$ = 667.4

**TSL6-SPCPHQVLDC-$NH_2$**

Scale: 18.3 mg
Yield: 2.4 mg (13.1%)
Purity: >95%

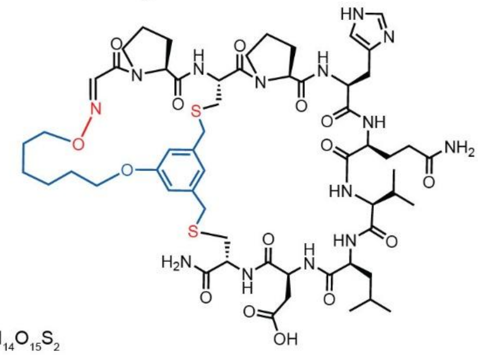

Chemical Formula: $C_{58}H_{84}N_{14}O_{15}S_2$
Exact Mass: 1280.57
Molecular Weight: 1281.51

**HPLC Purification Trace**

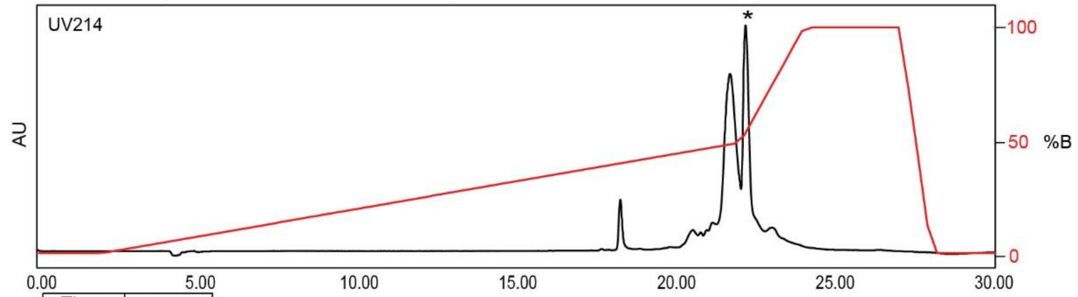

| Time (min) | Solvent B (%) |
|---|---|
| 0 | 2 |
| 2 | 2 |
| 22 | 50 |
| 24 | 100 |
| 27 | 100 |
| 28 | 2 |
| 30 | 2 |

Flow Rate: 13 mL / min
Phenomenex Kinetex EVO C18 Prep Column
(100 Å, 5 µm, 21.5 mm X 250mm)

Solvent A: $H_2O$ + 0.1% (v/v) TFA
Solvent B: MeCN + 0.1% (v/v) TFA

**LCMS Trace**

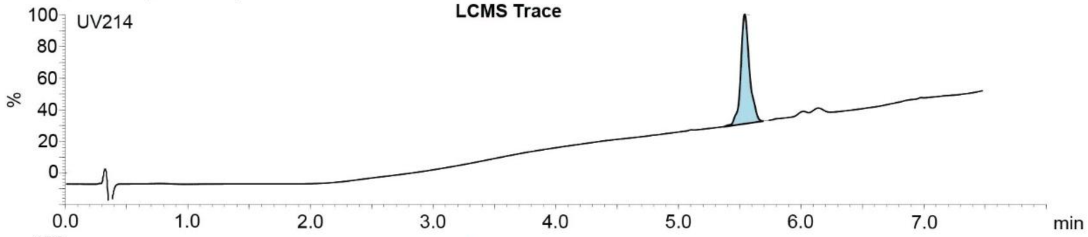

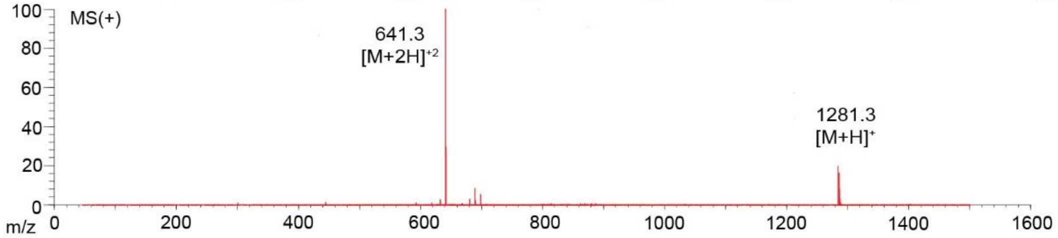

**Figure S126**. Data page for **52c**.

Expected: [M+1] = 1281.57, $[M+2]^{+2}$ = 641.285.

Observed: [M+1] = 1281.3, $[M+2]^{+2}$ = 641.3

### TSL6-SQCVVHMEEC-$NH_2$

Scale: 10.0 mg
Yield: 0.4 mg (4%)
Purity: >90%

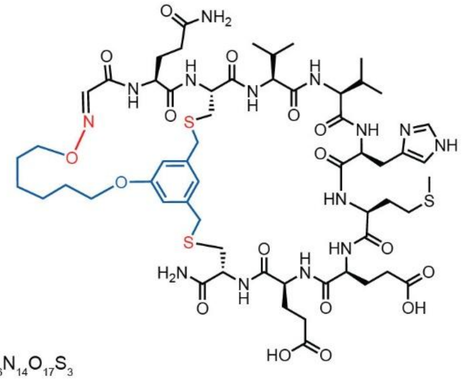

Chemical Formula: $C_{58}H_{86}N_{14}O_{17}S_3$
Exact Mass: 1346.55
Molecular Weight: 1347.58

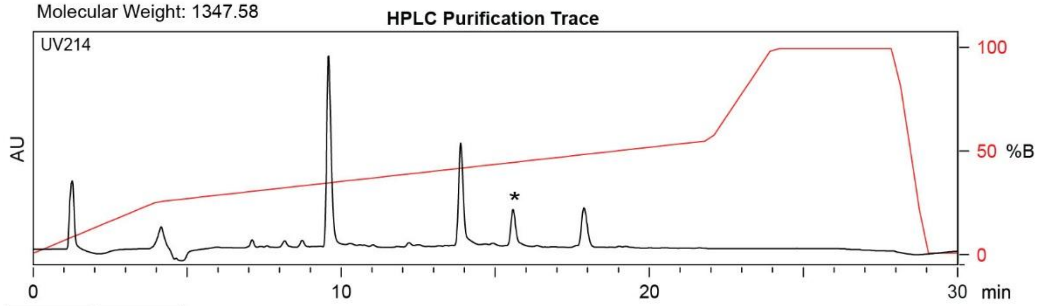

| Time (min) | Solvent B (%) |
|---|---|
| 0 | 0 |
| 4 | 25 |
| 22 | 55 |
| 24 | 100 |
| 28 | 100 |
| 29 | 0 |
| 30 | 0 |

Flow Rate: 13 mL / min
Phenomenex Kinetex EVO C18 Prep Column
(100 Å, 5 µm, 21.5 mm X 250 mm)

Solvent A: $H_2O$ + 0.1% (v/v) TFA
Solvent B: MeCN + 0.1% (v/v) TFA

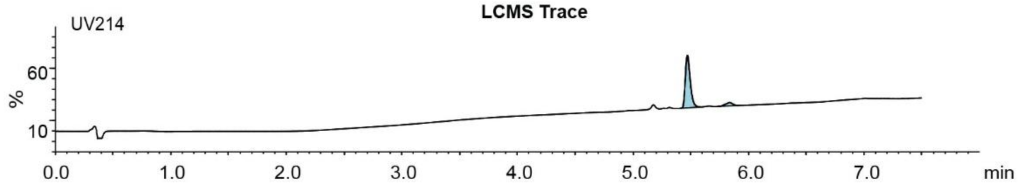

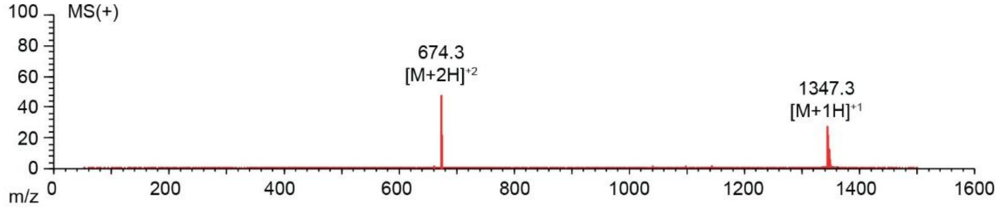

**Figure S127**. Data page for **53c**.

Expected: [M+1] = 1347.55, $[M+2]^{+2}$ = 674.275.

Observed: [M+1] = 1347.3, $[M+2]^{+2}$ = 674.3

### TSL6-SVQDHAHAC-$NH_2$

Scale: 10.0 mg
Yield: 1.1 mg (11%)
Purity: >95%

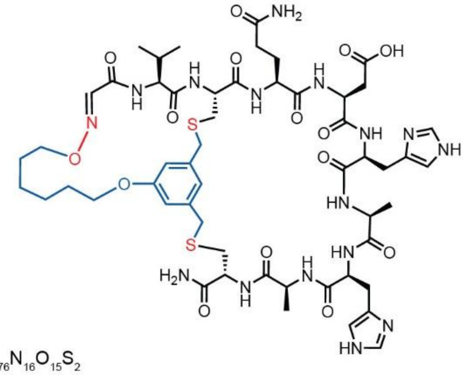

Chemical Formula: $C_{54}H_{76}N_{16}O_{15}S_2$
Exact Mass: 1252.51
Molecular Weight: 1253.41

**HPLC Purification Trace**

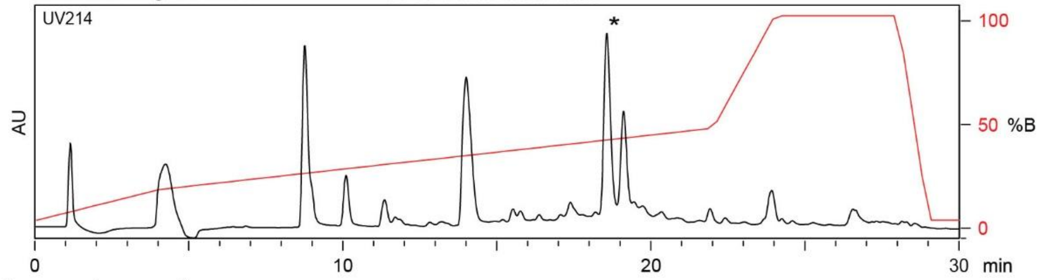

| Time (min) | Solvent B (%) |
|---|---|
| 0 | 0 |
| 4 | 15 |
| 22 | 45 |
| 24 | 100 |
| 28 | 100 |
| 29 | 0 |
| 30 | 0 |

Flow Rate: 13 mL / min
Phenomenex Kinetex EVO C18 Prep Column
(100 Å, 5 µm, 21.5 mm X 250 mm)

Solvent A: $H_2O$ + 0.1% (v/v) TFA
Solvent B: MeCN + 0.1% (v/v) TFA

**LCMS Trace**

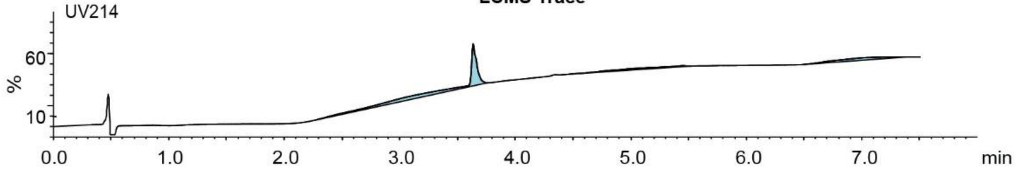

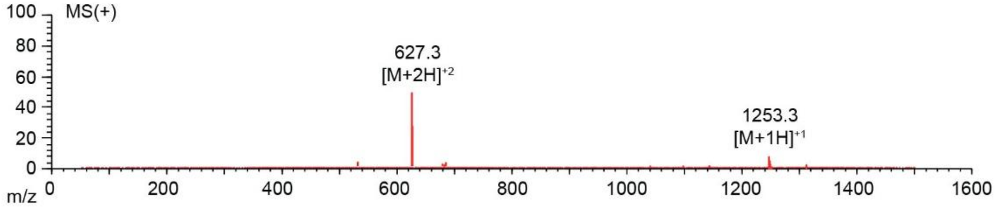

**Figure S128**. Data page for **54c**.

Expected: [M+1] = 1253.51, $[M+2]^{+2}$ = 627.255.

Observed: [M+1] = 1253.3, $[M+2]^{+2}$ = 627.3

**TSL6-STCTGIELDC-$NH_2$**

Scale: 11.0 mg
Yield: 1.0 mg (11%)
Purity: >95%

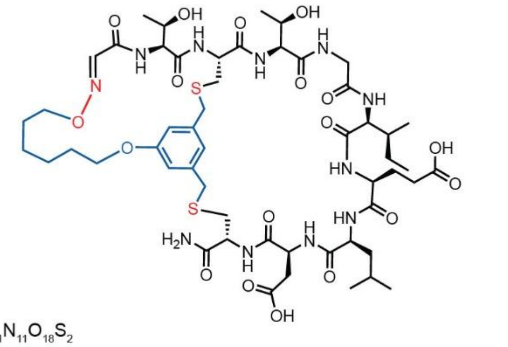

Chemical Formula: $C_{53}H_{81}N_{11}O_{18}S_2$
Exact Mass: 1223.52
Molecular Weight: 1224.41

HPLC Purification Trace

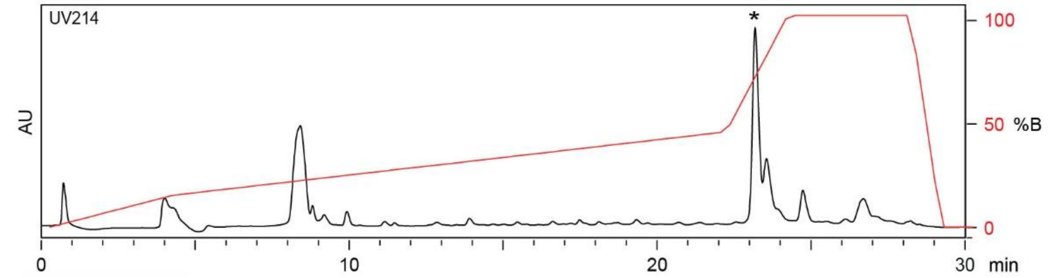

| Time (min) | Solvent B (%) |
|---|---|
| 0 | 0 |
| 4 | 15 |
| 22 | 45 |
| 24 | 100 |
| 27 | 100 |
| 29.5 | 0 |
| 30 | 0 |

Flow Rate: 13 mL / min
Phenomenex Kinetex EVO C18 Prep Column
(100 Å, 5 µm, 21.5 mm X 250 mm)

Solvent A: $H_2O$ + 0.1% (v/v) TFA
Solvent B: MeCN + 0.1% (v/v) TFA

LCMS Trace

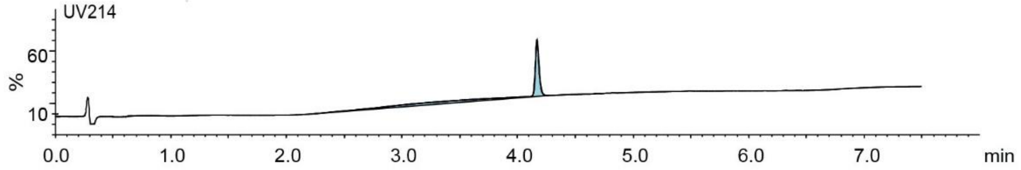

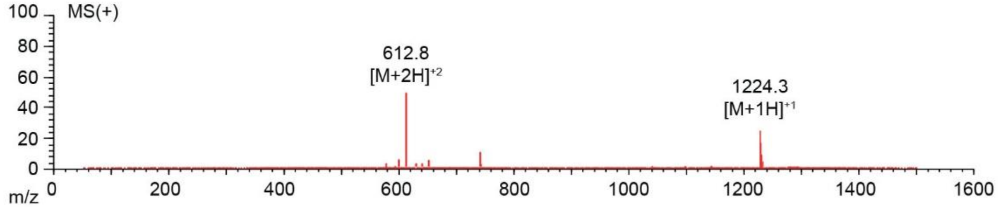

**Figure S129**. Data page for **55c**.

Expected: [M+1] = 1224.52, $[M+2]^{+2}$ = 612.76.

Observed: [M+1] = 1224.3, $[M+2]^{+2}$ = 612.8

### TSL6-SQCGKSYQEC-$NH_2$

Scale: 11.0 mg
Yield: 1.2 mg (10.9%)
Purity: > 90%

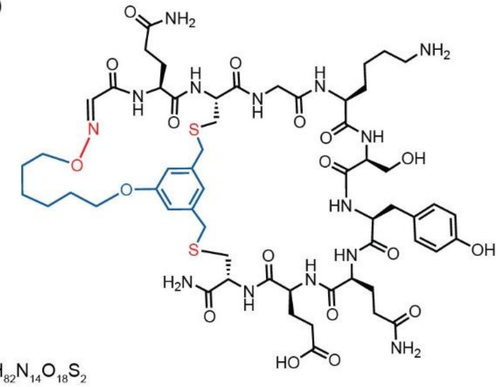

Chemical Formula: $C_{57}H_{82}N_{14}O_{18}S_2$
Exact Mass: 1314.54
Molecular Weight: 1315.46

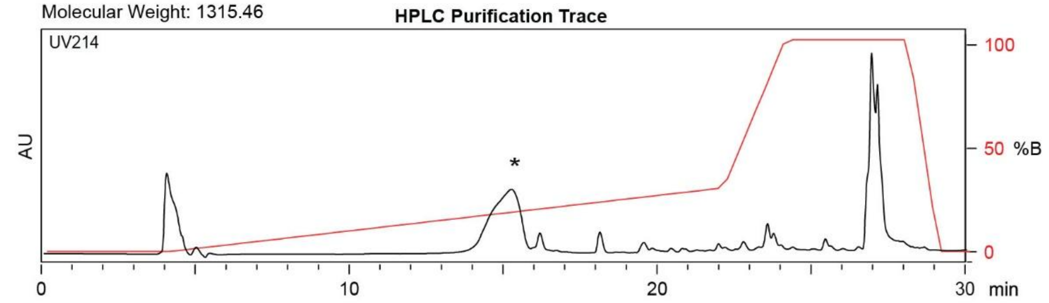

| Time (min) | Solvent B (%) |
|---|---|
| 0 | 0 |
| 4 | 0 |
| 22 | 30 |
| 24 | 100 |
| 27 | 100 |
| 29.5 | 0 |
| 30 | 0 |

Flow Rate: 13 mL / min
Phenomenex Kinetex EVO C18 Prep Column
(100 Å, 5 µm, 21.5 mm X 250 mm)

Solvent A: $H_2O$ + 0.1% (v/v) TFA
Solvent B: MeCN + 0.1% (v/v) TFA

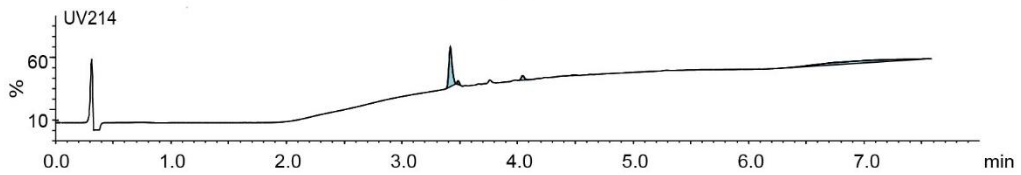

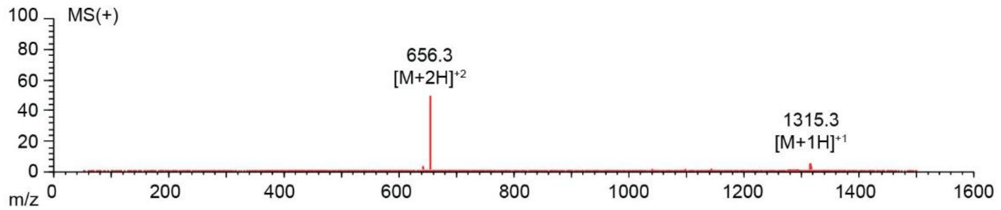

**Figure S130**. Data page for **56c**.

Expected: [M+1] = 1315.54, $[M+2]^{+2}$ = 658.27.

Observed: [M+1] = 1315.3, $[M+2]^{+2}$ = 656.3

**TSL1-SAAAAAWCRPATVNCBBX-$NH_2$**

Scale: 20.4 mg
Yield: 1.6 mg (7.8%)
Purity: > 90%

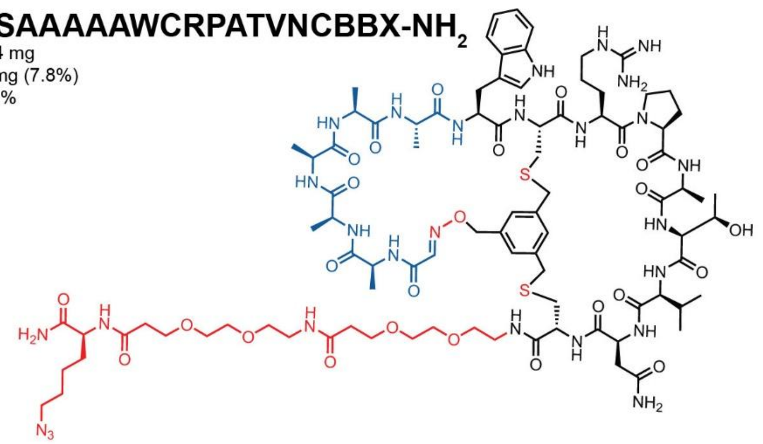

Chemical Formula: $C_{90}H_{137}N_{27}O_{25}S_2$
Exact Mass: 2059.97
Molecular Weight: 2061.37

**HPLC Purification Trace**

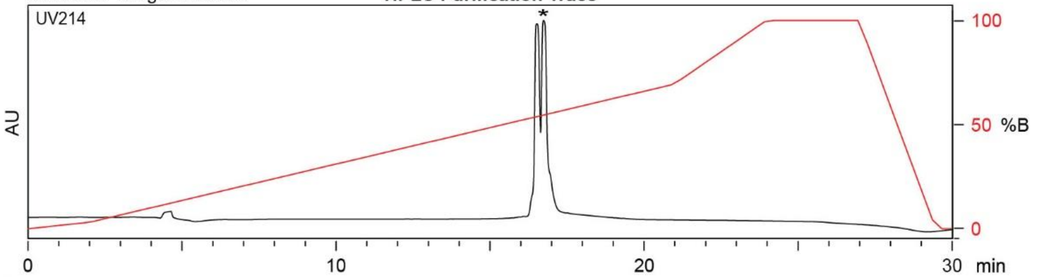

| Time (min) | Solvent B (%) |
|---|---|
| 0 | 2 |
| 2 | 5 |
| 21 | 70 |
| 24 | 100 |
| 27 | 100 |
| 29.5 | 2 |
| 30 | 2 |

Flow Rate: 13 mL / min
Phenomenex Kinetex EVO C18 Prep Column
(100 Å, 5 µm, 21.5 mm X 250 mm)

Solvent A: $H_2O$ + 0.1% (v/v) TFA
Solvent B: MeCN + 0.1% (v/v) TFA

**LCMS Trace**

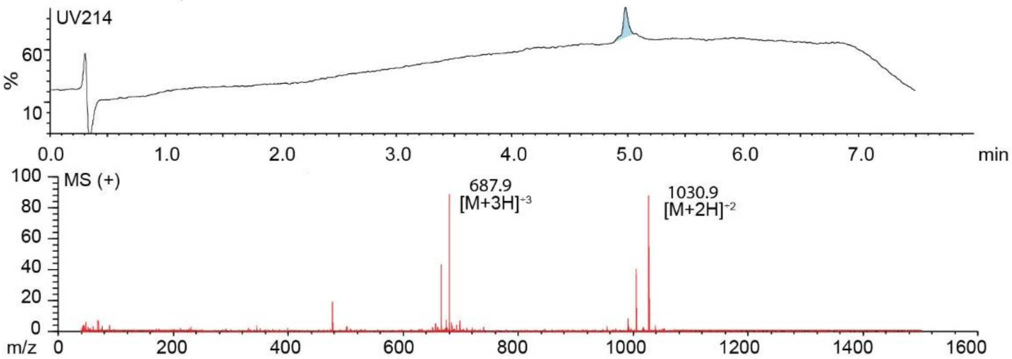

**Figure S131**. Data page for **57e**.

Expected: [M+1] = 2060.97, $[M+2]^{+2}$ = 1030.985.

Observed: [M+1] = N/A, $[M+2]^{+2}$ = 1030.9

Note: Detection limit is 1500

### TSL6-SFCHYPTHVGGGX-$NH_2$

Scale: 21.2 mg
Yield: 3.2 mg (15.1%)
Purity: > 95%

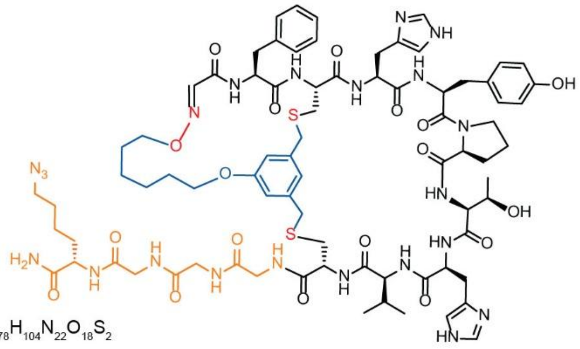

Chemical Formula: $C_{78}H_{104}N_{22}O_{18}S_2$
Exact Mass: 1700.73
Molecular Weight: 1701.95

**HPLC Purification Trace**

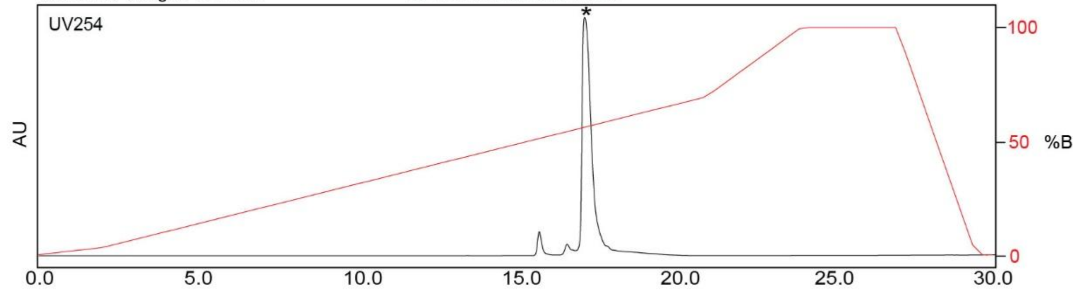

| Time (min) | Solvent B (%) |
|---|---|
| 0 | 2 |
| 2 | 5 |
| 21 | 70 |
| 24 | 100 |
| 27 | 100 |
| 30 | 2 |

Flow Rate: 13 mL / min
Phenomenex Kinetex EVO C18 Prep Column
(100 Å, 5 μm, 21.5 mm X 250 mm)

Solvent A: $H_2O$ + 0.1% (v/v) TFA
Solvent B: MeCN + 0.1% (v/v) TFA

**LCMS Trace**

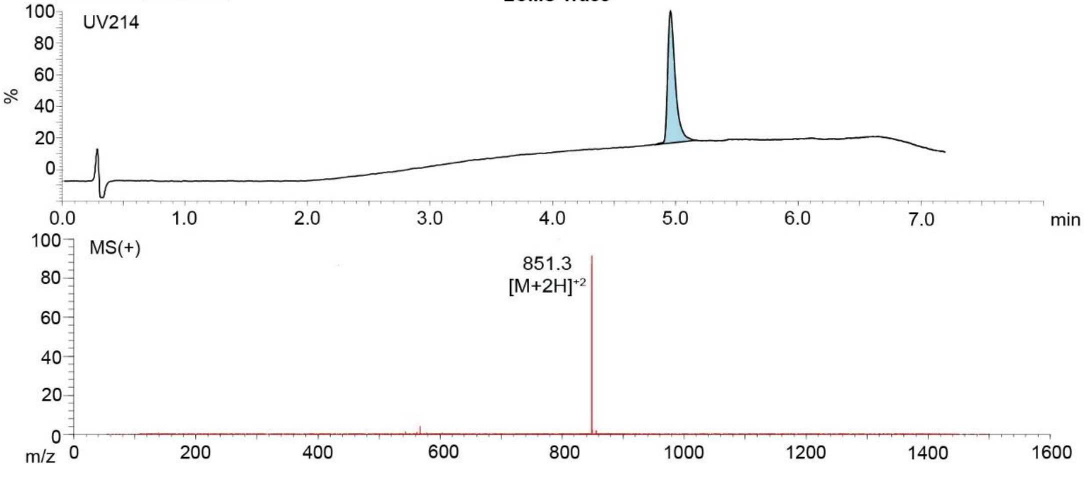

**Figure S132**. Data page for **58c**.

Expected: [M+1] = 1701.73, $[M+2]^{\div 2}$ = 851.365.

Observed: [M+1] = N/A, $[M+2]^{\div 2}$ = 851.3

Note: Detection limit is 1500
